# A foundational atlas of autism protein interactions reveals molecular convergence

**DOI:** 10.1101/2023.12.03.569805

**Authors:** Belinda Wang, Rasika Vartak, Yefim Zaltsman, Zun Zar Chi Naing, Kelsey M. Hennick, Benjamin J. Polacco, Ali Bashir, Manon Eckhardt, Mehdi Bouhaddou, Jiewei Xu, Nawei Sun, Micaela C. Lasser, Yuan Zhou, Justin McKetney, Keelan Z. Guiley, Una Chan, Julia A. Kaye, Nishant Chadha, Merve Cakir, Martin Gordon, Prachi Khare, Sam Drake, Vanessa Drury, David F. Burke, Silvano Gonzalez, Sahar Alkhairy, Reuben Thomas, Stephanie Lam, Montana Morris, Ethel Bader, Meghan Seyler, Tierney Baum, Rebecca Krasnoff, Sheng Wang, Presley Pham, Juan Arbalaez, Dexter Pratt, Shivali Chag, Nadir Mahmood, Thomas Rolland, Thomas Bourgeron, Steven Finkbeiner, Danielle L. Swaney, Sourav Bandyopadhay, Trey Ideker, Pedro Beltrao, Helen Rankin Willsey, Kirsten Obernier, Tomasz J. Nowakowski, Ruth Hüttenhain, Matthew W. State, A. Jeremy Willsey, Nevan J. Krogan

## Abstract

Translating high-confidence (hc) autism spectrum disorder (ASD) genes into viable treatment targets remains elusive. We constructed a foundational protein-protein interaction (PPI) network in HEK293T cells involving 100 hcASD risk genes, revealing over 1,800 PPIs (87% novel). Interactors, expressed in the human brain and enriched for ASD but not schizophrenia genetic risk, converged on protein complexes involved in neurogenesis, tubulin biology, transcriptional regulation, and chromatin modification. A PPI map of 54 patient-derived missense variants identified differential physical interactions, and we leveraged AlphaFold-Multimer predictions to prioritize direct PPIs and specific variants for interrogation in *Xenopus tropicalis* and human forebrain organoids. A mutation in the transcription factor FOXP1 led to reconfiguration of DNA binding sites and altered development of deep cortical layer neurons in forebrain organoids. This work offers new insights into molecular mechanisms underlying ASD and describes a powerful platform to develop and test therapeutic strategies for many genetically-defined conditions.

## Introduction

Autism spectrum disorder (ASD) is a highly heritable neurodevelopmental syndrome characterized by diverse etiology, marked inter-individual variability in symptom presentation, and a wide range of associated features apart from the defining impairments in social communication and highly restricted interests and/or repetitive behaviors^1,2^. Over the past decade and a half, whole-exome and whole-genome based approaches have resulted in the systematic and highly reliable identification of large effect risk genes based on the study of rare, often *de novo*, variations. The most recent large scale WES studies have identified more than 200 genes robustly associated with ASD^33–7^. However, translating the increasingly large lists of high confidence ASD (hcASD) genes generated by these approaches into a comprehensive understanding of the biological mechanisms underlying ASD - and potential therapeutic targets - has been hampered by extensive pleiotropy and locus heterogeneity coupled with the complexity of human brain development and a relative lack of understanding of the molecular interactions among the proteins these risk genes encode^7–10^. Adopting a convergent framework, rooted in the premise that identifying shared characteristics among hcASD genes will pinpoint core mechanisms underlying pathobiology, we have extended the molecular characterization of rare large-effect risk genes to address the proteins they encode, identify their direct interactors, and evaluate the consequences of syndrome associated mutations on this ASD proteomics landscape.

As the list of hcASD risk genes has expanded, strategies for determining the convergence of gene expression patterns or biological functions across various ASD risk genes have repeatedly highlighted neurogenesis in the human mid-gestational prefrontal cortex as an important nexus of pathobiology^7,11–17^. At the same time, gene ontology enrichment analyses have implicated broad functional categories of these genes, such as chromatin modification, transcriptional regulation,, cell signaling, and synaptic function^6,18–21^. However, gene ontology-based approaches may be incomplete as they are limited by *a priori* knowledge. For example, recent work by our group has suggested that many chromatin modifiers may also regulate tubulin and that disruption of microtubule dynamics may be another point of convergent biology underlying ASD^22^. Nonetheless, comprehensive molecular and functional data for ASD risk genes remain scarce^7^.

To employ convergent approaches for identifying core biological features of ASD, it is essential to generate large-scale biological datasets, particularly those derived from direct experimental evidence. Systematically mapping the physical interaction networks of ASD risk genes and their likely damaging missense variants provides opportunities to identify how these genes functionally converge at a molecular level and allows for identifying shifts in interactions or structural changes in key protein complexes in the presence of patient-derived variants. Recent studies that have profiled protein-protein interaction (PPI) networks of ASD genes have highlighted evidence of molecular convergence^23–25^. However, these studies focused on a narrow selection of genes, with varying degrees of association to ASD, and did not explore at scale how specific molecular pathways may be affected by missense variants derived from individuals with ASD.

In this study, we have generated the largest PPI networks of wildtype (WT) and mutant (mut) hcASD risk genes to date, using affinity purification-mass spectrometry (AP-MS) in HEK293T cells, a widely used *in vitro* system for proteomics studies. We have successfully applied similar approaches to cancer^26–29^, heart disease^30–33^, neurodegeneration^34^ and infectious diseases^35–45^. Our ASD networks here consist of 100 hcASD risk genes (ASD-PPI network) and 54 patient-derived variants^6^ affecting 30 hcASD genes (ASD_mut_-PPI network) (**Figure 1A**). This foundational ASD-PPI network, characterized in a non-neuronal cell line, contains over 1,000 ASD-PPI interactors and more than 1,800 interactions which we show to be highly relevant to the human brain and to ASD. Our networks highlight molecular convergence of ASD risk genes and their interactors and reveal interactions that change in the presence of patient-derived missense variants, thus highlighting a range of potentially targetable pathways. Using AlphaFold (AF) pairwise predictions to identify direct PPI interactions, we prioritize a subset for functional interrogation in *Xenopus tropicalis*, human induced pluripotent stem cells (iPSCs), and forebrain organoids. We identified DCAF7 as a central hub interacting with multiple hcASD proteins, with disruption of *DCAF7* leading to impaired neurogenesis and changes in telencephalon size in *Xenopus*. We utilize human iPSC-derived forebrain organoids to demonstrate that a missense mutation in FOXP1 (R513H) disrupts binding to FOXP4 as predicted by our HEK293T cell-derived data as well as AF modeling, disrupts neurogenesis, leads to re-wiring of transcription factor binding sites, and alters the differentiation trajectories of deep layer cortical glutamatergic neurons. Overall, this work and the resulting PPI resource offer valuable novel insights into the molecular mechanisms of ASD genetic risk and constitute a foundational platform for future therapeutics development.

**Figure 1.**
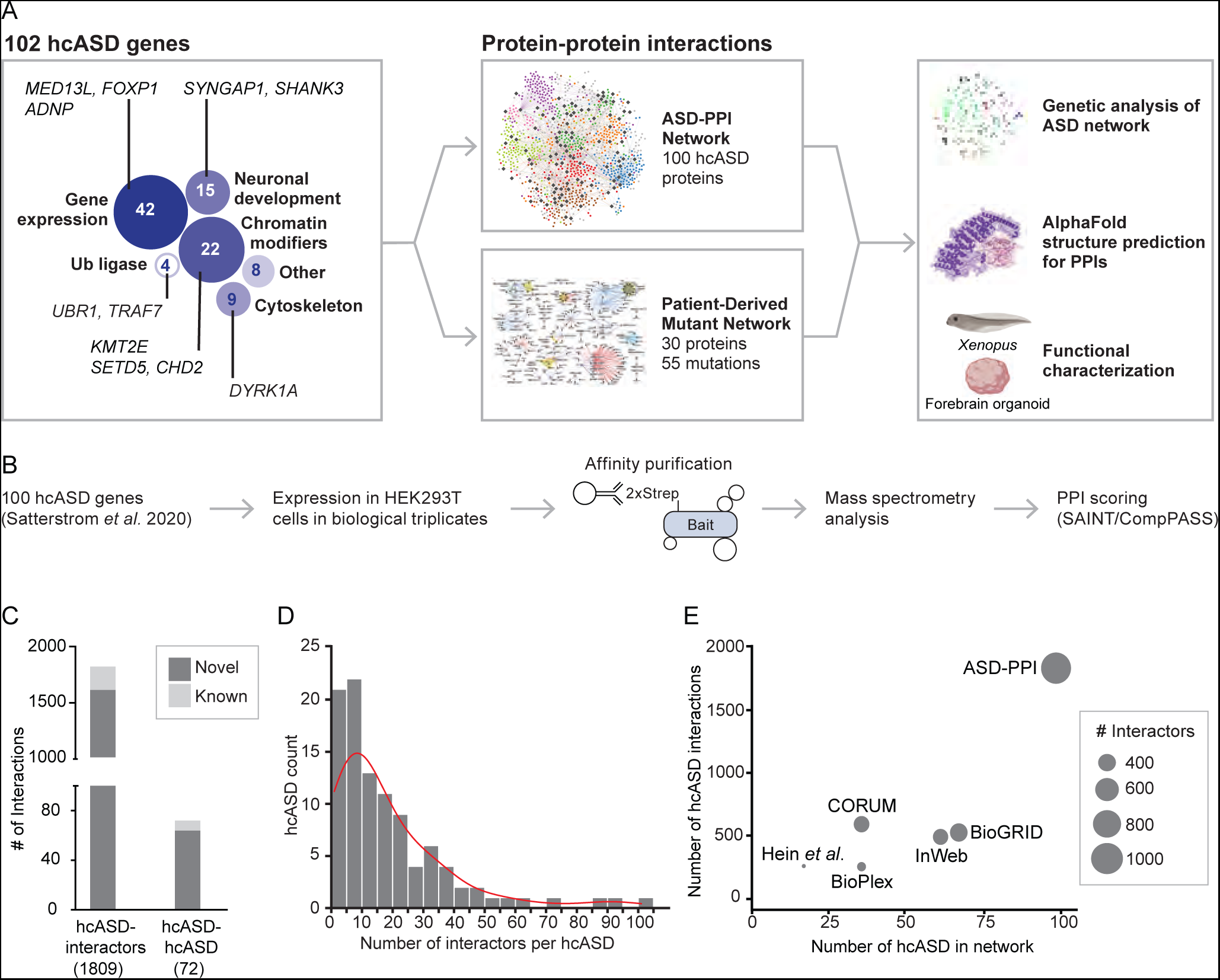
Protein interaction mapping of hcASD reveals novel interactors. (A) Overview of the ASD-PPI and patient derived mutant networks. (B) Workflow to generate PPI data for 100 hcASD^6^ via AP-MS in HEK293T cells. (C) ASD-PPI contains a total of 1,881 interactions of which 235 are known and 1,646 are novel. (D) The distribution of the number of interactors per hcASD bait. The median number of interactors for the baits is 11. (E) Comparison of the number of hcASD proteins reported as a bait and the number of hcASD-associated interactions in various large-scale PPI datasets, including BioGRID, BioPlex, CORUM, InWeb, and Hein *et al*. 2015^48–52^. Point size represents the number of unique hcASD interactors for each dataset.

## Results

### Protein-protein interaction mapping of hcASD risk genes reveals novel interactors

A recent large scale exome sequencing study identified 102 high-confidence ASD risk genes (hcASD102) with a false discovery rate (FDR) of 0.1 or less^6^. We generated a protein-protein interaction (PPI) network for 100 wildtype (WT) hcASD102 proteins (hereafter referred to as ‘hcASD’), selecting the highest brain expressed isoform where possible (**Table S1**). To generate this network, Strep-tagged proteins were individually overexpressed (as “bait” proteins) in HEK293T cells in biological triplicates and “interactor” proteins (preys) were identified by affinity purification followed by mass spectrometry (AP-MS) 48 hours after transfection (**Figure 1B**). We used two scoring algorithms to identify high-confidence interactors: SAINTexpress^46^ and CompPASS^47,48^, selecting score cutoffs based on recovery of “gold-standard” interactions^48–51^ (**Figures S1A-C**, **Table S1**, see Methods). This WT PPI network, termed ‘ASD-PPI’, consists of 1,074 unique interactors connecting 100 hcASD proteins (baits) via 1,881 interactions, of which 87% were novel, with a median of 11 interactors per hcASD (**Figures 1C-E**). The ASD-PPI network showed significant overlap with previously published ASD protein networks^23,24^. For example, interactors from our dataset overlapped significantly with those from a network generated via immunoprecipitation of 13 endogenous hcASD proteins from human iPSC-derived excitatory neurons (‘Pintacuda 2023 PPI’)^23^ (**Figure S1D**, **Table S1**; matching 13 baits in ASD-PPI: OR = 1.77, p.adj = 0.0011; all ASD-PPI interactors: OR = 1.79, p.adj = 3.8 × 10^−7^). Interactors from Pintacuda 2023 PPI and from the matching 13 baits in ASD-PPI were both significantly enriched for ASD-associated *de novo* damaging variants (**Figure S1E, Table S1**; Pintacuda 2023 PPI: OR 1.44, p.adj = 0.04; ASD-PPI subset: OR = 1.94, p.adj = 0.0012;). We similarly found significant overlap between interactors from ASD-PPI and those from protein networks generated from proximity labeling of seven ASD risk proteins (including three hcASD) in HEK293T cells (‘HEK-PPI’) and of 41 ASD risk proteins (including 17 hcASD) in mouse cortical neuron and glia cocultures (‘Mouse-PPI’)^24^ (**Figures S1F,G, Table S1**; HEK-PPI OR = 3.15, p = 8.95 × 10^−23^; Mouse-PPI OR = 1.61, p = 1.56 × 10^−5^). Mouse-PPI interactors, but not HEK-PPI interactors, were significantly enriched for ASD-associated *de novo* damaging variants (**Figures S1H,I**, **Table S1**; HEK-PPI OR 1.33, p = 0.144; Mouse-PPI OR 1.62, p = 0.0053). Additionally, ASD-PPI interactors trended towards capturing more ASD-associated genetic risk (**Figure S1E,H,I**, **Table S1**; Breslow-Day test comparing differences in OR between external PPI dataset and ASD-PPI: Pintacuda 2023 PPI, p = 0.90; HEK-PPI, p = 0.96; Mouse-PPI p = 0.15). Notably, the total number of hcASD and associated interactors in ASD-PPI was significantly larger than those in existing human protein interactomes such as CORUM^50^, BioGRID^49,50^, InWeb^51^, Hein *et al*. 2015^51,52^ and BioPlex^48,51,52^ (**Figure 1E**) or recently published ASD-relevant protein interactomes^23,24^, making this the most comprehensive PPI dataset for ASD to date.

### ASD-PPI interactors are expressed in the human brain and enriched for ASD genetic risk

We next sought to evaluate the relationship of ASD-PPI interactors identified in HEK293T cells to cells in the human brain and assessed their relevance to genetic risk for ASD. We first examined RNA-sequencing (‘RNAseq’) datasets of the developing (BrainSpan^53^) and adult human brain (Genotype-Tissue Expression; GTEx^54,55^). ASD-PPI interactors showed significantly higher expression in both prenatal and adult brain tissue compared to other HEK293T-expressed proteins (**Figures 2A**, **S2A**, **Table S2**), suggesting that the ASD-PPI network is enriched for brain-relevant genes. We next compared hcASD proteins and their interactors (excluding interactors that are encoded by hcASD genes) with respect to their relative expression levels in BrainSpan and observed highly significant spatiotemporal correlation (**Figures 2B-C**, **Table S2**, Pearson R^2^ = 0.81, p < 1 × 10^−15^ by comparison to 100,000 permuted genesets with similar HEK293T protein expression levels). Furthermore, both hcASD proteins and their interactors exhibited higher expression in prenatal compared to postnatal samples (**Figures 2C**, **S2B**, **Table S2**; T-test, p.adj < 4.4 × 10^−16^), consistent with prior observations for hcASD genes^14–16^. Finally, in adult brain samples from GTEx, interactor expression was significantly higher in the cerebellum and cortex (compared to 100,000 permuted genesets) (**Figure S2C**, **Table S2**), in line with previous analyses of hcASD genes^6^. Altogether, these data indicate a highly similar pattern of expression between hcASD proteins and their interactors, suggesting that the ASD-PPI network is highly relevant to human brain tissue.

**Figure 2.**
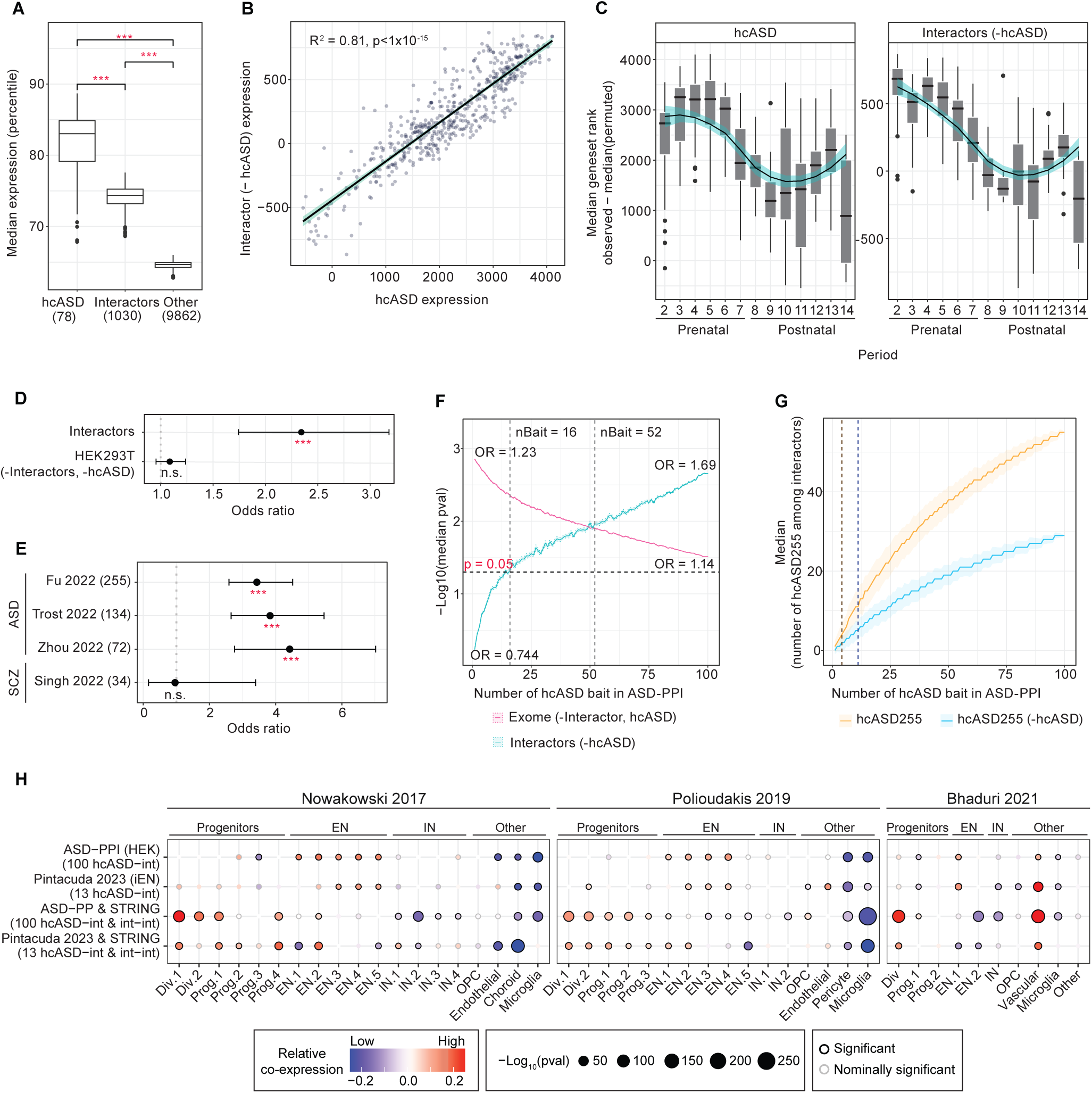
ASD-PPI interactors are expressed in the human brain and enriched for ASD genetic risk. (A) Differences in the median geneset expression percentile across n = 237 BrainSpan RNAseq prenatal brain samples^150^ for baits, interactors (-hcASD) and all other proteins expressed in HEK293T cells (‘Other’). (B) The relative expression levels of hcASD compared to interactors (-hcASD) within each of n = 524 BrainSpan RNAseq samples are significantly correlated (Pearson R^2^ = 0.81, p < 1 × 10^-15^). The relative expression within each brain sample was quantified by the difference between the median geneset rank of observed versus the median of 100,000 permuted genesets. (C) BrainSpan RNAseq samples were grouped by developmental period^150^, where periods 1-7 reflect prenatal stages of development and periods 8-15 reflect infancy through late adulthood. The relative expression levels of hcASD compared to interactors (-hcASD) across different periods are shown with an overlying Loess regression line, where gray shading reflects 1 standard error. The Spearman’s rho of the median rank difference of hcASD versus interactors (-hcASD) across periods was 0.946. (D) Geneset level burden test for *de novo* damaging variants in ASD probands compared with unaffected siblings from the Simons Simplex Collection^6^. (E) Enrichment of ASD-PPI interactors for three sets of ASD-associated risk genes and one set of SCZ-associated risk genes obtained from recent WES/WGS studies^3–5,60^. (F-G)The effect of increasing the number of baits used to construct the ASD-PPI network on the ability to capture interactors associated with ASD genetic risk (F) or identify interactors that are high-confidence ASD risk genes (‘hcASD255’) as defined by the latest ASD WES omnibus study^3^ (G). We downsampled the ASD-PPI network by selecting random sets of n = 1 to 100 baits and trimming the ASD-PPI network to include only the selected baits and associated interactors. In (F), for each downsampled network, we calculated the geneset-level burden of *de novo* damaging variants in ASD probands compared with unaffected siblings for interactors (teal) and all remaining genes in the exome (red); excluding hcASD from the analysis. Solid lines depict the median p-values across 1000 iterations; shaded regions indicate the median p-value +/- 1 standard error; the threshold for significance (p = 0.05) is labeled with a dashed black line. The median ORs for the genesets are labeled for ASD-PPI networks constructed using one bait (left) or 100 baits (right). Gray dashed lines indicate the ASD-PPI size at which the interactors captured a significant amount of ASD genetic risk (n = 16 baits) and when the interactors captured more ASD genetic risk than the remaining genes in the human exome (n = 52 baits). In (G), we defined ‘hcASD255’, brown, to be the n = 255 ASD risk genes with FDR < 0.1 and ‘hcASD255 (-hcASD)’, blue, to be the n = 174 hcASD255 genes that are not among the previously identified set of hcASD genes^6^. For each downsampled network, we calculated the median number of hcASD255 genes among the interactors, with shaded regions reflecting median number of hcASD255) +/- 1 standard error. We additionally calculated the the odds of interactors being enriched for hcASD255 genes and indicated the threshold for significance (median p < 0.05) with a dashed line (n = 4 baits for hcASD255, n = 11 baits for hcASD255 (-hcASD)). (H) Relative co-expression of ASD-PPI (generated in HEK cells) and a previously published ASD-relevant PPI network (‘Pintacuda 2023’^23^, generated in iENs) across cell types from three prenatal brain atlases (‘Nowakowski 2017’^62^, ‘Polioudakis 2019’^63^ and ‘Bhaduri 2021’^67^). hcASD-interactor (int) edges were defined by the indicated PPI network and int-int edges were extracted from STRING^66^ (see Methods). For each network, co-expression of two genes was measured by the proportion of cells in a given cell type with detected expression of both genes. To account for global differences in gene co-expression across cell types and differences in protein expression in the cell types that each PPI network was generated in, co-expression values were normalized by the cell type-specific average co-expression of all possible gene pairs. For each cell type, we evaluated whether the distribution of observed co-expression was significantly different from that of all cell types. Color reflects relative network co-expression, size reflects p-value, and edge color reflects significance. Abbreviations: div (dividing), EN (excitatory neuron), iENs (iPSC-derived EN), IN (inhibitory neuron), OPC (oligodendrocyte precursor cell), OR (odds ratio), prog (neural progenitor). Statistical tests: (A) T-test, (D, F, G) Fisher’s exact test (one sided, greater), (H) two-sample Wilcoxon rank sum. P-values corrected for multiple hypothesis testing (Bonferonni correction for: (A) three, (D) two, (E) four, (H) number of cell types in prenatal brain atlas × 2 tests). Boxplots in (A): boxes indicate first and third quartiles, line indicates the median and whiskers extend from the box to the highest or lowest value that is within the 1.5 × interquartile range of the box. Box and whisker plots in (D) (E): whiskers indicate 95% confidence interval.

We next evaluated whether the genes encoding interactor proteins tend to be highly evolutionarily constrained, like hcASD genes^3,5,6,19,20,56^, by comparing pLI (probability of being intolerant of a single loss-of-function variant), misZ (missense Z score, measures gene intolerance to missense variation), synZ (synonymous Z score, measures gene intolerance to synonymous variation, used as a negative control), and s_het (selective effect for heterozygous PTVs)^57,58^. The pLI and misZ scores of interactors (excluding interactors that are also hcASD) were in between those of hcASD and other HEK293T-expressed proteins, indicating that on average damaging mutations in interactors carry intermediate effect sizes (**Figure S2D**, **Table S2**). Consistent with this idea, the median s_het score of genes encoding interactors suggests they may act in an autosomal recessive or polygenic manner, in contrast to the majority of hcASD genes, which are thought to impart their major effects via haploinsufficiency or dominant-negative effects^58^ (**Figure S2D**).

Next, we examined interactors for enrichment of ASD genetic risk using data from the Simons Simplex Collection^6,59^. We observed a significantly greater burden of *de novo* damaging variants in genes encoding interactors (OR = 2.34, p.adj = 3.48 × 10^−7^, Fisher’s exact test comparing ASD probands to unaffected siblings) but not in the rest of the HEK293T proteome (OR = 1.08, p.adj = 0.30) (**Figure 2D**, **Table S2**). Moreover, interactors were enriched for ASD but not SCZ risk genes identified from recent whole exome (‘WES’) or whole genome sequencing (‘WGS’) studies^3–5,60^ as well as the highest confidence sets of ASD risk genes curated by SFARI Gene^61^ (**Figures 2E**, **S2E**, **Table S2**).

Finally, we evaluated the ASD-PPI network potential to identify additional ASD risk genes by creating networks with random sets of hcASD baits (varying from size 1 to 100, 1,000 iterations per size) and their associated interactors. We then assessed the genes encoding interactors in these networks (excluding interactors that are also hcASD) for enrichment of *de novo* damaging variants and for enrichment of an updated set of 255 high-confidence ASD genes (hcASD255)^3^. As the number of hcASD baits increased, the ASD-associated genetic risk captured by interactors also increased (**Figure 2F**, teal line), while the genetic risk attributable to the remainder of the exosome (excluding interactors and hcASD) from *de novo* damaging variants decreased (**Figure 2F**, red line). With n = 52 baits, genes encoding interactors were more significantly enriched for ASD-associated genetic risk than the remaining genes in the exome, and by n = 100 baits the interactors contained the majority of enrichment (**Figure 2F**, **Table S2**). Enlarging the ASD-PPI dataset also consistently increased the number of captured hcASD255 genes among the interactors (**Figure 2G**, **Table S2** brown line), even for those not previously identified in hcASD102 (**Figure 2G**, blue line). Thus, increasing the number of hcASD proteins used to construct the ASD-PPI appears to enrich for ASD rare variant genetic risk as well as increase the capture of novel ASD risk genes. Additionally, this effect does not appear to be plateauing, suggesting that continuing to build out ASD-PPI will be highly fruitful.

### ASD-PPI co-expression is greatest in dividing neural progenitor cells

Next, we assessed co-expression of the ASD-PPI network across cell types of the developing prenatal cortex (6-22 post-conceptual weeks, PCW; ‘Nowakowski 2017’)^62^ (see Methods). We observed that ASD-PPI bait-interactor interactions were highly co-expressed in excitatory neurons (**Figure 2H**, **Table S2**), consistent with prior findings^3,6,7,53,63,64,65^. Notably, after incorporating physical interactions from STRING^66^ to include interactor-interactor connections, co-expression became highly significant in neural progenitor cells and, to a lesser extent, excitatory neurons (**Figure 2H**, **Table S2**). Similar findings were obtained when analyzing two additional prenatal scRNAseq atlases from cortical samples aged 15-16 PCW (‘Pouliodakis 2019’)^63^ and 12-22 PCW (‘Bhaduri 2021’)^67^, or when assessing network co-expression of the Pintacuda *et al.* 2023 PPI network (‘Pintacuda 2023’)^23^, which consists of interaction data for 13 hcASD proteins from human iPSC-derived excitatory neurons (**Figure 2H**, **Table S2**).

### ASD-PPI reveals molecular convergence

Convergence of molecular and functional pathways has been previously observed among hcASD genes^7,68^. To determine whether this convergence is reflected in our data, we first arranged the proteins in the ASD-PPI network based on connectivity (using our derived PPIs) as well as similarity of Gene Ontology (GO) annotations^69,70^ (**Figure 3A**, **Table S3**) (see Methods). We then compared the ASD-PPI network to other PPI networks for baits with unified biological themes that have been previously generated using similar approaches (n = 90 tyrosine kinases^71^ or n = 39 breast cancer risk genes^26,27^). ASD-PPI baits were more likely to share interactors (**Figure 3B**), suggesting higher levels of molecular convergence compared to the other two PPI maps. Furthermore, 359 of the 1,043 (34.4%) interactors (that are not encoded by hcASD genes) interacted with more than one hcASD bait (**Figure S3A, Table S3**), and many baits converged on common complexes (**Figures 3C-F**). ASD and neurodevelopmental delay (NDD) share many risk genes^7^, and Satterstrom *et al*. 2020 stratified the hcASD102 genes into those more frequently mutated in ASD (ASD-predominant or ASD_P_, n = 53) and those more frequently mutated in NDD (ASD with NDD or ASD_NDD_, n = 49)^6^. Interestingly, the level of interactor overlap did not differ among all ASD-PPI bait pairs, ASD_P_ bait pairs, ASD_NDD_ bait pairs, or ASD_P_-ASD_NDD_ bait pairs (**Figures S3B**, **C**, **Table S3**, Kruskal-Wallis p = 0.15), suggesting that ASD_P_ and ASD_NDD_ baits have similar network properties in these data.

**Figure 3.**
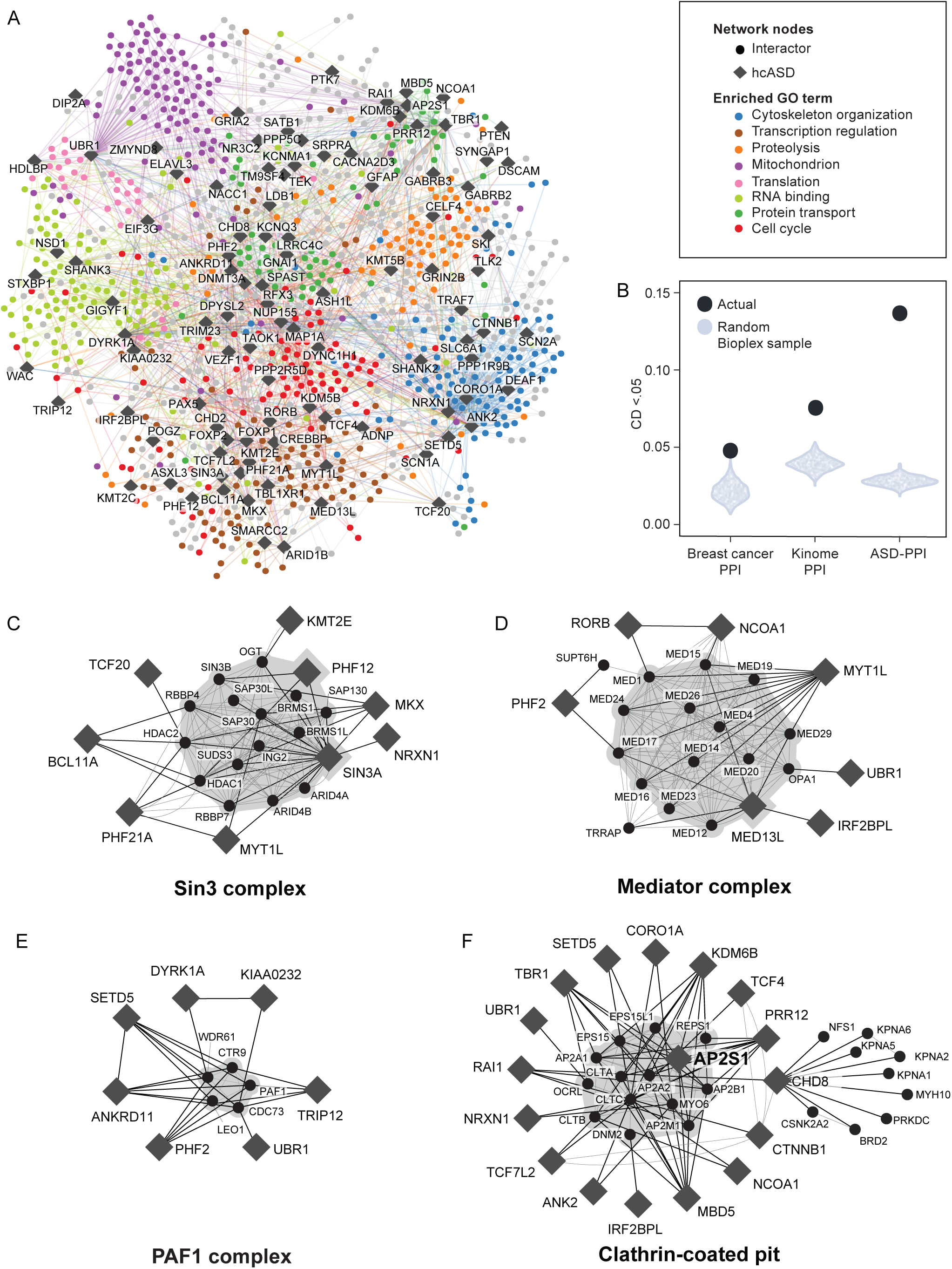
The ASD-PPI network demonstrates molecular convergence among hcASD. (A) Interactors and baits are arranged in two dimensions to best characterize their relative distances based on connectivity in our AP-MS data and shared GO annotations. Coloring of interactors is based on a selection of a small subset of the GO terms that are enriched in the full 1043 interactor set. GO terms are chosen to best cover the full set while balancing redundancy. See Methods for further details. (B) Interactor overlap with different baits in ASD-PPI, Breast cancer PPI and Kinome PPI as measured as proportion of significant overlap between the interactor-sets for all pairs of baits. Significant overlap is measured by p-values from hypergeometric tests, and the portion with p < 0.05 (CD < 0.05) is plotted. Each dataset was compared to 1000 random bait-int sub-networks of BioPlex, selected to have similar network size and degree, to establish a baseline. (C-F): PPI network for hcASD interaction with Mediator complex (C), Sin3 complex (D), PAF1 complex (E), and AP2-mediated clathrin-coated pit complex (F). Dark lines show AP-MS edges, thin gray lines indicate CORUM or STRING edges, and gray shading indicates CORUM complexes.

Notably, we identified several complexes that included multiple interactors with additional evidence for ASD. For example, several hcASD proteins, most of which have previously been connected to transcription or chromatin modification (TCF20, KMT2E, PHF12, MKX, MYT1L, PHF21A, BCL11A), interacted with the Sin3 complex (**Figure 3C**). Sin3-containing complexes act as transcriptional co-repressors, play a role in progenitor cell proliferation and differentiation^72,73^, processes that may be disrupted in ASD^7,1774^. We also found evidence that multiple hcASD proteins may functionally converge upon the mediator complex (**Figure 3D**), a multiprotein complex that functions as both a transcriptional activator and repressor, and plays a role in establishing neuronal identities by defining neuronal gene expression^75,76^. While we detected hcASD proteins (MYT1L, NCOA1) that are known to interact with the mediator complex, we also detected novel hcASD protein interactors such as PHF2, RORB and IRF2BPL (**Figure 3D**). The PAF1 complex also plays a role in chromatin modification and transcription regulation, and CTR9 (FDR = 0.07^3^) and LEO1 (FDR = 0.11) subunits are below or at the threshold for high confidence association (FDR < 0.1) in a recent omnibus WES study of ASD^3^ (**Figures S3D**, **E**). Along with known interactors of the PAF1 complex such as SETD5^77^, we identified interactions with other hcASD proteins such as DYRK1A, ANKRD11, TRIP12, PHF2 and KIAA0232 (**Figure 3E**). Besides complexes involved in gene expression regulation, we also identified the AP2-associated clathrin-mediated endocytosis complex as an interactor with a large number of hcASD proteins. Notably, subunits of the clathrin-mediated endocytosis complex interacted with multiple transcription factors such as TCF7L2, CHD8, TBR1, and RAI1, indicating a complex interplay between transcription factors, which may have pleiotropic functions outside of gene regulation^22^ and protein transport (**Figure 3F**). Additional work will help determine how these complexes are connected to the underlying biology of ASD.

### AlphaFold predicts direct PPIs

Recent developments in machine learning approaches have enabled accurate predictions of 3D structures of interacting proteins from amino acid sequences^78–80^, providing insights into the direct interactions between proteins^81,82^. ASD-PPI allowed for a unique opportunity to apply sequence-based PPI prediction algorithms to a constrained, relevant molecular search space that is enriched for molecular complexes (see **Figure 3**) with the goals of identifying and prioritizing direct PPIs for follow-up studies and identifying relevant 3D structural interaction interfaces. We therefore used AlphaFold-Multimer^78^ (referred to below as AF) to predict pairwise interactions of either hcASD-interactor (bait-int) or interactor-interactor (int-int) pairs (**Figure 4A, Table S4**). As a control, pairwise predictions of a random set were included, where each interactor in the bait-int set was replaced with a similarly sized protein (bait-random; see Methods). We examined the typical use of scores, either maximum ipTM (a predicted measure of topological accuracy in the interface^78^) or confidence (a measure which includes interface and overall topological accuracy) across nine generated models per pair and found only weak separation between observed and random sets (**Figure S4A**). In contrast, we found that mean ipTM, a statistic that summarizes the whole distribution of AF scores and favors protein pairs with consistently high scores across models, performed well to separate the observed from random sets, with an approximate 5 to 10-fold enrichment for score thresholds above a mean ipTM of 0.5 (**Figures 4B**, **S4B**). Of the 1,651 bait-int predictions completed by AF, 113 hcASD-int pairs had a score of mean ipTM > 0.5 (blue bars, **Figure 4C**), and 466 additional interactors connected to a hcASD protein indirectly via one or more int-int pairs with mean ipTM > 0.5 (purple bars, **Figure 4C**), accounting for 579 of 1,094 total AP-MS interactions where a hcASD-interactor is predicted by AF to have any interactions with mean ipTM > 0.5 (**Figure S4D**, **Table S4**). We observed that the portion of interactor pairs with high ipTM score was lower in the int-int set than bait-int regardless of applied threshold, which likely reflects the different portions of direct connections between bait-int versus int-int in the AP-MS datasets. We found that the ASD-PPI network showed higher rates of AF-predicted direct physical interactions between hcASD and interactor proteins as compared to a PPI network generated in human iPSC-derived excitatory neurons described above (‘Pintacuda 2023 PPI’^23^), with nearly twice the rate at mean ipTM > 0.5 (purple curve, **Figure 4B**). This demonstrates once again, in addition to the gene expression and genetic risk analysis described above, that AP-MS in HEK293T cells identifies ASD-relevant hcASD interactors and suggests that ASD-PPI may be relatively enriched for direct interactions.

**Figure 4.**
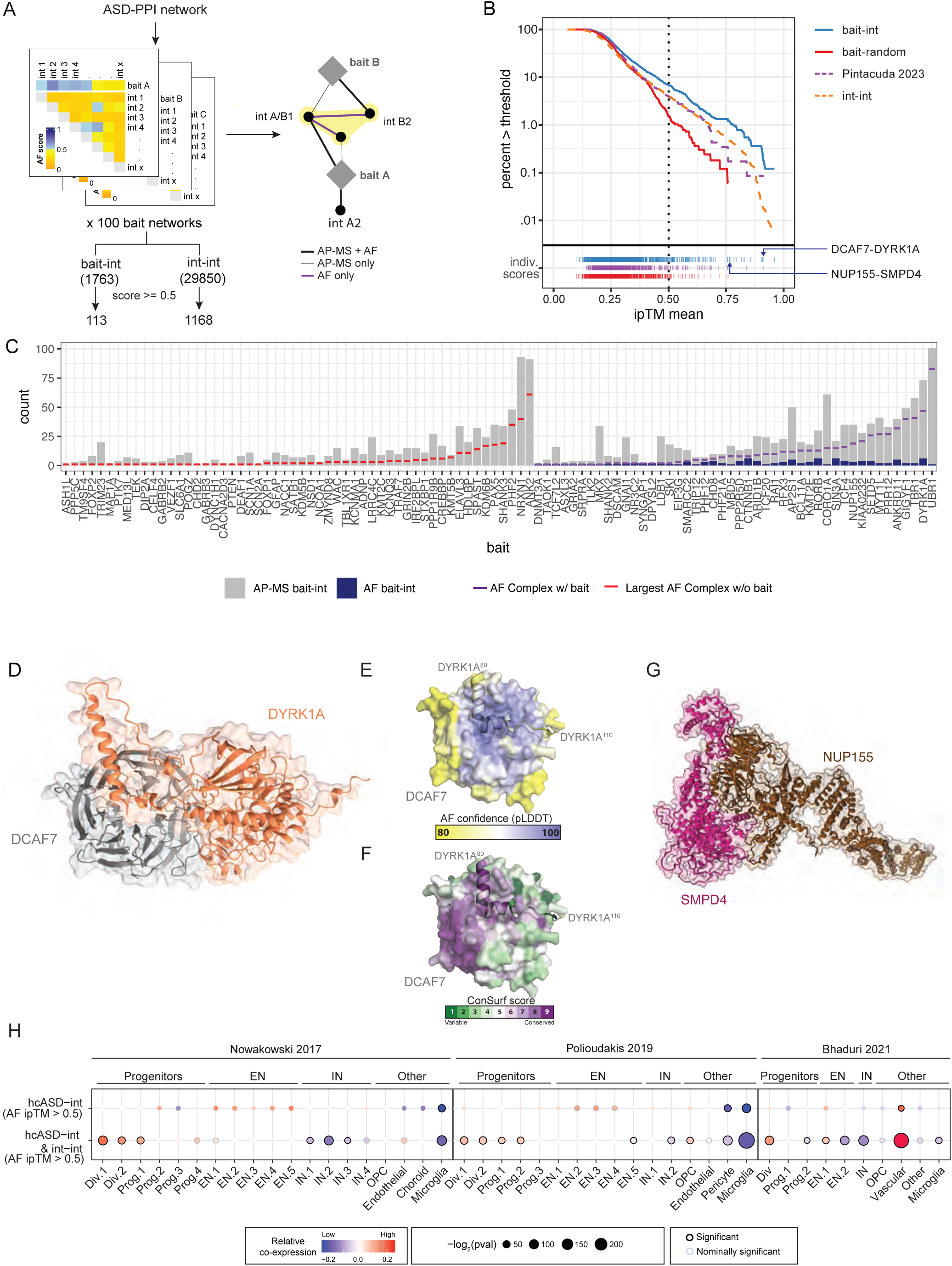
AlphaFold predicts PPI interaction interfaces. (A) Workflow of AF (Multimer) interface predictions for hcASD bait-int subnetworks. AF was run on every bait-int pair, and every int-int pair within a bait-int subnetwork. This resulted in two sets of completed predictions labeled bait-int (1763 runs) and int-int (29,850 runs). After filtering AF scores for direct interactions, the 113 bait-int pairs and 1168 int-int pairs provide a network of direct interactors overlaid on ASD-PPI (example subnetwork shown). (B) Cumulative distributions of mean ipTM scores in four different sets of AF runs allows for comparison of enrichment of high scores between sets. Higher lines show greater proportions of high scores, with separation between wider horizontal lines equal to a 10-fold increase in high scoring AF runs at a given threshold. Bait-int is enriched over bait-random approximately 10-fold at mean ipTM > 0.75 and is enriched ∼5-fold at mean ipTM > 0.50, the threshold we established (dotted vertical line). PPI found by Pintacuda *et al.* 2023 are included for comparison. Distribution of individual scores shown at bottom, except for int-int which is too numerous, with NUP155-SMPD4 and DCAF7-DYRK1A indicated. (C) Number of AP-MS bait-int pairs (gray) or AF-supported bait-int pairs (blue) for each hcASD protein. Horizontal bars within each gray column show the number of interactors joined via direct and indirect interactions (a “complex” if all pairwise interactions are simultaneous) either to the bait (purple) or in the largest interactor-only “complex” (red) for interactor sets with no AF bait-int connection. (D) AF predicted structure for DYRK1A-DCAF7. Regions with very low AF confidence are hidden for surfaces (pLDDT < 25) and ribbons (pLDDT < 20). (E) AF modeling confidence (pLDDT) per residue plotted for DYRK1A-DCAF7 interaction with pLDDT scores ranging from 80 (moderate confidence, yellow) to 100 (maximum confidence, blue). DCAF7 is drawn as a surface and residues 80-110 on DYRK1A are drawn as ribbons. (F) Same view as (E), but colored by ConSurf score, a measure of sequence conservation per residue. Scores range from 1 (low conservation, green) to 9 (high conservation, purple). (G) AF predicted structure for NUP155-SMPD4. Regions with very low AF confidence are hidden for surfaces (pLDDT < 25) and ribbons (pLDDT < 20). (H) Relative co-expression of ASD-PPI-AF network for hcASD-int (top) and for a denser network including predicted int-int connections (bottom) across cell types from three prenatal brain atlases (‘Nowakowski 2017’^62^, ‘Polioudakis 2019’^63^ and ‘Bhaduri 2021’^67^). Relative co-expression across cell types was evaluated as in Figure 2H.

The interaction of DYRK1A with DCAF7, well documented previously^83–85^, including a recognized DCAF7 binding motif, but with no available structure, provides an example of a high-confidence interface prediction by AF. The interaction received a high overall score (ipTM = 0.931, **Table S4**), the interface matches the recognized binding motif^84,85^ (**Figure 4D**), is modeled with high per-residue confidence (**Figure 4E**), and the residues involved directly in the interface are among the most conserved (**Figure 4F**). We overexpressed WT DYRK1A and DYRK1A^Δ80–100^ in HEK293T cells and confirmed that DYRK1A^Δ80–100^ lost interaction with DCAF7 but not with FAM54C, a protein that has been shown to bind the DYRK1A catalytic kinase domain (residues 156-479)^86^ **(Figure S4E, Table S4)** Together, these findings demonstrate that AF can identify a direct PPI between DYRK1A-DCAF7 and highlight a specific interface that mediates this interaction.

The interaction of hcASD NUP155 with SMPD4 represents an example of a direct interface prediction by AF (mean ipTM = 0.765, **Figure 4G**) that has not been previously described. Previous studies, though underpowered, have shown that loss-of-function variants of SMPD4, a sphingomyelinase, are associated with congenital microcephaly and developmental disorders^87,88^, underlining the power of this approach in identifying direct interactions potentially relevant to a broader range of neurodevelopmental disorders that warrant further research.

To determine if AF-supported direct interactions within ASD-PPI are enriched in specific cell types, we evaluated network co-expression in the three prenatal brain scRNAseq cell atlases^62,63,67^ as described above (see **Figure 2H**). We found that AF-supported direct bait-int connections trended towards higher co-expression in excitatory neurons (**Figure 4G**, **Table S4**). However, when AF int-int connections were considered, network co-expression was significantly higher in neural progenitor cells, similar to our observation with networks that incorporated int-int connections from STRING^66^ (**Figure 4G**, **Table S4**).

### DCAF7 physically interacts with DYRK1A and KIAA0232

As previously noted, the interaction between DCAF7 and the hcASD gene DYRK1A is well established^85,89–91^ and AF predicted it to be a direct interaction with a high ipTM score (**Figure 4B**). Of the seven interactors associated with eight or more hcASD proteins (**Figure 5A**, **S3A**), only DCAF7 had interactions in the public interactome BioGRID (the only considered interactome with entries for all seven interactors) that were enriched for the ASD risk genes identified from a recent ASD omnibus study (hcASD255)^3,92^ (**Figure 5B**, **Table S5**, Fisher’s exact test with Bonferroni correction: OR = 5.79, p = 0.018, p.adj = 0.126). These collective data suggest that proteins encoded by hcASD genes converge upon DCAF7 at a molecular level.

**Figure 5.**
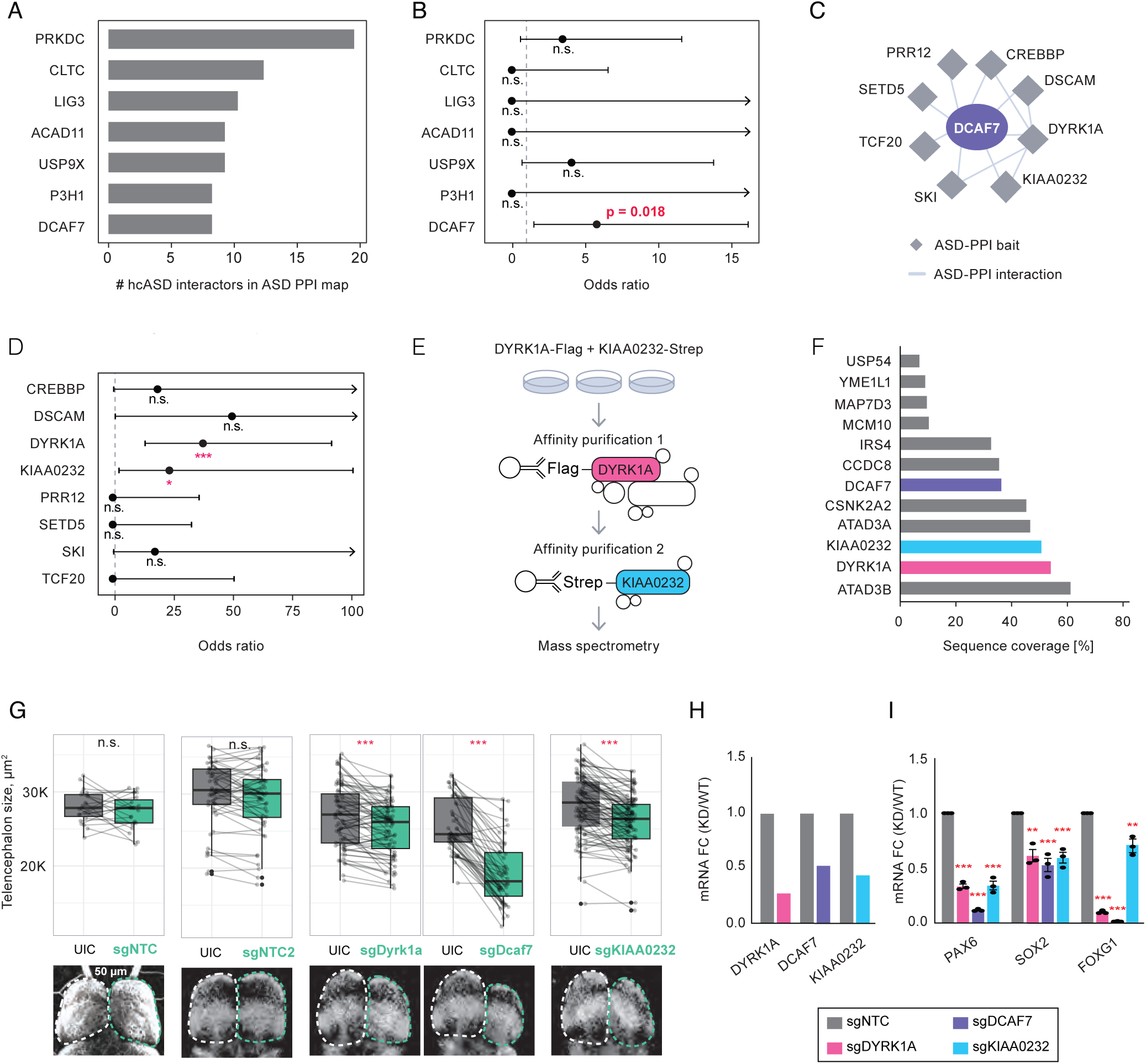
DCAF7 is a nexus for multiple hcASD. (A) The top 7 ASD-PPI interactors, ranked by the number of hcASD interactions. (B) Enrichment tests evaluating whether BioGRID^92^ interactors of the seven indicated proteins are enriched for hcASD255^3^. Fisher’s exact test (one sided, greater), p-values not adjusted for multiple hypothesis testing; whiskers indicate the 95% confidence interval, with upper interval cutoff set at OR = 15. (C) DCAF7 network showing interaction with hcASD genes. (D) Enrichment tests evaluating the overlap between DCAF7 BioGRID interactors and the ASD-PPI interactors for each of the 8 hcASD baits that bind DCAF7 in ASD-PPI. Fisher’s exact test (two-sided) gene universe was restricted to genes in the HEK293T proteome, p-values adjusted for 8 tests (Bonferroni). Whiskers indicate the 95% confidence interval, with upper interval cutoff set at OR = 100. (E) Workflow for sequential IP for DYRK1A (Flag-tag) and KIAA0232 (Strep-tag). (F) Sequence coverage by MS of significant interactors following sequential IP in (E) of DYRK1A and KIAA0232 highlighting DYRK1A, DCAF7 and KIAA0232. (G) Telencephalon sizes of Xenopus injected unilaterally with either guide RNAs against a non-targeted control or DCAF7, DYRK1A or KIAA0232. Data shows significant reduction in sizes plotted against the contralateral (noninjected) hemisphere for DYRK1A knockdown, DCAF7 knockdown or KIAA0232 knockdown. Scale bar: 50 μM. (H) CRISPR mediated knockdown shows reduced levels of DYRK1A, DCAF7 and KIAA0232 respectively. (I) Quantitative RT-PCR in NPCs with DCAF7, DYRK1A or KIAA0232 knockdown shows significant decrease in PAX6, SOX2 or FOXG1 expression as compared to non-targeted control (NTC). Data is shown as mean ± SEM, n = 3. n.s. nonsignificant, ∗p < 0.05, ∗∗p < 0.01, ∗∗∗p < 0.001

DCAF7 interaction with DYRK1A modulates its nuclear translocation and regulates interaction with other proteins such as Huntingtin-associated-protein 1 (HAP1), a protein implicated in delayed growth in Down syndrome^90,93^. Furthermore, a DCAF7 complex with AUTS2 and SKI regulates neuronal lineage specification *in vivo*^94,95^, pointing to a key role for DCAF7 in neurodevelopment. To prioritize the eight hcASD baits that interact with DCAF7 in the ASD-PPI (**Figure 5C**), we performed enrichment tests using the DCAF7 interactors defined in BioGRID and found significant overlap between the DCAF7 interactors and the ASD-PPI interactomes of DYRK1A and KIAA0232 (**Figure 5D**, **Table S5**, Fisher’s exact test with Bonferroni correction: DYRK1A interactome OR = 37.74, p.adj = 3.27×10^−8^; KIAA0232 interactome OR = 23.69, p.adj = 0.032). To further study the interactions shared between DCAF7 and DYRK1A, we co-expressed FLAG-tagged DCAF7 and Strep-tagged DYRK1A in HEK293T cells and performed single-tag as well as sequential (double) AP-MS, identifying 126 shared interactors between DYRK1A and DCAF7. These shared interactors were significantly enriched for hcASD255 (**Figures S5A,B**, 9 hcASD, OR = 4.19, p = 0.000565). Given the role of DCAF7 as a WD40 repeat containing scaffold protein^96,97^, these results present the intriguing possibility that DCAF7 functions as a scaffold, facilitating interactions with other hcASD proteins. In addition to capturing some well documented interactions, such as CREBBP^98^, TRAF2^98,99^ and TSC1^98–100^, we also found KIAA0232, a hcASD gene with unknown function, to interact with DCAF7 and DYRK1A. By performing sequential AP-MS of FLAG-tagged DYRK1A and Strep-tagged KIAA0232, we identified DCAF7 as a shared interactor, confirming that DYRK1A, DCAF7 and KIAA0232 interact with each other (**Figures 5E**, **F**).

### DCAF7, DYRK1A and KIAA0232 functionally overlap

We next evaluated whether DCAF7, DYRK1A, and KIAA0232 colocalize in human cells. We overexpressed DYRK1A, DCAF7 and KIAA0232 in HEK293T cells and found that all three proteins colocalize to the mitotic spindle (**Figure S5D**). Prior work has demonstrated that multiple hcASD proteins localize to the spindle during mitosis including in *Xenopus* and human iPSC-derived neural progenitor cells (NPCs)^22,101,102^. Consistent with these findings, we observed that ASD-PPI interactors are enriched for proteins associated with structures important for spindle organization (centriolar satellites^103^ and centrosomes^104^; **Figure S5C**, **Table S5**; two-sided Fisher’s exact tests with Bonferroni correction: HEK293T centriolar satellite, OR = 2.78, p.adj = 1 × 10^−19^; NPC centrosome, OR = 2.6, p.adj = 9.9 × 10^−19^; neuron centrosome, OR = 2.7, p.adj = 6 × 10^−21^). Together, these results provide further support for the potential involvement of tubulin biology in ASD pathogenesis.

Next, we targeted these three genes during brain development using CRISPR-Cas9 mutagenesis in *X. tropicalis* by injecting single guide RNAs (sgRNAs) unilaterally, which allowed for matched internal controls. We observed that knockout of *DCAF7, DYRK1A*, or *KIAA0232* resulted in significantly smaller telencephalon size, further confirming the role of DYRK1A, DCAF, and KIAA0232 in neurogenesis (**Figure 5G**, **Table S5**; paired T-test with Bonferroni correction: *DYRK1A*, p.adj = 4.14 × 10^−6^; *DCAF7*, p.adj = 2.09 × 10^−16^; *KIAA0232*, p.adj = 1.11 × 10^−12^).

To further evaluate functional outcomes of *DYRK1A, DCAF7* or *KIAA0232* perturbations, we performed CRISPRi mediated knockdown (KD) of each gene individually in human iPSC-derived NPCs with single guide RNAs (sgRNAs: sgDYRK1A, sgDCAF7, sgKIAA0232). We first confirmed the gene KD for each cell line by qPCR, showing a reduction in expression of *DYRK1A, DCAF7* or *KIAA0232* by 50-60%, respectively (**Figure 5H**). DCAF7 has been shown to be essential for survival^89,105–108^, with loss of DCAF7 leading to cell loss over time^89,106,108^, possibly due to cell death^109^. Consistent with this, we observed increased cell death of sgDCAF7 cells using the genetically-encoded death indicator (GEDI)^110^ during neuronal differentiation (**Figures S5E,F, Table S5**). DYRK1A has been shown to be involved in cell cycle regulation in a dose dependent manner, with moderate KD promoting neurogenesis and complete inhibition of DYRK1A inhibiting neurogenesis^111,112^. We observed a reduction in the proliferation marker Ki67 in both sgDCAF7 and sgKIAA0232 NPCs while a modest increase was observed in sgDYRK1A cells (**Figures S5G,H, Table S5**). DYRK1A further regulates PAX6^113^, which in turn maintains the balance between proliferation and neural specification^114,115^. Notably, mRNA expression of *PAX6* as well as *FOXG1* and *SOX2*, markers which are expressed in NPCs and whose deficits lead to proliferation defects and changes in neuronal differentiation patterns^116,117^, were significantly reduced in all three KD NPCs (**Figure 5I, Table S5;** One Way ANOVA with Dunnett correction: *PAX6*, all, p.adj <0.0001; *FOXG1*, sgDYRK1A, sgDCAF7, p.adj < 0.0001, sgKIAA0232 p.adj = 0.0012; *SOX2* sgDYRK1A, p.adj = 0.0012; sgDCAF7, p.adj = 0.0003; sgKIAA0232, p.adj = 0.0009).

### ASD missense mutations affect protein-protein interactions

We next assessed how ASD-associated *de novo* missense mutations impact the ASD-PPI network. ASD patient-derived *de novo* missense mutations that were predicted to be highly deleterious (Missense badness, Polyphen-2, and Constraint (MPC) score ≥ 2)^118^ were selected, resulting in a list of 87 damaging *de novo* missense mutations across 43 hcASD genes^6^. Using the same methods as for ASD-PPI, we mapped PPIs for 54 of these missense mutations across 30 hcASD proteins (ASD_mut_-PPI), including 13 hcASD proteins encoded by genes with more than one patient-derived mutation, and identified 1,070 interactions (**Figure 6A, Table S6**). To investigate differential interactions between mutant and WT hcASD proteins, i.e., strengthened or weakened interactions, we normalized interactor intensities by bait intensity (accounting for changes in interactor intensity derived from differences in bait expression) and calculated the fold change in interactor intensity between mutant and WT baits. We defined significant interaction changes as those with a log_2_(fold-change) ≥ 1 and p ≤ 0.05. Of the 1,070 interactions identified from WT or mutant baits, we found that the majority were preserved (817 interactions, 324 unique interactors), while a subset were strengthened (117 interactions, 69 unique interactors) or weakened (136 interactions, 95 unique interactors) for at least one mutant hcASD bait (**Figures 6B,C, Table S6**). We observed significant enrichment of hcASD255 genes among the 95 lost interactors but not the 69 gained interactors (lost: OR = 3.63, p.adj = 0.0171; gained: OR = 1.58, p.adj = 0.738; two-sided Fisher’s exact test with Bonferroni correction) (**Figure S6A**, **Table S6**).

**Figure 6.**
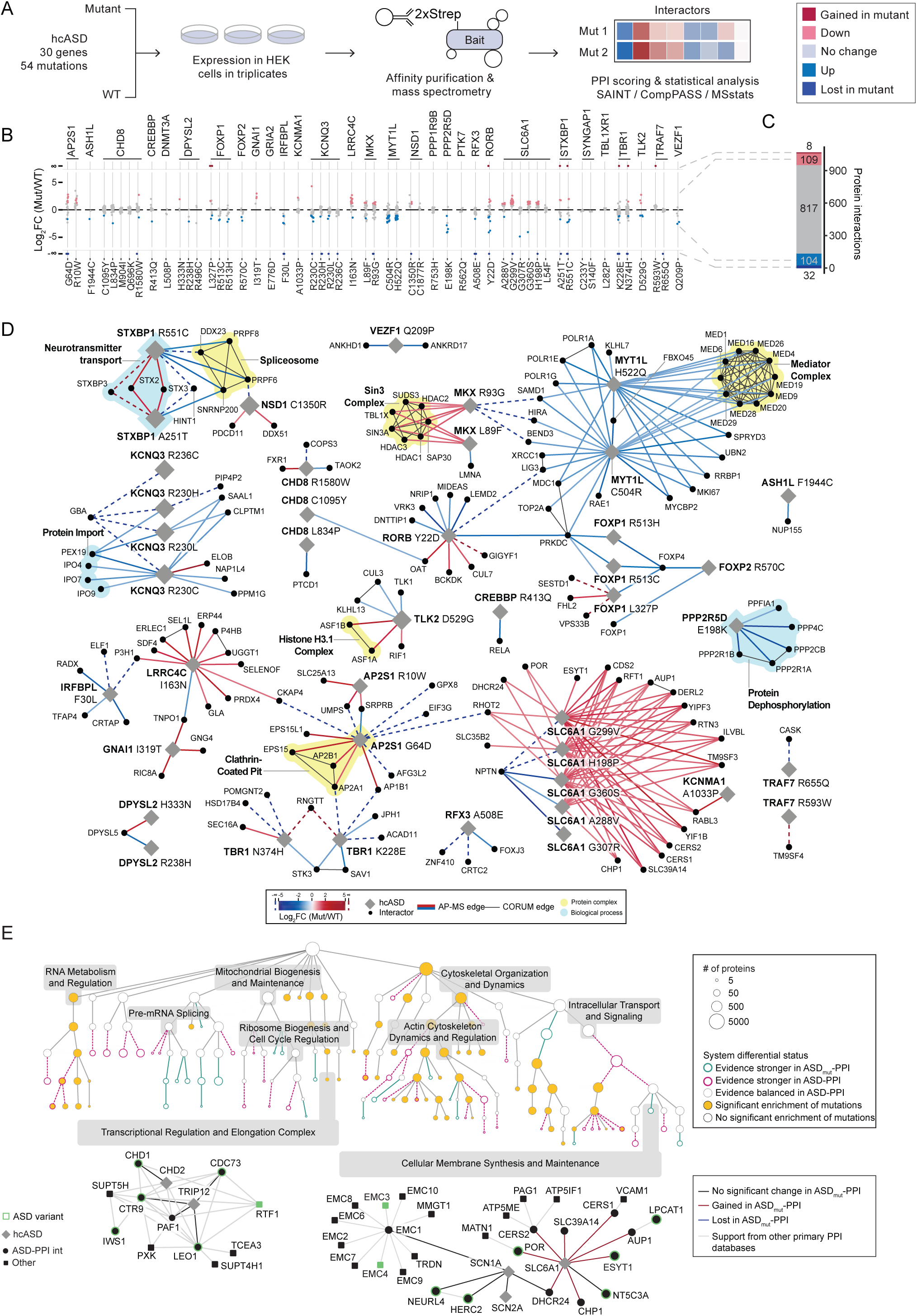
Patient-derived hcASD missense mutations alter protein interactions. (A) Overview of the generation of the ASD mutant interactome (ASD-ASD_mut_-PPI). We prioritized damaging *de novo* missense mutations in hcASD with (MPC ≥ 2) that were observed in ASD individuals^6^. Strep-tagged WT and mutant variants were overexpressed in parallel in HEK293T cells and subjected to AP-MS. High confidence interactions were identified using SAINT and CompPASS, and log_2_(fold-change) of the interactor intensity between mutant and wildtype baits was calculated using MSstats. (B) Dot-plot visualization of the loss and gain of interactors in hcASD mutant variants, grouped by parent hcASD. Each dot represents a high-confident interactor in either the wildtype or the mutant baits. (C) Quantification of the differential protein interactors identified 117 interactors (red) to have higher affinity for the mutant hcASD and 136 interactors (blue) for the wildtype hcASD. (D) Differential interactions identified from the ASD_mut_-PPI interactome. Diamond shapes denote the mutant baits and circles denote their respective interactors. The color scale of the edges corresponds to the specificity of the interactions, where blue edges have stronger affinity for the wildtype hcASD and red edges have higher affinity to the mutant hcASD. CORUM protein complexes among interactors are highlighted in yellow. (E) Multi-scale hierarchical layout of protein systems annotated based on whether the system has stronger evidence in ASD_mut_-PPI (green border) or ASD-PPI (pink border) as well as with mutant enrichment status (yellow fill: systems enriched for functionally disrupting mutated genes). Circle size corresponds to the number of proteins included in the respective system. Examples of systems that were either not affected by ASD_mut_-PPI (lower left) or had differential interactions from ASD_mut_-PPI (lower right) are shown below. Diamond shapes denote the mutant baits and circles denote their respective interactors in ASD_mut_-PPI; squares represent protein interactors not found in this study. Green highlights on nodes indicate ASD-related genes from Satterstrom *et al.* 2020^6^ not included in our hcASD list (see Methods). Edge color corresponds to the source and specificity of interactions (black = no change between ASD-PPI and ASD_mut_-PPI, blue = stronger affinity for WT hcASD, red = higher affinity for mutant hcASD, gray = from other PPI databases).

### Patient-derived mutations result in convergent interaction changes

We next aimed to determine if multiple different missense mutations in the same hcASD gene demonstrated similar effects on PPIs, and whether missense mutations in different hcASD genes converged on similar changes in the PPI network. For the 13 hcASD genes with PPI data for more than one mutant bait, 52 of the 115 altered interactions were replicated in two or more mutant baits for the same hcASD protein. For example, all five mutant SLC6A1 baits lost interactions with NPTN, all four KCNQ3 mutant baits lost interactions with GBA, and all three FOXP1 mutant baits lost interactions with FOXP4 and PRKDC (**Figures 6D**, **S6B**, **Table S6**). We additionally found that 15 interactors showed consistent changes for two different mutant hcASD baits; for example, the interactor FOXP4 lost interaction with all three FOXP1 mutant baits as well as one FOXP2 mutant bait, which was validated by Western blot (**Figures 6D**, **S6B,C**, **Table S6**). Notably, we identified changes in patient-derived mutation driven interactions that converged on key complexes or biological processes mentioned above (**Figures 3C-F, Table S6**). Among the differential interactors, we observed enhanced interaction of transcription factor MKX mutants (MKX^R93G^ and MKX^L89F^) with the transcriptional repressor Sin3 complex. Similarly, MYT1L mutations H522Q and C504R showed decreased interaction with the mediator complex, another transcriptional regulator. We also observed an increase in gained interactions with vesicle mediated transport and clathrin-mediated endocytic complex in STXBP1 mutants (STXBP1^R551C^, STXBP1^A215T^) and AP2S1 mutants (AP2S1^R10W^, AP2S1^G64D^) (**Figure 6D**). Given the key roles that Sin3 and mediator complex play in gene expression regulation, the ASD_mut_-PPI network provides key insights into convergent mechanisms that may be dysregulated in the context of patient-derived mutations.

To gain a higher-level understanding of biological processes most affected by ASD-related mutations, we generated a multi-scale map of protein systems by integrating ASD-PPI with other PPI networks^23,48,52,92,119,120^, and analyzed network reorganization upon including vs. excluding differential interactions from ASD_mut_-PPI (for details, see Methods). This multi-scale map consists of 422 protein systems, with network reorganization highlighted by 22 and 36 protein systems showing stronger evidence in ASD_mut_-PPI or ASD-PPI, respectively. We also identified 61 systems with an enrichment of functionally disrupting mutations described in Satterstrom *et al*.^6^ that were not part of ASD_mut_-PPI (i.e. variants not limited to hcASD; see Methods for details) (**Figure 6E, Table S6**). For example, the protein system “Cellular Membrane Synthesis and Maintenance”, which consisted of 36 proteins including three hcASD (SLC6A1, SCN2A, and SCN1A), had differential interactions from ASD_mut_-PPI (i.e. contained interactions gained or strengthened compared to ASD-PPI; see also **Figure 6D**) and contained two additional genes (ER membrane protein complex subunit 3 & 4; EMC3 and EMC4) that were described as *de novo* damaging variants in ASD probands^6^. Conversely, the protein system “Transcription Regulation and Elongation Complex” was not affected by ASD_mut_-PPI. This system consisted of 13 proteins, including two hcASD (TRIP12 and CHD2), the PAF1 complex (see also **Figure 3E**), and RTF1^121^, another gene with *de novo* damaging variants^6^ not included in ASD_mut_-PPI.

We next leveraged the ability of AF to predict direct PPIs and prioritized mutations predicted to be located at the interaction interface (**Figure 7A**). Using the differential PPIs from our ASD_mut_- PPI network, we mapped mutations onto the AF predicted structures for ASD-PPI. We found 34 mutations across 22 hcASD proteins that mapped to the interfaces (< 10 Å) with 198 different interactors, corresponding to 216 pairwise interactions. Out of these, 73 interactions were impacted in ASD_mut_-PPI, showing differential interactions in the presence of mutations (**Table S7**). The interactions that decreased in the mutant state were enriched for hcASD mutants in interfaces (46% located in interface); in contrast, only 18% of hcASD mutations that led to an increase in interactions were located in the interface, indicating that the position of mutations in the interaction interface tended to correlate with the direction of differential interactions (**Figure S7A**, **Table S7**). For example, the interaction between PPP2R5D and PPP4C was lost in the presence of the PPP2R5D mutation E198K, which directly contacts the interaction interface (1.6 Å) (**Figures 7B,C**, **S7B**). In contrast, a mutation in GNAI1 (I319T) more distant to its interface with its known interactor RIC8A^122^ (6.1 Å) strengthened the PPI (**Figures 7D,E**, **S7C**).

**Figure 7.**
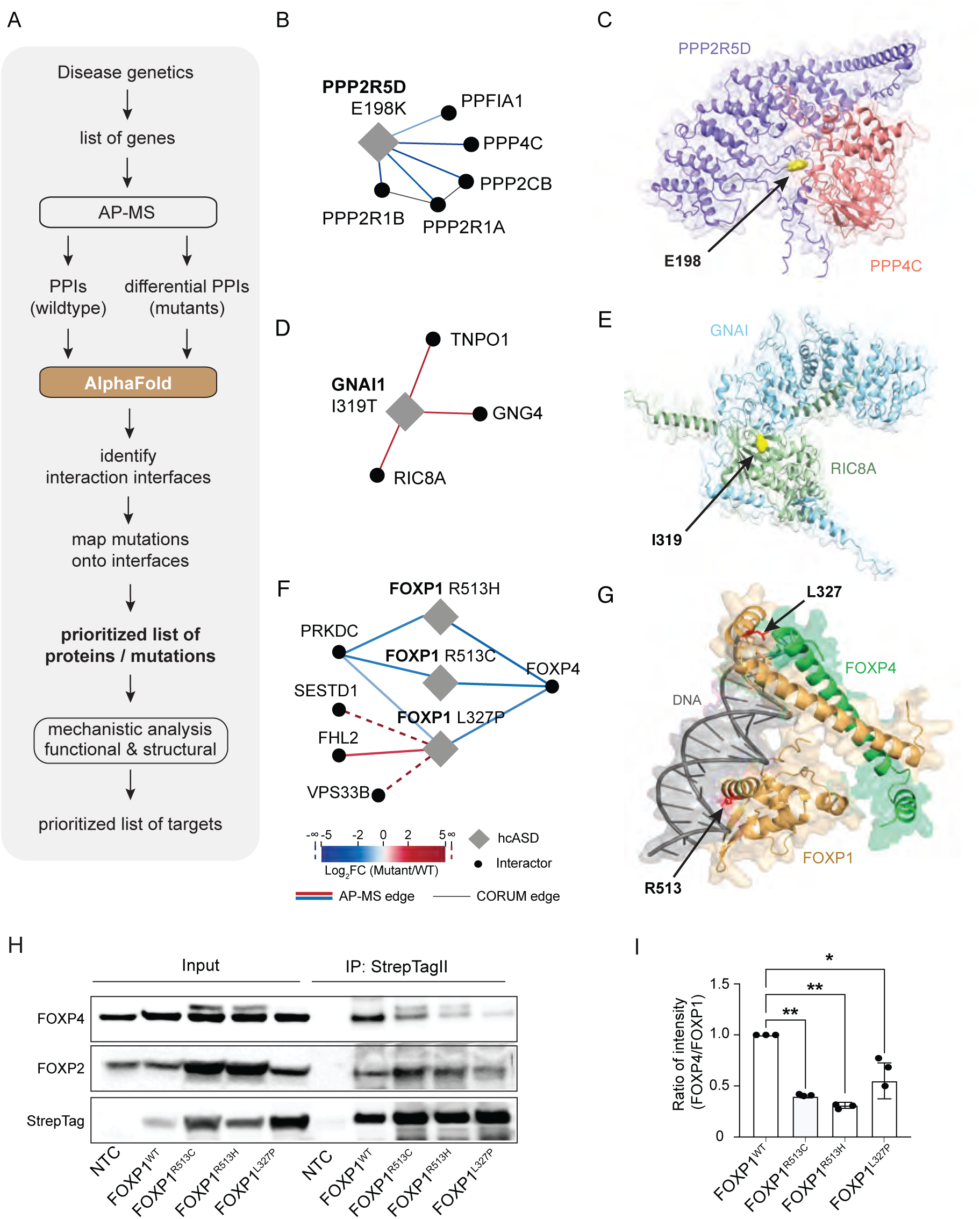
Mapping mutations using AlphaFold. (A) Workflow for prioritizing disease-relevant mutations based on AP-MS and AF structure predictions. (B,D,F) Differential ASD_mut_-PPI networks for PPP2R5D^E198K^ (B), GNAI1I^319T^ (D), and FOXP1^R513H^, FOXP1^R513C^, FOXP1^L327P^ (F); blue and red lines indicate loss and gain of interaction with mutants, respectively [abs (Log2FC > 1)]. (C,E,G) AF structures for PPP2R5D interaction with PPP2CA with the E198 at the interaction interface (C), for GNAI1 interaction with Ric8A with the I319 at the interaction interface, and (G) for FOXP1 showing interaction with DNA and FOXP4. (H)IP-Western blot of either non-transfected control (NTC), Strep-tagged FOXP1^WT^, FOXP1^R513C^, FOXP1^R513H^ or FOXP1^L327P^ in HEK293T cells show loss of FOXP4 interaction in cells transfected with FOXP1 mutants. (I) Quantification of IP-Western. Data is shown as mean ± SEM [n = 3, One way ANOVA, F = 16.34, p = 0.0009; Dunnett correction, adj. p = 0.0.0011 (vs. FOXP1^R513C^), adj p = 0.0018 (vs. FOXP1^R513H^), adj p = 0.0141 (vs. FOXP1^L327P^).

### FOXP1 R513H mutation alters the differentiation of deep layer cortical neurons in a human forebrain organoid model

We prioritized for further study the interaction changes of variants with the highest predicted pathogenicity based on Evolutionary model of Variant Effect (EVE), a deep learning model that predicts the likelihood that a human missense variant is pathogenic based on patterns of sequence variation across evolution^123^ (**Figure S7E**). All three FOXP1 mutants in ASD_mut_-PPI (FOXP1^L327P^, FOXP1^R513C^ and FOXP1^R513H^) had an EVE score of 1, indicating high likelihood of pathogenicity.

FOXP1 is a forkhead-box (FOX) family transcription factor that is important in the early development of multiple organ systems, including the brain^124–133^. The transcriptional activity of FOXP1 is regulated by homo- and heterodimerization with other FOX proteins^134,135^. Our ASD_mut_- PPI interaction study showed that all three FOXP1 mutants had significantly lower binding affinity to FOXP4 (**Figures 6D**, **7F**). An AF-predicted structure showed that the FOXP1 R513 residue is located in the Forkhead domain that interacts with DNA, while the L327 residue is located at the interface with FOXP4 (**Figures 7G**, **S7D**). Using immunoprecipitation followed by Western blot, we confirmed that all three FOXP1 mutations lost interaction with FOXP4 (**Figures 7 H,I, Table S7**).

To evaluate the effects of mutant FOXP1 in a model of human brain development, we generated clonally derived isogenic iPSC lines with either wildtype or heterozygous FOXP1^R513H/WT^ (**Figures 8A,B**, **S8A**). Consistent with data shown above, immunoprecipitation of FOXP1 from FOXP1^R513H/WT^ iPSC-derived neural progenitor cells showed decreased interaction with FOXP4, (**Figures 8C,D**, **S8B, Table S8**). We sought to determine whether the differential interactions of FOXP1^R513H^ affect neuronal development. We differentiated iPSCs into forebrain organoids (**Figure 8A**) and, to determine if the mutation impacts DNA binding of FOXP1, we performed CUT&Tag^136^ in FOXP1^R513H/WT^ and FOXP1^WT/WT^ forebrain organoids **(Figure S8C)**. Of 11,527 binding sites found in FOXP1^WT/WT^ organoids, only 6,939 were detected in FOXP1^R513H/WT^ organoids, suggesting that the heterozygous mutation leads to the loss of approximately 40% of binding events. Interestingly, organoids carrying the FOXP1^R513H^ mutant allele also exhibited 1,266 new binding sites not found in FOXP1^WT/WT^ organoids (**Figure 8E,F**).

**Figure 8.**
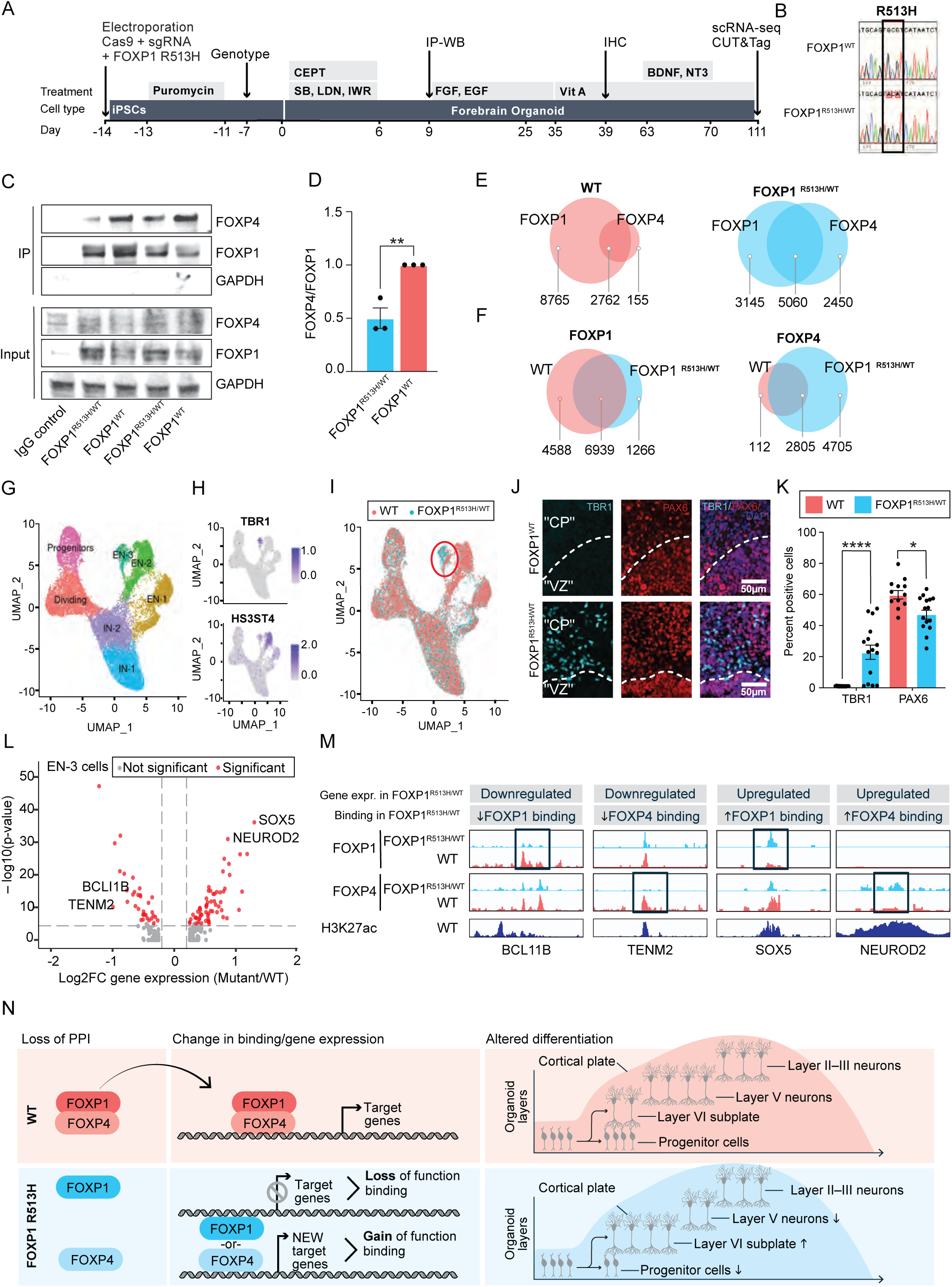
Impact of FOXP1 mutation on neuronal development and differentiation. (A) Genome editing timeline for generation of FOXP1 R513H/WT. A correctly targeted clone and WT control were expanded and used for forebrain organoid differentiation. (B) Sanger sequencing of clonal iPSC lines with wildtype (FOXP1^WT^) or heterozygous FOXP1 R513H mutation (FOXP1 ^R513H/WT^). Mutated bases have two peaks, one for the WT alleles (CGC) and one for the mutant alleles (ACA). (C) Co-immunoprecipitation of FOXP1 for WT and FOXP1^R513H/+^ NPCs. Representative western blot is shown for FOXP1, FOXP4, and GAPDH as a loading control. (D) Quantification of (C). Interaction of FOXP4 and FOXP1 is significantly reduced (*p* = 0.0065, t = 5.205, df = 4). (E) Overlap of FOXP1 and FOXP4 DNA binding regions in WT (Left panel) and FOXP1^R513H/WT^ (right panel). (F) Overlap of FOXP1 binding regions in WT and FOXP1^R513H/+^ (left) and FOXP4 binding regions in WT and FOXP1^R513H/+^ (right) (G) scRNA-seq UMAP from day 111 forebrain organoids. n = 2 technical replicates for both WT and FOXP1 ^R513H/+^. IN: inhibitory neurons; EN: excitatory neurons. (H) Feature plots of deep layer/subplate markers TBR1 and HS3ST4. (I) UMAP showing distribution of cells from either WT or FOXP1^R513H/WT^ forebrain organoids. FOXP1 R513H/WT cells are enriched in the EN-3 cluster, outlined in red. (J) Immunohistochemistry of day 39 forebrain organoids. TBR1 is expressed in subplate/deep layer neurons, and PAX6 is expressed in radial glia progenitors. VZ- and CP-like regions surrounding organoid rosette are shown. (K) Quantification of (J). n = 12 rosettes (WT), n = 15 rosettes (FOXP1 R513H/WT). Significance determined by Mann Whitney U tests with adjusted p-value calculated by Bonferroni correction for multiple comparisons. * = p < 0.05; **** = p < 0.0001. (L) Volcano plot of differentially expressed genes (DEGs) between WT and FOXP1 R513H/WT in the EN-3 cluster. Genes with positive log2fc value are enriched in FOXP1 R514H/WT cells, genes with negative log2fc value are enriched in WT cells. Significance determined by Wilcoxon Rank Sum test. Significant genes satisfy both log2FC cutoff = 0.2 and p-value cutoff = 1 × 10^−6^. (M) IGV browser tracks of CUT&Tag data for FOXP1 and FOXP4, in both WT and FOXP1 R513H/WT forebrain organoids. Two genes downregulated in FOXP1 R513H/+ (*BCL11B* and *TENM2*), and two genes upregulated in FOXP1 R513H/WT (*SOX5, NEUROD2*) are shown. These examples are shown to demonstrate the four patterns observed for changes in binding in the FOXP1 R513H/WT mutant: loss of FOXP1 binding (shown in *BCL11B*); loss of FOXP4 binding (shown in *TENM2*); gain of FOXP1 binding (shown in *SOX5*); and gain of FOXP4 binding (shown in *NEUROD2*). H3K27ac marks putative enhancers. (N) Schematic representation of mechanism by which FOXP1^R513H/WT^ alters differentiation by changing DNA binding of FOXP1 and FOXP4.

Given that the presence of a mutant allele of FOXP1 led to the loss of binding between FOXP1 and FOXP4, we also examined the landscape of FOXP4 DNA binding. Consistent with the heterodimerization model between FOXP1 and FOXP4, almost all FOXP4 binding sites (2,805 out of 2,817) overlapped with FOXP1 sites in the FOXP1^WT/WT^ organoids. By contrast, we found over 4,705 new binding sites for FOXP4 in FOXP1^R513H/WT^ not found in FOXP1^WT/WT^ organoids (**Figures 8E,F**). This suggests that changes in DNA binding of FOXP1 and FOXP4 occur in FOXP1^R513H/WT^ forebrain organoids, consistent with the proposed model of PPI disruption caused by the presence of the FOXP1^R513H^ allele.

To examine if the apparent reconfiguration of FOXP1 and FOXP4 binding underlies transcriptional differences in the developing forebrain, we performed single cell RNAseq (scRNA-seq) analysis on organoids and identified cell type-specific differentially expressed genes (**Figures 8G,H**, **S8D**-**F, Table S8**). Glutamatergic cortical neurons (EN-3) showed the most striking transcriptomic differences, with FOXP1^R513H/WT^ cells demonstrating relative upregulation of genes related to deep cortical layer neuron (TBR1) and subplate neuron identities (SOX5, NFIB, NFIA), and downregulation of genes associated with layer V glutamatergic neurons (TENM2, BCL11B, FOXP1) (**Figures 8I-L, S8G,H, Table S8**). Of note, genes with decreased FOXP1 binding in FOXP1^R513H/WT^ organoids and genes that were transcriptionally upregulated in FOXP1^R513H/WT^ EN-3 were both significantly enriched for hcASD255 (**Figure S8I**, **Table S8**; two-sided Fisher’s enrichment tests with Bonferroni correction: decreased FOXP1 binding, OR = 4.99, p.adj = 0.0038; EN-3 upregulated transcripts, OR = 4.93, p.adj = 0.012).

We observed that genes that were nominally differentially expressed in FOXP1^R513H/WT^ versus FOXP1^WT/WT^ EN-3 cells positively correlated with genes that were nominally differentially bound by FOXP1^R513H/WT^ versus FOXP1^WT/WT^ organoids (**Figure S8J**, Pearson R = 0.42, p = 0.18). While this correlation is not statistically significant, it suggests that genes with increased FOXP1 binding in FOXP1^R513H/WT^ organoids are relatively more highly expressed and vice versa. Of all differentially expressed genes in the EN-3 cluster (n = 239), 43% had unique FOXP1 binding sites that were gained in FOXP1^R513H/WT^ organoids, and not seen in FOXP1^WT^. Additionally, 32% of the differentially expressed genes had a gained FOXP4 binding site unique to FOXP1^R513H/WT^ organoids. 21% of the genes differentially expressed exhibited both loss of FOXP1 binding and gain of FOXP4 binding in FOXP1^R513H/WT^, suggesting a partial gain-of-function of FOXP4 in the FOXP1^R513H/WT^ organoids to compensate for loss of FOXP1 binding at these genes (**Table S8**).

Together, our results suggest that expression of the FOXP1^R513H^ allele during human forebrain differentiation alters gene regulatory relationships involving FOXP1 and FOXP4 transcription factors. While our scRNA-seq data did not suggest major differences in progenitor cell abundance, we detected a modest, but statistically significant decrease in the abundance of PAX6 positive cells, consistent with the proposed role for FOXP1 in human radial glia development^130^ and with findings from FOXP1 knockout organoids^133^. In addition, our findings implicate the FOXP1^R513H^ allele in altered differentiation of deep cortical layer and subplate neurons, which we confirmed by immunostaining for TBR1 (**Figures 8J,K**, **S8K**).

## Discussion

While the understanding of ASD genetic architecture and the identification of large-effect ASD risk genes has advanced in the past decade, the translation of this knowledge into molecular mechanisms underlying ASD, and the attendant identification of tractable treatment targets, has been challenging^7,10,137^. To date, efforts at identifying convergent biology from the growing list of hcASD risk genes have relied predominantly on gene expression analyses and traditional model systems studies. Resources characterizing the proteomic landscape of ASD have been strikingly limited. Prior to the present analyses, only approximately 30% of hcASD genes have been the subject of empirically generated, systemic PPIs investigation, including in resources such as BioPlex^48,51,52^. Compilation datasets like BioGRID^49,50^ and InWeb^51^ provide interaction data for a larger number of hcASD but suffer from challenges inherent in the aggregation of data from various experimental and analytic approaches. Here we confirm that the creation of large-scale empirically derived systematic data on PPI interactions related to large-effect ASD risk genes, and ASD-associated coding mutations therein, offer important insights into the biological mechanisms underlying ASD and offer new potential to generate and test highly specific mechanistic therapeutic hypotheses.

In this study, we adopted an integrated approach that combines human genetics, protein-protein interaction mapping and artificial intelligence (using AlphaFold-Multimer) to characterize the convergence of hcASD genes at the molecular and cellular level. We employed AP-MS in HEK293T cells to map PPIs for the vast majority of hcASD risk genes identified in the study by Satterstrom *et al*. 2020^6^, revealing 1,043 unique proteins connected to 100 hcASD through 1,881 interactions, of which 87% were not previously reported. Using AF for pairwise predictions of direct PPIs, we found that 113 of the 1,881 interactions (or 84 of 1,043 interactors) were predicted as direct physical interactions. Furthermore, AF additionally connected 377 interactors to hcASD via one or more intervening direct interactors.

We describe this dataset as “foundational” due to the scale of the investigation, the systematic approach to data generation, and the fact that it was derived in non-neuronal cells. We anticipate that additional studies currently underway in neural progenitor cells and iPSC-derived excitatory neurons will augment and further refine the understanding of ASD related PPIs as well as the consequences of ASD-associated mutations, capturing important context dependent data. We relied on HEK293T cells for this initial exploration as they have been very widely used to characterize the proteomic landscape of other human disorders^28,36,38,39,42^, facilitating multiple technical aspects of this study and allowing for key comparisons across datasets. Importantly, multiple lines of evidence point to the relevance of the PPI data generated in these cells for the human brain and for ASD. For example, genes encoding interactors are tightly co-expressed with hcASD genes, and the indicators identified here are highly enriched for independent sets of *bona fide* ASD risk genes, but not for a set of schizophrenia (SCZ) risk genes carrying similar rare, protein-disrupting coding mutations. Moreover, we demonstrate, via the in-depth study of FOXP1, that the consequences of disorder-associated mutation on binding to FOXP4, identified in HEK293T cells, are recapitulated in human forebrain organoids and result in a developmental phenotype consistent with a decade of analyses of convergence of human brain gene expression data^14,17^.

Notably, as the number of baits used to construct the ASD-PPI network increased, enrichment of ASD-associated genetic risk, including *de novo* variants and newly identified high-confidence ASD risk genes^3^, steadily increased, highlighting the potential of ASD-PPI in identifying and prioritizing additional ASD genes. Furthermore, we observed that ASD-PPI interactors tended to be more tolerant to damaging variants and to have lower selection coefficients than hcASD genes, potentially indicating that they contribute to ASD through recessive or polygenic inheritance. This suggests that PPI network-based gene discovery may be valuable for identifying additional ASD genes that may be missed by contemporary studies emphasizing haploinsufficient genes and rare heterozygous variants carrying large effect sizes. Furthermore, analyzing proteins in the context of their complexes can help interpret genomic data more effectively. For example, individual components of the PAF1 complex were not originally considered hcASD; however, our ASD-PPI map identified it as a central hub, and many of its components were just below the threshold used to define hcASD. Therefore, re-analysis of genomic data using modules or complexes will provide better insights into the underlying biology.

Significant overlap in interactors was found between ASD-PPI baits, suggesting that despite significant phenotypic and genetic heterogeneity across individuals with ASD, a smaller number of underlying molecular pathways may be driving pathology and that patients can be stratified accordingly, enabling more approaches for targeted treatment. Large-scale studies have repeatedly demonstrated that deleterious genetic variants in the same gene can contribute to different conditions, including ASD, SCZ, epilepsy, and other forms of neurodevelopmental delay (NDD) including intellectual disability (ID)^7^. Thus, there is an ongoing debate about the specificity of rare variant risk, especially with respect to ASD and NDD^138,1397^. In our study, we observed enrichment of NDD risk genes within the ASD-PPI network as well as no distinction between ASD-predominant and NDD-predominant hcASDs in the PPI network space, which could change as our power to distinguish ASD-specific versus shared genetic risk increases. Importantly, we do not observe enrichment of rare damaging variants identified in SCZ.

The ASD-PPI network also revealed substantial convergence at both the cellular and molecular pathway levels. Specifically, we found that co-expression of both ASD-PPI-AF and ASD-PPI-STRING networks were significantly increased in dividing neural progenitors, consistent with previous work in ASD^7^. We focused on DCAF7, which interacted with seven hcASD in the ASD-PPI network, including DYRK1A and KIAA0232. Knocking down *Dcaf7*, *Dyrk1a* or *Kiaa0232* in *Xenopus tropicalis* resulted in reduced telencephalon size, corroborating previous findings for DYRK1A while unveiling a novel function for DCAF7 and KIAA0232^17,101^. Our data suggest that all three genes may influence neurogenesis - consistent with longstanding observations in ASD - and underscore the potential functional overlap of DYRK1A/DCAF7 and other hcASD genes.

Our data also revealed that among the 1,070 high-confidence interactions identified in the mutant ASD PPI network (ASD_mut_-PPI), which encompassed 54 patient-derived *de novo* damaging missense mutations across 30 hcASD risk genes, 117 interactions had higher affinity for mutant proteins, while 136 had higher affinity for wildtype proteins. This mutant network suggests that damaging hcASD variants share common changes in molecular function, with some protein interactors being commonly gained or lost across multiple mutant hcASD. Using AF, we provide molecular-level details of the interaction interfaces of PPIs, enabling mechanistic hypotheses on gain- and loss-of-functions by ASD-related mutations. As we observed that mutations in the interface are more likely to result in loss rather than gain of interaction, even for low-scoring AF predictions, this structural knowledge from AF appears to extend beyond the 113 pairs that pass the high-confidence score threshold. AF was not initially built as a tool to detect PPIs but has been used as such effectively^81,82,140^. In our study, based on the increased performance of mean AF score (instead of the common practice of using the maximum from several models), we determined that consistency in AF results is important for identifying true interactors, in line with previous findings suggesting that aggregate conclusions from models with similar predictions were more likely to be accurate^141^. Our data suggest that directly incorporating model variance into a score may help identify true direct interactors, but there is some indication that low variance among models may be partly due to AF modeling an already solved structure. While our approach did not make use of template structures, we did find low variance among model scores more often when there are homologous structures in the training set for AF. The association of low-variance with good AF models may be the result of training data availability and not a highly confident *de novo* elucidation of structure, and warrants further investigation. Even so, AF remains a valuable tool for applying all available structural knowledge (via its training set) to the problem of predicting presence of interaction interfaces and their make-up.

Notably, our ASD_mut_-PPI highlighted the loss of interaction with FOXP4 for patient-derived mutations in the hcASD proteins FOXP1 and FOXP2. Given the central role for FOXP1 and FOXP2 in neurodevelopment, we hypothesized that these changes could impact neuronal differentiation. Prior studies have suggested that mutations in the forkhead box domain of FOXP1 result in loss of transcription factor activity^142,143^. We and others have found that overexpression of mutant FOXP1 R513H in HEK293T cells (data not shown) resulted in the formation of protein condensates containing FOXP1, FOXP2, or FOXP4, potentially changing DNA-binding ability and resulting in transcriptional dysregulation during neural development^143^. Interestingly, contrary to the anticipated loss-of-function^144^, the FOXP1 R513H mutation not only lost but also gained a large number of new DNA interactions, which correlated with changes in gene expression. Additionally, FOXP4, which failed to interact with FOXP1 R513H on a protein level, gained numerous DNA interactions in heterozygous FOXP1^R513H/WT^ organoids. Our findings suggest that the presence of a mutant form of FOXP1 leads to rewiring of DNA binding for several transcription factors, including FOXP2 and FOXP4. These findings emphasize the importance of investigating the functions of patient mutations.

Examining the impact of FOXP1^R513H/WT^ in forebrain organoids revealed several phenotypic characteristics, including reduced abundance of radial glia, mirroring the effects of null alleles^133^. Interestingly, no major differences in deep cortical layer neuron differentiation were observed in organoids with homozygous null mutations in FOXP1, while heterozygous FOXP1^R513H/WT^ organoids exhibited specific transcriptional dysregulation related to the differentiation of deep cortical layer neurons. This finding further underscores that patient-derived mutations do not merely replicate the effects of null mutations. The specific impact on the differentiation of glutamatergic neurons, with increased specification of TBR1+ deep cortical layer / subplate-like neurons, raises intriguing possibilities for the role of these mutations. The differential expression patterns of FOXP1, FOXP2 (broad expression) and FOXP4 (highly enriched in NPCs and early postmitotic neurons during deep cortical layer neurogenesis)^62,145^ in the developing forebrain and their potential impact on deep layer-like cell differentiation due to changes in DNA binding could be significant, but will require further confirmation using orthogonal models. Notably, similar alterations in deep cortical layer and subplate neuron differentiation have been reported in the study of idiopathic ASD patient-derived brain organoids^146^. It is important to note, however, that studies of ASD mutations using organoids are inherently limited by the virtue of their *in vitro* nature. While access to post mortem tissue from patients with FOXP1 R513H mutation is currently impractical and hampers direct validation of our findings, our results are consistent with previous systems biology work^14,17^ as well as observations in post mortem tissue from ASD patients that identified supernumerary neurons in deep cortical layers and the subplate^146,147^.

Previously, we have used PPI mapping to gain insight into a variety of different disease areas, including cancer^26,27^, infectious diseases^35,36,38–40^, neurodegeneration^34,148,149^ and heart disease^30,32^. In this study, we built a PPI map to study ASD, the largest such dataset focused on a neuropsychiatric disorder, which can be used as a resource for the identification of ASD biomarkers and therapeutic targets. Importantly, in this study, we describe a pipeline that combines the PPI data (wildtype and mutant) with AF, which allowed us to narrow in on specific interaction interfaces regulated by individual mutations that we analyzed in greater detail using a suite of tools including CRISPR-based genetics, stem cell differentiation and generation of organoids. This strategy (**Figure 7A**) represents a blue-print that can be used to interpret the genomic information from any genetically-defined disease or disorder, efforts that will rapidly uncover not only novel targets but inform specific therapeutic strategies.

## Acknowledgements

This research was funded by grants from the National Institutes of Health to N.J.K, A.J.W, M.W.S, T.I. and T.J.N (U01MH115747), A.J.W. and M.W.S (U01MH116487), N.J.K and T.I. (1OT2OD032742, U54CA274502), and T.J.N (R01MH128364, R01NS123263, SF810018). This study was also supported by the Weill Institute for Neurosciences (Startup Funding to A.J.W.), the Quantitative Biosciences Institute (QBI) at UCSF, the Overlook International Foundation (to M.W.S. and A.J.W.), gifts from Schmidt Futures and the William K. Bowes Jr Foundation (to T.J.N), the Sergey Brin Family Foundation (to M.W.S) and the Sorensen Foundation Career Award in Child & Adolescent Psychiatry (to B.W.). T.J.N. is a New York Stem Cell Foundation Robertson Neuroscience Investigator; H.R.W. is a Chan Zuckerberg Biohub - San Francisco Investigator. The authors would like to thank Tami Tolpa and Alex Olmsted for graphic design support for figures, and all members of the Psychiatric Cell Map Initiative (PCMI; U01MH115747) for their invaluable discussions and support.

## Author contributions

Conceptualization: T.I., P.B., H.R.W., R.H., K.O., T.J.N., M.W.S., A.J.W. and N.J.K.; Methodology: B.W., R.V., Z.Z.C.N., K.M.H., B.J.P., A.B., M.B., U.C., J.A.K., R.T., D.L.S., T.J.N., R.H. and A.J.W.; Software: B.W., Z.Z.C.N., B.J.P., A.B., M.B., Y.Z., J.M., K.Z.G., M.C., M.G., D.F.B., S.A., P.P., and D.P.; Validation: B.W., R.V. and B.J.P.; Formal Analysis: B.W., R.V., Z.Z.C.N., K.M.H., B.J.P., M.B., M.C.L., Y.Z., J.M., U.C., J.A.K., M.C., M.G., S.A., S.L., S.W. and T.R.; Investigation: B.W., R.V., Y.Z., Z.Z.C.N., K.M.H., M.B., J.X., N.S., M.C.L., U.C., J.A.K., N.C., P.K., S.D., V.D., S.G., M.M., E.B., M.S., T.B., R.K., J.A. and T.R.; Data Curation: B.W., Z.Z.C.N., B.J.P., A.B., M.B., K.Z.G., U.C., M.C., M.G., S.A. and T.R.; Writing - Original Draft: B.W., R.V., Z.Z.C.N., K.M.H., B.J.P., A.B., M.E., K.O. and T.J.N.; Writing - Review & Editing: B.W., R.V., M.E., S.C., K.O., M.W.S., A.J.W and N.J.K.; Visualization: B.W., R.V., Z.Z.C.N., K.M.H., B.J.P., A.B., M.E., M.B., M.C.L., U.C., J.A.K., S.A. and D.P.; Supervision: J.A.K., T.B., S.F., S.B., N.M., T.I., P.B., H.R.W., K.O., T.J.N., R.H., M.W.S., A.J.W. and N.J.K.; Project Administration: A.J.W. and N.J.K.; Funding Acquisition: T.I.,T.J.N., M.W.S., A.J.W. and N.J.K.

## Supplemental Figures and Tables

**Supplemental Figure 1. Defining ASD-PPI high-confidence interactions, related to Figure 1**.

(A) Workflow to determine SAINTexpress^46^ and CompPASS^151^ scoring cutoffs by maximizing sensitivity and specificity metrics for recovery of gold standard interactions from public databases^48–52^.

(B) Precision (red) and recall (blue) analysis of known interactions using different combinations of SAINTexpress and CompPASS cutoffs. The dotted lines show the precision and recall values at the selected scoring cutoffs for ASD-PPI.

(C) The kernel density plot displaying the scoring distribution of known interactions observed in the unfiltered PPI dataset. The dotted lines show the scoring cutoffs selected for ASD-PPI and highlight the high density of known interactions at these cutoffs.

(D) Overlap between the interactors identified for 13 hcASD proteins using endogenous IP-MS in iPSC-derived excitatory neurons (iENs)^23^ and the combined interactors for the matched set of 13 hcASD baits (ASD-PPI subset) or all ASD-PPI. Odds ratios were calculated using Fisher’s exact test, with p-values adjusted for 2 tests (Bonferroni); the gene universe was restricted to the n = 2,552 proteins that are expressed in both HEK293T and iNs.

(E) Geneset-level burden tests for enrichment of *de novo* damaging variants in ASD probands compared with unaffected siblings from the Simons Simplex Collection^6^. Genesets include the combined network generated for 13 hcASD proteins using endogenous IP-MS in iPSC-derived excitatory neurons (iNs)^23^ and the matching network for 13 hcASD baits from ASD-PPI (ASD-PPI subnetwork). Statistical significance was calculated using Fisher’s exact test (one sided, greater), and p-values were adjusted for 2 tests (Bonferroni).

(F) Overlap between the interactors identified for 7 ASD risk proteins (3 of which were hcASD) using proximity labeling with baits overexpressed in HEK293T cells (‘HEK-PPI’)^24^ and the ASD-PPI interactors. The p-value was calculated using Fisher’s exact test (one sided, greater).

(G) Overlap between the interactors identified for 41 ASD risk proteins (17 of which were hcASD) using proximity labeling with baits overexpressed in mouse cortical neurons (‘Mouse-PPI’)^24^ and the ASD-PPI interactors. The p-value was calculated using Fisher’s exact test (one sided, greater).

(H) Geneset level burden tests for (F) HEK-PPI

(I) Geneset level burden tests for (G) Mouse-PPI.

n.s. not significant, ∗p < 0.05, ∗∗∗p < 0.001

**Supplemental Figure 2. Interactors are expressed in contexts associated with ASD and enriched for ASD genetic risk, related to Figure 2.**

(A) The median geneset expression percentile for adult brain tissue samples from 13 brain regions in GTEx^54^ for three genesets - baits, interactors (excluding hcASD, ‘Interactors – hcASD)’), and all other proteins expressed in HEK293T cells (‘Other’). Differences in the median expression percentile between genesets was assessed by T-test; p-values were adjusted for multiple hypothesis testing (Bonferroni, 3 tests).

(B) The relative expression levels of hcASD compared to interactors (-hcASD) in prenatal versus postnatal brain samples from BrainSpan^150^. The relative expression within each brain sample was quantified by the difference between the median geneset rank of observed versus median of 100,000 permuted genesets. Differences in relative expression between prenatal and postnatal samples were assessed by T-tests.

(C) The relative expression levels of interactors (-hcASD) in GTEx brain samples across 13 brain regions^54^ compared to that of permuted genesets selected from the HEK293T proteome. The relative expression within each brain region was quantified by the difference between the median geneset rank of observed versus median of 100,000 permuted genesets. Red dashed line shows nominally significant p-value of 0.05, and orange dashed line shows significance adjusted for 13 tests (Bonferroni).

(D) The distribution of evolutionary constraint metrics (pLI, misZ, shet, and synZ) for baits, interactors (-hcASD), and all other proteins expressed in HEK293T cells, The difference in score distribution between different genesets was assessed by T-test, with p-values corrected for 3 tests (Bonferroni).

(E) ASD-PPI interactors are enriched for ASD and DD risk genes but not SCZ risk genes. The 8 sets of ASD-associated risk genes were obtained from two recent WES studies (Fu *et al*. 2022^3^: n = 255 genes; Zhou *et al.* 2022^5^: n = 72 genes; and SFARI^152^: n = 230 syndromic genes, n = 92 syndromic & category 1 genes, n = 206 category 1, n = 219 category 2, n = 514 category 3, and n = 1020 SFARI genes (‘SFARI all’)). SCZ-associated risk genes were obtained from Singh *et al.* 2022^60^: n = 34 genes. Risk genes associated with developmental disorders (DD) were obtained from Kaplanis *et al.* 2020^153^: n = 285 genes. Enrichment was calculated using Fisher’s exact test (one sided, greater), with p-values adjusted for 10 tests (Bonferroni); the gene universe was restricted to the HEK293T proteome.

∗p < 0.05, ∗∗p < 0.01, ∗∗∗p < 0.001

**Supplemental Figure 3. ASD-PPI convergence metrics and comparison with other PPI data, related to Figure 3.**

(A) Distribution of the number of hcASD interactions for ASD-PPI interactors.

(B) The distribution of number of interactors in ASD-PPI for all baits (green), baits that were more frequently mutated in ASD (orange, ASD-predominant; ASD_P_) and baits that were more frequently mutated in NDD (blue, ASD_NDD_)^6^. Shaded gray regions indicate baits with extremely large numbers of interactors (> 45).

(C) Interactor overlap for all ASD-PPI bait pairs, ASD_P_ bait pairs, ASD_NDD_ bait pairs, and ASD_P_ – ASD_NDD_ bait pairs as measured as proportion of significant overlap between the interactor sets for all pairs of baits. Significant overlap is measured by p-values from hypergeometric tests, and the portion with p < 0.05 is shown.

(D) Genome-wide rank and FDR for the five proteins of the PAF complex scored for their genetic association with ASD by Fu *et al.* 2022 (green) or Satterstrom *et al.* 2020 (orange). Ranks are based on FDR, with lowest FDR receiving the lowest rank, so points in the top right corner are the most associated with ASD. Curved lines are the FDR and rank association for all genes measured in each study. Dashed horizontal line is at FDR = 0.1, the threshold applied by Satterstrom *et al.* 2020.

(E) Genome-wide rank for ASD-int (blue) scored for their genetic association with ASD by Fu *et al.* 2022 (y-axis) and Satterstrom *et al.* 2020 (x-axis). Members of the PAF complex are highlighted in red. Gray dots are for non-ASD-int genes. Dots are shown for proteins that rank in the top 8000 for both studies. Outside of this range, the linear relationship between gene ranks in the two genetics studies decreases. The blue horizontal and vertical violin plots and line segments show the distribution of all ASD-int, even those with rank less than 8000 in one or both studies.

**Supplemental Figure 4. Comparison of different metrics for thresholding AF structure predictions, related to Figure 4.**

(A) Evaluation of three scoring metrics, confidence (red), ipTM (green) and pDOCkQ (blue), for AF dimer models for their ability to discriminate the bait-int from bait-random sets of proteins. Confidence is 0.8⋅ipTM + 0.2⋅pTM. Scores were evaluated by the ratio that passed threshold in bait-int versus bait-random sets (y-axis) across a sliding threshold and compared based on the portion of bait-int they recall (x-axis). Greater areas under the curve reflect better performance. For thresholds that exclude all bait-random pairs, no value is shown as the ratio is undefined. The upper panel shows the use of the maximum score across all AF models and the lower panel shows the use of the arithmetic mean.

(B) A comparison of different summary statistics for ipTM scores and their ability to discriminate the bait-int from bait-random sets. Summary statistics are evaluated by the ratio that passes threshold in the bait-int versus bait-random (y-axis) across a sliding threshold and compared based on the number of bait-int dimers they recall (x-axis). Mean and max are as in (A). Min is minimum, assigning the lowest score of any AF to the dimer, and adjMean is the arithmetic mean adjusted by subtracting two standard deviations for the model. At stringent thresholds, where bait-int counts are below 100, min and adjMean perform similarly and better than mean, but they become more equivalent at bait-int counts greater than 100. Dotted vertical line at count = 113 corresponds to the mean ipTM threshold of 0.5.

(C) Estimates of false discovery rate (FDR) of direct interactors in the bait-int set when using different thresholds of mean ipTM as the classifier. Estimates are made by assuming that all bait-random pairs are true non-interactors or non-direct interactors, and thus the bait-random pass rate is an estimate of false positive rate (1-specificity). We estimate the FDR, defined as (non-direct-interactors above threshold)/(all pairs above threshold), separately assuming that different portions (10, 25, 50%) of the bait-int sets are true direct interactors. The model used is that the bait-int dimers passing any threshold are composed of: (*total number false in bait-int*) × (*bait-random pass rate at threshold*) + (*true positives*). FDR rates are shown (colored horizontal lines and numbers) corresponding to the mean ipTM threshold of 0.5 (black vertical line).

(D) Scatterplots showing relation between mean ipTM and standard deviation ipTM for all dimer pairs compared by availability of prior knowledge. A dimer pair gets classified as known instead of novel based on presence in a CORUM complex, a STRING-DB combined score greater than 0.5, presence in HumanNet gold standard set, or having a HumanNet score greater than 2.0. A dimer pair is classified as “homologous pdb” based on exhaustive sequence searches of PDB sequence records by BLAST. If both proteins in a dimer pair have detectable similarity (BLAST expect < 0.001) to different chains in a single PDB record, the pair is labeled “in pdb”. Numbers in each plot region are the number of AF models above and below the threshold of mean ipTM = 0.5.

(E) Amount of DYRK1A, FAM53C, and DCAF7 detected in AP-MS with Strep-tagged DYRK1A or DYRK1A^Δ^^80–100^. Significant interactions (SAINT 1-BFDR ≳ 0.95) are colored in red.

**Supplemental Figure 5. DCAF7, DYRK1A and KIAA0232 overlap functionally, related to Figure 5.**

(A) Interactors shared between DCAF7 AP-MS, DYRK1A AP-MS and sequential AP-MS for DCAF7 and DYRK1A. Red bars denote hcASD255^3^ while gray bars denote interactors other than hcASD255.

(B) Overlap between hcASD255 genes and interactors of DCAF7-DYRK1A sequential IP reveal 9 shared hcASD genes.

(C) ASD-PPI interactor enrichment for three sets of proteins associated with structures important for spindle organization (centriolar satellites^103^ and centrosomes^104^, two-sided Fisher’s exact tests with Bonferroni correction).

(D) Representative images showing immunostaining of DYRK1A (green), DCAF7 (blue), KIAA0232 (magenta) show co-localization of the three proteins with alpha tubulin on HEK293T cell mitotic spindles. Scale bar = 5 μM.

(E) Representative images showing increased cell death in cortical neurons with *DCAF7* knockdown transduced with the red GEDI (RGEDI) biosensor at differentiation day 13. Images show morphology (GFP, left), cell death (RFP, middle) and overlay (right). Arrows point to a dying or dead cell (scale bar = 100 µm).

(F) Fraction of dead cells (RFP+GFP+/GFP+) in NPCs with *DCAF7* knockdown versus control on differentiation days 7 to 11. (n = 3, p < 0.00001).

(G) Representative image of Ki67 staining in NPCs containing control (top) or *DCAF7* gRNAs (n = 4 per condition) (scale bar = 100 µm). Images show nuclei (Hoechst, left), dividing cells (Ki-67, middle) and overlay (right).

(H) Differences in Ki67 staining in NPCs with sgRNAs targeting *DYRK1A, DCAF7,* or *KIAA0232*.

**Supplemental Figure 6. Validation and metrics of convergence for ASDmut-PPI differential interactions, related to Figure 6.**

(A) hcASD255 enrichment in interactors with decreased (‘Down’) or increased (‘Up’) binding to mutant versus WT hcASD baits in ASD_mut_-PPI was assessed by two-sided Fisher’s exact tests with Bonferroni correction.

(B) Convergent changes in ASD_mut_-PPI. X-axis labels indicate individual interactors and in parentheses the number of significant interaction changes involving the interactor in ASD_mut_-PPI y-axis labels indicate the parent hcASD baits and in parentheses the number of variants in ASDmut-PPI. Point size reflects the number of hcASD variant baits with significantly changed binding to the indicated interactor and color reflects the proportion of hcASD baits variants with significantly changed binding to the indicated interactor. For example: there were four changed interactions with the interactor FOXP4 (highlighted in red text): three from FOXP1 variants (100%) and one from FOXP2 variants (100%).

(C) Differential interaction of FOXP2-R570C and FOXP1-R513C, R513H and L327P with FOXP4.

(D) Representative Western blot evaluating immunoprecipitation of Strep-tagged FOXP2-WT or FOXP2-R570C with FOXP4.

**Supplemental Figure 7. Mapped mutations onto AF models relate to functional changes based on their presence in interaction interfaces, related to Figure 7.**

(A) Distribution of mutant bait-int pairs from the ASD_mut_-PPI network based on distance of the mutated residue to the interaction interface (y axis). The mutant bait-int interactions are classified as down (interactions that are weakened), no change, and up (interactions that are enhanced) in presence of a mutation.

(B-D) Confidence (pLDDT) per residue plotted for PPP2R5D-PPP4C (B), GNAI1-RIC8A (C), and FOXP1-FOXP4 (D) with pLDDT scores ranging from 0 (red) to 100 (blue).

(E) EVE scores^123^ of hcASD missense mutations in ASD_mut_-PPI (higher scores reflect higher predicted pathogenicity).

Supplementary Figure 8. FOXP1^R513H^ differentially regulates neuronal gene expression, related to Figure 8.

(A) Confirmation of forebrain identity by IHC staining for FOXG1 (green) in both FOXP1 R513H/WT and WT organoids.

(B) Additional quantifications from IP-WB from Figure 8C, showing no change in FOXP4 (left) or FOXP1 (right) expression between FOXP1^R513H/WT^ and WT NPCs. Abbreviations: IP-Immunoprecipitation

(C) CUT&Tag estimated library size for replicates 1 and 2, for both FOXP1 R513H/WT and WT organoids at D111. Libraries for FOXP1 binding are shown in orange, and libraries for FOXP4 are shown in green.

(D) Gene and UMI count, and mitochondrial percentage for FOXP1 R513H/WT and WT organoid scRNA-seq at D111.

(E) Gene and UMI count, and mitochondrial percentage across clusters.

(F) Cell-type specific genes used to annotate scRNA-seq clusters, shown in Figure 8G.

(G) Cluster proportion graph showing distribution of cells from either FOXP1 R513H/WT or WT forebrain organoids in each cluster. FOXP1 R513H/WT cells are enriched in EN-3.

(H) Volcano plot from Figure 8L with additional genes annotated.

(I) Enrichment of hcASD255 (Fu *et al.* 2022) in FOXP1 differentially bound genes (DB, left panel) and cell-type specific differentially expressed genes (DEGs, middle panel: upregulated in FOXP1^R413H/WT^, right panel: upregulated in FOXP1^R514H/WT^). Two-sided Fisher’s exact tests, p values adjusted for multiple hypothesis testing (Bonferroni correction, 16 tests).

(J) Scatter plot showing correlation of CUT&Tag FOXP1 differential peaks between FOXP1 R513H/WT and WT (x-axis) and scRNA-seq DEGs between FOXP1 R513H/WT and WT in EN-3 cluster (y-axis).

(K) Full images that were cropped around rosettes and quantified for Figure 8J-K. While minimal TBR1 expression (was seen in the cropped images for the WT as compared to FOXP1^R513H/WT^, full images show that TBR1 is expressed in these organoids.

**Supplemental Table 1. Related to Figure 1 and S1**

**Supplemental Table 2. Related to Figure 2 and S2**

**Supplemental Table 3. Related to Figure 3 and S3**

**Supplemental Table 4. Related to Figure 4 and S4**

**Supplemental Table 5. Related to Figure 5 and S5**

**Supplemental Table 6. Related to Figure 6 and S6**

**Supplemental Table 7. Related to Figure 7 and S7**

**Supplemental Table 8. Related to Figure 8 and S8**

## Methods

### Cloning hcASD risk genes

The coding sequence of each hcASD risk gene was cloned into a pcDNA4 plasmid with either N- or C-terminal 2xStrep tags, which encode the bait proteins for the AP-MS study **(Table S1)**. The terminus position of the tags was determined so that the tag will not interfere with the protein function based on the prior reported plasmids that have been used in functional studies. The isoforms for the hcASD risk gene were chosen based on high brain expression levels as well as the high frequency of mutations observed in ASD using Clonotator (http://ec2-52-91-98-53.compute-1.amazonaws.com/login/). All constructs were sequence validated.

### Cell culture

HEK293T cells were cultured in Dulbecco’s modified Eagle’s medium (DMEM; Corning) supplemented with 10% fetal bovine serum (FBS; Gibco, Life Technologies) and 1% penicillin– streptomycin (Corning). All cells were maintained in a humidified incubator at 37 °C with 5% CO_2_. hiPSCs (WTC11, 13234) were cultured on Matrigel (Corning #354230) coated plates in StemFlex Medium (Gibco #A3349401) with 100ug/ml Primocin (Invivogen #ANTPM1). Cells were passaged 1:10 every 3 days using ReLeSR (StemCell Technologies #05873). Cells were maintained in a humidified incubator at 37°C with 5% CO_2_.

### Transfection

Each transfection (102 baits, one GFP control and one empty vector control) was carried out in a 15cm dish with 10 million HEK293T cells (70-80% confluency), with three biological replicates per bait. Transfections were split in 15 batches, with three biological replicates of GFP and empty vector controls included in each batch. For each transfection, 15 μg of Strep-tagged plasmids was combined with PolyJet Transfection Reagent (SignaGen Laboratories) at a 1:3 μg:μl ratio of plasmid:transfection reagent, incubated at room temperature for 10 mins, and added dropwise to HEK293T cells. About 48h post transfection, cells were resuspended at room temperature using 10 ml Dulbecco’s phosphate-buffered saline without calcium and magnesium (DPBS) supplemented with 10 mM EDTA, followed by centrifugation at 200g, 4 °C for 5 min. Cell pellets were frozen on dry ice and stored at −80 °C.

### Affinity purification

The cell pellets were thawed on ice and then lysed with 1 ml ice-cold lysis buffer (IP buffer (50 mM Tris-HCl, pH 7.4, 150 mM NaCl, 1 mM EDTA, 0.5% Nonidet P40 substitute (NP40; Fluka Analytical), cOmplete mini EDTA-free protease and PhosSTOP phosphatase inhibitor cocktails (Roche)). Samples were then flash-frozen on dry ice for about 10 min and partially thawed at 37°C for 30-45s before incubation for 30 min at 4 °C on a tube rotator. Lysates were centrifuged at 13,000g, 4 °C for 15 min to clarify lysate and pellet debris. A 50 μl lysate was reserved at this point for future experiments such as Western blot. The remaining lysates either underwent automated affinity purification on the KingFisher Flex Purification System (Thermo Scientific), which was first equilibrated to 4°C in the cold room or were processed for immunoprecipitation for validation. First, MagStrep ‘type3’ beads (30 μl; IBA Lifesciences) were equilibrated twice with 1 ml wash buffer (IP buffer supplemented with 0.05% NP40) and incubated with 0.95 ml lysate for 2 h. Beads were washed 3 times with 1 ml wash buffer and once with 1 ml IP buffer, then resuspended in 50 μl denaturation–reduction buffer (2 M urea, 50 mM Tris-HCl pH 8.0, 1 mM DTT) and 50 μl 1× buffer BXT (IBA Lifesciences) and dispensed into a single 96-well KingFisher microtitre plate. Purified proteins were eluted at room temperature for 30 min with constant shaking at 1,100 rpm on a ThermoMixer C incubator.

For immunoprecipitation, the lysate was incubated with MagStrep ‘type3’ beads (30 μl; IBA Lifesciences) for 2 h at 4C, followed by washing with 1ml wash buffer (IP buffer supplemented with 0.05% NP40) three times. The resulting washed beads were mixed with 6X Laemmli SDS Sample buffer (Boston Bioproducts, Cat#BP-111R), heated at 95C for 5 mins and loaded onto SDS-PAGE for Western blot.

### On-bead digestion

Bead-bound proteins were denatured and reduced at 37 °C for 30 min, brought back to room temperature, alkylated in the dark with 3 mM iodoacetamide for 45 min and quenched with 3 mM DTT for 10 min. To offset evaporation, 15 μl 50 mM Tris-HCl, pH 8.0 were added and proteins were digested with 1.5 μl trypsin (0.5 μg/μl; Promega) with constant shaking at 1,100 rpm and incubation at 37 °C on a ThermoMixer C incubator for 4 h, and again for 1–2 h with 0.5 μl additional trypsin. Resulting peptides were combined with 50 μl 50 mM Tris-HCl, pH 8.0 to rinse beads and then, acidified with trifluoroacetic acid (0.5% final, pH < 2.0). Desalting was conducted in a BioPureSPE Mini 96-Well Plate (20 mg PROTO 300 C18; The Nest Group) according to standard protocols.

### MS data acquisition and analysis

To prepare for mass spectrometry, samples were resuspended in 4% formic acid, 2% acetonitrile solution, and separated by a reversed-phase gradient over a Nanoflow C18 column (Dr Maisch).

Each sample was directly injected via an Easy-nLC 1200 (Thermo Fisher Scientific) into a Q-Exactive Plus mass spectrometer (Thermo Fisher Scientific) and analyzed with a 75 min acquisition, with all MS1 and MS2 spectra collected in the orbitrap; data were acquired using the Thermo software Xcalibur (4.2.47) and Tune (2.11 QF1 Build 3006). For all acquisitions, QCloud was used to control instrument longitudinal performance. All proteomic data were searched against the human proteome (UP000005640_9606, downloaded in October 2021) using the default settings for MaxQuant software (version 1.6.12.0) (Cox & Mann, 2008) with match-between-runs (MBR) feature turned on. Briefly, the MBR algorithm annotates unidentified peaks by assessing and comparing the retention times of the identified peaks in an MS1 spectrum. Detected peptides and proteins were filtered to 1% false-discovery rate in MaxQuant, and identified proteins were then subjected to protein–protein interaction scoring with both SAINTexpress (v.3.6.3)^46^ and CompPASS (version 0.0.0.9000)^47,48^.

### Removal of carryover effect and identification of high-confidence interactors

A crucial step in our AP-MS study is the probabilistic scoring of all quantified proteins in the dataset to identify high-confidence interaction proteins (HCIP) of the ASD risk genes. To do this, the scoring outputs from existing computational tools such as CompPASS (Comparative Proteomic Analysis Software Suite)^47,48^ and SAINTexpress^46^ were systematically combined. These two scoring algorithms are widely used in the proteomics community to score the quality of protein-protein interactions as distinct from background.

Input files - bait, prey and interaction files - for SAINTexpress were made using artmsEvidenceToSaintExpress function in an open-sourced R package, artMS^154^. The interaction input file for SAINTexpress was reformatted to be used as a CompPASS input file. As CompPASS requires all baits to have the same number of preys, we defined spectral counts for the union of all identified proteins across all AP-MS experiments for each bait; if the bait-prey interaction was not detected, its spectral count was assigned to be zero. This bait-prey spectral count matrix was used to search for carryover effect, which was defined as proteins with a continuous decrease in spectral counts in replicates where such proteins were used as baits in the previous sample injections. The resulting bait-prey spectral count data table after carryover removal was reformatted and used as the input file for SAINTexpress and CompPASS.

To determine the scoring cutoffs that capture the highest number of true interactions, a gold standard set of protein-protein interactions was manually defined. Gold standard interactions were extracted from publicly available large-scale PPI databases, Hein *et al.*^52^, InWeb^51^, CORUM (Core Corum Complexes, Corum 3.0)^50^, BioPlex^48^, and BioGrid^49,92^, with some additional filtering steps for BioGrid and InWeb. As BioGrid contains interactions from various sources, including both experimental and predicted, the dataset was filtered to only interactions that were attained using another experimental method in addition to the AP-MS method. Similarly, InWeb databases were filtered for interactions that have a high-confidence score of > 0.95 and are identified through an experimental method.

SAINTexpress (version 3.6.3) was run batch-wise, each batch with its own empty-vector and GFP replicates as controls. The outputs were then concatenated to a single SAINTexpress output. CompPASS was run using the R package, cRomppass (https://github.com/dnusinow/cRomppass). The scoring outputs from SAINTexpress and CompPASS were merged for each interaction. The SAINTexpress Bayesian False Discovery Rate (BFDR) and CompPASS WD score (rank_WD, ranked from 0 to 1 across the dataset) were used as prediction confidence scores. Using the ranked CompPASS WD score (rank_WD) as a predictive value and interactions found in the previously described manually curated gold standard PPI database as true positives, the optimal rank_WD score cutoff for each SAINTexpress BFDR increment was determined by calculating Youden’s index. The best performing composite score of SAINTexpress and CompPASS (out of 78 composite scores) was determined by calculating precision, recall and F1 scores.

### Selection and cloning of Missense Mutations for AP-MS studies

Satterstrom *et al.* considered combined deleterious effects of both protein truncating variants (PTVs) and missense variants to generate a list of genes that are highly associated with ASD in family studies. To quantitatively measure functional effects of these variants, they calculated “probability of loss-of-function intolerance” (pLI) scores for PTVs and an integrated score called MPC for missense mutations. There are three tiers for MPC score (≥2, 1-2, 0-1) which are in the order of decreasing functional impact. When evaluating the MPC scores of *de novo* missense variants, Satterstrom *et al.* found that MPC ≥ 2 are 2.2-fold more enriched in cases. To study protein interaction changes using AP-MS, we focused on these missense mutations in hcASD risk genes with MPC≥ 2. This resulted in 87 *de novo* missense mutations in 43 hcASD risk genes. The missense mutations were introduced in the wildtype version of the pcDNA4 construct using Q5 site-directed mutagenesis and used in the usual AP-MS pipeline described above. After checking for expression of the variants in HEK293T cells and further quality control, 33 variants across 13 hcASD were removed.

### PPI scoring for Missense Mutations in AP-MS studies

Following the identification and quantification of proteins using MaxQuant, high confidence interacting proteins (hcIP) were identified by running both SAINTexpress and CompPASS after carryover effects were removed. ASD-PPI refers to the AP-MS dataset generated from 102 hcASD risk genes. ASD_mut_-PPI refers to AP-MS dataset generated from 87 *de novo* missense mutations in 43 hcASD risk genes, along with repeated AP-MS of their respective wildtype constructs as controls for differential interaction analysis, and the usual empty-vector and GFP controls. SAINTexpress was run batch-wise, using empty vector and GFP constructs as controls. To have better specificity in identifying hcIPs using CompPASS, a larger input dataset was created by combining the ASD-PPI CompPASS input with ASD_mut_-PPI input. CompPASS was then run using the CompPASS input for ASD-PPI WT network as the background stats table. hcIPs were identified using the scoring cutoffs previously optimized from the wildtype ASD-PPI network: SAINTexpress score (1-BFDR) ≥ 0.95 and CompPASS score (rank_WD) ≥ 0.971. We removed new hcIPs identified for ASD-PPI; these were likely a result of rerunning CompPASS on the ASD-PPI input together with ASD_mut_-PPI creating a different background model. We retained only hcIPs from the mutant AP-MS experiments (including from their WT quantitative controls) to build the ASD_mut-PPI interactome. As a first step for quantifying differential interactions, we normalized interactor peptide intensity values to baits levels to account for variation in bait expression. This bait normalization was completed within each AP-MS run using a Tukey Median Polish normalization method on all bait peptides to calculate and remove the bait-variation across all peptides in the run. Following normalization, we quantified the changes observed in interactors between wildtype and mutant baits, using the R package MSstats (Choi *et al.* 2014). Using the default parameters, except to disable further normalization, we used the functions *dataProcess* and *groupComparison* on the normalized intensity values to run differential analysis between wildtype and mutant groups. Differential interactors were then identified using thresholds *p* < 0.05 and Log2FC ≥ 1.

### ASD-PPI protein expression in HEK293T cells

We defined the HEK293T proteome to be the n=11,133 proteins with detected protein expression in a published global quantitative mass spectrometry dataset^155^. We defined the protein expression level to be the average intensity based absolute quantification (iBAQ) scores for the two HEK293T experimental replicates reported in Supplementary Table 7. For subsequent analyses, we defined HEK293T proteome genes to be the union of proteins with non-zero iBAQ scores in Bekker-Jensen *et al.* 2017 and ASD-PPI interactors.

### Network Layout of Baits and interactors

The total network showing AP-MS detected interactions between 100 hcASD baits and their connections to 1043 prey plus connections to other baits was laid out using t-SNE (R package Rtsne) on a weighted combination of distance based on shared Gene Ontology annotations (weight = 4) and distance based on shared AP-MS connections (weight = 1). Binary distances, equivalent to 1-Jaccard similarity, were used for both distances. For GO distances, to create a limited set of 141 GO terms that maximally cover and describe the set of 1043 interactors, the GO terms were limited to those significantly (FDR < 0.05) enriched in the full set of 1043 interactors and further limited to only those terms that were the top-enriched term (by p value) per at least one interactor. Enrichment was computed and scored by the function enricher in the R clusterProfiler package using GO annotations from all GO in org.Hs.eg.db (R package) and limiting GO terms to those with at least 20 and at most 500 genes. Interactors were assigned to terms according to their GO annotations. Baits were assigned to these GO terms based on significant (p value < 0.05) enrichment in their set of interactors using the set of 1043 as background (universe), or when no significant enrichment existed, simply the one term (of 141) with highest number of interactors per bait. For network distances, a symmetric adjacency matrix was built where each bait and interactor were connected to itself and to each protein that it connects to by a direct AP-MS interaction, and binary distances were computed on rows of the matrix. After layout by t-SNE (default settings except is_distance = TRUE and theta = 0.0), protein two dimensional coordinates were adjusted to avoid overlapping nodes using an iterative, repulsive algorithm. The final image was formatted and drawn using the R package ggplot2, mostly with functions geom_point and geom_segment. Coloring of interactors was based on a subset of the GO terms chosen to best cover the 1043 interactors in a non-redundant fashion with a small number of terms. For this term selection, an additional GO enrichment run was completed, allowing terms to have at most 2000 genes, and including additional terms that described interactors previously undescribed by the 141 terms. Clustering rows and columns of the annotation matrix, 1043 genes × 186 terms, was performed to aid in this manual selection. In the network view, where a gene is annotated to more than one of the chosen terms, it is assigned the color whose median 2D location on the plot it is closest to.

### Comparing convergence in prey sets

Convergence in an AP-MS network, the interaction of multiple baits with the same interactor, was measured by the portion of of all possible paired baits that had a statistically significant (p < 0.05) number of overlapping (convergent) genes as measured by a hypergeometric test with a background size of 11,169 proteins (the number of proteins in a HEK293T proteome; Bekker Jensen, 2017). To calculate a baseline, considering a study’s network size and degree distribution of the baits, we used random samples of baits from BioPlex to match each study. For each actual bait in a study, a random bait was chosen from BioPlex that had an identical number of interactors (degree), or from among the smallest balanced window of degree around the actual degree that included at least 100 different baits. 1000 randomly assembled networks were used for each study, and the full all-pair hypergeometric tests were done per randomly assembled network.

### Comparison of ASD-PPI and Pintacuda *et al.* 2023 PPI network

Pintacuda *et al.* 2023 reported a PPI network for 13 ASD risk genes in using IP-MS of endogenous proteins in human excitatory neurons derived from iPS cells^23^. Interactors from Pintacuda 2023 were extracted from Supplemental Table S3 (’Interaction_Annotations’ sheet).

We used Fisher’s enrichment tests (one-sided, greater) to assess whether there was greater than expected overlap in the combined interactors for all n=13 hcASD index proteins in Pintacuda 2023 and 1) the subset of ASD-PPI interactors from the matched set of n=13 hcASD baits; and 2) all ASD-PPI interactors from n=100 hcASD baits. We defined the universe of genes to be the set of n=2552 genes that are in the HEK293T proteome and in the set of detected proteins in Pintacuda *et al.* 2023^23,156^ (Pintacuda 2023 Table S3, union of the interactor and non-interactor proteins in the combined network). The 2×2 contingency table was defined as: X, the number of ASD-PPI interactors that are Pintacuda 2023 combined network interactors; Y; the number of ASD-PPI interactors that are not Pintacuda 2023 combined network interactors; Z, the number of Pintacuda 2023 interactors that are not ASD-PPI interactors; W, the number of genes that are not interactors in either ASD-PPI or Pintacuda 2023 combined network.

We additionally assessed whether damaging *de novo* variants in Pintacuda 2023 interactors or ASD-PPI subset interactors are associated with ASD. We focused on *de novo* damaging variants identified from the Simon’s Simplex Collection (Satterstrom *et al.* 2020^6^ Supplementary Table 1). We defined “damaging” variants to be variants that resulted in a damaging missense mutation (Polyphen Mis3 (damaging); or MPC Mis B (MPC>=2)) or PTV (frameshift, stop gained, or canonical splice site disruption).

(ssc_enrichment/scripts/01_format_satterstrom2020_SSC_variants.R). We defined 2 genesets of interest:

1. Pintacuda 2023 interactors: interactors from combined network in Pintacuda *et al.*2023 (n=979 genes).
2. ASD-PPI subnetwork interactors - hcASD13: ASD-PPI_13 interactors, excluding hcASD13 (n=222 genes)

For each Fisher’s enrichment test, the universe of genes was defined to be the geneset members, and the 2×2 contingency table was: X, the number of ASD probands with a damaging variant in at least one gene in the geneset; Y; the number of control siblings with a damaging variant in at least one gene in the geneset; Z, the number of ASD probands with no damaging variants in any gene in the geneset; W, the number of control siblings with no damaging variants in any gene in the geneset. We calculated the enrichment odds ratio with Fisher’s exact test (one sided, alternative = greater), and adjusted p values for multiple hypothesis testing (Bonferroni correction, p.adj = p*number genesets assessed).

We assessed whether there is significant difference in ORs for the Pintacuda 2023 complete network and the ASD-PPI_13 subnetwork (baits + interactors) using the Breslow Day test (R DescTools::BreslowDayTest). We found no significant difference (p=0.085).

### Comparison of ASD-PPI and Murtaza *et al.* 2021 PPI networks

Murtaza *et al.* 2021^24^ report ASD-relevant PPI networks generated by overexpressing baits and identifying interactors via *in vitro* proximity labeling (BioID2), including:

1. HEK-PPI: 7 ASD risk genes in HEK293T cells (3 baits overlapping with ASD-PPI baits)
2. Mouse-PPI: 41 ASD risk genes in mouse primary neurons co cultured with glia (17 baits overlapping with ASD-PPI baits). ASD risk genes were selected to have non-nuclear cellular localization.

HEK-PPI data was obtained from Murtaza *et al.* 2023 Table S2. We converted “Gene” column from mouse to human ontology (GRCm39 to GRCh38.p13) and removed those that did not map to human genes, self-interactions, and non-significant interactors (“HEK_Biotinylation” column = “NS”). This resulted in n=539 unique interactors (original 710).

Mouse-PPI data was obtained from Murtaza *et al.* 2023 Table S1. We trimmed Table S1 to the 41 ASD risk genes, converted “Prey” interactors from mouse to human ontology (GRCm39 to GRCh38.p13). We removed interactors that did not map to human genes and removed self-interactions, resulting in n=807 unique interactors (original 1107).

We defined the mouse brain proteome set of background genes using data from Murtaza *et al.* 2023 Table S5. We started with the list of mouse brain proteome genes in sheet “Mouse Brain Protein List”, converted the genes from mouse to human ontology (GRCm39 to GRCh38.p13), and removed genes that did not map to human genes. The final list consists of 8,678 genes (originally 11,992).

We conducted a Fisher’s exact test (one sided, greater) to evaluate the overlap between Murtaza 2023 HEK-PPI (7 baits) and combined interactors from ASD-PPI (100 baits) and HEK-PPI (7 baits). We defined the universe of genes to be the HEK293T proteome (union of Bekker Jensen 2017, ASD-PPI interactors, and HEK-PPI interactors, n=11,181 genes). The 2×2 contingency table was defined by ASD-PPI interactor (yes/no) and HEK-PPI interactor (yes/no). We conducted a second Fisher’s exact test (one sided, greater) to evaluate the overlap between ASD-PPI (100 baits) and Mouse-PPI (41 baits). We defined the universe of genes to be the intersection of the HEK293T proteome and mouse brain proteome (n=7049 genes). The 2×2 contingency table was defined by ASD-PPI interactor (yes/no) and Mouse-PPI interactor (yes/no).

We determined the association between having a damaging variant in different genesets of interest with ASD status. We defined the following genesets:

- ASD-PPI interactors: all ASD-PPI interactors from n=100 hcASD baits (n=1,074)
- HEK-PPI interactors: all interactors from Murtaza 2023 HEK-PPI that map to human genes (536 interactors from 7 baits)
- Mouse-PPI interactors: all interactors from Murtaza mouse-PPI that map to human genes (792 interactors from 41 baits)

Fisher’s enrichment tests were conducted as described in “Comparison of ASD-PPI and Pintacuda *et al.* 2023 PPI network”.

### Interactor expression in adult brain tissue (GTEx)

The Genotype-Tissue Expression (GTEx) project includes RNA-sequencing data from 54 non-diseased tissue sites across nearly 1,000 adult donors^54^. We downloaded GTEx v8 data (dbGaP Accession phs000424.v8.p2) from the GTEx website (https://www.gtexportal.org/home/datasets), which contains median gene-level TPM for samples from 54 tissue types, including 13 brain tissue sites. (gtex/scripts/01_downloadGtexData.R)

We assessed whether ASD-PPI genes are more highly expressed in GTEx brain tissues compared to other HEK293T expressed proteins. We defined HEK293T expressed proteins as the union of proteins detected by global quantitative mass spectrometry^155^ and ASD-PPI interactors. We defined genesets of interest to be baits, interactors (excluding hcASD102), and HEK293T expressed proteins that are not baits or interactors. For each geneset, we found the median geneset expression percentile within individual brain tissue samples and performed T-tests to assess whether these were significantly different across genesets. P values were adjusted for multiple hypothesis testing (Bonferroni correction, p.adj = p × 3 tests). (**Figure S2B**, gtex/scripts/02_GTEx_asdPPI100_analysis.R)

We evaluated whether interactors are more highly expressed in GTEx brain samples compared to permuted genesets from the HEK293T proteome. We created 100,000 permuted genesets from a universe of n=10,984 genes that are in the HEK293T proteome, measured in GTEx RNA-sequencing data, and not ASD-PPI baits. The probability of gene selection was weighted by HEK293T iBAQ scores^155^ and permuted genesets were required to have a median HEK293T iBAQ rank within 1 quantile of that of ASD-PPI interactors. We calculated the median geneset expression rank for interactors (observed) and for the 100,000 permuted genesets (null distribution) in GTEx brain tissue samples, where higher rank reflects higher relative expression within a sample. We then calculated the permuted significance of the observed ASD-interactor GTEx expression rank within each tissue type. P values were adjusted for multiple hypothesis testing (Bonferroni correction, p.adj = p × 13 brain regions). We found that 3 brain tissues - cerebellum (p.adj = 0.0023), cerebellar hemisphere (p.adj = 0.0043), and cortex (p.adj = 0.626, p = 0.048) - had at least nominally significantly higher median genset rank for interactor genes compared to permuted genesets. (**Figure S2C**, gtex/scripts/02_GTEx_asdPPI100_analysis.R).

### hcASD102 and interactor expression in human brain tissue (BrainSpan RNAseq data)

The BrainSpan developmental RNAseq dataset (‘bsRNAseq’) profiled human brain tissue RNA expression across the full course of human brain development, from early prenatal stages through late adulthood^150^. BsRNAseq data (n=524 samples) was downloaded from brainspan.org. The original 52,376 genes were trimmed to a final set of 18,552 protein-coding genes associated with an HGNC symbol; if a HGNC symbol was associated with multiple Ensembl Gene IDs, only one was kept. Expression values were reported in reads per kilobase million (RPKM). (bsRNAseq/scripts/ 01_download_format_bsRNAseq.R)

#### Expression of ASD-PPI genes and other HEK293T proteins in prenatal brain tissue

We assessed whether ASD-PPI genes are expressed at relatively higher levels in BrainSpan prenatal samples compared to 293T background genes. We defined 3 genesets of interest:

- Baits: ASD-PPI baits;
- Interactors: ASD-PPI interactors, excluding hcASD102;
- Other: n= 10,046 genes that are in the HEK293T proteome^155^ that are not in the ASD-PPI network.

We restricted our analysis to the n=237 prenatal brain samples as the prenatal period has been previously implicated in ASD^14–16^. For each geneset, we calculate the median geneset expression percentile within individual brain tissue samples. We performed T-tests to assess whether the median genset expression percentiles are significantly different between different genesets, adjusting p values for multiple hypothesis testing (Bonferroni correction, p.adj = p * 3 comparisons) (**Figure 2A**; bsRNAseq/scripts/02_asdPPI_bsRNAseq_prenatal_expression.R).

#### Calculating the relative expression of ASD-PPI genes compared to permuted genesets

We created 100,000 permuted genesets from HEK293T proteome genes, matching the number of interactor genes. To generate the permuted genesets, we selected genes from a set of n=10,902 genes that are 1) in the HEK293T proteome^155^ (described above), 2) measured in BrainSpan RNAseq, and 3) not an ASD-PPI bait. The probability of gene selection was weighted by HEK293T protein expression level (median iBAQ rank), and only genesets with a median iBAQ rank within 1 quantile of the median iBAQ rank of ASD-PPI interactors were retained. We calculated the median geneset rank (‘medRank’) within each bsRNAseq brain tissue sample for the interactors and each of the 100,000 permuted genesets (higher medRank reflects higher geneset expression within a sample). For each brain tissue sample, we calculated the ‘normalized medRank’, defined as the observed medRank – median (100,000 permuted medRanks). The normalized medRank reflects the sample-level expression enrichment of interactors, with positive values reflecting higher than expected expression. As a comparator, we also calculated the sample-level expression enrichment of hcASD102 genes^6^ compared to permuted genesets. (03_asdPPI_bsRNAseq_permutations.R)

#### Comparing expression levels of ASD-PPI baits and interactors in brain samples across development

To assess whether the relative expression of interactors mirror that of hcASD102 genes across samples, we calculated the Spearman and Pearson correlations of the normalized medRanks for interactors versus hcASD102 genes across n=524 brain samples (Spearman rho = 0.884, Pearson R = 0.897, indicating high correlation between interactors and hcASD102 expression across samples). To compare relative interactor (or hcASD102) expression in prenatal versus postnatal brain tissue, we grouped bsRNAseq brain samples by prenatal versus postnatal status (n=237 prenatal, n=287 postnatal), and assessed whether the normalized medRank of interactors (or hcASD102) were significantly different between prenatal versus postnatal samples (T-test, p<2.2e-16 for both interactors and hcASD102, prenatal samples with significantly higher expression of interactors and hcASD102 compared to postnatal samples). We also grouped bsRNAseq brain samples by developmental period (as defined in Kang *et al.* 2011^150^), where periods 1-7 reflect prenatal stages of development and periods 8-15 reflect late infancy through late adulthood, resulting in 13 sample groups ranging from period 2-14. For each period, we calculated the median(normalized medRank) across samples for interactors and hcASD102; we subsequently calculated the correlation of median(normalized medRank) for interactors versus hcASD102 across the 13 period groups (Spearman rho = 0.946, Pearson R = 0.901, indicating high correlation between interactors and hcASD102 expression across developmental periods) (**Figures 2B, 2C**, **S2D**; 04_make_bsRNAseq_asdPPI_hcASD102_permutation_plots.R).

### Interactor enrichment for ASD genetic risk

If ASD-PPI has successfully identified ASD network genes that are ASD-relevant, interactor genes should be enriched for ASD genetic risk. We used *de novo* genetic variants previously identified from simplex family studies as a measure of ASD genetic risk^6^ and conducted enrichment tests (Fisher’s exact) to assess whether interactors are enriched for ASD genetic risk compared to other genes in the HEK293T proteome or other exome genes.

We focused on *de novo* damaging variants identified from the Simon’s Simplex Collection (Satterstrom *et al.* 2020 Supplementary Table 1) We defined “damaging” variants to be variants that resulted in a damaging missense mutation (Polyphen Mis3 (damaging); or MPC Mis B (MPC>=2)) or PTV (frameshift, stop gained, or canonical splice site disruption). (ssc_enrichment/scripts/01_format_satterstrom2020_SSC_variants.R)

We determined the association between having a damaging variant in different genesets of interest with ASD status. We defined the following genesets:

- Interactors: ASD-PPI interactors (n=1,074)
- HEK293T (-Interactors, -hcASD102): HEK293T proteome genes, excluding interactors and hcASD102 (n = 10,045)
- Exome: all autosomal genes measured in Satterstrom *et al.* 2020 (n= 17,332)
- Exome (-hcASD102): autosomal genes measured in Satterstrom *et al.* 2020, excluding hcASD102 (n= 17,230)
- Exome (-Interactors, hcASD102): autosomal genes measured in Satterstrom 2020, excluding hcASD102 and interactors (n= 16,234)

For each Fisher’s enrichment test, the universe of genes was defined to be the geneset members, and the 2×2 contingency table was: X, the number of ASD probands with a damaging variant in at least one gene in the geneset; Y; the number of control siblings with a damaging variant in at least one gene in the geneset; Z, the number of ASD probands with no damaging variants in any gene in the geneset; W, the number of control siblings with no damaging variants in any gene in the geneset. We calculated the enrichment odds ratio with Fisher’s exact test (one sided, alternative = greater), and adjusted p values for multiple hypothesis testing (Bonferroni correction, p.adj = p*number genesets assessed) (**Figures 2D, S2E**; ssc_enrichment/scripts/03_asdPPI_geneticRisk_Satterstrom2020_SSC.R).

#### Downsampling analyzes

To determine whether increasing the number of baits used to create ASD-PPI is associated with increased ability to identify interactors associated with ASD genetic risk, we downsampled the ASD-PPI network. Specifically, we randomly selected sets of baits (ranging from n = 1-100 baits, 1000 iterations for each set size) and trimmed the ASD-PPI network to include only interactors associated with the downsampled baits. For each downsampled network, we calculated the association between having a damaging variant in different genesets of interest with ASD status as described above. We evaluated 2 genesets: 1) Interactors (-hcASD102), and 2) Exome (-Interactors, -hcASD102), which allows us to compare the amount of ASD-associated genetic risk in interactors versus the remaining genes in the human exome. For each bait set size, we calculated the median and standard deviation of ORs across the 1000 iterations, as well as the median and standard error of p-values across the 1000 iterations (**Figure 2F**; ssc_enrichment/scripts/04_asdPPI_prey_ASDrisk_downsamplingAnalysis.R).

We additionally assessed whether increasing the number of baits used to create ASD-PPI is associated with increased ability to identify interactors that are novel hcASD genes. We defined novel hcASD genes as the n=255 genes with FDR < 0.1 in the latest omnibus WES study^3^. We downsampled the ASD-PPI network as described above. For each downsampled network, we 1) counted the number of hcASD255 genes in the interactors and 2) conducted a Fisher’s exact test (one sided, greater) to assess for enrichment of hcASD255 in interactors. The counts of the two-by-two contingency tables were: X, the number of hcASD255 that are interactors; Y, the number of hcASD255 genes that are not interactors; Z, the number of interactors that are not in hcASD255; and W the number of genes that are not interactors or hcASD255. We defined the universe of possible genes to consist of those with 1) proteins that are detected in the HEK293T proteome^155^ or ASD-PPI interactors); 2) measured in Satterstrom *et al.* 2020^6^; and 3) measured in Fu *et al.* 2022^3^. For each bait set size, we calculated the median and standard deviation of OR, -log10(pvalue), and number of hcASD255 genes among interactors across the 1,000 iterations. We repeated this analysis after excluding hcASD102 genes (Satterstrom *et al.* 2020^6^, FDR <0.1) from the gene universe to assess the ability of ASD-PPI to identify truly novel hcASD genes (**Figure 2G**; risk_gene_enrichment/scripts/02_asdPPI_downsampling_hcASD255discovery.R).

### Interactor enrichment for ASD, DD, or SCZ risk genes

We evaluated whether interactors are enriched for risk genes that have been implicated in ASD, DD, or schizophrenia. We defined several sets of risk genes, including:

- ASD, Fu 2022^3^: n=255 autosomal genes with TADA FDR <0.1
- ASD, Trost 2022^4^: n=134 autosomal genes with TADA FDR<0.1
- ASD, Zhou 2022^5^: n=72 genes with study-wide significance (based on 5,754 constrained genes, p<8.69E-6)
- ASD, SFARI^152^: SFARI genes are a database that endeavors to include all genes associated with ASD risk, regardless of the level or nature of evidence. SFARI genes are further divided into categories, including syndromic, category 1 (high confidence), category 2 (strong candidate), and category 3 (suggestive evidence). SFARI genes were downloaded from gene.sfari.org (09/02/2021 release), and included a total of n=1,020 genes, of which there were n=230 syndromic genes, n = 92 syndromic and category 1 genes, n=206 category 1 genes, n=219 category 2 genes, and n=514 category 3 genes.
- DD, Kaplanis 2022^153^: n=285 genes that are significantly associated with DD after one-sided Bonferroni correction
- SCZ, Singh 2022^60^: n=34 genes with TADA FDR<0.1.

We defined the HEK293T expressed genes to be the union of interactors and proteins that were detected by global quantitative mass spectrometry^155^. We conducted two sets of enrichment tests.

For the first set of enrichment tests, we assessed only risk genes from genetic studies that reported the set of genes that were evaluated as possible risk genes^3–5,60^. We defined the universe of possible genes to be those that were 1) HEK293T expressed genes; 2) measured in Satterstrom *et al.* 2020^6^, and 3) measured in the genetic study that generated the risk gene set of interest ^3–5,60^ (**Figure 2E**). For the second set of enrichment tests (**Figure S2G**), we simply defined the gene universe to be HEK293T expressed genes, as the exact gene universe considered is unavailable for the SFARI risk genesets. For both sets of enrichment tests, we conducted Fisher’s exact tests (one sided, greater), in which the contingency tables were set up by interactor status (yes/no) and disease geneset status (yes/no). We corrected p values for multiple hypothesis testing (Bonferroni correction, p.adj = p * number of genesets assessed). (**Figures 2E**, **S2G**; risk_gene_enrichment/01_asdPPI_prey_riskGeneEnrichment.R; data/geneAnnotation_hek293T_ASD_DD_SCZ.csv).

### Evolutionary constraint metrics of ASD-PPI genes

We evaluated whether ASD-PPI genes are more evolutionary constrained than expected by assessing several metrics, including the pLI (probability of being intolerant of a single loss-of-function variant), misZ (missense Z score, measures gene intolerance to missense variation), synZ (synonymous Z score, measures gene intolerance to synonymous variation, used as negative control), and s_het (selective effect for heterozygous PTVs)^57,58^.

We obtained pLI, misZ, and synZ scores from the Genome Aggegation Database (ExAC dataset, https://storage.googleapis.com/gcp-public-data--gnomad/legacy/exac_browser/forweb_cleaned_exac_r03_march16_z_data_pLI_CNV-final.txt.gz) and s_het scores from Cassa *et al.* 2017 Supplementary Table 1^58^. We assessed whether the pLI, misZ, s_het, and synZ scores of ASD-PPI bait genes, interactor genes, and 293T proteome genes^155^ were significantly different from ASD-PPI baits, interactors (excluding hcASD102), and other HEK293T proteome genes using t-tests, adjusting p values for multiple hypothesis testing (Bonferroni correction, p.adj = pvalue * number of t-tests) (**Figure S2H**, 02_eval_asdPPI_geneConstraintMetrics.R).

### Comparison of interactor overlap between bait pairs predominantly associated with ASD or ASD/NDD

Satterstrom *et al.* 2020^6^ categorized hcASD into those that were more frequently mutated in ASD (ASD-predominant; ASD_P_) and those that were more frequently mutated in NDD (ASD_NDD_). If ASD_P_ vs ASD_NDD_ genes have separable molecular functions, we would expect ASD_P_ baits to have greater interactor overlap with other ASD_P_ baits (and the same for ASD_NDD_ bait pairs). We categorized ASD-PPI baits as ASD_P_ or ASD_NDD_ as defined in Satterstrom *et al.* 2020^6^ (Supplementary Table 2, Sheet 2, ASD_vs_DDID column). We evaluated the interactor overlap for the following four groups of bait pairs: 1) all ASD-PPI bait-bait pairs, 2) ASD_P_ - ASD_P_ bait pairs; 3) ASD_NDD_ - ASD_NDD_ bait pairs; and 4) ASD_P_ - ASD_NDD_ bait pairs. We determined the significance of interactor overlap using Fishers exact tests (one sided, greater), where the 2×2 contingency table was defined by: bait #1 interactor (yes/no) and bait #2 interactor (yes/no). The universe of genes was defined to be the HEK293T proteome. For each of the four baits groupings, we calculated the proportion of bait pairs whose interactors had nominally significant overlap (pval<0.05). We additionally used a Kruskal-Wallis rank-sum test to evaluate whether the distribution in bait-bait interactor overlap p values is significantly different across the 4 groups of bait pairs (R kruskal.test). We conducted these analyses using a trimmed dataset that includes only baits with fewer than 45 interactors (excludes 9 baits). Analyses using the full ASD-PPI dataset showed similar findings (data not shown).

### Prioritizing ASD-PPI interactors by number of hcASD interactions and shared interactors with hcASD

We were interested in ASD-PPI interactors that interact with large numbers of hcASD genes as they may be members of molecular complexes upon which hcASD genes functionally converge. We defined a set of n=7 ASD-PPI interactors (“prioritized interactors”) that interacted with at least 8 baits/hcASD102 genes (**Figure 5A**, asdppi_prioritizedPrey/scripts/ 01_asdPPIprey_topDegreeBait.R).

We assessed whether BioGRID interactors of these prioritized interactors are enriched for hcASD genes. Interactors were defiend using BioGRID) human physical interactions^92^. The file “BIOGRID-MV-Physical-4.2.191.tab2.txt” was downloaded from https://downloads.thebiogrid.org/BioGRID/Release-Archive/BIOGRID-4.2.191/. “Official.Symbol.Interactor.A” and interactors were defined using “Official.Symbol.Interactor.B”. Interactions were trimmed to human data (Organism.Interactor.A and Organism.Interactor.B = 9606). Bait and interactor symbols were updated using limma:alias2GeneSymbolTable. For each of the n=7 prioritized interactors, we conducted one-sided Fishers exact tests (greater) to assess whether interactors associated with individual baits are enriched for hcASD255 genes^3^. We restricted the gene universe to genes that were measured in Fu *et al.* 2022. The counts of the two-by-two contingency table were: X the number of interactors that are hcASD; Y, the number of interactors that are not hcASD; Z, the number of not.interactors that are hcASD; and W, the number of not.interactors that are not hcASD. We corrected p values for multiple hypothesis testing (Bonferroni correction, p.adj = pval* 7 genes). We found that DCAF7 BioGRID interactors to be nominally enriched for hcASD255 (**Figure 5B**, asdppi_prioritizedPrey/scripts/02_asdPPIprey_bioGRIDinteractors_hcASD255enrichment.R).

We were interested in identifying hcASD with interactomes that overlap with that of DCAF7. Therefore, we conducted two-sided Fisher’s exact tests to evaluate whether there is significant overlap between DCAF7 interactors and the interactors of each of the 8 hcASD that bound to DCAF7 in ASD-PPI. We restricted the gene universe to the HEK293T proteome. The counts in the 2×2 contingency table were: DCAF7 interactor in BioGRID (yes/no) and hcASD interactor in ASD-PPI (yes/no). We corrected p values for multiple hypothesis testing (Bonferroni correction, p.adj = pval* 7 genes).

### Effect of DYRK1A delta94-105 on DCAF7 and FAM54C interactions

We expressed Strep-tagged DYRK1A or DYRK1A^Δ80–100^ or empty vector (control) in HEK293T cells and conducted AP-MS as discussed above (3 replicates each for control and DYRK1A; 2 replicates for DYRK1A^Δ80–100^). We calculated SAINT scores (spectral counts). We defined significnat interactions to be those with SAINT 1-BFDR >=0.95. We evaluated the strength of binding (Log2FC(DYRK1A/control) spectral counts) of DYRK1A or DYRK1A^Δ80–100^ to DCAF7 and FAM54C.

### Enrichment of ASD risk genes among DCAF7-DYRK1A shared interactors

Co-expression of FLAG-tagged DCAF7 and Strep-tagged DYRK1A in HEK293T cells followed by sequential (double) AP-MS identified 126 shared interactors between DYRK1A and DCAF7 (see above). We evaluated whether these shared interactors are enriched for the n=255 ASD risk genes found to have FDR<0.1 in Fu *et al.* 2022^3^ using a two-sided Fisher’s exact test. The setup of the 2×2 contingency table was DCAF7-DYRK1A shared interactor (yes/no) and hcASD 55(yes/no). The gene universe was restricted to the HEK293T proteome (union of Bekker-Jensen 2017, ASD-PPI interactors and single/doubleIP interactors, n=11180 genes). (asdppi_prioritizedPrey/scripts/03_DCAF7interactors_hcASDenrichment_FET.R)

### Network co-expression in prenatal brain cells

We compared relative network co-expression in cells from three prenatal brain atlases. Briefly, for each cell type within a prenatal brain atlas, we evaluated the observed coexpression of network gene pairs and normalized this value by a background coexpression value to adjust for differences in the expected coexpression of genes present in the context from which the network was derived (i.e. for the 100 hcASD network we used all gene pairs present in the proteome of HEK293T cells^155^ and for the iEN network we used all gene pairs present in the iEN proteome^23^). We then compared the distribution of observed network co-expression across cell types using Wilcoxon rank sum tests.

We defined 2 sets of networks of interest:

1. Set 1: ASD-relevant PPI networks from experimental data and STRING^66^

a. ASD-PPI (HEK) hcASD-int: 1,143 nodes and 1,879 edges.
b. Pintacuda 2023 (iEN) hcASD-int: 1034 nodes and 1343 edges. 1343 index-interactor interactions identified for 13 hcASD index genes from iEN cells in Pintacuda *et al.* 2023. Interactions were extracted from Supplemental Table S3 (’Interaction_Lists’ sheet, selected rows where IsInteractor column is ’T’ and Index protein column was not ’COMBINED’).
c. ASD-PPI (HEK) hcASD-int & int-int: 1143 nodes and 10,191 edges. Union of interactions from ASD-PPI and STRINGv11.5 (experimental score >0) for genes in ASD-PPI
d. Pintacuda 2023 (iEN) hcASD-int & int-int: 1034 nodes and 11,706 edges. Union of interactions from Pintacuda *et al.* 2023 and STRINGv11.5 (experimental score >0) for genes in Pintacuda 2023 (iEN) hcASD-int.
2. Set 2: ASD-relevant PPI networks trimmed to direct interactions supported by AF

a. hcASD-int (AF iPTM > 0.5): 291 nodes and 284 edges. ASD-PPI hcInteractions with AF iPTM>0.5
b. hcASD-int & int-int (AF iPTM>0.5): 933 nodes and 2,350 edges. Union of and ASD-PPI hcASD-int and int-int interactions with AF iPTM>0.5.

We used data from three prenatal brain cell atlases:

1. The Nowakowski *et al.* 2017^62^ scRNAseq data (n = 4,261 cells) was downloaded from https://cortex-dev.cells.ucsc.edu/. The original set of n=56,864 genes was trimmed to a final set of 18,803 genes by keeping only protein coding genes associated with a HGNC symbol. If a HGNC symbol was associated with multiple Ensembl gene IDs, only the first was kept. Expression values were not altered from the original data download and are in the form of TPM. We grouped cells into the 19 cell types defined in Satterstrom *et al.* 2020^6^ and excluded unassigned cells from the analysis; see Supplemental Table 2 for the names of cell type in original publication and the simplified names used in this manuscript.
2. The Polioudakis *et al.* 2019^63^ scRNAseq data (n = 33976 cells) was downloaded from http://solo.bmap.ucla.edu/shiny/webapp/. Genes were trimmed to a final set of 16548 genes by keeping only protein coding genes associated with a HGNC symbol. If a HGNC symbol was associated with multiple Ensembl gene IDs, only the first was kept. Expression values were not altered from the original data download and are in the form of UMI counts. Cell types were defined in Polioudakis *et al.* 2019, and consisted of 16 cell types: vRG, ventricular radial glia; oRG, outer radial glia; PgS, cycling progenitors (S phase); PgG2M, Cycling progenitors(G2/M phase); IP, intermediate progenitor; ExN, migrating excitatory; ExM, maturing excitatory; ExM-U, maturing excitatory upper enriched; ExDp1, excitatory deep layer 1; ExDp2, excitatory deep layer 2; InMGE, interneuron MGE; InCGE, interneuron MGE; OPC, oligodendrocyte precursor; End, endothelial cell; Per, pericyte; Mic, microglia. See Supplemental Table 2 for the names of cell type in original publication and the simplified names used in this manuscript.
3. The Bhaduri *et al.* 2021^67^ scRNAseq data (neocortex, GW15-25, n = 404,218 cells) was downloaded from https://cells-test.gi.ucsc.edu/?ds=dev-brain-regions+neocortex. Cell types were defined using the “ConsensusCellType-Final” column in the meta.tsv file (n=12 cell types). We excluded ’Excitatory Neurons’ from our analyses as this cluster contains few cells (n=61) and this cluster is not included in the Bhaduri *et al.* 2021 main text Figure 2. See Supplemental Table 2 for the names of cell type in original publication and the simplified names used in this manuscript.

scRNAseq data is limited by dropouts and resultant sparse and heterogeneous gene expression across cells. Therefore, we define gene pairs to be ’co-expressed’ in a cell if there is at least 1 read of each gene detected. We define the co-expression between gene pairs as the proportion of cells in a cell type that co-express both genes (range 0-1).

We defined two sets of background genes:

1. HEK: 11185 genes in the HEK293T proteome (union of Bekker Jensen 2017^155^ and ASD-PPI genes). We used this background for ASD-PPI based networks.
2. iEN: 3274 genes in the iEN proteome as defined in Pintacuda *et al.* 2023^23^ (Pintacuda *et al.* 2023 Supplemental table 3, sheet ’Interaction_Lists’, ’Gene symbol’ column). We used this background for Pintacuda *et al.* 2023^23^ based networks.

For each cell type context, we determined the average background co-expression by calculating the connectivity between all possible gene-gene pairs. We use this background co-expression as a normalization factor to remove biases that have to do with global differences in co-expression in each cell type. Let i and j denote genes and k denote cell type context. Define r_ijk_ as the co-expression between genes i and j in cell type k, and define avg(r_k_) as the average co-expression over all pairs of background genes measured in cell type k. Then define a normed co-expression as:

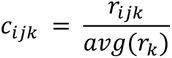

Within each prenatal brain atlas, for each cell type, we calculated the normed co-expression of all observed network edges. We then evaluated whether the distribution of observed network co-expression within each individual cell type was significantly different from the base distribution (concatenated observed network co-expression of all cell types) using a two-sample Wilcoxon rank sum test (R wilcox.test). We used the Wilcoxon estimator for the difference in location as a measure of relative difference in network co-expression between in the cell type versus base distribution). Within each single cell atlas, p-values were corrected for multiple hypothesis testing (Bonferroni correction, p.adj = pval * number of cell types * number of networks evaluated).

### Generating a random set of ASD-PPI bait-int

We generated a set of random interactors to use as a baseline benchmark for the ASD-PPI AlphaFold-Multimer (hereafter also referred to as AF) pairwise interaction analysis. For each of the baits in ASD-PPI, we generated a set of random interactors that matched true interactors in protein size and for which odds of selection were weighed by HEK293T protein expression levels. Specifically, we defined HEK293T protein expression level by average iBAQ expression (Bekker-Jensen 2017^155^ Supplementary table 7, average of iBAQ scores for the two HEK293T experimental replicates) and used cDNA length reported in Satterstrom *et al.* 2020 as a proxy for protein size (Satterstrom *et al.* 2020^6^ Supplementary Table 2, “Autosomal” sheet, “cDNA” column). We divided the roughly 10,000 HEK293T expressed proteins into 5 groups based on cDNA length (cDNA_quantile). For each observed ASD-PPI interactor, we selected a random interactor from HEK293T-expressed proteins in the same cDNA quantile, with probability of selection weighted by HEK293T protein expression level. Random interactors were required to be unique at a bait level but could be duplicated across different baits.

### Running AlphaFold to predict direct pairwise interactions

Pairwise protein interactions and protein complexes were predicted using AlphaFold-Multimer^157^ (AlphaFold version 2.3.1) via ColabFold^158^ (version 1.5.2). As input to protein structure prediction, sequences for each protein pair were extracted from uniprot (https://ftp.uniprot.org/pub/databases/uniprot/current_release/knowledgebase/reference_proteo mes/Eukaryota/UP000005640/UP000005640_9606.fasta.gz). In all cases, we utilized the wildtype protein sequence even when examining interactions defined by mutant-specific baits. Multiple sequence alignments (MSA) were generated from ColabFold using MMseqs2^78^ (Release 14), with the following databases: Colabfold Env DB (2021_08), UniRef30 (2021_03). All input complexes were run without templates or amber relaxation for 10 recycles and with at least three random seeds. For bait-prey interactions three model replicates were run for each seed, bringing the total number of replicates to at least nine. Only one model per seed was run for the int-int set, so only three models per pair were produced. Some inputted sequences failed to return structures due to memory limits (particularly for large complexes) or MSA failures. The five different sets of pairs modeled were: bait-int produced 1654 structures of 1879 attempted pairs, bait-random produced 1677 for 1862, bait-int-mutant (those pairs only observed in the mutant network) produced 126 for 148 pairs, bait-int-Pintacuda produced 1133 for 1303, and int-int produced 32,890 for 35,897.

From each replicate, multiple statistics were generated. AF predicted scores (pTM, ipTM, pLDDT) were extracted directly from replicate json files and PDB files. The pDockQ score is based on parameters fit to match true DockQ values for simulated protein complexes with resolved crystal structures, and it was calculated as described previously^79^. The Confidence score was a weighted combination of pTM and ipTM, 0.8 × ipTM + 0.2 × pTM.

### Searching Protein Data Bank (PDB) for homologous complex structures

As a proxy for possible presence of a complexed structure for any pair in the AF training set, we searched the full PDB for structures with detectably similar sequences. BLAST+ v2.15 was used to run blastp with default parameters against a custom database built based on all sequences deposited to PDB (downloaded on 11/17/2023). Possible matches were assembled in an iterative process over all sequences in any AF pair. For each protein, all individual chains that return a BLAST match were identified. For each matching chain, the other chains within each corresponding PDB entry were then queried to identify the subset that matches to a protein that forms an AF pair to the starting protein.

### Visualizing protein structure files

ChimeraX or PyMOL were used to generate all 3D figures of protein structures.

### Calculating distances from mutations to PPI interface

Inter-chain distances in an AF model were calculated using the center-center inter-atom distances between the three-dimensional coordinates of atoms in the AF structure file. Per residue, we used the minimum distance across all atoms to any atom in the opposite chain with plDDT at least 25.0. Only wildtype sequences were modeled, so the distance is from any atom in the wildtype residue.

### Evaluating residue-level evolutionary conservation with ConSurf

We used the ConSurf^159–161^ web server (https://consurf.tau.ac.il/consurf_index.php) to estimate the evolutionary conservation of amino acid positions for DCAF7 and DYRK1A based on the phylogenetic relations between homologous sequences. We ran ConSurf on each protein separately with default parameters (one HMMER iteration, maximum 150 homologs, E-value cutoff of 0.0001). The sequences found were clustered by their level of identity using CD-HIT and the cutoff specified to be between 35% to 95% identity)

### Evaluating whether Mut-PPI changed interactors are enriched for ASD risk genes

We evaluated whether the interactors that demonstrate increased (n=69) or decreased (n=95) interaction with hcASD in Mut-PPI are enriched for the n=255 ASD risk genes with FDR < 0.1 in Fu *et al.* 2022^3^. We defined the gene universe to be HEK293T expressed proteins measured in Fu *et al.* 2022. The 2×2 contingency table used for the two-sided Fisher’s enrichment tests were defined by: hcASD255 (yes/no), changed interactor (yes/no). P values were corrected for multiple hypothesis testing (Bonferroni correction, padj = pval * 2 tests).

### Protein System Enrichment

#### *De novo* mutations

A comprehensive list of exome-wide *de novo* somatic mutations identified in the ASD cohort considered in this research was obtained from the large-scale whole exome sequencing study from Satterstrom *et al.* 2020^6^. We used the 5287 filtered probands from the analyses and considered only functionally disrupting missense and protein truncating variant mutations. Specifically, we only kept mutations that met any of the following criteria: (1) VEP_functional_class_canonical_simplified equal to one of [“frameshift_variant”, “stop_gained”, “splice_acceptor_variant”, “splice_donor_variant”] AND pLI >= 0.995; (2) MPC >= 2; or (3) Polyphen_prediction == “probably_damaging’’.

#### Integrated protein network

We compiled a diverse set of experimental and curated protein-protein interaction datasets (**Table S6**). We processed each PPI network as follows: (1) standardized the naming of all the gene symbols by mapping to their standard symbols via HUGO Gene Nomenclature Committee (HGNC) ID; (2) removed interactions with no documented score; (3) removed self-interactions; and (4) removed duplicates. In addition to aforementioned processing, for the case of Mentha and BioGRID, we only kept interactions in humans (Taxon A/B == 96096 in Mentha & Organism Name Interactor A/B == “Homo sapiens” in BioGRID). We further filtered BioGRID to include only physical interactions.

The ASD-PPI and other PPI networks were integrated into a single interaction network. The integration was performed using BIONIC^162^, a graph convolutional neural network that uses graph attention networks. We used the following parameter settings {epochs: 1000, batch_size: 512, learning_rate: 0.00005, embedding_size: 512}, with the remaining parameters using the default values. BIONIC ran until the reconstruction loss stabilized.

We computed an interaction score between each pair of proteins by calculating the cosine similarity score between their embeddings. We then focused the network on the 102 identified high-risk ASD-genes and the interactors of ASD-PPI and ASD_mut_-PPI by obtaining their top associated proteins. We further retained all ASD-PPI interactions.

#### Multi-scale map of protein systems

To identify a multi-scale map of systems of tightly connected proteins, we applied Hierarchical community Decoding Framework (HiDeF)^163^ to the integrated network. HiDeF determines connected proteins across various thresholds and identifies systems from most stringent thresholds to least stringent, where the smaller more stringent identified systems were part of the larger less stringently identified systems. We used the following parameter setting {maxres: 50, tau: 0.75, chi: 5, alg: ‘leiden’}.

#### Differential proteins systems

To identify systems that have stronger evidence in ASD-PPI, ASD_mut_-PPI, or balanced, we modified the computed WT integrated network according to interactions found to have stronger support in either ASD-PPI or ASD_mut_-PPI. We then generated another multi-scale map using the modified network using the same multiscale community detection algorithm described previously. We computationally assessed which systems have stronger evidence in ASD_mut_-PPI or ASD-PPI by aligning the original and modified multi-scale maps using the align function developed by Dutkowski & Kramer *et al.*^164^.

#### Systems enriched with mutated genes and disrupted interactions

To identify systems that have significant enrichment for functionally disrupting mutations, we performed hypergeometric tests on the number of mutated genes in each system. To compute the number of mutated genes, we identified genes with functionally disruptive *de novo* mutations as described in section “Mutation data”. The significance of the number of mutated genes was computed using the hypergeom.sf function from the scipy package in Python. We used a cutoff of 0.05 to determine significance.

#### Protein system names

The protein systems were labeled using a GPT-4-based pipeline developed by Hu *et al.* 2023^165^. This pipeline assigns a set of genes with succinct literature-driven names that summarizes their consensus functions as well as providing supportive analysis text. We used the latest version of GPT-4 (gpt-4-1106-preview) for this analysis.

### Evaluating whether FOXP1-mutant differentially bound genes or DEGs are enriched for ASD risk genes

We evaluated whether FOXP1-differentially bound genes or cell type-specific differentially expressed genes (DEGs) from FOXP1^WT/WT^ vs FOXP1^R513H/WT^ organoids are enriched for a set of n=255 ASD risk genes with FDR < 0.1 in Fu *et al.* 2022^3^. We defined genes that are more strongly (CUT&Tag pval<0.05 & log2FoldChange>0) or weakly (pval<0.05 & log2FoldChange<0) bound by FOXP1 in FOXP1^R513H/WT^ organoids. For each cell type, we defined DEGs that are upregulated in FOXP1^R513H/WT^ (scRNAseq p_val_dj<0.05 and pct.1-pct.2 >0) or downregulated (val_adj<0.05 and pct.1-pct.2<0) in FOXP1^R513H/WT^ organoids. We defined the gene universe to be the intersection of genes expressed in each cell type and measured in Fu *et al*. 2022. We conducted two-sided Fisher’s enrichment tests, defining the 2×2 contingency table by the parameters: hcASD255 (yes/no) and differentially bound/DEG (yes/no). We corrected p values for multiple hypothesis testing (Bonferroni correction, p.adj = pval* #comparisons).

### HEK293T microscopy

HEK293T cells were transfected with DYRK1A, DCAF7, KIAA032 plasmids using Lipofectamine 3000 (Invitrogen, #L3000001) and fixed after 30h with 4% paraformaldehyde (EMS, #50980487). After permeabilization with 0.1% TritonX-100, cells were stained with anti-Flag (1:1000, Cell Signaling, 14793), anti-Strep (1:1000, IBA, 2-1507), anti-ɑ-Tubulin (1:500, Santa Cruz, sc-53029) and anti-DYRK1A (1:500, R&D systems, AF5407). Secondary fluorescence-conjugated antibodies were used at 1:1000 (Abcam ab150177, Abcam ab150159, Thermo Fisher A31553, Thermo Fisher A32732). Cells were imaged on a Zeiss 980 LSM confocal microscope with fast airyscan with a 63X oil objective and processed in ImageJ and Adobe Illustrator.

### Obtaining *Xenopus tropicalis* Embryos and Tadpoles

Ovulation was induced by injection of human chorionic gonadotropin (Sigma) into the dorsal lymph sac according to standard procedure^166^ and in accordance with approved UCSF IACUC protocols. Natural matings and *in vitro* fertilizations were performed. Embryos and tadpoles were staged by ^167^. Clutch mates were always used as matched controls.

### Xenopus tropicalis Microinjections

*Xenopus tropicalis* embryonic microinjections were performed as described before^166^. Microinjections were performed at the 2-cell stage using a Narishige micromanipulator, Parker Picospritzer, and Zeiss Stemi microscopes. Injection volume was calibrated with an eye-piece micrometer. Embryos were grown between 22–25°C in 1/9 Modified Ringer’s (MR) solution, which was refreshed daily. Male and female embryos were analyzed.

### *Xenopus* CRISPR/Cas9 Genome Editing

High-efficiency sgRNAs were designed, synthesized, and validated as in ^168^. For each embryo, 3 ng of purified Cas9-NLS protein (Macrolabs, UC Berkeley), 800 pg sgRNA, and a dextran dye conjugated with Alexa-555 (Invitrogen) were injected into 1 cell of a 2-cell stage embryo. The day following injection at stages 14–20, embryos were sorted left from right according to the dye.

### *Xenopus* Whole Mount Immunofluorescence Staining

*Xenopus* immunostaining was carried out as previously described^102^ with a primary antibody against beta-tubulin (1:100, DSHB E7) and a secondary goat anti-mouse fluorescent antibody (1:250, Life Technologies A32723).

### Microscopy and *Xenopus* Brain Size Measurements

*Xenopus* tadpole images were acquired on a Zeiss Axio Zoom.V16 microscope with apotome and a 1X objective. Images were processed using Fiji^169^. Telencephalon size was calculated from stereoscope images of brain immunostainings using the freehand select and measure functions in Fiji^169^. The injected side was compared to the noninjected side (internal control). These measurements were from two-dimensional images taken from a dorsal perspective and reflect relative size differences. GraphPad Prism software version 8.3 was used to graph data and determine statistical significance using a student’s paired t-test. p values less than 0.05 were considered significant.

### Differentiation of iPSCs to NPCs

Dorsal forebrain patterned NPCs were generated from iPSCs using a small molecule protocol adapted from a published method^170^. Specifically, we plated iPSCs on Matrigel coated plates in mTeSR plus medium (STEMCELL Technologies, Cat#100-0276). For the next 3 days, we fed cells with the KSR medium (15% Knockout Serum Replacement in Knockout DMEM, 1xGlutaMAX, 1xMEM-NEAA, 0.1mM BME) containing small molecules 250nM LDN193189 (Tocris, Cat No. 6053), 10uM SB431542 (Tocris, Cat#1614) and 5uM XAV939 (Tocris, Cat#3748) (LDN/SB/XAV) for dual SMAD inhibition and Wnt inhibition. On day 4 and day 5, we fed cells with in ⅔ KSR + ⅓ N2 (DMEM/F12, 1x N2, 1x B27 -Vitamin A, 1x GlutaMAX, 1x MEM-NEAA) + LDN/SB/XAV and ⅓ KSR + ⅔ N2 + LDN/SB/XAV respectively. On day 6, we passaged cells at 1:2 using EDTA and plated cells onto Matrigel coated plates. For day 6 and day 7, cells were cultured in NPC medium (DMEM/F12, 1xN2, 1xB27 -Vitamin A, 1x GlutaMAX, 1x MEM-NEAA, 10ng/ml FGF2, 10ng/ml EGF) supplemented with 5uM XAV. On day 8, we passaged the cells at 1:3 using Accutase (STEMCELL Technologies, Cat#07920) and cultured cells in NPCs medium onwards.

### CRISPR/Cas9 mediated DCAF7, DYRK1A, KIAA0232 knockdowns in neuronal progenitor cells (NPCs)

We designed non-targeting control and *DCAF7*, *DYRK1A* or *KIAA0232* targeting single guide RNA (sgRNA) using a bioinformatics pipeline developed by Martin Kampamann’s lab at UCSF (https://github.com/mhorlbeck/CRISPRiaDesign). Individual sgRNA was cloned into lentiviral vector pMK1334 expressing BFP (Addgene Cat# 127965). Lentiviruses carrying sgRNAs were produced in Lenti-X 293T cells (Takara Bio, Cat#632180) by transfection and concentrated using Lenti-X concentrator (Takara Bio, Cat# 631231). We transduced NPCs with the concentrated lentiviruses and selected the transduced NPCs using 4ug/ml puromycin until the BFP+ cells reached greater than 90%. We then cultured the NPCs in NPC medium (DMEM/F12, 1xN2, 1xB27 -Vitamin A, 1x GlutaMAX, 1x MEM-NEAA.) on tissue culture plates coated with Martrigel (Fisher Scientific, Cat# 08-774-552). The knockdown efficiency of sgRNA was measured by qPCR.

### Cortical neuron differentiation

Cortical neuronal differentiation was carried out as described before170. NPCs were thawed and grown to confluency in a 6-well dish. The cells then were dissociated using Accutase for 5 minutes at 37°C to single cell resolution and plated at ∼10,000 cells/well in NPC media and 10 µM ROCKi. Lentiviral constructs of either RGEDI^2^ or pHR-hSyn:EGFP (Addgene #114215)^3^ were added to the cell suspension at 5 MOI at the time of passage and removed after 48 hours. The NPCs were cultured in a 12-well dish to confluency in NPC media and then dissociated using Accutase for 5 minutes at 37°C to single cell resolution and plated on 5 µg/mL human laminin and 5µg/mL fibronectin-coated 384 well plates (Perkin Elmer Cat #6057308) at 12,000 cells/well in NPC media and 10 µM ROCKi. 24 hours later, media was replaced with Neuronal Differentiation media consisting of: 1:1 DMEM/F12:Neurobasal Media (Gibco 21103049), 1x N2, 1x B27 -Vitamin A, 1x GlutaMAX, 0.5 mM dibutyryl cAMP (Sigma D0627), 0.2 mM ascorbic acid (Sigma A4403-100MG), 1 μM PD0325901 (Selleck Chem S1036), 5 μM SU5402 (Selleck Chem S7667), 10 μM DAPT (R&D v), 20 ng/mL BDNF (R&D 248-BD). Media was refreshed at 75% every other day up to Day 18 of differentiation.

### Immunocytochemistry

Fixed cells were permeabilized using a 0.1% Triton-X/PBS solution for 20 minutes at room temperature. Permeabilization solution was removed, and a 1 M glycine solution added and incubated at room temperature for 20 minutes. A blocking solution of 0.1% Triton-X/PBS, 2% FBS, and 3% BSA was added after removal of the glycine solution and incubated at room temperature for 1.5 hours. MAP2 (1:1000 Abcam chicken-anti MAP2 # ab5392) and Ki67 (1:200 Millipore mouse-anti Ki67 # mab4190) antibodies were diluted in blocking solution and incubated overnight at 4°C. Primary antibody was removed by washing cells 3 times with 0.1%Triton-X/PBS. Secondary antibodies (goat anti-chicken 488 Abcam # ab96947), goat anti-mouse 555 Invitrogen # A21426) were added to 1:1000 in blocking solution and incubated at room temperature and covered for 1.5 hours. Cells were spun down at 8000 RPM in a cold centrifuge during the washes. Cells were washed in PBS for 5 minutes, then once with PBS plus Hoechst at a dilution of 1:1000 in PBS and incubated for 10 minutes at room temperature, covered. The Hoechst was washed out with PBS and the cells covered in PBS for imaging.

### Longitudinal imaging

Cells that contained the red GEDI biosensor were imaged on an ImageXpress Micro Confocal High-Content Imaging System from Molecular Devices for 7-10 days starting on differentiation day 7. Montages of 9 tiles were imaged in RFP (200 ms) and GFP (100 ms) channels at 20x magnification per well. Ki67-stained NPCs were imaged at 9 tiles per well in the RFP (100 ms), GFP (50 ms), and DAPI (15ms) channels at 20x per well. Images were processed using a custom workflow in Galaxy^171^.

### Odds ratio of cell death (GEDI)

Cells from each line were assessed for being alive or dead by the GEDI^110^ biosensor on different plates at multiple points in time. The change in odds of cell death (OR-CD) at any point in time is modeled using a generalized linear model using the glm function in R, between the cell line and time using the binomial probability distribution as the family argument to this function. The modeled changes include the plate on which the cell was assayed, the time (as a continuous variable) at which the cell status was ascertained, the cell-line from which the cell was derived and the interaction between the cell-line and time. These changes, as odds ratios are derived from the model fits to the data. Custom -built scripts in R were developed to model the data.

### Cell Profiler

The Ki67 proliferation assay used a modified Cell Profiler^172^ example pipeline for colocalization (https://cellprofiler.org/examples). Images were pooled in CellProfiler and analyzed on a per-image basis. Background values were calculated per image then subtracted from the whole image. Following background correction, all objects in each DAPI image were identified as nuclei using minimum and maximum diameters per object and filtering out excessive intensity values using a Minimum Cross-Entropy thresholding method. This method identifies all possible nuclear objects within the appropriate size range. Intensity values were calculated per object then used to filter out non-nuclear objects or dead cells that display bright nuclei that have much higher intensities than live cells, which were relabeled as nuclei segments. Images that contained Ki67 staining in the FITC channel were run through a feature enhancement step that increases the signal-to-noise ratio. Enhanced images were run through segmentation to identify all FITC objects that fit Ki67 signal criteria in terms of size and signal intensity again using a Minimum Cross-Entropy thresholding method. These identified objects were renamed as Ki67 segments. Ki67 segments were related to the nuclei segments to determine how many nuclei were Ki67-positive. Finally, the number of Ki67-positive nuclei was divided by the number of total nuclei to calculate the percent of Ki67-positive nuclei. The segmentation overlays and math were exported from the program and then plotted in Prism10.

### Statistical Analysis

We developed a statistical package called RMeDPower^173^ in R, a complete package of statistical tools that allow a scientist to understand the effect size and variance contribution of a set of variables within a dataset to a given response. RMeDPower uses linear mixed models on repeated measures data such as those described here. Outliers were removed and data was log transformed for statistical analysis. All p values and estimates for each comparison of NPCs or cortical neurons that contain scrambled versus ASD gRNAs for both the nerite analysis and percent Ki67-positive staining were calculated in RMeDPower.

### hiPSC CRISPR/Cas9 genome editing plasmid design

The closest PAM sequence to FOXP1 R514H for CRISPR/Cas9 genome editing was identified, and the corresponding sgRNA sequence (GTGCGAGTAGAAAACGTTAA) was cloned into the pX459 plasmid (Addgene #62988) using BbsI Golden Gate cloning. To construct the donor plasmid, an IDT gBlock was ordered to create the silent R5414H mutation, as well as create a silent mutation in the PAM site to prevent further editing. ∼500bp homology arms to the FOXP1 cut site were generated using PCR, and the homology arms and gBlock were inserted into the pUC18 plasmid (Addgene #50004) linearized with HindIII and BamHI.

### hiPSC CRISPR/Cas9 genome editing and genotyping

eWT-1323.4 hiPSCs^174^ were incubated for one hour in StemFlex media supplemented with the CEPT small molecule cocktail^175^ containing: Chroman 1 (MedChem Express #HY-15392, 50nM); Emricasan (SelleckChem #S7775, 5µM); Polyamine Supplement (Sigma Aldrich #P8483, 1:1000 diluted); and trans-ISRIB (Tocris #5284, 0.7 µM). For electroporation using the Invitrogen Neon Transfection System, cells were enzymatically lifted using TrypLE (Gibco #12605010), quenched with 10% FBS, and counted using a hemocytometer. 100,000 cells per electroporation reaction were pelleted and resuspended in 5uL Buffer R per reaction. Separately, 2µg pX459, 1µg pUC18, and Buffer R were mixed to 7µL. 5µL cells were added to the plasmid mix, and 10µL cells and plasmid mix were electroporated using a 10µL pipet tip (Invitrogen #MPK1025) with the following settings: 1100 V, 30ms, 1 pulse. Electroporated cells were added to plates with prewarmed StemFlex media with CEPT and incubated overnight. Starting 16 hours after electroporation, cells were incubated with media supplemented with 0.5µg/ml Puromycin for 72 hours to select for cells that received the pX459 plasmid and were more likely to be genome edited.

Seven days after electroporation, individual colonies were manually passaged, with one colony per one well of a 12 well plate. After reaching confluency, cells were passaged once more, with remaining cells lysed for gDNA extraction using the NEB Monarch Genomic DNA Isolation Kit (#T3010S). Clonal gDNA was PCR amplified to contain the CRISPR/Cas9 edit site, PCR products were purified using the Macherey Nagel NucleoSpin PCR clean up kit (#740609) and submitted for Sanger sequencing. The correctly targeted clone and two untargeted clones were expanded.

### hiPSC-derived cortical organoid differentiation

hiPSCs were lifted using ReLeSR and resuspended in Neural Induction Media containing GMEM (Gibco #11710035), 10% Knockout Serum Replacement (Gibco #10828028), NEAA diluted 1:100 (Gibco #11140050), Sodium Pyruvate diluted 1:100 (Gibco #11360070), 5mM 2-mercaptoethanol (Sigma Aldrich #M6250), and 100µg/mL Primocin supplemented with 5µM SB431542 (Tocris #1614), 100nM LDN-193189 (Sigma Aldrich #SML0559), and 3µM IWR1-endo (Cayman Chemicals #13659). Cells in Neural induction media were moved to 6-well low attachment plates (Corning #3471), with a media change on day 3 with CEPT, and day 6 without CEPT. From day 9-25, organoids were cultured in Maintenance Media 1: 50% DMEM/F12 with Glutamax (Gibco #10565042) and 50% Neurobasal (Gibco #21103049) with B27 without vitamin A (Gibco #12587001), N2 (Gibco #17502048), NEAA diluted 1:100, Glutamax diluted 1:200 (Gibco #35050061), and 55µM 2-mercaptoethanol supplemented with 10ng/mL each FGF (Peprotech #100-18B) and EGF (Peprotech #100-47). Media was changed every 2-3 days. From days 26-35, media was changed without FGF and EGF. From day 55 onward, organoids were cultured in Maintenance Media 2: Maintenance Media 1 supplemented with B27 with vitamin A (Gibco #17504001), instead of B27 without vitamin A. From day 63-70, organoids were cultured in Maintenance Media 2 supplemented with 10ng/mL each of BDNF (Alomone #B-250) and NT3 (Alomone #N-260).

### Cortical organoid fixation, embedding, cryosectioning

Organoids were fixed in 4% PFA in PBS for 30 minutes at room temperature, then washed with PBS three times. Organoids were then dehydrated in 30% sucrose in PBS at 4C overnight.

Cryomolds were filled with a 1:1 mixture of 30% sucrose in PBS and OCT, and organoids were embedded in the cryomolds and frozen on dry ice before being stored at -80C. Fixed and frozen organoids were cryosectioned to 18µm thickness and mounted on SuperFrost Plus microscope slides (Fisher Scientific #12-550-15).

### Immunohistochemistry

Sectioned organoids were first rehydrated in PBS for 10 minutes and then treated with boiling 10mM sodium citrate solution pH 6 for 15 minutes. Slides were washed once with PBS and blocked for one hour at room temperature in blocking buffer of PBS with 1% Normal Donkey Serum (Jackson ImmunoResearch #017-000-121), 0.1% Gelatin, and 1% Triton X-100. Primary antibodies were diluted in blocking buffer and added to slides in a hybridization chamber, where slides were incubated at 4C overnight. Slides were then washed three times with PBS with 1% Triton X-100 and incubated with secondary antibodies diluted 1:1000 in blocking buffer in a hybridization chamber at room temperature for 3 hours. Slides were washed three times in PBS with 1% Triton X-100 and coverslips were mounted using ProLong Gold Antifade Mountant (Invitrogen #P36930).

The following primary antibodies and dilutions were used: FOXP1 (Abcam #ab227649 1:100), FOXP2 (R&D Systems #AF5647 1:500), FOXG1 (Abcam #ab18259 1:1000), DLX2 (Santa Cruz Biotechnology #sc-393879 1:50), PAX6 (Biolegend #901301 1:200), Ki67 (Dako #M7240 1:200), CTIP2 (Abcam #ab18465 1:500).

### Imaging and quantification

Prepared slides were imaged on a Leica SP8 laser scanning confocal microscope using a 20x air objective. Images were processed using Fiji and quantified using CellProfiler^172^. Mann-Whitney U test was used to determine significant differences in expression, with FDR q = 0.01.

### Cortical organoid dissociation

Organoids were dissociated using 20 units of Papain (Worthington #LK003178) with 5% Trehalose in HBSS for 30 minutes at 37C. DNase was added, and organoids were incubated for another 15 minutes at 37C. Papain was quenched using Albumin-ovomucoid inhibitor and cells were filtered through a 40µm cell strainer. Cells were spun down, resuspended, and counted.

### CUT&Tag

CUT&Tag was performed on dissociated organoids as previously described^136^ with some modifications. Briefly, 200,000 cells per reaction were pelleted at 600g for 3 minutes at room temperature and resuspended and fixed in PBS with 0.1% formaldehyde for 2 minutes. 1.25M glycine was added to double the molar concentration of formaldehyde and stop cross linking, and cells were spun down at 1300g at 4C for 3 minutes. Cells were resuspended in wash buffer (20mM HEPES pH 7.5, 150mM NaCl, 0.5mM spermidine, and 1 EDTA-free complete protease inhibitor tablet). Concanavalin A-coated beads (Fisher Scientific #NC1526856) were prepared by adding 10µL beads per reaction to bead-binding buffer (20mM HEPES pH 7.9, 10mM KCl, 1mM CaCl_2_, and 1mM MnCl_2_). Using a magnetic rack, binding buffer was removed, and beads were resuspended in binding buffer once more before removing the binding buffer again and finally resuspending in enough binding buffer for 10µL per reaction. 10µL beads were then added to cells and incubated on an end over end rotator for 10 minutes at room temperature. Wash buffer was removed from cells using a magnetic rack, and cells were resuspended in enough antibody buffer (wash buffer with 2mM EDTA, 0.1% BSA, and 0.05% digitonin) for 50µL per reaction. Primary antibodies (FOXP1 Cell Signaling Technologies #2005S, FOXP2 Abcam #ab16046, FOXP4 Millipore #ABE74, H3K27me3 Cell Signaling Technologies #9733, IgG Epicypher #13-0042) were added at 1:50 dilution and samples were nutated overnight at 4C.

Using a magnetic rack, primary antibody was removed, and cells were resuspended in 100µL secondary antibody (Antibodies Online #ABIN101961) diluted in wash buffer with 0.05% digitonin. Cells were then nutated for one hour at room temperature. Secondary antibody mix was removed using a magnetic rack, and cells were washed with wash buffer with 0.05% digitonin three times. pA-Tn5 preloaded with Nextera adapters (Epicypher #15-1117) was diluted 1:20 in dig-300 buffer (20mM HEPES pH7.5, 300mM NaCl, 0.5mM spermidine, 0.015% digitonin with 1 EDTA-free complete protease inhibitor tablet) and cells were resuspended in 50µL of pA-Tn5 mix. Cells were nutated for one hour at room temperature. Using a magnetic rack, pA-Tn5 mix was removed, and cells were washed three times in dig-wash buffer. Cells were resuspended in 300µL of tagmentation buffer (dig-300 buffer with 10mM MgCl_2_) and incubated in a 37C water bath for one hour. Cells were released from beads with addition of 10µL 0.5M EDTA, 3µL 10% SDS, and 2.5µL 20mg/ml proteinase K. Samples were vortexed and incubated in a heat block at 55C for one hour.

Fragments were purified by adding 300µL phenol:chloroform:isoamyl alcohol (25:24:1 v/v) and sample to a phase lock tube (Qiagen #129046), and spun down at 16,000g for 3 minutes. 300µL chloroform was added to each sample and spun down once more at 16,000g for 3 minutes. The aqueous layer was added to a 1.5ml lo-bind tube with 750µL 100% ethanol and mixed by pipetting. Samples were cooled on ice and spun down at 16,000g for 15 minutes at 4C. Supernatant was decanted, and samples were washed once more with 1ml 100% ethanol and spun down at 16,000g for 1 minute at 4C. Ethanol was decanted, the pellet was air dried, and resuspended in 22µL water. Libraries were amplified by mixing 21µL sample, 2µL each i5 and i7 primers (Illumina #FC-131-2001), and 25µL NEBNext High Fidelity 2X Master Mix (NEB #M0541S) and using the following PCR cycle settings: 72C for 5 minutes, 98C for 30 seconds, 98C for 10 seconds, 61C for 10 seconds, repeat steps 3-4 15x, and a final 72C incubation for 1 minute. Libraries were purified using SPRI Select Reagent (Beckman Coulter #B23317) and eluted in 20µL water.

Libraries were pooled to 2nM and diluted to a final concentration of 750pM with 2% PhiX spike in for sequencing using the Illumina NextSeq 2000 with a targeted read depth of ∼10 million reads per sample. 2 technical replicates were used per sample.

### CUT&Tag analysis

Reads were trimmed using TrimGalore, aligned to reference genomes for hg38 and *E. coli* using bowtie2 with parameters “--end-to-end --very-sensitive --no-mixed --no-discordant --phred33 -I 10 -X 700.” Duplicates were marked and removed with picard. Peaks were called using MACS2 using the corresponding IgG sample as a control, with q = 0.01.

DESeq2 was used to determine differential peaks between FOXP1 R514H and WT. Wald test was used to determine significance with Benjamini-Hochberg correction.

### scRNA-seq

Dissociated organoids as described above were used as input for Fluent Biosciences PIP-seq T2 V4.0 kit^176^ with 20,000 cells per reaction as input with two technical replicates per sample. Libraries were pooled to 2nM and sequenced at 750pM final concentration with 2% PhiX spike-in. Libraries were paired-end sequenced with a targeted sequencing depth of 20,000 reads per cell on a NextSeq 2000.

### scRNA-seq analysis

Cell by gene matrices for each sample/replicate were generated using PIPseeker and were integrated into one object using Seurat. High quality cells were subsetted by removing those with less than 500 or greater than 10,000 genes; less than 1000 UMIs; and greater than 10% mitochondrial reads. Reads were normalized using SCTransform and batch correction was done with Harmony. Cells were clustered using resolution 0.2. Clusters were manually annotated using cluster markers. Differentially expressed genes between FOXP1 R514H/+ and WT in each cluster were determined using FindMarkers with the default parameters.

During the preparation of this work the authors used chatGPT-3 to shorten text sections. After using this tool/service, the authors reviewed and edited the content as needed and take full responsibility for the content of the publication.

