## Supplemental Figures and Tables for "A foundational atlas of autism protein interactions reveals molecular convergence"

Supplementary Figure 1

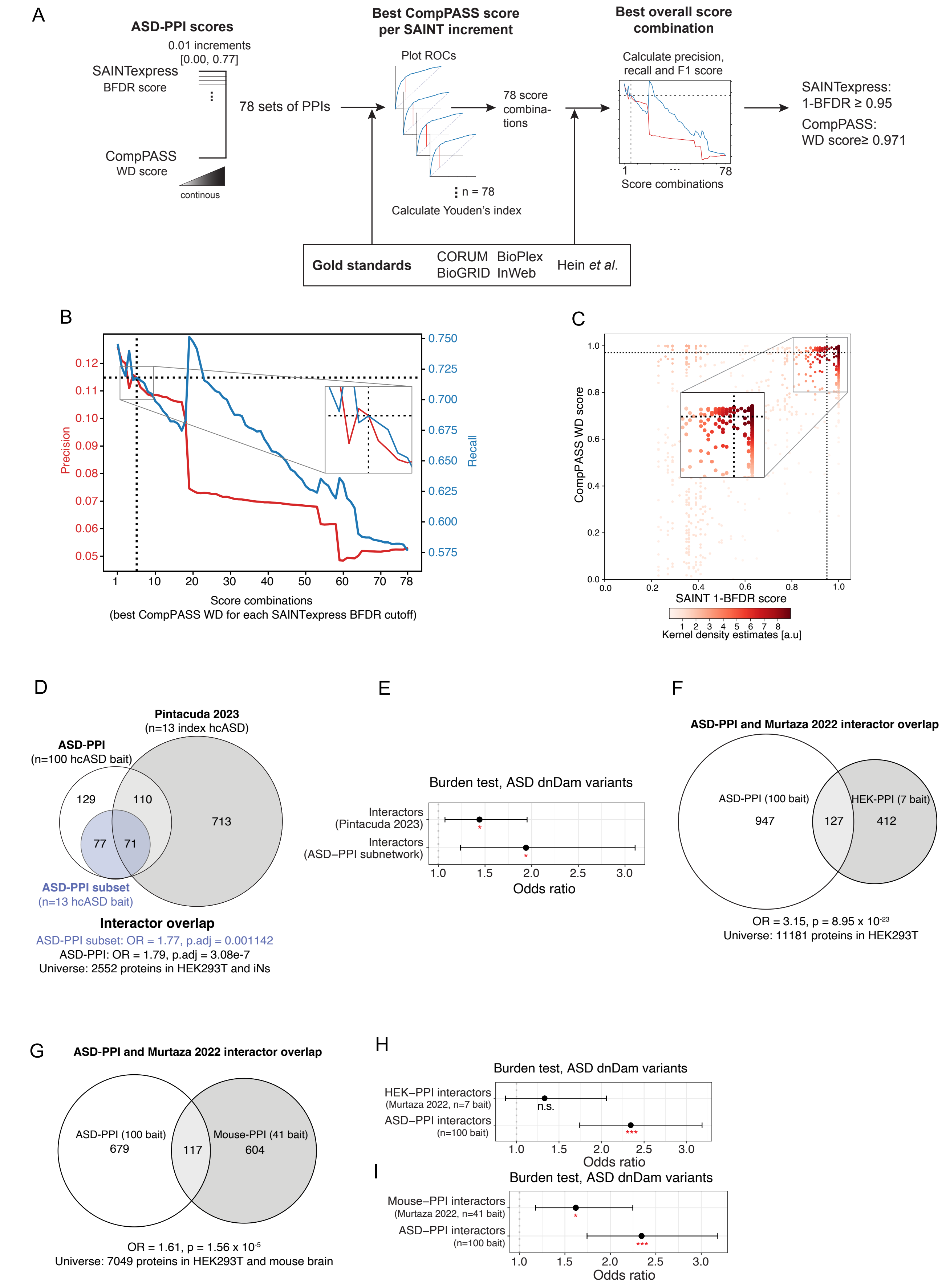

Supplementary Figure 2

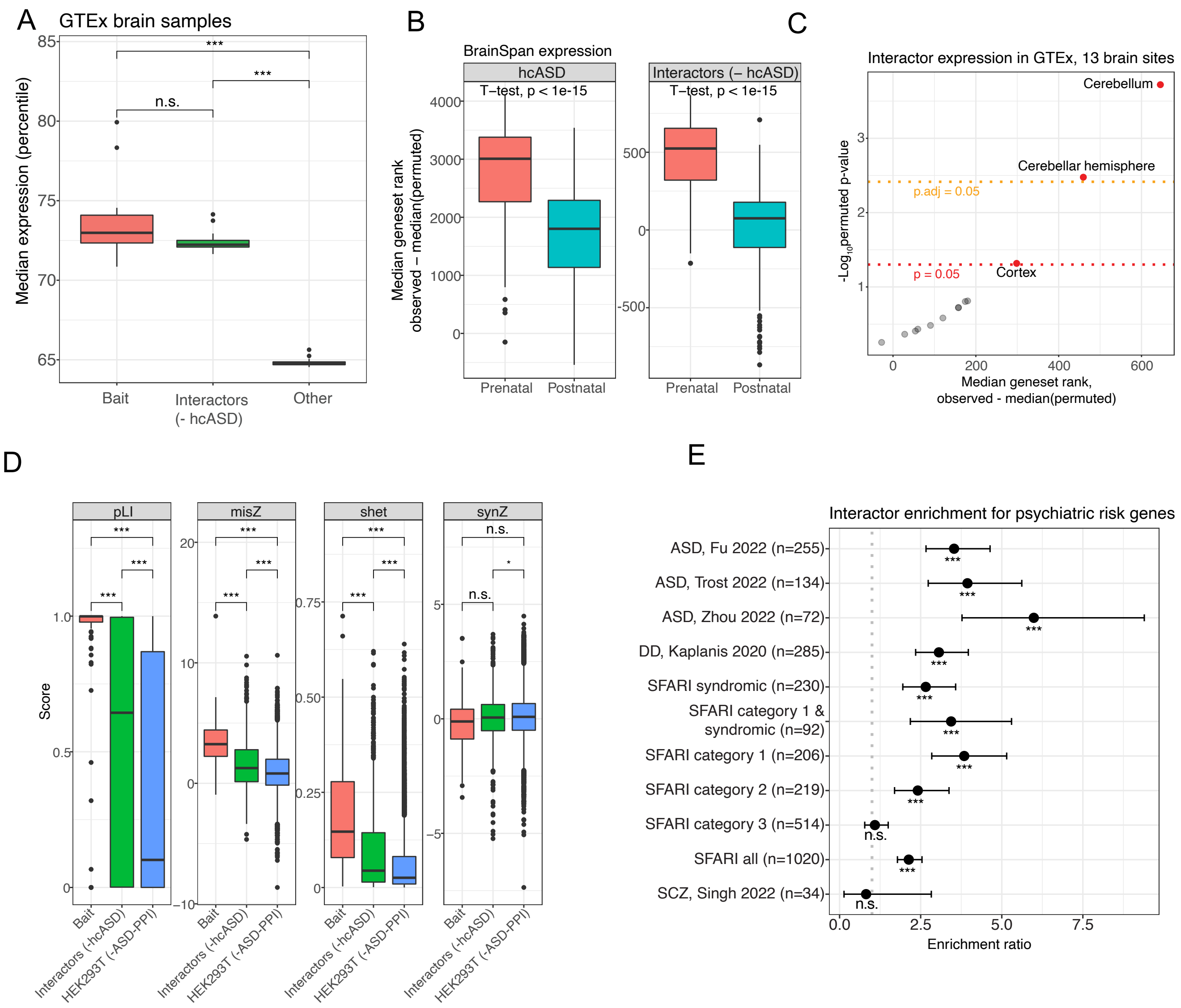

Supplementary Figure 3

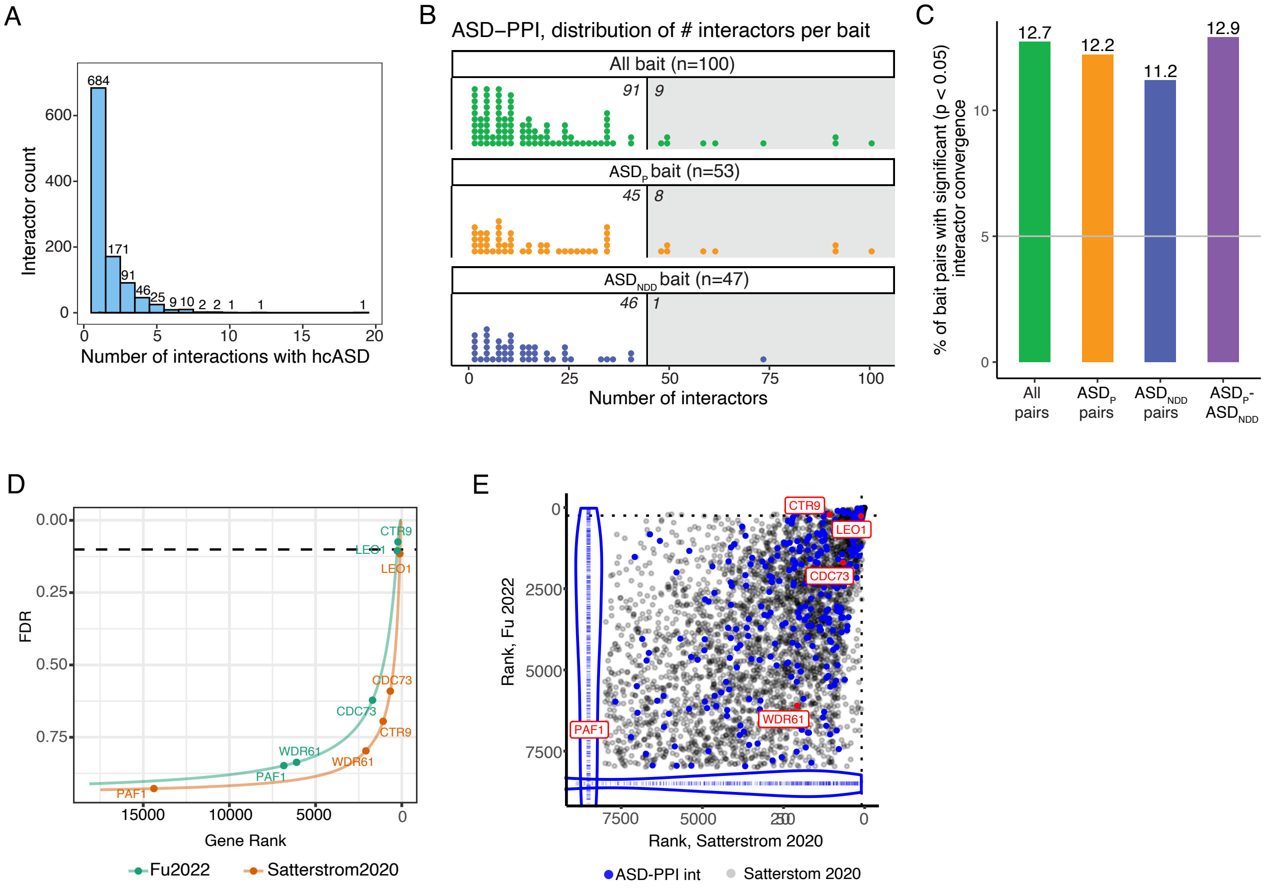

### Supplementary Figure 4

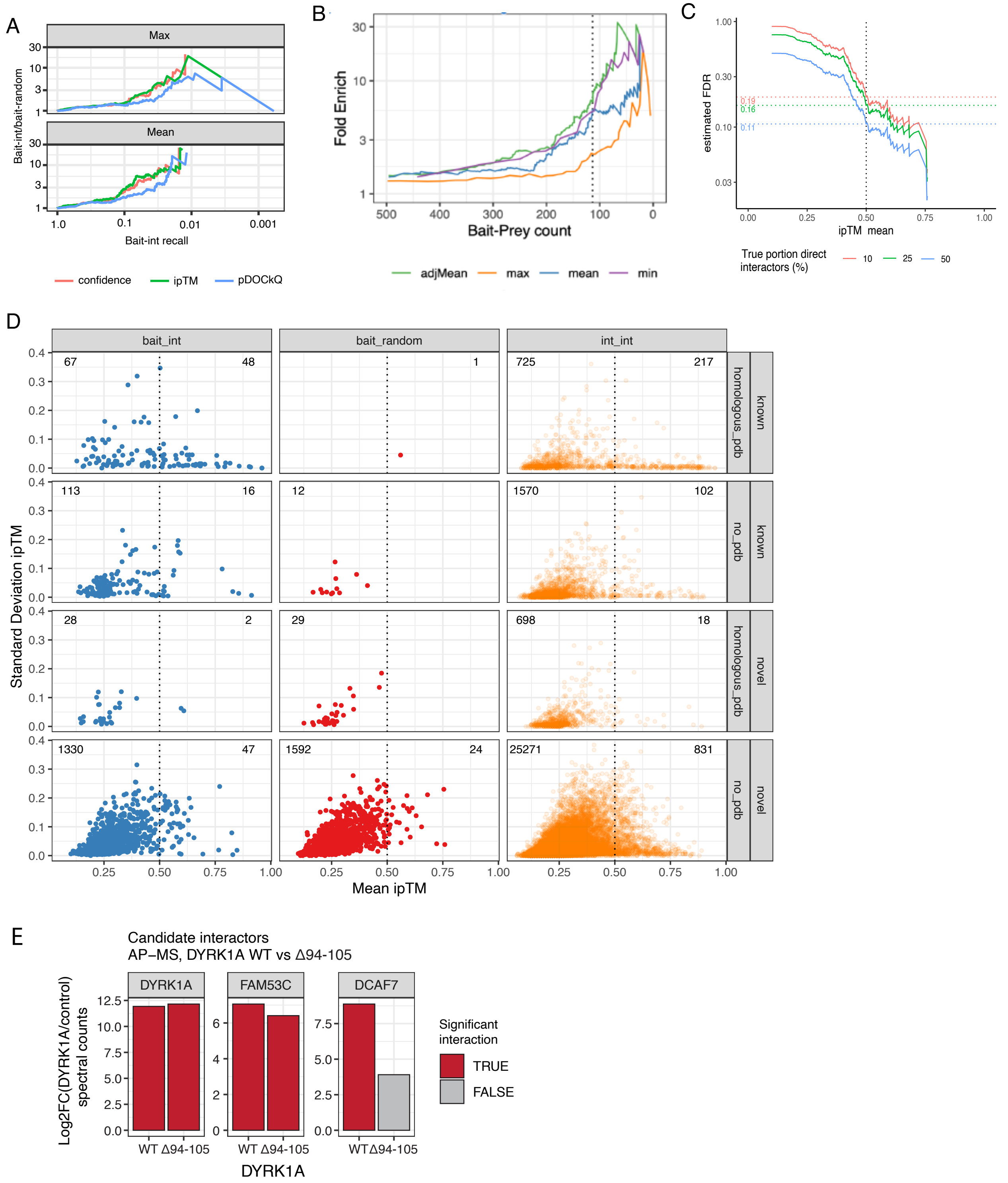

Supplementary Figure 5

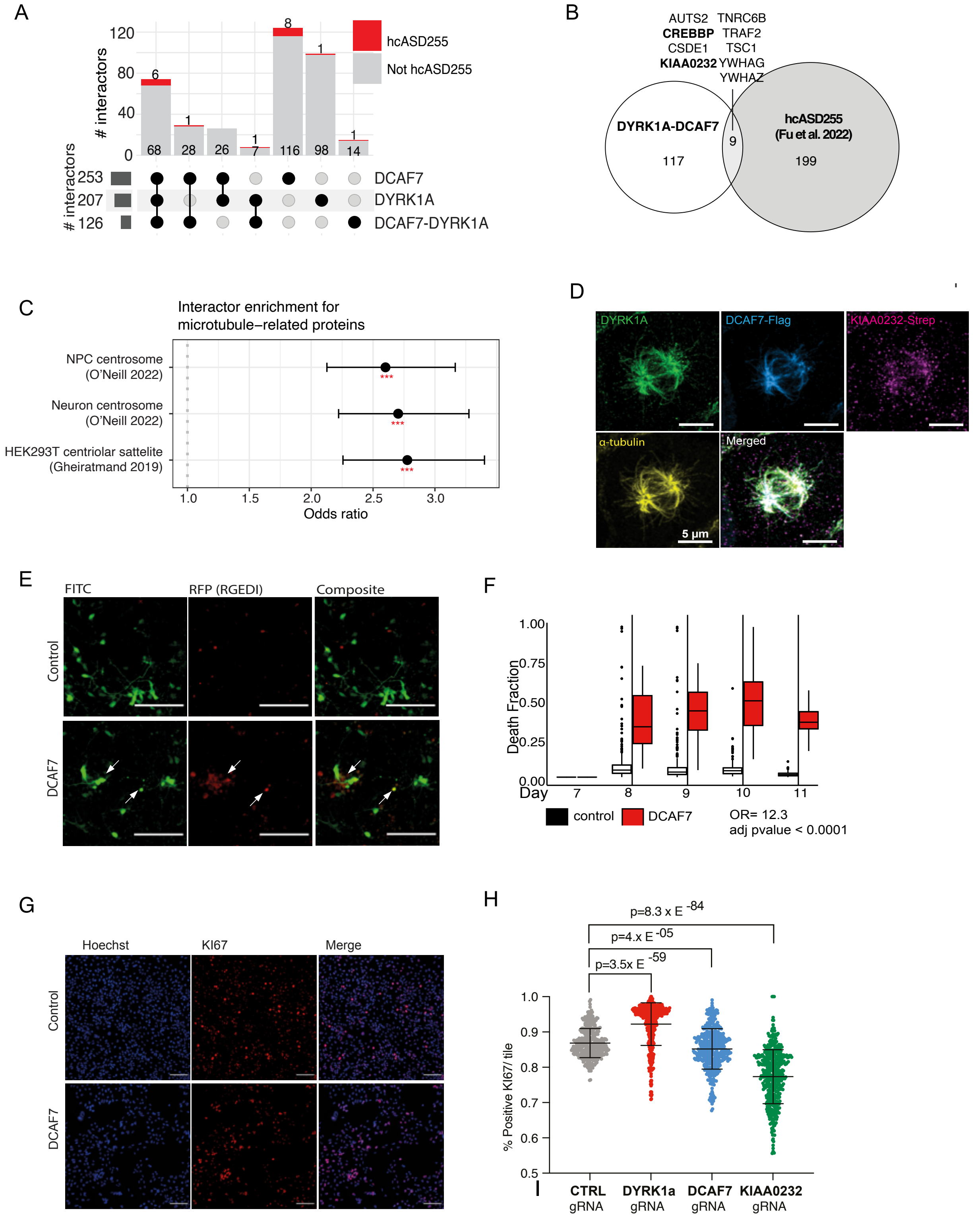

Supplementary Figure 6

A

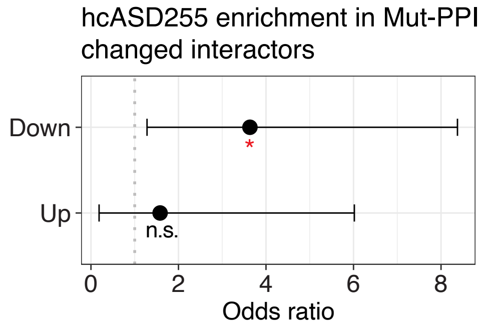

B

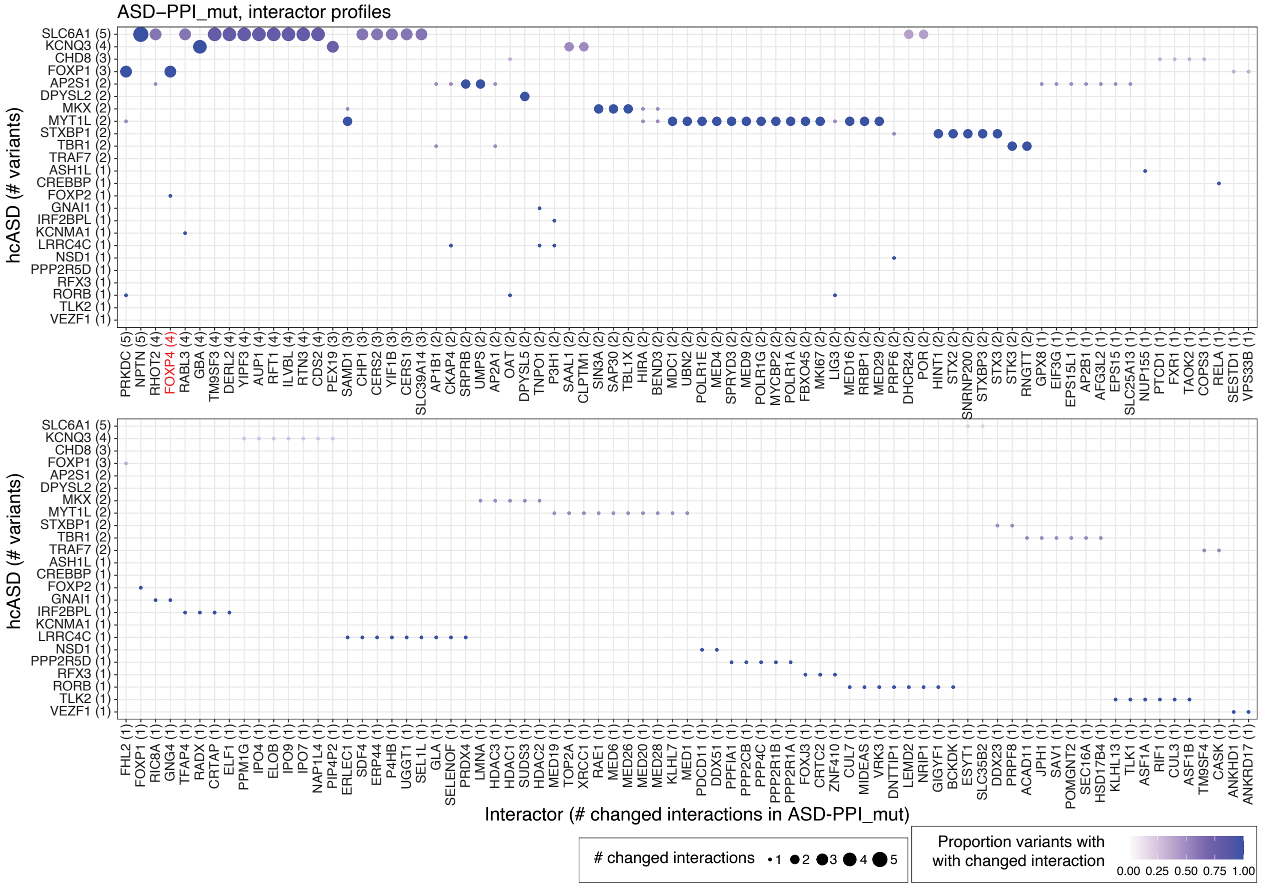

C

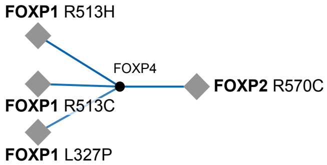

D

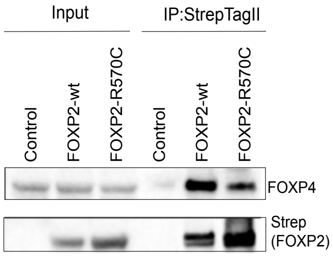

Supplementary Figure 7

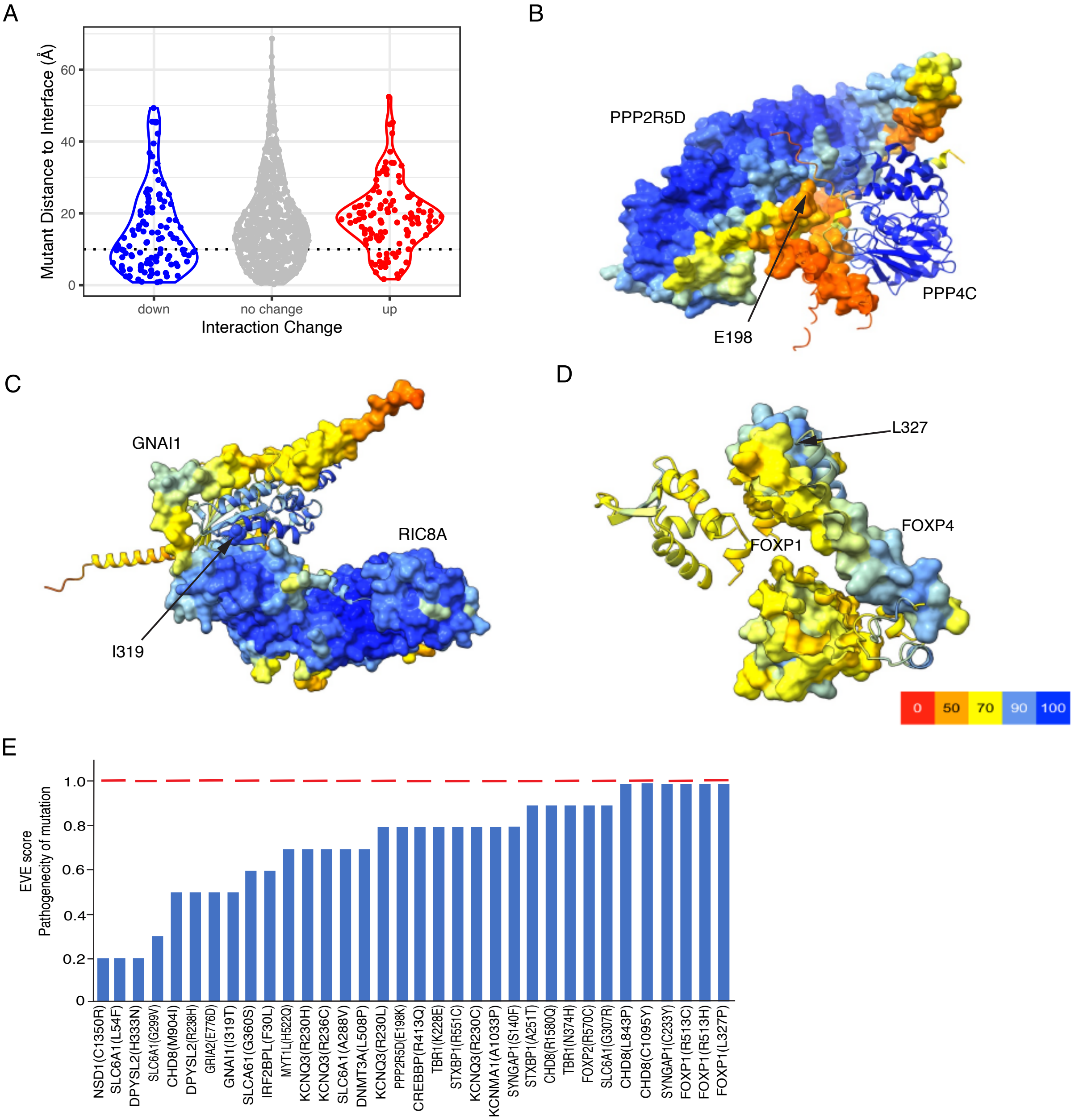

Supplementary Figure 8

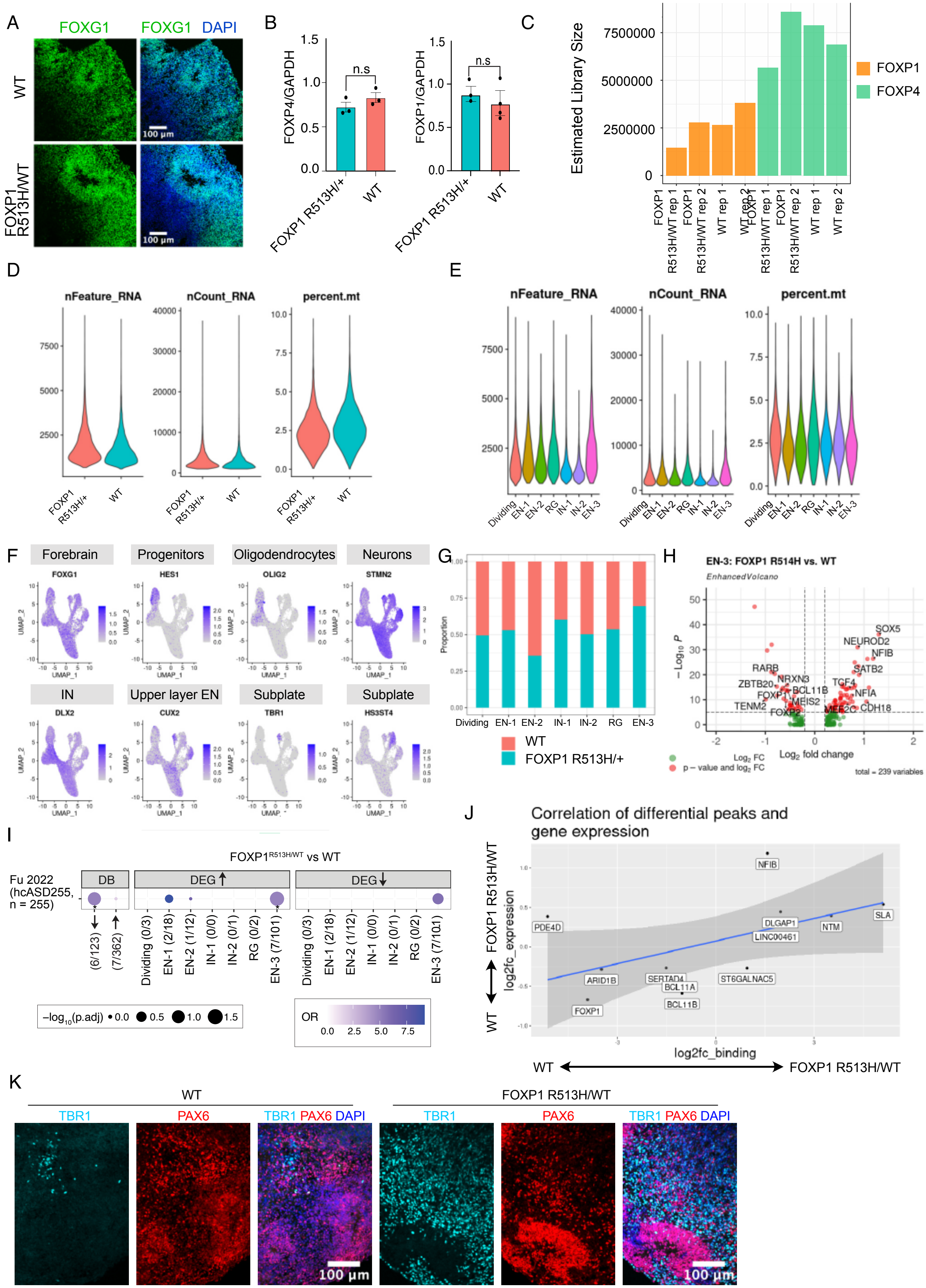

### **Supplemental Table: High confidence interactors of hcASD by AP-MS scores**
#
### The full set of interactors (columns Interactor_uniprot and Interactor) of 100 different
### hcASD (column Bait). Interactions are labeled as bait_int unless the interactor is another
### hcASD then it is labeled bait_bait (column type). Scores used for thresholding are given
### for AP-MS scoring algorithms SAINTexpress (SAINT_BFDR) and CompPASS (CompPASS_rank_wd)
#
### Confidence scores from either SAINT or CompPASS are given.
#
### Scoring is based on counts of peptide-matched spectra per protein and replicate, and are
### described per replicate in column Spec, with AvgSpec averaging across replicates. Column
### ctrlCounts shows the same spectral counts for the Interactor protein in the control AP-MS
### samples.
#
### The interactions that were considered in the gold standard set are indicated, along with
### the source(s) of the knowledge, by columns is_gold_standard and gold_standard_source.

| Bait | Interactor_unipro | Interactor | type | SAINT_BFDR | CompPASS_rank_W | Spec | AvgSpec | #Replicates | ctrlCounts | gold_std? |
|---|---|---|---|---|---|---|---|---|---|---|
| FOXP1 | Q13363 | CTBP1 | bait_int | 0 | 0.980475735 | 3\|2\|4 | 3 | 3 | 0\|0\|0\|0\|0\|0 | FALSE |
| FOXP1 | Q8IVH2 | FOXP4 | bait_int | 0 | 0.996957347 | 17\|18\|19 | 18 | 3 | 0\|0\|0\|0\|0\|0 | TRUE |
| FOXP1 | O15409 | FOXP2 | bait_bait | 0 | 0.997795983 | 14\|15\|14 | 14.33 | 3 | 0\|0\|0\|0\|0\|0 | TRUE |
| FOXP1 | P56545 | CTBP2 | bait_int | 0 | 0.975866705 | 13\|15\|7 | 11.67 | 3 | 0\|0\|0\|0\|0\|0 | FALSE |
| FOXP1 | O75694 | NUP155 | bait_bait | 0 | 0.984388212 | 3\|10\|7 | 6.67 | 3 | 0\|0\|0\|0\|0\|0 | FALSE |
| ASH1L | O75694 | NUP155 | bait_bait | 0 | 0.988376929 | 10\|11\|13 | 11.33 | 3 | 0\|0\|0\|0\|0\|0 | FALSE |
| RFX3 | Q13445 | TMED1 | bait_int | 0 | 0.975305305 | 3\|2\|3 | 2.67 | 3 | 0\|0\|0\|0\|0\|0 | FALSE |
| RFX3 | Q9P0L0 | VAPA | bait_int | 0.01 | 0.971153713 | 2\|3\|2 | 2.33 | 3 | 0\|0\|0\|0\|0\|0 | FALSE |
| RFX3 | Q86VK4 | ZNF410 | bait_int | 0 | 0.987971473 | 3\|2\|3 | 2.67 | 3 | 0\|0\|0\|0\|0\|0 | FALSE |
| RFX3 | Q9NX40 | OCIAD1 | bait_int | 0 | 0.981335163 | 4\|3\|4 | 3.67 | 3 | 0\|0\|0\|0\|0\|0 | FALSE |
| RFX3 | O14662 | STX16 | bait_int | 0 | 0.99375875 | 4\|4\|2 | 3.33 | 3 | 0\|0\|0\|0\|0\|0 | FALSE |
| RFX3 | Q9NRW7 | VPS45 | bait_int | 0 | 0.996312776 | 11\|7\|5 | 7.67 | 3 | 0\|0\|0\|0\|0\|0 | FALSE |
| RFX3 | Q53ET0 | CRTC2 | bait_int | 0 | 0.993471119 | 6\|4\|4 | 4.67 | 3 | 0\|0\|0\|0\|0\|0 | FALSE |
| RFX3 | Q9UPW0 | FOXJ3 | bait_int | 0 | 0.994327082 | 4\|4\|3 | 3.67 | 3 | 0\|0\|0\|0\|0\|0 | FALSE |
| RFX3 | P28074 | PSMB5 | bait_int | 0.01 | 0.975936014 | 2\|2\|2 | 2 | 3 | 0\|0\|0\|0\|0\|0 | FALSE |
| RFX3 | P22670 | RFX1 | bait_int | 0 | 0.979831164 | 6\|6\|7 | 6.33 | 3 | 0\|0\|0\|0\|0\|0 | TRUE |
| RFX3 | Q02086 | SP2 | bait_int | 0.01 | 0.982568858 | 2\|2\|2 | 2 | 3 | 0\|0\|0\|0\|0\|0 | FALSE |
| RFX3 | O75915 | ARL6IP5 | bait_int | 0 | 0.972512164 | 2\|3\|3 | 2.67 | 3 | 0\|0\|0\|0\|0\|0 | FALSE |
| RFX3 | Q07065 | CKAP4 | bait_int | 0 | 0.975229065 | 12\|10\|3 | 8.33 | 3 | 0\|0\|0\|0\|0\|0 | FALSE |
| RFX3 | P52294 | KPNA1 | bait_int | 0 | 0.97113292 | 3\|3\|2 | 2.67 | 3 | 0\|0\|0\|0\|0\|0 | FALSE |
| RFX3 | O75694 | NUP155 | bait_bait | 0 | 0.986020432 | 6\|9\|10 | 8.33 | 3 | 0\|0\|0\|0\|0\|0 | FALSE |
| RFX3 | P08240 | SRPRA | bait_bait | 0 | 0.993055267 | 6\|3\|2 | 3.67 | 3 | 0\|0\|0\|0\|0\|0 | FALSE |
| RFX3 | Q92688 | ANP32B | bait_int | 0 | 0.980974758 | 2\|5\|3 | 3.33 | 3 | 0\|0\|0\|0\|0\|0 | FALSE |
| RFX3 | Q99959 | PKP2 | bait_int | 0.04 | 0.973617638 | 5\|3\|6 | 4.67 | 3 | 0\|0\|1\|1\|2\|2 | FALSE |
| RFX3 | Q9BTT0 | ANP32E | bait_int | 0 | 0.977474668 | 5\|6\|3 | 4.67 | 3 | 0\|0\|0\|0\|0\|0 | FALSE |
| RFX3 | P25787 | PSMA2 | bait_int | 0 | 0.9718468 | 5\|5\|3 | 4.33 | 3 | 0\|0\|0\|0\|0\|0 | FALSE |
| RFX3 | Q14157 | UBAP2L | bait_int | 0 | 0.9739122 | 11\|12\|8 | 10.33 | 3 | 2\|1\|3\|2\|0\|2 | FALSE |
| RFX3 | P78406 | RAE1 | bait_int | 0 | 0.985646165 | 21\|15\|9 | 15 | 3 | 0\|0\|0\|0\|0\|0 | FALSE |
| RFX3 | Q13724 | MOGS | bait_int | 0 | 0.971469068 | 5\|3\|2 | 3.33 | 3 | 0\|0\|0\|0\|0\|0 | FALSE |
| NUP155 | P78406 | RAE1 | bait_int | 0 | 0.973946854 | 5\|6\|3 | 4.67 | 3 | 0\|0\|0\|0\|0\|0 | FALSE |
| NUP155 | Q5T9A4 | ATAD3B | bait_int | 0 | 0.979713339 | 43\|41\|41 | 41.67 | 3 | 0\|0\|0\|0\|0\|0 | FALSE |
| NUP155 | Q9ULW0 | TPX2 | bait_int | 0.05 | 0.972893362 | 12\|20\|5 | 12.33 | 3 | 2\|2\|4\|2\|3\|3 | FALSE |
| NUP155 | Q9P2N5 | RBM27 | bait_int | 0 | 0.988106625 | 12\|13\|7 | 10.67 | 3 | 1\|1\|1\|2\|0\|3 | FALSE |
| NUP155 | Q9UPQ9 | TNRC6B | bait_int | 0 | 0.994503819 | 13\|12\|9 | 11.33 | 3 | 0\|0\|0\|0\|0\|0 | FALSE |
| NUP155 | Q8WUM0 | NUP133 | bait_int | 0 | 0.978659847 | 3\|3\|3 | 3 | 3 | 0\|0\|1\|0\|0\|0 | FALSE |
| NUP155 | Q9NRG9 | AAAS | bait_int | 0 | 0.99825342 | 22\|25\|26 | 24.33 | 3 | 0\|0\|0\|0\|0\|0 | FALSE |
| NUP155 | Q9BY89 | KIAA1671 | bait_int | 0 | 0.981938149 | 3\|6\|6 | 5 | 3 | 0\|0\|0\|0\|0\|0 | FALSE |
| NUP155 | Q99959 | PKP2 | bait_int | 0.04 | 0.973617638 | 3\|6\|5 | 4.67 | 3 | 0\|0\|1\|1\|2\|2 | FALSE |
| NUP155 | Q9UBU9 | NXF1 | bait_int | 0 | 0.990088854 | 16\|17\|16 | 16.33 | 3 | 1\|0\|0\|0\|0\|0 | FALSE |
| NUP155 | P78527 | PRKDC | bait_int | 0 | 0.982769853 | 98\|89\|97 | 94.67 | 3 | 9\|2\|5\|4\|2\|4 | FALSE |
| NUP155 | P51617 | IRAK1 | bait_int | 0 | 0.988439306 | 23\|29\|18 | 23.33 | 3 | 0\|0\|0\|0\|0\|0 | FALSE |
| NUP155 | P49790 | NUP153 | bait_int | 0 | 0.997594988 | 27\|40\|17 | 28 | 3 | 0\|0\|0\|0\|0\|1 | TRUE |
| NUP155 | Q96N67 | DOCK7 | bait_int | 0 | 0.993942418 | 11\|12\|11 | 11.33 | 3 | 0\|0\|0\|0\|1\|1 | FALSE |
| NUP155 | Q9H078 | CLPB | bait_int | 0.02 | 0.974016163 | 4\|7\|2 | 4.33 | 3 | 0\|1\|0\|0\|1\|0 | FALSE |
| NUP155 | Q8WWK9 | CKAP2 | bait_int | 0 | 0.978070723 | 14\|16\|7 | 12.33 | 3 | 0\|0\|0\|1\|1\|1 | FALSE |
| NUP155 | Q9BTX1 | NDC1 | bait_int | 0 | 0.99880789 | 24\|29\|27 | 26.67 | 3 | 0\|0\|0\|0\|0\|0 | FALSE |
| NUP155 | Q9UH99 | SUN2 | bait_int | 0 | 0.993630529 | 7\|5\|7 | 6.33 | 3 | 0\|0\|0\|0\|0\|0 | FALSE |
| NUP155 | Q15058 | KIF14 | bait_int | 0 | 0.99809401 | 25\|22\|21 | 22.67 | 3 | 0\|0\|0\|0\|0\|0 | FALSE |
| NUP155 | Q8NBJ5 | COLGALT1 | bait_int | 0 | 0.988494753 | 3\|3\|3 | 3 | 3 | 0\|0\|0\|0\|0\|0 | FALSE |
| NUP155 | Q9BRJ2 | MRPL45 | bait_int | 0 | 0.989329923 | 6\|7\|6 | 6.33 | 3 | 0\|0\|0\|0\|0\|0 | FALSE |
| NUP155 | A8CG34 | POM121C | bait_int | 0 | 0.989832412 | 2\|4\|3 | 3 | 3 | 0\|0\|0\|0\|0\|0 | FALSE |
| NUP155 | Q8N3R3 | TCAIM | bait_int | 0 | 0.997425181 | 8\|9\|9 | 8.67 | 3 | 0\|0\|0\|0\|0\|0 | FALSE |
| NUP155 | Q8NFH5 | NUP35 | bait_int | 0 | 0.998586102 | 23\|26\|23 | 24 | 3 | 0\|0\|0\|0\|0\|0 | TRUE |
| NUP155 | Q6UUV7 | CRTC3 | bait_int | 0 | 0.990906697 | 3\|5\|2 | 3.33 | 3 | 0\|0\|0\|0\|0\|0 | FALSE |
| NUP155 | Q9NXE4 | SMPD4 | bait_int | 0 | 0.997976186 | 20\|26\|38 | 28 | 3 | 0\|0\|0\|0\|0\|0 | FALSE |

| | | | | | | | | | | |
|---|---|---|---|---|---|---|---|---|---|---|
| NUP155 | Q5SNT2 | TMEM201 | bait_int | 0 | 0.98131437 | 4\|5\|9 | 6 | 3 | 0\|0\|0\|0\|0\|0 | FALSE |
| NUP155 | Q92621 | NUP205 | bait_int | 0 | 0.979415312 | 15\|16\|7 | 12.67 | 3 | 0\|0\|0\|0\|0\|0 | FALSE |
| NUP155 | O60568 | PLOD3 | bait_int | 0.01 | 0.979685616 | 2\|2\|9 | 4.33 | 3 | 0\|0\|0\|0\|0\|0 | FALSE |
| NUP155 | Q9NRA8 | EIF4ENIF1 | bait_int | 0.01 | 0.989770034 | 8\|8\|10 | 8.67 | 3 | 1\|0\|4\|3\|1\|3 | FALSE |
| NUP155 | Q96SK2 | TMEM209 | bait_int | 0 | 0.987877906 | 11\|14\|7 | 10.67 | 3 | 0\|0\|0\|0\|0\|0 | FALSE |
| NUP155 | P52948 | NUP98 | bait_int | 0 | 0.98767691 | 4\|5\|2 | 3.67 | 3 | 0\|0\|0\|0\|0\|0 | TRUE |
| NUP155 | Q7Z4S6 | KIF21A | bait_int | 0 | 0.99356122 | 7\|8\|14 | 9.67 | 3 | 0\|0\|0\|0\|0\|0 | FALSE |
| NUP155 | P39060 | COL18A1 | bait_int | 0 | 0.998121734 | 19\|18\|22 | 19.67 | 3 | 0\|0\|0\|0\|0\|0 | FALSE |
| NUP155 | Q6IQ23 | PLEKHA7 | bait_int | 0 | 0.996666251 | 17\|14\|28 | 19.67 | 3 | 0\|0\|0\|0\|0\|0 | FALSE |
| KMT5B | Q5TAQ9 | DCAF8 | bait_int | 0 | 0.977772695 | 2\|2\|2 | 2 | 3 | 0\|0\|0\|0\|0\|0 | FALSE |
| KMT5B | Q13617 | CUL2 | bait_int | 0 | 0.975617194 | 3\|2\|4 | 3 | 3 | 0\|0\|0\|0\|0\|0 | FALSE |
| KMT5B | Q96JK2 | DCAF5 | bait_int | 0 | 0.993166161 | 4\|3\|2 | 3 | 3 | 0\|0\|0\|0\|0\|0 | FALSE |
| KMT5B | P48634 | PRRC2A | bait_int | 0.02 | 0.987212542 | 4\|2\|4 | 3.33 | 3 | 0\|0\|2\|0\|0\|1 | FALSE |
| KMT5B | P17655 | CAPN2 | bait_int | 0 | 0.974840936 | 5\|5\|5 | 5 | 3 | 0\|0\|0\|0\|0\|0 | FALSE |
| KMT5B | Q93009 | USP7 | bait_int | 0 | 0.977301396 | 36\|41\|22 | 33 | 3 | 0\|0\|0\|0\|0\|0 | FALSE |
| KMT5B | Q16531 | DDB1 | bait_int | 0 | 0.971306192 | 17\|14\|9 | 13.33 | 3 | 0\|0\|0\|0\|0\|0 | FALSE |
| KMT5B | P13489 | RNH1 | bait_int | 0 | 0.982513411 | 11\|9\|4 | 8 | 3 | 0\|0\|0\|0\|0\|0 | FALSE |
| KMT5B | O94906 | PRPF6 | bait_int | 0 | 0.98576399 | 7\|6\|7 | 6.67 | 3 | 0\|0\|0\|0\|0\|0 | FALSE |
| AP2S1 | Q13724 | MOGS | bait_int | 0 | 0.971469068 | 4\|4\|2 | 3.33 | 3 | 0\|0\|0\|0\|0\|0 | FALSE |
| AP2S1 | Q8NFH3 | NUP43 | bait_int | 0 | 0.97844499 | 3\|3\|3 | 3 | 3 | 0\|0\|0\|0\|0\|0 | FALSE |
| AP2S1 | Q9BRK5 | SDF4 | bait_int | 0 | 0.972248791 | 9\|9\|9 | 9 | 3 | 0\|0\|0\|0\|0\|0 | FALSE |
| AP2S1 | O95782 | AP2A1 | bait_int | 0 | 0.993990934 | 34\|36\|40 | 36.67 | 3 | 0\|0\|0\|0\|0\|0 | TRUE |
| AP2S1 | Q9UJS0 | SLC25A13 | bait_int | 0 | 0.972671574 | 21\|10\|10 | 13.67 | 3 | 0\|0\|0\|0\|0\|0 | FALSE |
| AP2S1 | P63010 | AP2B1 | bait_int | 0 | 0.988723472 | 10\|12\|22 | 14.67 | 3 | 0\|0\|0\|0\|0\|0 | TRUE |
| AP2S1 | Q7Z6Z7 | HUWE1 | bait_int | 0 | 0.984052065 | 21\|20\|7 | 16 | 3 | 0\|0\|0\|0\|0\|0 | FALSE |
| AP2S1 | Q96CW1 | AP2M1 | bait_int | 0 | 0.985542202 | 11\|11\|20 | 14 | 3 | 0\|0\|0\|0\|0\|0 | TRUE |
| AP2S1 | P08240 | SRPRA | bait_bait | 0 | 0.99674249 | 21\|8\|4 | 11 | 3 | 0\|0\|0\|0\|0\|0 | FALSE |
| AP2S1 | P09543 | CNP | bait_int | 0 | 0.972886431 | 13\|9\|9 | 10.33 | 3 | 0\|0\|0\|0\|0\|0 | FALSE |
| AP2S1 | Q9Y5M8 | SRPRB | bait_int | 0 | 0.98003909 | 23\|15\|18 | 18.67 | 3 | 0\|0\|0\|0\|0\|0 | FALSE |
| AP2S1 | O75821 | EIF3G | bait_bait | 0.01 | 0.990033407 | 6\|3\|3 | 4 | 3 | 2\|0\|0\|0\|0\|1 | FALSE |
| AP2S1 | P51570 | GALK1 | bait_int | 0 | 0.97892322 | 6\|4\|4 | 4.67 | 3 | 0\|0\|0\|0\|0\|0 | FALSE |
| AP2S1 | Q07065 | CKAP4 | bait_int | 0 | 0.980545044 | 17\|15\|8 | 13.33 | 3 | 0\|0\|0\|0\|0\|0 | FALSE |
| AP2S1 | Q9BW92 | TARS2 | bait_int | 0 | 0.977751903 | 2\|5\|2 | 3 | 3 | 0\|0\|0\|0\|0\|0 | FALSE |
| AP2S1 | Q14204 | DYNC1H1 | bait_bait | 0.01 | 0.992653276 | 5\|8\|3 | 5.33 | 3 | 0\|0\|0\|1\|1\|1 | FALSE |
| AP2S1 | Q6PD74 | AAGAB | bait_int | 0 | 0.997525679 | 11\|13\|13 | 12.33 | 3 | 0\|0\|0\|0\|0\|0 | TRUE |
| AP2S1 | Q7Z2K6 | ERMP1 | bait_int | 0 | 0.99626426 | 5\|5\|6 | 5.33 | 3 | 0\|0\|0\|0\|0\|0 | FALSE |
| AP2S1 | Q86VU5 | COMTD1 | bait_int | 0 | 0.982839162 | 4\|7\|4 | 5 | 3 | 0\|0\|0\|0\|0\|0 | FALSE |
| AP2S1 | Q53S58 | TMEM177 | bait_int | 0 | 0.975679572 | 2\|2\|2 | 2 | 3 | 0\|0\|0\|0\|0\|0 | FALSE |
| AP2S1 | Q13445 | TMED1 | bait_int | 0 | 0.97355526 | 2\|3\|2 | 2.33 | 3 | 0\|0\|0\|0\|0\|0 | FALSE |
| AP2S1 | Q8WWC4 | MAIP1 | bait_int | 0 | 0.972910689 | 8\|5\|6 | 6.33 | 3 | 0\|0\|0\|0\|0\|0 | FALSE |
| AP2S1 | Q8TED1 | GPX8 | bait_int | 0 | 0.986540248 | 9\|7\|5 | 7 | 3 | 0\|0\|0\|0\|0\|0 | FALSE |
| AP2S1 | Q8IXI1 | RHOT2 | bait_int | 0 | 0.980572767 | 8\|6\|6 | 6.67 | 3 | 0\|0\|0\|0\|0\|0 | FALSE |
| AP2S1 | Q8NBN7 | RDH13 | bait_int | 0 | 0.997186066 | 16\|28\|24 | 22.67 | 3 | 0\|0\|0\|0\|0\|0 | FALSE |
| AP2S1 | Q8TEB1 | DCAF11 | bait_int | 0 | 0.99626426 | 9\|3\|4 | 5.33 | 3 | 0\|0\|0\|0\|0\|0 | FALSE |
| AP2S1 | Q13232 | NME3 | bait_int | 0 | 0.981598536 | 7\|3\|5 | 5 | 3 | 0\|0\|0\|0\|0\|0 | FALSE |
| AP2S1 | Q10567 | AP1B1 | bait_int | 0 | 0.990795803 | 11\|7\|11 | 9.67 | 3 | 0\|0\|0\|0\|0\|0 | TRUE |
| AP2S1 | O75688 | PPM1B | bait_int | 0 | 0.991461166 | 6\|7\|8 | 7 | 3 | 0\|0\|0\|0\|0\|0 | FALSE |
| AP2S1 | O43929 | ORC4 | bait_int | 0 | 0.994122621 | 2\|8\|4 | 4.67 | 3 | 0\|0\|0\|0\|0\|0 | FALSE |
| AP2S1 | O94973 | AP2A2 | bait_int | 0 | 0.995550381 | 51\|46\|56 | 51 | 3 | 0\|0\|0\|0\|0\|0 | TRUE |
| AP2S1 | A0FGR8 | ESYT2 | bait_int | 0 | 0.972234929 | 9\|9\|7 | 8.33 | 3 | 0\|0\|0\|0\|0\|0 | FALSE |
| AP2S1 | O14975 | SLC27A2 | bait_int | 0 | 0.992639414 | 3\|6\|5 | 4.67 | 3 | 0\|0\|0\|0\|0\|0 | FALSE |
| AP2S1 | O15173 | PGRMC2 | bait_int | 0 | 0.972532956 | 6\|4\|3 | 4.33 | 3 | 0\|0\|0\|0\|0\|0 | FALSE |
| AP2S1 | O43169 | CYB5B | bait_int | 0 | 0.972526025 | 5\|4\|4 | 4.33 | 3 | 0\|0\|0\|0\|0\|0 | FALSE |
| AP2S1 | P10155 | RO60 | bait_int | 0 | 0.989624485 | 5\|2\|2 | 3 | 3 | 0\|0\|0\|0\|0\|0 | FALSE |
| AP2S1 | P11172 | UMPS | bait_int | 0 | 0.988425445 | 11\|9\|10 | 10 | 3 | 0\|0\|0\|0\|0\|0 | FALSE |
| AP2S1 | P17066 | HSPA6 | bait_int | 0 | 0.992091876 | 35\|23\|25 | 27.67 | 3 | 0\|0\|0\|0\|0\|0 | FALSE |
| AP2S1 | P42566 | EPS15 | bait_int | 0 | 0.996326638 | 6\|9\|20 | 11.67 | 3 | 0\|0\|0\|0\|0\|0 | TRUE |
| AP2S1 | Q96TA2 | YME1L1 | bait_int | 0 | 0.982250038 | 14\|13\|9 | 12 | 3 | 0\|0\|0\|0\|0\|0 | FALSE |
| AP2S1 | Q9HBH5 | RDH14 | bait_int | 0 | 0.996936555 | 11\|5\|5 | 7 | 3 | 0\|0\|0\|0\|0\|0 | FALSE |
| AP2S1 | Q9HC21 | SLC25A19 | bait_int | 0 | 0.986096672 | 5\|2\|5 | 4 | 3 | 0\|0\|0\|0\|0\|0 | FALSE |
| AP2S1 | Q9NP72 | RAB18 | bait_int | 0 | 0.984273853 | 6\|2\|2 | 3.33 | 3 | 0\|0\|0\|0\|0\|0 | FALSE |
| AP2S1 | Q9H4I3 | TRABD | bait_int | 0 | 0.988030385 | 3\|4\|3 | 3.33 | 3 | 0\|0\|0\|0\|0\|0 | FALSE |
| AP2S1 | Q99707 | MTR | bait_int | 0 | 0.995099874 | 5\|8\|16 | 9.67 | 3 | 0\|0\|0\|0\|0\|0 | FALSE |
| AP2S1 | Q9BV29 | CCDC32 | bait_int | 0 | 0.997501421 | 7\|9\|11 | 9 | 3 | 0\|0\|0\|0\|0\|0 | FALSE |
| AP2S1 | Q9H4P4 | RNF41 | bait_int | 0 | 0.998392038 | 18\|18\|20 | 18.67 | 3 | 0\|0\|0\|0\|0\|0 | TRUE |
| AP2S1 | Q9UBC2 | EPS15L1 | bait_int | 0 | 0.991613645 | 7\|4\|20 | 10.33 | 3 | 0\|0\|0\|0\|0\|0 | TRUE |
| AP2S1 | Q9Y2W6 | TDRKH | bait_int | 0 | 0.994053312 | 3\|3\|2 | 2.67 | 3 | 0\|0\|0\|0\|0\|0 | FALSE |
| AP2S1 | Q9Y4W6 | AFG3L2 | bait_int | 0 | 0.986325391 | 12\|16\|13 | 13.67 | 3 | 0\|0\|0\|0\|0\|0 | FALSE |
| CORO1A | P35222 | CTNNB1 | bait_bait | 0 | 0.994781054 | 8\|2\|8 | 6 | 3 | 0\|0\|0\|0\|0\|0 | FALSE |
| CORO1A | P28290 | ITPRID2 | bait_int | 0 | 0.986845206 | 35\|19\|34 | 29.33 | 3 | 0\|0\|0\|0\|0\|0 | FALSE |
| CORO1A | P53355 | DAPK1 | bait_int | 0 | 0.974057748 | 2\|2\|9 | 4.33 | 3 | 0\|0\|0\|0\|0\|0 | FALSE |
| CORO1A | P07197 | NEFM | bait_int | 0.05 | 0.979463828 | 29\|1\|10 | 13.33 | 3 | 0\|0\|0\|0\|0\|0 | FALSE |

| | | | | | | | | | | |
|---|---|---|---|---|---|---|---|---|---|---|
| CORO1A | P12814 | ACTN1 | bait_int | 0 | 0.981439126 | 20\|2\|23 | 15 | 3 | 0\|0\|0\|0\|1\|0 | FALSE |
| CORO1A | Q14573 | ITPR3 | bait_int | 0 | 0.990878973 | 27\|8\|36 | 23.67 | 3 | 0\|0\|0\|0\|0\|0 | FALSE |
| CORO1A | Q14571 | ITPR2 | bait_int | 0 | 0.992563175 | 45\|15\|58 | 39.33 | 3 | 0\|0\|0\|0\|0\|0 | FALSE |
| CORO1A | Q14254 | FLOT2 | bait_int | 0 | 0.9734409 | 6\|4\|16 | 8.67 | 3 | 0\|0\|0\|0\|0\|0 | FALSE |
| CORO1A | Q5VT25 | CDC42BPA | bait_int | 0 | 0.980066814 | 8\|16\|37 | 20.33 | 3 | 0\|0\|0\|0\|0\|0 | FALSE |
| CORO1A | Q69YQ0 | SPECC1L | bait_int | 0 | 0.992833479 | 66\|28\|62 | 52 | 3 | 0\|0\|0\|0\|0\|0 | FALSE |
| CORO1A | Q5M775 | SPECC1 | bait_int | 0 | 0.984953078 | 3\|9\|26 | 12.67 | 3 | 0\|0\|0\|0\|0\|0 | FALSE |
| CORO1A | P67936 | TPM4 | bait_int | 0 | 0.9755063 | 4\|3\|4 | 3.67 | 3 | 0\|0\|0\|0\|0\|0 | FALSE |
| CORO1A | Q13045 | FLII | bait_int | 0 | 0.983185706 | 4\|4\|16 | 8 | 3 | 0\|0\|0\|0\|0\|0 | FALSE |
| CORO1A | Q12792 | TWF1 | bait_int | 0 | 0.975922152 | 9\|2\|4 | 5 | 3 | 0\|0\|0\|0\|0\|0 | FALSE |
| CORO1A | Q07157 | TJP1 | bait_int | 0 | 0.988279896 | 51\|8\|30 | 29.67 | 3 | 0\|0\|0\|0\|0\|0 | FALSE |
| CORO1A | Q9UHB6 | LIMA1 | bait_int | 0 | 0.976324143 | 88\|46\|96 | 76.67 | 3 | 0\|0\|0\|0\|0\|0 | FALSE |
| CORO1A | O75955 | FLOT1 | bait_int | 0 | 0.974667665 | 2\|5\|15 | 7.33 | 3 | 0\|0\|0\|0\|0\|0 | FALSE |
| CORO1A | O60784 | TOM1 | bait_int | 0 | 0.979969781 | 10\|2\|4 | 5.33 | 3 | 0\|0\|0\|0\|0\|0 | FALSE |
| CORO1A | O14974 | PPP1R12A | bait_int | 0 | 0.988058108 | 30\|27\|41 | 32.67 | 3 | 0\|0\|0\|0\|0\|0 | FALSE |
| CORO1A | O15020 | SPTBN2 | bait_int | 0 | 0.982541135 | 15\|9\|32 | 18.67 | 3 | 0\|0\|0\|0\|0\|0 | FALSE |
| CORO1A | Q9NTK5 | OLA1 | bait_int | 0 | 0.984634258 | 6\|3\|7 | 5.33 | 3 | 0\|0\|0\|0\|0\|0 | FALSE |
| CORO1A | Q5SW79 | CEP170 | bait_int | 0 | 0.994136483 | 10\|13\|5 | 9.33 | 3 | 0\|0\|0\|0\|0\|0 | FALSE |
| CORO1A | P26640 | VARS1 | bait_int | 0 | 0.98600657 | 10\|14\|5 | 9.67 | 3 | 0\|0\|0\|0\|0\|0 | FALSE |
| CORO1A | Q96SB3 | PPP1R9B | bait_bait | 0 | 0.996811799 | 30\|8\|26 | 21.33 | 3 | 0\|0\|0\|0\|0\|0 | FALSE |
| CORO1A | Q99959 | PKP2 | bait_int | 0 | 0.981099513 | 9\|4\|14 | 9 | 3 | 0\|0\|0\|1\|0\|0 | FALSE |
| CORO1A | Q9BR76 | CORO1B | bait_int | 0 | 0.994794916 | 25\|25\|21 | 23.67 | 3 | 0\|0\|0\|0\|0\|0 | TRUE |
| CORO1A | Q9NQX4 | MYO5C | bait_int | 0 | 0.988182864 | 21\|2\|13 | 12 | 3 | 0\|0\|0\|0\|0\|0 | FALSE |
| CORO1A | Q96PY5 | FMNL2 | bait_int | 0 | 0.986865999 | 7\|7\|21 | 11.67 | 3 | 0\|0\|0\|0\|0\|0 | FALSE |
| CORO1A | Q7Z406 | MYH14 | bait_int | 0 | 0.996804868 | 90\|38\|52 | 60 | 3 | 0\|0\|0\|0\|0\|0 | FALSE |
| CORO1A | Q96IZ0 | PAWR | bait_int | 0 | 0.983719383 | 10\|3\|8 | 7 | 3 | 0\|0\|0\|0\|0\|0 | FALSE |
| CORO1A | Q92614 | MYO18A | bait_int | 0 | 0.991090365 | 21\|8\|18 | 15.67 | 3 | 0\|0\|0\|0\|0\|0 | FALSE |
| CORO1A | Q9ULJ8 | PPP1R9A | bait_int | 0 | 0.992389903 | 20\|2\|13 | 11.67 | 3 | 0\|0\|0\|0\|0\|0 | FALSE |
| CORO1A | Q9UM54 | MYO6 | bait_int | 0 | 0.990116577 | 85\|16\|34 | 45 | 3 | 0\|0\|0\|0\|0\|0 | FALSE |
| CORO1A | Q9P0K7 | RAI14 | bait_int | 0 | 0.994413718 | 47\|45\|78 | 56.67 | 3 | 0\|0\|0\|0\|0\|0 | FALSE |
| CORO1A | Q9UEY8 | ADD3 | bait_int | 0 | 0.976781581 | 7\|3\|5 | 5 | 3 | 0\|0\|0\|0\|0\|0 | FALSE |
| CORO1A | Q9Y608 | LRRFIP2 | bait_int | 0 | 0.983005503 | 4\|3\|10 | 5.67 | 3 | 0\|0\|0\|0\|0\|0 | FALSE |
| CORO1A | Q9Y4I1 | MYO5A | bait_int | 0 | 0.985306552 | 19\|2\|7 | 9.33 | 3 | 0\|0\|0\|0\|0\|0 | FALSE |
| CORO1A | Q00610 | CLTC | bait_int | 0 | 0.975728088 | 25\|16\|24 | 21.67 | 3 | 0\|0\|0\|0\|0\|0 | FALSE |
| CORO1A | Q14160 | SCRIB | bait_int | 0 | 0.981910425 | 3\|3\|17 | 7.67 | 3 | 0\|0\|0\|0\|0\|0 | FALSE |
| CORO1A | Q01082 | SPTBN1 | bait_int | 0 | 0.983462941 | 95\|63\|114 | 90.67 | 3 | 0\|0\|0\|0\|0\|0 | FALSE |
| CORO1A | P21333 | FLNA | bait_int | 0 | 0.974175573 | 34\|7\|31 | 24 | 3 | 0\|1\|2\|0\|1\|0 | FALSE |
| CORO1A | P35580 | MYH10 | bait_int | 0 | 0.992237424 | 246\|205\|25 | 236.33 | 3 | 0\|0\|0\|0\|0\|0 | FALSE |
| CORO1A | P35579 | MYH9 | bait_int | 0 | 0.98981855 | 128\|97\|158 | 127.67 | 3 | 1\|0\|0\|0\|0\|0 | FALSE |
| CORO1A | P35221 | CTNNA1 | bait_int | 0 | 0.99022054 | 10\|6\|17 | 11 | 3 | 0\|0\|0\|0\|0\|0 | FALSE |
| CORO1A | Q96N67 | DOCK7 | bait_int | 0 | 0.99435134 | 32\|2\|3 | 12.33 | 3 | 0\|0\|0\|0\|0\|0 | FALSE |
| CORO1A | Q8WWI1 | LMO7 | bait_int | 0 | 0.988175933 | 27\|22\|52 | 33.67 | 3 | 0\|0\|0\|0\|0\|0 | FALSE |
| CORO1A | Q15149 | PLEC | bait_int | 0 | 0.992909718 | 61\|2\|16 | 26.33 | 3 | 0\|0\|1\|0\|0\|0 | FALSE |
| CORO1A | Q16643 | DBN1 | bait_int | 0 | 0.971985417 | 96\|51\|99 | 82 | 3 | 0\|0\|0\|0\|0\|0 | FALSE |
| CORO1A | Q13813 | SPTAN1 | bait_int | 0 | 0.984558018 | 113\|72\|143 | 109.33 | 3 | 0\|0\|0\|0\|0\|0 | FALSE |
| CORO1A | Q14204 | DYNC1H1 | bait_bait | 0.01 | 0.994171137 | 5\|3\|14 | 7.33 | 3 | 0\|0\|0\|1\|1\|1 | FALSE |
| CORO1A | Q6WCQ1 | MPRIP | bait_int | 0 | 0.992930511 | 49\|44\|72 | 55 | 3 | 0\|0\|0\|0\|0\|0 | FALSE |
| CORO1A | O43707 | ACTN4 | bait_int | 0 | 0.973094357 | 41\|16\|47 | 34.67 | 3 | 0\|0\|0\|0\|0\|0 | FALSE |
| CORO1A | O43795 | MYO1B | bait_int | 0 | 0.974660734 | 64\|58\|77 | 66.33 | 3 | 0\|0\|0\|0\|0\|0 | FALSE |
| CORO1A | O95425 | SVIL | bait_int | 0 | 0.98059356 | 18\|5\|20 | 14.33 | 3 | 0\|0\|0\|0\|0\|0 | FALSE |
| CORO1A | O75083 | WDR1 | bait_int | 0 | 0.974393895 | 17\|10\|26 | 17.67 | 3 | 0\|0\|0\|0\|0\|0 | FALSE |
| CORO1A | O75369 | FLNB | bait_int | 0 | 0.990449259 | 91\|33\|89 | 71 | 3 | 0\|0\|0\|0\|0\|0 | FALSE |
| CORO1A | Q9ULV4 | CORO1C | bait_int | 0 | 0.973073564 | 51\|45\|45 | 47 | 3 | 0\|0\|0\|0\|0\|0 | TRUE |
| CORO1A | Q9P2M7 | CGN | bait_int | 0.05 | 0.980489597 | 16\|0\|5 | 7 | 3 | 0\|0\|0\|0\|0\|0 | FALSE |
| CORO1A | Q14643 | ITPR1 | bait_int | 0 | 0.994628575 | 9\|4\|14 | 9 | 3 | 0\|0\|0\|0\|0\|0 | FALSE |
| CORO1A | Q15154 | PCM1 | bait_int | 0 | 0.989250218 | 16\|2\|5 | 7.67 | 3 | 0\|0\|0\|0\|0\|0 | FALSE |
| CORO1A | Q8NBT0 | POC1A | bait_int | 0 | 0.997487559 | 9\|14\|9 | 10.67 | 3 | 0\|0\|0\|0\|0\|0 | TRUE |
| GFAP | Q9H2C0 | GAN | bait_int | 0.05 | 0.994954326 | 1\|10\|9 | 6.67 | 3 | 0\|0\|0\|0\|0\|0 | FALSE |
| GFAP | Q9Y5X1 | SNX9 | bait_int | 0 | 0.992985958 | 36\|8\|6 | 16.67 | 3 | 0\|0\|0\|0\|0\|0 | FALSE |
| GFAP | O15027 | SEC16A | bait_int | 0 | 0.990761148 | 6\|13\|12 | 10.33 | 3 | 0\|0\|0\|0\|0\|0 | FALSE |
| GFAP | O75153 | CLUH | bait_int | 0 | 0.992251286 | 4\|3\|4 | 3.67 | 3 | 0\|0\|0\|0\|0\|0 | FALSE |
| GFAP | Q5JSZ5 | PRRC2B | bait_int | 0.01 | 0.989076947 | 5\|6\|4 | 5 | 3 | 0\|2\|0\|0\|2\|2 | FALSE |
| GFAP | P48634 | PRRC2A | bait_int | 0.01 | 0.987212542 | 3\|4\|3 | 3.33 | 3 | 0\|0\|2\|0\|0\|1 | FALSE |
| GFAP | P78527 | PRKDC | bait_int | 0 | 0.973427039 | 44\|46\|29 | 39.67 | 3 | 1\|1\|0\|0\|0\|0 | FALSE |
| GFAP | Q10713 | PMPCA | bait_int | 0 | 0.974653803 | 11\|5\|3 | 6.33 | 3 | 0\|0\|0\|0\|1\|0 | FALSE |
| GFAP | Q16352 | INA | bait_int | 0 | 0.977734575 | 3\|4\|4 | 3.67 | 3 | 0\|0\|0\|0\|0\|0 | TRUE |
| GFAP | P07197 | NEFM | bait_int | 0 | 0.977218225 | 13\|8\|12 | 11 | 3 | 0\|0\|0\|0\|0\|0 | TRUE |
| GFAP | P07196 | NEFL | bait_int | 0 | 0.982575789 | 7\|3\|3 | 4.33 | 3 | 0\|0\|0\|0\|0\|0 | TRUE |
| ELAVL3 | Q96DH6 | MSI2 | bait_int | 0 | 0.993977073 | 10\|11\|12 | 11 | 3 | 0\|0\|0\|0\|0\|0 | FALSE |
| ELAVL3 | Q13325 | IFIT5 | bait_int | 0 | 0.995973164 | 5\|9\|9 | 7.67 | 3 | 0\|0\|0\|0\|0\|0 | FALSE |
| ELAVL3 | Q9BV44 | THUMPD3 | bait_int | 0 | 0.992577037 | 4\|6\|7 | 5.67 | 3 | 0\|0\|0\|0\|0\|0 | FALSE |
| ELAVL3 | O14561 | NDUFAB1 | bait_int | 0 | 0.993824594 | 9\|8\|6 | 7.67 | 3 | 0\|0\|0\|0\|0\|0 | FALSE |

| | | | | | | | | | | |
|---|---|---|---|---|---|---|---|---|---|---|
| ELAVL3 | Q9Y5J7 | TIMM9 | bait_int | 0 | 0.979145008 | 6\|5\|5 | 5.33 | 3 | 0\|0\|0\|0\|0\|0 | FALSE |
| ELAVL3 | Q96EB6 | SIRT1 | bait_int | 0 | 0.996652389 | 21\|26\|25 | 24 | 3 | 0\|0\|0\|0\|0\|0 | FALSE |
| ELAVL3 | O43347 | MSI1 | bait_int | 0 | 0.989908651 | 3\|8\|5 | 5.33 | 3 | 0\|0\|0\|0\|0\|0 | FALSE |
| ELAVL3 | O15212 | PFDN6 | bait_int | 0 | 0.975672641 | 3\|3\|2 | 2.67 | 3 | 0\|0\|0\|0\|0\|0 | FALSE |
| ELAVL3 | Q969Y2 | GTPBP3 | bait_int | 0 | 0.998426692 | 27\|27\|33 | 29 | 3 | 0\|0\|0\|0\|0\|0 | FALSE |
| ELAVL3 | P10155 | RO60 | bait_int | 0 | 0.998655411 | 53\|61\|59 | 57.67 | 3 | 0\|0\|0\|0\|0\|0 | FALSE |
| ELAVL3 | P52298 | NCBP2 | bait_int | 0 | 0.98942349 | 3\|2\|3 | 2.67 | 3 | 0\|0\|0\|0\|0\|0 | FALSE |
| ELAVL3 | Q32P28 | P3H1 | bait_int | 0 | 0.993512704 | 45\|34\|42 | 40.33 | 3 | 0\|0\|0\|0\|0\|0 | FALSE |
| ELAVL3 | Q99471 | PFDN5 | bait_int | 0 | 0.974750835 | 4\|2\|2 | 2.67 | 3 | 0\|0\|0\|0\|0\|0 | FALSE |
| ELAVL3 | Q15020 | SART3 | bait_int | 0 | 0.996132574 | 20\|20\|16 | 18.67 | 3 | 0\|0\|0\|0\|0\|0 | FALSE |
| ELAVL3 | O75718 | CRTAP | bait_int | 0 | 0.991225517 | 20\|21\|21 | 20.67 | 3 | 0\|0\|0\|0\|0\|0 | FALSE |
| ELAVL3 | Q9NXA8 | SIRT5 | bait_int | 0 | 0.997903411 | 11\|10\|15 | 12 | 3 | 0\|0\|0\|0\|0\|0 | FALSE |
| ELAVL3 | Q8N5D0 | WDTC1 | bait_int | 0 | 0.996402878 | 5\|14\|8 | 9 | 3 | 0\|0\|0\|0\|0\|0 | FALSE |
| ELAVL3 | Q7L8W6 | DPH6 | bait_int | 0 | 0.997702416 | 12\|9\|9 | 10 | 3 | 0\|0\|0\|0\|0\|0 | FALSE |
| ELAVL3 | Q6DKK2 | TTC19 | bait_int | 0 | 0.997026656 | 9\|9\|15 | 11 | 3 | 0\|0\|0\|0\|0\|0 | FALSE |
| ELAVL3 | Q12926 | ELAVL2 | bait_int | 0 | 0.997584591 | 11\|10\|10 | 10.33 | 3 | 0\|0\|0\|0\|0\|0 | TRUE |
| ELAVL3 | O60220 | TIMM8A | bait_int | 0 | 0.971562634 | 12\|12\|12 | 12 | 3 | 0\|0\|0\|0\|0\|0 | FALSE |
| ELAVL3 | Q86X55 | CARM1 | bait_int | 0 | 0.981189615 | 36\|37\|39 | 37.33 | 3 | 0\|0\|0\|0\|0\|2 | FALSE |
| ELAVL3 | Q8TEQ6 | GEMIN5 | bait_int | 0 | 0.986068948 | 3\|19\|6 | 9.33 | 3 | 0\|0\|0\|0\|0\|0 | FALSE |
| ELAVL3 | Q9HCE1 | MOV10 | bait_int | 0 | 0.972976532 | 7\|10\|6 | 7.67 | 3 | 0\|0\|0\|0\|0\|0 | FALSE |
| ELAVL3 | Q9H9Y6 | POLR1B | bait_int | 0 | 0.996763283 | 12\|13\|15 | 13.33 | 3 | 0\|0\|0\|0\|0\|0 | FALSE |
| ELAVL3 | Q15293 | RCN1 | bait_int | 0 | 0.974619149 | 6\|7\|7 | 6.67 | 3 | 0\|0\|0\|0\|0\|0 | FALSE |
| ELAVL3 | Q8IZH2 | XRN1 | bait_int | 0.05 | 0.994385994 | 1\|11\|12 | 8 | 3 | 0\|0\|0\|0\|0\|0 | FALSE |
| ELAVL3 | Q09161 | NCBP1 | bait_int | 0 | 0.984883769 | 7\|4\|3 | 4.67 | 3 | 0\|0\|0\|0\|0\|0 | FALSE |
| ELAVL3 | Q9BYD3 | MRPL4 | bait_int | 0 | 0.971236883 | 2\|3\|3 | 2.67 | 3 | 0\|0\|0\|0\|1\|0 | FALSE |
| ELAVL3 | O15355 | PPM1G | bait_int | 0 | 0.978971736 | 23\|24\|25 | 24 | 3 | 0\|0\|0\|0\|0\|0 | FALSE |
| ELAVL3 | O43395 | PRPF3 | bait_int | 0 | 0.988744265 | 7\|2\|3 | 4 | 3 | 0\|0\|0\|0\|0\|0 | FALSE |
| ELAVL3 | Q9Y5J9 | TIMM8B | bait_int | 0 | 0.972796329 | 12\|13\|14 | 13 | 3 | 0\|0\|0\|0\|0\|0 | FALSE |
| ELAVL3 | Q9NXV6 | CDKN2AIP | bait_int | 0.01 | 0.980586629 | 3\|6\|4 | 4.33 | 3 | 0\|0\|0\|2\|0\|1 | FALSE |
| ELAVL3 | P62072 | TIMM10 | bait_int | 0 | 0.971992348 | 6\|5\|5 | 5.33 | 3 | 0\|0\|0\|0\|0\|0 | FALSE |
| PPP1R9B | P28289 | TMOD1 | bait_int | 0.05 | 0.975000347 | 1\|8\|5 | 4.67 | 3 | 0\|0\|0\|0\|0\|0 | FALSE |
| PPP1R9B | P46940 | IQGAP1 | bait_int | 0 | 0.971430948 | 2\|4\|3 | 3 | 3 | 0\|0\|0\|0\|0\|0 | FALSE |
| PPP1R9B | P12814 | ACTN1 | bait_int | 0 | 0.986928376 | 11\|41\|36 | 29.33 | 3 | 0\|0\|0\|0\|1\|0 | TRUE |
| PPP1R9B | Q69YQ0 | SPECC1L | bait_int | 0 | 0.974889453 | 2\|10\|5 | 5.67 | 3 | 0\|0\|0\|0\|0\|0 | FALSE |
| PPP1R9B | Q13555 | CAMK2G | bait_int | 0 | 0.983968894 | 2\|3\|3 | 2.67 | 3 | 0\|0\|0\|0\|0\|0 | FALSE |
| PPP1R9B | Q9Y490 | TLN1 | bait_int | 0 | 0.97932521 | 4\|18\|14 | 12 | 3 | 0\|0\|0\|0\|0\|0 | FALSE |
| PPP1R9B | Q86VM9 | ZC3H18 | bait_int | 0 | 0.990899766 | 6\|11\|12 | 9.67 | 3 | 0\|0\|0\|0\|0\|0 | FALSE |
| PPP1R9B | Q7Z406 | MYH14 | bait_int | 0 | 0.978334096 | 3\|2\|3 | 2.67 | 3 | 0\|0\|0\|0\|0\|0 | FALSE |
| PPP1R9B | Q86VP6 | CAND1 | bait_int | 0.05 | 0.975086982 | 1\|10\|10 | 7 | 3 | 0\|0\|0\|0\|0\|0 | FALSE |
| PPP1R9B | P28838 | LAP3 | bait_int | 0 | 0.990206679 | 5\|15\|12 | 10.67 | 3 | 0\|0\|0\|0\|0\|0 | FALSE |
| PPP1R9B | Q14008 | CKAP5 | bait_int | 0.05 | 0.977530115 | 1\|15\|13 | 9.67 | 3 | 0\|0\|0\|0\|0\|0 | FALSE |
| PPP1R9B | Q01082 | SPTBN1 | bait_int | 0 | 0.971548773 | 18\|39\|36 | 31 | 3 | 0\|0\|0\|0\|0\|0 | FALSE |
| PPP1R9B | P35580 | MYH10 | bait_int | 0 | 0.988453168 | 76\|146\|152 | 124.67 | 3 | 0\|0\|0\|0\|0\|0 | FALSE |
| PPP1R9B | O43707 | ACTN4 | bait_int | 0 | 0.975145895 | 28\|49\|45 | 40.67 | 3 | 0\|0\|0\|0\|0\|0 | FALSE |
| PPP1R9B | Q96GY0 | ZC2HC1A | bait_int | 0 | 0.984357023 | 2\|6\|3 | 3.67 | 3 | 0\|0\|0\|0\|0\|0 | FALSE |
| PPP1R9B | Q9NZQ3 | NCKIPSD | bait_int | 0 | 0.992549313 | 3\|6\|4 | 4.33 | 3 | 0\|0\|0\|0\|0\|0 | FALSE |
| PPP1R9B | Q7L590 | MCM10 | bait_int | 0 | 0.994787985 | 14\|25\|25 | 21.33 | 3 | 0\|0\|0\|0\|0\|0 | FALSE |
| PPP1R9B | Q9BUZ4 | TRAF4 | bait_int | 0 | 0.98766998 | 2\|4\|3 | 3 | 3 | 0\|0\|0\|0\|0\|0 | FALSE |
| PPP1R9B | Q9UPN4 | CEP131 | bait_int | 0.05 | 0.991100761 | 1\|6\|6 | 4.33 | 3 | 0\|0\|0\|0\|0\|0 | FALSE |
| PPP1R9B | Q9P2B7 | CFAP97 | bait_int | 0.05 | 0.995432556 | 1\|7\|12 | 6.67 | 3 | 0\|0\|0\|0\|0\|0 | FALSE |
| PPP1R9B | Q8IWC1 | MAP7D3 | bait_int | 0 | 0.992868133 | 17\|29\|25 | 23.67 | 3 | 0\|0\|0\|0\|1\|0 | FALSE |
| PPP1R9B | Q9Y5X1 | SNX9 | bait_int | 0 | 0.989679932 | 7\|9\|11 | 9 | 3 | 0\|0\|0\|0\|0\|0 | FALSE |
| PPP1R9B | P63172 | DYNLT1 | bait_int | 0 | 0.987101648 | 2\|4\|3 | 3 | 3 | 0\|0\|0\|0\|0\|0 | FALSE |
| PPP1R9B | Q8TAF3 | WDR48 | bait_int | 0 | 0.989125463 | 5\|12\|7 | 8 | 3 | 0\|0\|0\|0\|0\|0 | FALSE |
| PPP1R9B | Q6IPR3 | TYW3 | bait_int | 0 | 0.986533317 | 3\|4\|3 | 3.33 | 3 | 0\|0\|0\|0\|0\|0 | FALSE |
| PPP1R9B | Q5SW79 | CEP170 | bait_int | 0.05 | 0.974127057 | 0\|6\|10 | 5.33 | 3 | 0\|0\|0\|0\|0\|0 | FALSE |
| PPP1R9B | Q15154 | PCM1 | bait_int | 0.05 | 0.980406426 | 0\|26\|19 | 15 | 3 | 0\|0\|0\|0\|0\|0 | FALSE |
| PPP5C | Q16543 | CDC37 | bait_int | 0 | 0.981251993 | 21\|17\|23 | 20.33 | 3 | 0\|0\|0\|1\|0\|1 | TRUE |
| PPP5C | O94966 | USP19 | bait_int | 0 | 0.997789052 | 5\|9\|25 | 13 | 3 | 0\|0\|0\|0\|0\|0 | FALSE |
| PPP5C | Q6IA69 | NADSYN1 | bait_int | 0 | 0.997747467 | 14\|16\|6 | 12 | 3 | 0\|0\|0\|0\|0\|0 | FALSE |
| PHF21A | Q9NP66 | HMG20A | bait_int | 0 | 0.995931578 | 15\|13\|13 | 13.67 | 3 | 0\|0\|0\|0\|0\|0 | TRUE |
| PHF21A | P37198 | NUP62 | bait_int | 0 | 0.996853384 | 14\|11\|11 | 12 | 3 | 0\|0\|0\|0\|0\|0 | TRUE |
| PHF21A | Q9P0W2 | HMG20B | bait_int | 0 | 0.998385107 | 17\|19\|17 | 17.67 | 3 | 0\|0\|0\|0\|0\|0 | TRUE |
| PHF21A | O60341 | KDM1A | bait_int | 0 | 0.998419761 | 65\|70\|61 | 65.33 | 3 | 0\|0\|0\|0\|0\|0 | TRUE |
| PHF21A | Q13547 | HDAC1 | bait_int | 0 | 0.98115496 | 13\|16\|14 | 14.33 | 3 | 0\|0\|0\|0\|0\|0 | TRUE |
| PHF21A | Q9UKL0 | RCOR1 | bait_int | 0 | 0.997199928 | 24\|22\|16 | 20.67 | 3 | 0\|0\|0\|0\|0\|0 | TRUE |
| PHF21A | Q92769 | HDAC2 | bait_int | 0 | 0.98640163 | 18\|13\|13 | 14.67 | 3 | 0\|0\|0\|0\|0\|0 | TRUE |
| PHF21A | Q9P2K3 | RCOR3 | bait_int | 0 | 0.997927669 | 22\|21\|19 | 20.67 | 3 | 0\|0\|0\|0\|0\|0 | TRUE |
| PHF21A | Q9H3G5 | CPVL | bait_int | 0 | 0.97256068 | 22\|17\|21 | 20 | 3 | 0\|0\|0\|0\|0\|0 | FALSE |
| PHF21A | Q8IZ40 | RCOR2 | bait_int | 0 | 0.994150344 | 5\|5\|5 | 5 | 3 | 0\|0\|0\|0\|0\|0 | FALSE |
| NACC1 | Q709F0 | ACAD11 | bait_int | 0 | 0.977561304 | 3\|12\|10 | 8.33 | 3 | 0\|0\|0\|0\|0\|0 | FALSE |
| NACC1 | Q9Y4Y9 | LSM5 | bait_int | 0 | 0.972470578 | 2\|3\|4 | 3 | 3 | 0\|0\|0\|0\|0\|0 | FALSE |

| | | | | | | | | | | |
|---|---|---|---|---|---|---|---|---|---|---|
| NACC1 | Q93009 | USP7 | bait_int | 0 | 0.972449786 | 27\|20\|19 | 22 | 3 | 0\|0\|0\|0\|0\|0 | FALSE |
| NACC1 | O95163 | ELP1 | bait_int | 0 | 0.996673182 | 17\|26\|27 | 23.33 | 3 | 0\|0\|0\|0\|0\|0 | FALSE |
| NACC1 | Q9Y4C2 | TCAF1 | bait_int | 0 | 0.995987025 | 20\|24\|22 | 22 | 3 | 0\|0\|0\|0\|0\|0 | FALSE |
| NACC1 | Q8IWV8 | UBR2 | bait_int | 0 | 0.993658253 | 8\|20\|20 | 16 | 3 | 0\|0\|0\|0\|0\|0 | FALSE |
| NACC1 | Q8TBX8 | PIP4K2C | bait_int | 0 | 0.981397541 | 2\|7\|4 | 4.33 | 3 | 0\|0\|0\|0\|0\|0 | FALSE |
| NACC1 | Q9NQH7 | XPNPEP3 | bait_int | 0 | 0.99785143 | 37\|48\|46 | 43.67 | 3 | 0\|0\|0\|0\|0\|0 | FALSE |
| NACC1 | Q9Y333 | LSM2 | bait_int | 0 | 0.980510389 | 3\|5\|4 | 4 | 3 | 0\|0\|0\|0\|0\|0 | FALSE |
| DNMT3A | O00505 | KPNA3 | bait_int | 0 | 0.975769673 | 10\|10\|8 | 9.33 | 3 | 0\|0\|0\|0\|0\|0 | FALSE |
| DNMT3A | P46013 | MKI67 | bait_int | 0 | 0.983764434 | 40\|9\|29 | 26 | 3 | 0\|0\|0\|0\|0\|0 | FALSE |
| BCL11A | O95292 | VAPB | bait_int | 0 | 0.984523364 | 15\|22\|20 | 19 | 3 | 0\|0\|0\|0\|0\|0 | FALSE |
| BCL11A | Q709F0 | ACAD11 | bait_int | 0 | 0.988224449 | 20\|33\|31 | 28 | 3 | 0\|0\|0\|0\|0\|0 | FALSE |
| BCL11A | Q92769 | HDAC2 | bait_int | 0 | 0.990047269 | 23\|28\|23 | 24.67 | 3 | 0\|0\|0\|0\|0\|0 | TRUE |
| BCL11A | O94776 | MTA2 | bait_int | 0 | 0.998918784 | 66\|77\|69 | 70.67 | 3 | 0\|0\|0\|0\|0\|0 | FALSE |
| BCL11A | Q09028 | RBBP4 | bait_int | 0 | 0.981792601 | 28\|38\|44 | 36.67 | 3 | 4\|3\|3\|0\|1\|1 | TRUE |
| BCL11A | Q16576 | RBBP7 | bait_int | 0 | 0.976365728 | 23\|33\|27 | 27.67 | 3 | 3\|1\|0\|0\|0\|0 | TRUE |
| BCL11A | Q13547 | HDAC1 | bait_int | 0 | 0.984710497 | 21\|22\|23 | 22 | 3 | 0\|0\|0\|0\|0\|0 | TRUE |
| BCL11A | Q14839 | CHD4 | bait_int | 0 | 0.998676204 | 117\|114\|1[illegible] | 115 | 3 | 0\|0\|0\|0\|0\|0 | TRUE |
| BCL11A | Q9P0L0 | VAPA | bait_int | 0 | 0.990962144 | 16\|22\|23 | 20.33 | 3 | 0\|0\|0\|0\|0\|0 | FALSE |
| BCL11A | Q9BTC8 | MTA3 | bait_int | 0 | 0.995089478 | 6\|5\|4 | 5 | 3 | 0\|0\|0\|0\|0\|0 | FALSE |
| BCL11A | Q9NYL2 | MAP3K20 | bait_int | 0 | 0.99471521 | 11\|7\|9 | 9 | 3 | 0\|0\|0\|0\|0\|0 | FALSE |
| BCL11A | Q9UBB5 | MBD2 | bait_int | 0 | 0.996032076 | 6\|4\|5 | 5 | 3 | 0\|0\|0\|0\|0\|0 | FALSE |
| BCL11A | Q9Y5J7 | TIMM9 | bait_int | 0 | 0.974834006 | 3\|5\|3 | 3.67 | 3 | 0\|0\|0\|0\|0\|0 | FALSE |
| BCL11A | Q92575 | UBXN4 | bait_int | 0 | 0.995869201 | 5\|5\|4 | 4.67 | 3 | 0\|0\|0\|0\|0\|0 | TRUE |
| BCL11A | O14519 | CDK2AP1 | bait_int | 0 | 0.996617735 | 6\|7\|8 | 7 | 3 | 0\|0\|0\|0\|0\|0 | FALSE |
| BCL11A | O95983 | MBD3 | bait_int | 0 | 0.998835614 | 29\|29\|33 | 30.33 | 3 | 0\|0\|0\|0\|0\|0 | FALSE |
| BCL11A | Q12873 | CHD3 | bait_int | 0 | 0.99856531 | 24\|27\|29 | 26.67 | 3 | 0\|0\|0\|0\|0\|0 | FALSE |
| BCL11A | Q13330 | MTA1 | bait_int | 0 | 0.999168295 | 56\|63\|69 | 62.67 | 3 | 0\|0\|0\|0\|0\|0 | TRUE |
| BCL11A | Q86YP4 | GATAD2A | bait_int | 0 | 0.99904354 | 40\|40\|38 | 39.33 | 3 | 0\|0\|0\|0\|0\|0 | FALSE |
| BCL11A | Q8WXI9 | GATAD2B | bait_int | 0 | 0.999112848 | 41\|56\|53 | 50 | 3 | 0\|0\|0\|0\|0\|0 | FALSE |
| TBR1 | P51659 | HSD17B4 | bait_int | 0 | 0.982527273 | 3\|3\|4 | 3.33 | 3 | 0\|0\|0\|0\|0\|1 | FALSE |
| TBR1 | Q14690 | PDCD11 | bait_int | 0 | 0.984564949 | 64\|45\|53 | 54 | 3 | 0\|0\|0\|0\|0\|0 | FALSE |
| TBR1 | Q709F0 | ACAD11 | bait_int | 0 | 0.977561304 | 2\|10\|13 | 8.33 | 3 | 0\|0\|0\|0\|0\|0 | FALSE |
| TBR1 | P78406 | RAE1 | bait_int | 0 | 0.971971556 | 5\|4\|3 | 4 | 3 | 0\|0\|0\|0\|0\|0 | FALSE |
| TBR1 | Q12797 | ASPH | bait_int | 0 | 0.972193344 | 9\|8\|8 | 8.33 | 3 | 0\|0\|0\|0\|0\|0 | FALSE |
| TBR1 | Q6DN90 | IQSEC1 | bait_int | 0 | 0.993533497 | 3\|16\|15 | 11.33 | 3 | 0\|0\|0\|0\|0\|0 | FALSE |
| TBR1 | Q5T9A4 | ATAD3B | bait_int | 0 | 0.979332141 | 35\|42\|43 | 40 | 3 | 0\|0\|0\|0\|0\|0 | FALSE |
| TBR1 | P28340 | POLD1 | bait_int | 0 | 0.988966053 | 14\|3\|9 | 8.67 | 3 | 0\|0\|0\|0\|0\|0 | FALSE |
| TBR1 | O43823 | AKAP8 | bait_int | 0 | 0.973544864 | 5\|2\|3 | 3.33 | 3 | 1\|0\|0\|0\|0\|0 | FALSE |
| TBR1 | Q93008 | USP9X | bait_int | 0 | 0.980018298 | 12\|25\|22 | 19.67 | 3 | 0\|0\|0\|0\|0\|0 | FALSE |
| TBR1 | Q9UBW7 | ZMYM2 | bait_int | 0.03 | 0.992396834 | 1\|9\|8 | 6 | 3 | 0\|0\|0\|0\|0\|0 | FALSE |
| TBR1 | P53680 | AP2S1 | bait_bait | 0 | 0.988321482 | 3\|2\|2 | 2.33 | 3 | 0\|0\|0\|0\|0\|0 | FALSE |
| TBR1 | Q6PI48 | DARS2 | bait_int | 0 | 0.98416989 | 4\|2\|5 | 3.67 | 3 | 0\|0\|0\|0\|0\|0 | FALSE |
| TBR1 | Q8IVS2 | MCAT | bait_int | 0 | 0.987129372 | 8\|5\|4 | 5.67 | 3 | 0\|0\|0\|0\|0\|0 | FALSE |
| TBR1 | O95470 | SGPL1 | bait_int | 0 | 0.975430061 | 9\|4\|3 | 5.33 | 3 | 0\|0\|0\|0\|0\|0 | FALSE |
| TBR1 | O95782 | AP2A1 | bait_int | 0 | 0.988522477 | 16\|12\|12 | 13.33 | 3 | 0\|0\|0\|0\|0\|0 | FALSE |
| TBR1 | P19447 | ERCC3 | bait_int | 0.05 | 0.992438419 | 9\|1\|2 | 4 | 3 | 0\|0\|0\|0\|0\|0 | FALSE |
| TBR1 | P63010 | AP2B1 | bait_int | 0 | 0.981584674 | 7\|4\|7 | 6 | 3 | 0\|0\|0\|0\|0\|0 | FALSE |
| TBR1 | Q96T37 | RBM15 | bait_int | 0 | 0.981224269 | 28\|13\|11 | 17.33 | 3 | 0\|0\|0\|0\|0\|0 | FALSE |
| TBR1 | Q96CW1 | AP2M1 | bait_int | 0 | 0.981931218 | 10\|8\|9 | 9 | 3 | 0\|0\|0\|0\|0\|0 | FALSE |
| TBR1 | Q9BW92 | TARS2 | bait_int | 0 | 0.982811439 | 8\|2\|5 | 5 | 3 | 0\|0\|0\|0\|0\|0 | FALSE |
| TBR1 | Q9NNW5 | WDR6 | bait_int | 0 | 0.972775537 | 22\|37\|33 | 30.67 | 3 | 0\|0\|0\|0\|0\|0 | FALSE |
| TBR1 | Q9H078 | CLPB | bait_int | 0 | 0.984363954 | 11\|14\|10 | 11.67 | 3 | 0\|0\|0\|0\|0\|0 | FALSE |
| TBR1 | Q5VZL5 | ZMYM4 | bait_int | 0 | 0.992216631 | 3\|10\|11 | 8 | 3 | 0\|0\|0\|0\|0\|0 | FALSE |
| TBR1 | Q96BR5 | COA7 | bait_int | 0 | 0.986422423 | 16\|12\|14 | 14 | 3 | 0\|0\|0\|0\|0\|0 | FALSE |
| TBR1 | O95248 | SBF1 | bait_int | 0 | 0.994327082 | 3\|4\|4 | 3.67 | 3 | 0\|0\|0\|0\|0\|0 | FALSE |
| TBR1 | O75150 | RNF40 | bait_int | 0 | 0.994288962 | 5\|2\|3 | 3.33 | 3 | 0\|0\|0\|0\|0\|0 | FALSE |
| TBR1 | Q6ZW49 | PAXIP1 | bait_int | 0.03 | 0.993117645 | 1\|6\|8 | 5 | 3 | 0\|0\|0\|0\|0\|0 | FALSE |
| TBR1 | Q13643 | FHL3 | bait_int | 0.05 | 0.979678685 | 6\|2\|1 | 3 | 3 | 0\|0\|0\|0\|0\|0 | FALSE |
| TBR1 | Q96L58 | B3GALT6 | bait_int | 0 | 0.993921626 | 5\|4\|3 | 4 | 3 | 0\|0\|0\|0\|0\|0 | FALSE |
| TBR1 | Q9BZH6 | WDR11 | bait_int | 0 | 0.9963405 | 6\|5\|8 | 6.33 | 3 | 0\|0\|0\|0\|0\|0 | FALSE |
| TBR1 | Q01968 | OCRL | bait_int | 0 | 0.989243287 | 4\|4\|3 | 3.67 | 3 | 0\|0\|0\|0\|0\|0 | FALSE |
| TBR1 | Q07864 | POLE | bait_int | 0 | 0.99626426 | 8\|4\|4 | 5.33 | 3 | 0\|0\|0\|0\|0\|0 | FALSE |
| TBR1 | Q5BJF6 | ODF2 | bait_int | 0 | 0.998773236 | 13\|37\|26 | 25.33 | 3 | 0\|0\|0\|0\|0\|0 | FALSE |
| TBR1 | Q6DKJ4 | NXN | bait_int | 0 | 0.99697814 | 10\|8\|6 | 8 | 3 | 0\|0\|0\|0\|0\|0 | FALSE |
| TBR1 | Q16625 | OCLN | bait_int | 0 | 0.996479117 | 4\|9\|7 | 6.67 | 3 | 0\|0\|0\|0\|0\|0 | FALSE |
| TBR1 | O95163 | ELP1 | bait_int | 0.05 | 0.98711551 | 6\|2\|1 | 3 | 3 | 0\|0\|0\|0\|0\|0 | FALSE |
| TBR1 | Q9Y4C2 | TCAF1 | bait_int | 0 | 0.991163139 | 11\|6\|3 | 6.67 | 3 | 0\|0\|0\|0\|0\|0 | FALSE |
| TBR1 | Q969N2 | PIGT | bait_int | 0 | 0.9749449 | 2\|4\|3 | 3 | 3 | 0\|0\|0\|0\|0\|0 | FALSE |
| TBR1 | Q13613 | MTMR1 | bait_int | 0.01 | 0.981674776 | 3\|2\|2 | 2.33 | 3 | 0\|0\|0\|1\|0\|0 | FALSE |
| TBR1 | Q9Y2G8 | DNAJC16 | bait_int | 0 | 0.989492799 | 10\|5\|4 | 6.33 | 3 | 0\|0\|0\|0\|0\|0 | FALSE |
| TBR1 | Q53S58 | TMEM177 | bait_int | 0.03 | 0.98210449 | 6\|1\|4 | 3.67 | 3 | 0\|0\|0\|0\|0\|0 | FALSE |
| TBR1 | Q10567 | AP1B1 | bait_int | 0 | 0.983102535 | 4\|3\|3 | 3.33 | 3 | 0\|0\|0\|0\|0\|0 | FALSE |

| | | | | | | | | | | |
|---|---|---|---|---|---|---|---|---|---|---|
| TBR1 | O94973 | AP2A2 | bait_int | 0 | 0.989347251 | 17\|12\|11 | 13.33 | 3 | 0\|0\|0\|0\|0\|0 | FALSE |
| TBR1 | O60942 | RNGTT | bait_int | 0.03 | 0.975180549 | 5\|4\|1 | 3.33 | 3 | 0\|0\|0\|0\|0\|0 | FALSE |
| TBR1 | Q9UQB8 | BAIAP2 | bait_int | 0 | 0.996416739 | 18\|6\|7 | 10.33 | 3 | 0\|0\|0\|0\|0\|0 | FALSE |
| TBR1 | O14874 | BCKDK | bait_int | 0 | 0.990601738 | 7\|11\|9 | 9 | 3 | 0\|0\|0\|0\|0\|0 | FALSE |
| TBR1 | Q9UKB1 | FBXW11 | bait_int | 0 | 0.997913808 | 4\|22\|19 | 15 | 3 | 0\|0\|0\|0\|0\|0 | FALSE |
| TBR1 | Q9Y297 | BTRC | bait_int | 0 | 0.996732094 | 2\|10\|10 | 7.33 | 3 | 0\|0\|0\|0\|0\|0 | FALSE |
| TBR1 | Q13188 | STK3 | bait_int | 0 | 0.993547359 | 2\|10\|11 | 7.67 | 3 | 0\|0\|0\|0\|0\|0 | FALSE |
| TBR1 | Q9H4B6 | SAV1 | bait_int | 0 | 0.993263193 | 2\|16\|8 | 8.67 | 3 | 0\|0\|0\|0\|0\|0 | FALSE |
| TBR1 | Q658Y4 | FAM91A1 | bait_int | 0 | 0.991908208 | 3\|3\|3 | 3 | 3 | 0\|0\|0\|0\|0\|0 | FALSE |
| TBR1 | Q8NAT1 | POMGNT2 | bait_int | 0 | 0.991204724 | 3\|2\|4 | 3 | 3 | 0\|0\|0\|0\|0\|0 | FALSE |
| TBR1 | Q2M1P5 | KIF7 | bait_int | 0 | 0.994864224 | 14\|28\|26 | 22.67 | 3 | 0\|0\|0\|1\|0\|0 | FALSE |
| TBR1 | Q99569 | PKP4 | bait_int | 0 | 0.982797577 | 2\|12\|5 | 6.33 | 3 | 0\|0\|0\|0\|0\|0 | FALSE |
| TBR1 | O14936 | CASK | bait_int | 0 | 0.992528521 | 6\|4\|9 | 6.33 | 3 | 0\|0\|0\|0\|0\|0 | FALSE |
| TBR1 | Q9HDC5 | JPH1 | bait_int | 0 | 0.988688818 | 5\|7\|8 | 6.67 | 3 | 0\|0\|0\|0\|0\|0 | FALSE |
| TBR1 | Q8NHY2 | COP1 | bait_int | 0.03 | 0.979186593 | 0\|9\|6 | 5 | 3 | 0\|0\|0\|0\|0\|0 | FALSE |
| TM9SF4 | Q00765 | REEP5 | bait_int | 0 | 0.981667845 | 7\|5\|7 | 6.33 | 3 | 0\|0\|0\|0\|0\|0 | FALSE |
| TM9SF4 | Q9NQC3 | RTN4 | bait_int | 0 | 0.981446057 | 3\|3\|5 | 3.67 | 3 | 0\|0\|0\|0\|0\|0 | FALSE |
| TM9SF4 | O95197 | RTN3 | bait_int | 0 | 0.971493326 | 5\|4\|6 | 5 | 3 | 0\|0\|0\|0\|0\|0 | FALSE |
| TM9SF4 | Q7L7X3 | TAOK1 | bait_bait | 0.04 | 0.978957874 | 0\|3\|9 | 4 | 3 | 0\|0\|0\|0\|0\|0 | FALSE |
| SMARCC2 | Q9NQH7 | XPNPEP3 | bait_int | 0 | 0.993692907 | 13\|11\|6 | 10 | 3 | 0\|0\|0\|0\|0\|0 | FALSE |
| SMARCC2 | Q92925 | SMARCD2 | bait_int | 0 | 0.992570106 | 4\|7\|3 | 4.67 | 3 | 0\|0\|0\|0\|0\|0 | TRUE |
| SMARCC2 | Q12824 | SMARCB1 | bait_int | 0 | 0.994164206 | 7\|7\|9 | 7.67 | 3 | 0\|0\|0\|0\|0\|0 | TRUE |
| SMARCC2 | Q96GM5 | SMARCD1 | bait_int | 0 | 0.994621644 | 7\|5\|9 | 7 | 3 | 0\|0\|0\|0\|0\|0 | TRUE |
| SMARCC2 | Q9UJA5 | TRMT6 | bait_int | 0.05 | 0.976033046 | 3\|3\|1 | 2.33 | 3 | 0\|0\|0\|0\|0\|0 | FALSE |
| SMARCC2 | Q92922 | SMARCC1 | bait_int | 0 | 0.992098807 | 3\|2\|3 | 2.67 | 3 | 0\|0\|0\|0\|0\|0 | TRUE |
| SMARCC2 | Q969G3 | SMARCE1 | bait_int | 0 | 0.994178068 | 8\|7\|6 | 7 | 3 | 0\|0\|0\|0\|0\|0 | TRUE |
| SMARCC2 | P51532 | SMARCA4 | bait_int | 0.03 | 0.992944373 | 4\|1\|5 | 3.33 | 3 | 0\|0\|0\|0\|0\|0 | TRUE |
| SMARCC2 | Q9Y5B9 | SUPT16H | bait_int | 0 | 0.975007277 | 35\|37\|32 | 34.67 | 3 | 0\|0\|0\|0\|0\|0 | TRUE |
| SMARCC2 | O15381 | NVL | bait_int | 0 | 0.97733605 | 6\|3\|4 | 4.33 | 3 | 0\|0\|0\|0\|0\|0 | FALSE |
| SMARCC2 | Q01518 | CAP1 | bait_int | 0 | 0.977245949 | 11\|13\|9 | 11 | 3 | 0\|0\|0\|0\|0\|0 | FALSE |
| SMARCC2 | Q14690 | PDCD11 | bait_int | 0 | 0.979546998 | 26\|41\|26 | 31 | 3 | 0\|0\|0\|0\|0\|0 | FALSE |
| SMARCC2 | P46013 | MKI67 | bait_int | 0 | 0.982998572 | 25\|24\|23 | 24 | 3 | 0\|0\|0\|0\|0\|0 | FALSE |
| ZMYND8 | P46013 | MKI67 | bait_int | 0 | 0.986567971 | 29\|26\|58 | 37.67 | 3 | 0\|0\|0\|0\|0\|0 | FALSE |
| ZMYND8 | P13489 | RNH1 | bait_int | 0 | 0.979858887 | 8\|6\|4 | 6 | 3 | 0\|0\|0\|0\|0\|0 | FALSE |
| ZMYND8 | Q14690 | PDCD11 | bait_int | 0 | 0.977322188 | 25\|19\|33 | 25.67 | 3 | 0\|0\|0\|0\|0\|0 | FALSE |
| ZMYND8 | Q9Y5B9 | SUPT16H | bait_int | 0 | 0.975852844 | 43\|45\|24 | 37.33 | 3 | 0\|0\|0\|0\|0\|0 | FALSE |
| ZMYND8 | P17655 | CAPN2 | bait_int | 0 | 0.977620216 | 10\|7\|2 | 6.33 | 3 | 0\|0\|0\|0\|0\|0 | FALSE |
| ZMYND8 | Q6AZZ1 | TRIM68 | bait_int | 0 | 0.994614713 | 11\|9\|5 | 8.33 | 3 | 0\|0\|0\|0\|0\|0 | FALSE |
| ZMYND8 | Q8IWV8 | UBR2 | bait_int | 0 | 0.986872929 | 10\|3\|2 | 5 | 3 | 0\|0\|0\|0\|0\|0 | FALSE |
| ZMYND8 | Q9NQH7 | XPNPEP3 | bait_int | 0 | 0.991447305 | 9\|7\|3 | 6.33 | 3 | 0\|0\|0\|0\|0\|0 | FALSE |
| HDLBP | P31939 | ATIC | bait_int | 0 | 0.980226224 | 9\|10\|10 | 9.67 | 3 | 0\|0\|0\|0\|0\|0 | FALSE |
| HDLBP | P26639 | TARS1 | bait_int | 0 | 0.977031092 | 5\|8\|4 | 5.67 | 3 | 0\|0\|0\|0\|1\|0 | FALSE |
| HDLBP | P50213 | IDH3A | bait_int | 0 | 0.983563438 | 2\|2\|3 | 2.33 | 3 | 0\|0\|0\|0\|0\|0 | FALSE |
| HDLBP | P06737 | PYGL | bait_int | 0 | 0.976951387 | 9\|13\|4 | 8.67 | 3 | 0\|0\|0\|0\|0\|0 | FALSE |
| HDLBP | Q14690 | PDCD11 | bait_int | 0 | 0.973628034 | 18\|22\|16 | 18.67 | 3 | 0\|0\|0\|0\|0\|0 | FALSE |
| HDLBP | Q9UBE0 | SAE1 | bait_int | 0 | 0.982090628 | 4\|7\|5 | 5.33 | 3 | 0\|0\|0\|0\|0\|0 | FALSE |
| HDLBP | Q9NTZ6 | RBM12 | bait_int | 0 | 0.990192817 | 2\|2\|2 | 2 | 3 | 0\|0\|0\|0\|0\|0 | FALSE |
| HDLBP | P49588 | AARS1 | bait_int | 0 | 0.988702679 | 9\|12\|8 | 9.67 | 3 | 0\|0\|0\|0\|0\|0 | FALSE |
| HDLBP | Q14166 | TTLL12 | bait_int | 0 | 0.982721337 | 5\|7\|4 | 5.33 | 3 | 0\|0\|0\|0\|1\|0 | FALSE |
| HDLBP | Q15631 | TSN | bait_int | 0 | 0.977925174 | 6\|5\|3 | 4.67 | 3 | 0\|0\|0\|0\|0\|0 | FALSE |
| HDLBP | P30566 | ADSL | bait_int | 0 | 0.978652916 | 5\|8\|5 | 6 | 3 | 0\|0\|0\|0\|0\|0 | FALSE |
| HDLBP | Q86VP6 | CAND1 | bait_int | 0 | 0.976078097 | 10\|11\|2 | 7.67 | 3 | 0\|0\|0\|0\|0\|0 | FALSE |
| HDLBP | P07954 | FH | bait_int | 0 | 0.976899405 | 4\|4\|2 | 3.33 | 3 | 0\|0\|0\|0\|0\|0 | FALSE |
| HDLBP | P28838 | LAP3 | bait_int | 0 | 0.983878793 | 4\|7\|2 | 4.33 | 3 | 0\|0\|0\|0\|0\|0 | FALSE |
| HDLBP | P11310 | ACADM | bait_int | 0.03 | 0.98290154 | 5\|7\|1 | 4.33 | 3 | 0\|0\|0\|0\|0\|0 | FALSE |
| HDLBP | P41227 | NAA10 | bait_int | 0.03 | 0.978403404 | 4\|4\|3 | 3.67 | 3 | 1\|2\|0\|1\|1\|1 | FALSE |
| HDLBP | Q9BXJ9 | NAA15 | bait_int | 0 | 0.994885017 | 6\|8\|5 | 6.33 | 3 | 0\|0\|0\|0\|0\|0 | FALSE |
| HDLBP | Q9NX55 | HYPK | bait_int | 0 | 0.985098626 | 3\|2\|2 | 2.33 | 3 | 0\|0\|0\|0\|0\|0 | FALSE |
| HDLBP | Q16650 | TBR1 | bait_bait | 0.04 | 0.975173618 | 5\|3\|0 | 2.67 | 3 | 0\|0\|0\|0\|0\|0 | FALSE |
| HDLBP | Q7L7X3 | TAOK1 | bait_bait | 0.04 | 0.986671934 | 0\|3\|26 | 9.67 | 3 | 0\|0\|0\|0\|0\|0 | FALSE |
| KCNQ3 | Q7Z4Q2 | HEATR3 | bait_int | 0.04 | 0.974355775 | 4\|8\|1 | 4.33 | 3 | 0\|0\|0\|0\|0\|0\|0\|( | FALSE |
| KCNQ3 | Q96P70 | IPO9 | bait_int | 0 | 0.974965692 | 6\|9\|5 | 6.67 | 3 | 0\|0\|0\|0\|0\|0\|0\|( | FALSE |
| KCNQ3 | Q99733 | NAP1L4 | bait_int | 0.01 | 0.976552862 | 2\|4\|2 | 2.67 | 3 | 0\|0\|0\|0\|0\|0\|0\|( | FALSE |
| KCNQ3 | Q8TEX9 | IPO4 | bait_int | 0 | 0.975537489 | 5\|5\|3 | 4.33 | 3 | 0\|0\|0\|0\|0\|0\|0\|( | FALSE |
| KCNQ3 | Q92973 | TNPO1 | bait_int | 0 | 0.974896383 | 8\|12\|4 | 8 | 3 | 0\|0\|0\|0\|0\|0\|0\|( | FALSE |
| KCNQ3 | Q9BXJ9 | NAA15 | bait_int | 0.05 | 0.978784602 | 3\|0\|13 | 5.33 | 3 | 0\|0\|0\|0\|0\|0\|0\|( | FALSE |
| KCNQ3 | Q9H4A6 | GOLPH3 | bait_int | 0.04 | 0.989111601 | 4\|5\|1 | 3.33 | 3 | 0\|0\|0\|0\|0\|0\|0\|( | FALSE |
| KCNQ3 | O96005 | CLPTM1 | bait_int | 0 | 0.977959829 | 3\|3\|3 | 3 | 3 | 0\|0\|0\|0\|0\|0\|0\|( | FALSE |
| KCNQ3 | Q9H3P7 | ACBD3 | bait_int | 0 | 0.984613465 | 7\|8\|7 | 7.33 | 3 | 0\|0\|0\|0\|0\|0\|0\|( | FALSE |
| KCNQ3 | O95905 | ECD | bait_int | 0 | 0.99228594 | 10\|11\|10 | 10.33 | 3 | 0\|0\|0\|0\|0\|0\|0\|( | FALSE |
| KCNQ3 | Q96ER3 | SAAL1 | bait_int | 0 | 0.972567611 | 3\|6\|2 | 3.67 | 3 | 0\|0\|0\|0\|0\|0\|0\|( | FALSE |
| KCNQ3 | Q8N8L6 | ARL10 | bait_int | 0 | 0.981643587 | 2\|4\|3 | 3 | 3 | 0\|0\|0\|0\|0\|0\|0\|( | FALSE |

| | | | | | | | | | | |
|---|---|---|---|---|---|---|---|---|---|---|
| KCNQ3 | P40855 | PEX19 | bait_int | 0 | 0.987413537 | 5\|12\|6 | 7.67 | 3 | 0\|0\|0\|0\|0\|0\|0\|( | FALSE |
| WAC | Q9NTZ6 | RBM12 | bait_int | 0 | 0.998800959 | 42\|42\|37 | 40.33 | 3 | 0\|0\|0\|0\|0\|0\|0\|( | FALSE |
| WAC | Q96T37 | RBM15 | bait_int | 0 | 0.97948462 | 14\|8\|22 | 14.67 | 3 | 0\|0\|0\|0\|0\|0\|0\|( | FALSE |
| WAC | Q5T200 | ZC3H13 | bait_int | 0 | 0.998135596 | 38\|23\|42 | 34.33 | 3 | 1\|0\|0\|0\|0\|0\|1\|( | FALSE |
| WAC | O75150 | RNF40 | bait_int | 0 | 0.999189088 | 72\|63\|78 | 71 | 3 | 0\|0\|0\|0\|0\|0\|0\|( | TRUE |
| WAC | P49459 | UBE2A | bait_int | 0.01 | 0.971954229 | 2\|2\|2 | 2 | 3 | 0\|0\|0\|0\|0\|0\|0\|( | FALSE |
| WAC | P24928 | POLR2A | bait_int | 0 | 0.987226404 | 9\|7\|3 | 6.33 | 3 | 0\|0\|0\|0\|0\|0\|0\|( | FALSE |
| WAC | A6NGB9 | WIPF3 | bait_int | 0.01 | 0.994559266 | 2\|5\|2 | 3 | 3 | 0\|0\|0\|0\|0\|0\|0\|( | FALSE |
| WAC | O00401 | WASL | bait_int | 0 | 0.998613826 | 19\|28\|15 | 20.67 | 3 | 0\|0\|0\|0\|0\|0\|0\|( | FALSE |
| WAC | O43516 | WIPF1 | bait_int | 0 | 0.99626426 | 7\|7\|2 | 5.33 | 3 | 0\|0\|0\|0\|0\|0\|0\|( | FALSE |
| WAC | Q5VTR2 | RNF20 | bait_int | 0 | 0.999244535 | 66\|62\|81 | 69.67 | 3 | 0\|0\|0\|0\|0\|0\|0\|( | TRUE |
| WAC | Q69YN4 | VIRMA | bait_int | 0 | 0.99912671 | 55\|29\|48 | 44 | 3 | 0\|0\|0\|0\|0\|0\|0\|( | FALSE |
| WAC | Q75N03 | CBLL1 | bait_int | 0 | 0.997702416 | 10\|9\|11 | 10 | 3 | 0\|0\|0\|0\|0\|0\|0\|( | FALSE |
| WAC | Q8TF74 | WIPF2 | bait_int | 0 | 0.998007374 | 13\|14\|9 | 12 | 3 | 0\|0\|0\|0\|0\|0\|0\|( | FALSE |
| WAC | Q9NX55 | HYPK | bait_int | 0 | 0.990830457 | 7\|5\|4 | 5.33 | 3 | 0\|0\|0\|0\|0\|0\|0\|( | FALSE |
| WAC | Q15007 | WTAP | bait_int | 0 | 0.9984059 | 29\|14\|29 | 24 | 3 | 0\|0\|0\|0\|0\|0\|0\|( | FALSE |
| RORB | Q6ZU65 | UBN2 | bait_int | 0 | 0.97677465 | 7\|8\|2 | 5.67 | 3 | 0\|0\|0\|0\|0\|0\|0\|( | FALSE |
| RORB | P54198 | HIRA | bait_int | 0 | 0.981896564 | 10\|14\|4 | 9.33 | 3 | 0\|0\|0\|0\|0\|0\|0\|( | FALSE |
| RORB | Q8TBC3 | SHKBP1 | bait_int | 0 | 0.988044247 | 3\|2\|6 | 3.67 | 3 | 0\|0\|0\|0\|0\|0\|0\|( | FALSE |
| RORB | Q9NRA8 | EIF4ENIF1 | bait_int | 0 | 0.989354181 | 6\|10\|8 | 8 | 3 | 0\|1\|0\|0\|0\|0\|1\|( | FALSE |
| RORB | P49792 | RANBP2 | bait_int | 0 | 0.993886971 | 21\|12\|15 | 16 | 3 | 0\|0\|0\|0\|0\|0\|0\|( | FALSE |
| RORB | Q9H147 | DNTTIP1 | bait_int | 0 | 0.994489957 | 4\|7\|5 | 5.33 | 3 | 0\|0\|0\|0\|0\|0\|0\|( | FALSE |
| RORB | Q15596 | NCOA2 | bait_int | 0 | 0.997560333 | 25\|25\|22 | 24 | 3 | 0\|0\|0\|0\|0\|0\|0\|( | FALSE |
| RORB | P48552 | NRIP1 | bait_int | 0 | 0.998870268 | 26\|32\|27 | 28.33 | 3 | 0\|0\|0\|0\|0\|0\|0\|( | FALSE |
| RORB | P02794 | FTH1 | bait_int | 0 | 0.99570286 | 3\|4\|6 | 4.33 | 3 | 0\|0\|0\|0\|0\|0\|0\|( | FALSE |
| RORB | Q13557 | CAMK2D | bait_int | 0 | 0.987261058 | 3\|2\|3 | 2.67 | 3 | 0\|0\|0\|0\|0\|0\|0\|( | FALSE |
| RORB | Q15648 | MED1 | bait_int | 0 | 0.989291804 | 3\|4\|5 | 4 | 3 | 0\|0\|0\|0\|0\|0\|0\|( | FALSE |
| RORB | Q15788 | NCOA1 | bait_bait | 0 | 0.998301936 | 13\|24\|20 | 19 | 3 | 0\|0\|0\|0\|0\|0\|0\|( | FALSE |
| RORB | Q6PJG2 | MIDEAS | bait_int | 0 | 0.997685089 | 20\|25\|20 | 21.67 | 3 | 0\|0\|0\|0\|0\|0\|0\|( | FALSE |
| RORB | Q92878 | RAD50 | bait_int | 0 | 0.990130439 | 18\|25\|18 | 20.33 | 3 | 0\|0\|0\|0\|0\|0\|0\|( | FALSE |
| RORB | P04637 | TP53 | bait_int | 0 | 0.978181617 | 27\|29\|29 | 28.33 | 3 | 0\|0\|0\|1\|0\|0\|0\|( | FALSE |
| RORB | Q8IV63 | VRK3 | bait_int | 0 | 0.975762742 | 9\|16\|9 | 11.33 | 3 | 0\|0\|0\|0\|0\|0\|1\|( | FALSE |
| RORB | P78527 | PRKDC | bait_int | 0 | 0.973794375 | 44\|26\|53 | 41 | 3 | 3\|4\|2\|2\|3\|0\|3\|: | FALSE |
| RORB | Q9UEG4 | ZNF629 | bait_int | 0 | 0.997629642 | 14\|14\|20 | 16 | 3 | 0\|0\|0\|0\|0\|0\|0\|( | FALSE |
| RORB | P16278 | GLB1 | bait_int | 0 | 0.984849115 | 8\|11\|9 | 9.33 | 3 | 0\|0\|0\|0\|0\|0\|0\|( | FALSE |
| RORB | P10619 | CTSA | bait_int | 0 | 0.988037316 | 8\|9\|9 | 8.67 | 3 | 0\|0\|0\|0\|0\|0\|0\|( | FALSE |
| RORB | O75420 | GIGYF1 | bait_bait | 0 | 0.997040518 | 9\|11\|6 | 8.67 | 3 | 0\|0\|0\|0\|0\|0\|0\|( | FALSE |
| RORB | O60934 | NBN | bait_int | 0 | 0.981480711 | 2\|4\|4 | 3.33 | 3 | 0\|0\|0\|0\|0\|0\|0\|( | FALSE |
| RORB | P49959 | MRE11 | bait_int | 0 | 0.975970669 | 5\|3\|4 | 4 | 3 | 0\|0\|0\|0\|0\|0\|0\|( | FALSE |
| RORB | Q8NC56 | LEMD2 | bait_int | 0 | 0.978354888 | 12\|15\|9 | 12 | 3 | 0\|0\|0\|0\|0\|0\|0\|( | FALSE |
| RORB | Q9Y6Q9 | NCOA3 | bait_int | 0 | 0.986062017 | 13\|17\|19 | 16.33 | 3 | 2\|0\|0\|0\|0\|0\|0\|( | FALSE |
| RORB | Q13555 | CAMK2G | bait_int | 0 | 0.987025409 | 2\|5\|5 | 4 | 3 | 0\|0\|0\|0\|0\|0\|0\|( | FALSE |
| RORB | P18887 | XRCC1 | bait_int | 0 | 0.973773582 | 13\|11\|3 | 9 | 3 | 0\|0\|0\|0\|0\|0\|0\|( | FALSE |
| RORB | P04181 | OAT | bait_int | 0 | 0.997893015 | 46\|36\|37 | 39.67 | 3 | 0\|0\|0\|0\|0\|0\|0\|( | FALSE |
| RORB | P49916 | LIG3 | bait_int | 0 | 0.982548066 | 25\|28\|11 | 21.33 | 3 | 0\|0\|0\|0\|0\|0\|0\|( | FALSE |
| RORB | P43243 | MATR3 | bait_int | 0 | 0.983622351 | 87\|90\|97 | 91.33 | 3 | 0\|0\|0\|0\|2\|0\|0\|( | FALSE |
| MKX | P49916 | LIG3 | bait_int | 0 | 0.980905449 | 16\|16\|21 | 17.67 | 3 | 0\|0\|0\|0\|0\|0\|0\|( | FALSE |
| MKX | P02545 | LMNA | bait_int | 0 | 0.986450146 | 28\|50\|30 | 36 | 3 | 0\|0\|0\|0\|0\|0\|0\|( | FALSE |
| MKX | P18887 | XRCC1 | bait_int | 0 | 0.973336937 | 9\|7\|10 | 8.67 | 3 | 0\|0\|0\|0\|0\|0\|0\|( | FALSE |
| MKX | P14735 | IDE | bait_int | 0 | 0.992327525 | 8\|16\|8 | 10.67 | 3 | 0\|0\|0\|0\|0\|0\|0\|( | FALSE |
| MKX | Q92769 | HDAC2 | bait_int | 0 | 0.977862796 | 7\|5\|5 | 5.67 | 3 | 0\|0\|0\|0\|0\|0\|0\|( | FALSE |
| MKX | Q9UL15 | BAG5 | bait_int | 0.02 | 0.971715113 | 6\|3\|2 | 3.67 | 3 | 1\|1\|0\|0\|0\|0\|1\|( | FALSE |
| MKX | Q8NC56 | LEMD2 | bait_int | 0 | 0.974477066 | 8\|8\|10 | 8.67 | 3 | 0\|0\|0\|0\|0\|0\|0\|( | FALSE |
| MKX | Q01433 | AMPD2 | bait_int | 0 | 0.996160297 | 23\|20\|21 | 21.33 | 3 | 0\|0\|0\|0\|0\|0\|0\|( | FALSE |
| MKX | Q99733 | NAP1L4 | bait_int | 0 | 0.994011727 | 26\|30\|27 | 27.67 | 3 | 0\|0\|0\|0\|0\|0\|0\|( | FALSE |
| MKX | P82933 | MRPS9 | bait_int | 0 | 0.975267185 | 3\|4\|5 | 4 | 3 | 0\|0\|0\|0\|0\|0\|0\|( | FALSE |
| MKX | Q9BZK7 | TBL1XR1 | bait_bait | 0 | 0.992583968 | 11\|8\|8 | 9 | 3 | 0\|0\|0\|0\|0\|0\|0\|( | TRUE |
| MKX | Q9Y618 | NCOR2 | bait_int | 0 | 0.990275987 | 8\|7\|14 | 9.67 | 3 | 0\|0\|0\|0\|0\|0\|0\|( | FALSE |
| MKX | P19447 | ERCC3 | bait_int | 0 | 0.990185886 | 3\|2\|3 | 2.67 | 3 | 0\|0\|0\|0\|0\|0\|0\|( | FALSE |
| MKX | O15379 | HDAC3 | bait_int | 0.01 | 0.978743017 | 2\|2\|4 | 2.67 | 3 | 0\|0\|0\|0\|0\|0\|0\|( | FALSE |
| MKX | O60907 | TBL1X | bait_int | 0 | 0.986311529 | 4\|4\|5 | 4.33 | 3 | 0\|0\|0\|0\|0\|0\|0\|( | TRUE |
| MKX | O75376 | NCOR1 | bait_int | 0 | 0.984866442 | 2\|3\|6 | 3.67 | 3 | 0\|0\|0\|0\|0\|0\|0\|( | FALSE |
| MKX | Q8WWY3 | PRPF31 | bait_int | 0 | 0.985077834 | 4\|5\|5 | 4.67 | 3 | 0\|0\|0\|0\|0\|0\|0\|( | FALSE |
| MKX | Q9UNF1 | MAGED2 | bait_int | 0 | 0.988605647 | 12\|13\|12 | 12.33 | 3 | 0\|0\|0\|0\|0\|0\|0\|( | FALSE |
| MKX | P78527 | PRKDC | bait_int | 0 | 0.983927309 | 101\|75\|143 | 106.33 | 3 | 3\|4\|2\|2\|3\|0\|3\|: | FALSE |
| MKX | P13010 | XRCC5 | bait_int | 0 | 0.97367655 | 63\|58\|61 | 60.67 | 3 | 1\|0\|0\|0\|0\|0\|0\|( | FALSE |
| MKX | Q13547 | HDAC1 | bait_int | 0 | 0.973482486 | 6\|6\|10 | 7.33 | 3 | 0\|0\|0\|0\|0\|0\|1\|( | FALSE |
| MKX | Q14839 | CHD4 | bait_int | 0 | 0.986200635 | 3\|4\|4 | 3.67 | 3 | 1\|0\|0\|0\|0\|0\|0\|( | FALSE |
| MKX | Q9H1B7 | IRF2BPL | bait_bait | 0.03 | 0.991377996 | 3\|5\|6 | 4.67 | 3 | 2\|3\|0\|0\|0\|0\|1\|: | FALSE |
| MKX | O75182 | SIN3B | bait_int | 0 | 0.994365201 | 3\|7\|9 | 6.33 | 3 | 0\|0\|0\|0\|0\|0\|0\|( | FALSE |
| MKX | O75446 | SAP30 | bait_int | 0.01 | 0.987812062 | 2\|3\|2 | 2.33 | 3 | 0\|0\|0\|0\|0\|0\|0\|( | FALSE |
| MKX | Q96ST3 | SIN3A | bait_bait | 0 | 0.998378176 | 37\|30\|48 | 38.33 | 3 | 0\|0\|0\|0\|0\|0\|0\|( | FALSE |

| | | | | | | | | | | |
|---|---|---|---|---|---|---|---|---|---|---|
| MKX | Q6SPF0 | SAMD1 | bait_int | 0.04 | 0.97868757 | 6\|7\|1 | 4.67 | 3 | 0\|0\|0\|0\|0\|0\|0\|( | FALSE |
| MKX | Q8IU81 | IRF2BP1 | bait_int | 0 | 0.981300509 | 3\|2\|7 | 4 | 3 | 0\|0\|0\|0\|0\|0\|0\|( | FALSE |
| MKX | P17066 | HSPA6 | bait_int | 0 | 0.990178955 | 17\|22\|21 | 20 | 3 | 0\|0\|0\|0\|0\|1\|0\|( | FALSE |
| MKX | Q13227 | GPS2 | bait_int | 0 | 0.978645985 | 3\|2\|3 | 2.67 | 3 | 0\|0\|0\|0\|0\|0\|0\|( | FALSE |
| MKX | Q8IWV7 | UBR1 | bait_bait | 0 | 0.996971209 | 8\|9\|12 | 9.67 | 3 | 0\|0\|0\|0\|0\|0\|0\|( | FALSE |
| MKX | P54198 | HIRA | bait_int | 0 | 0.978472713 | 7\|7\|6 | 6.67 | 3 | 0\|0\|0\|0\|0\|0\|0\|( | FALSE |
| MKX | Q5T5X7 | BEND3 | bait_int | 0 | 0.973849822 | 2\|4\|7 | 4.33 | 3 | 0\|0\|0\|0\|0\|0\|0\|( | FALSE |
| MKX | Q6ZU65 | UBN2 | bait_int | 0 | 0.971424017 | 3\|6\|2 | 3.67 | 3 | 0\|0\|0\|0\|0\|0\|0\|( | FALSE |
| PPP2R5D | Q13136 | PPFIA1 | bait_int | 0 | 0.998599964 | 35\|34\|40 | 36.33 | 3 | 0\|0\|0\|0\|0\|0\|0\|( | TRUE |
| PPP2R5D | P15735 | PHKG2 | bait_int | 0 | 0.984786737 | 4\|6\|9 | 6.33 | 3 | 0\|0\|0\|0\|0\|0\|0\|( | FALSE |
| PPP2R5D | P46020 | PHKA1 | bait_int | 0.04 | 0.984942682 | 1\|4\|4 | 3 | 3 | 0\|0\|0\|0\|0\|0\|0\|( | FALSE |
| PPP2R5D | Q9BWN1 | PRR14 | bait_int | 0 | 0.997764794 | 9\|9\|13 | 10.33 | 3 | 0\|0\|0\|0\|0\|0\|0\|( | TRUE |
| PPP2R5D | Q5FBB7 | SGO1 | bait_int | 0 | 0.99626426 | 5\|6\|5 | 5.33 | 3 | 0\|0\|0\|0\|0\|0\|0\|( | FALSE |
| PPP2R5D | P60510 | PPP4C | bait_int | 0 | 0.998475208 | 19\|19\|22 | 20 | 3 | 0\|0\|0\|0\|0\|0\|0\|( | TRUE |
| PPP2R5D | P30154 | PPP2R1B | bait_int | 0 | 0.998849475 | 33\|31\|32 | 32 | 3 | 0\|0\|0\|0\|0\|0\|0\|( | TRUE |
| PPP2R5D | P62714 | PPP2CB | bait_int | 0 | 0.991419581 | 5\|2\|5 | 4 | 3 | 0\|0\|0\|0\|0\|0\|0\|( | TRUE |
| PPP2R5D | P67775 | PPP2CA | bait_int | 0 | 0.992965165 | 4\|3\|7 | 4.67 | 3 | 0\|0\|0\|0\|0\|0\|0\|( | TRUE |
| PPP2R5D | P30153 | PPP2R1A | bait_int | 0 | 0.977634078 | 76\|79\|88 | 81 | 3 | 1\|0\|1\|1\|1\|0\|3\|3 | TRUE |
| PPP2R5D | O75145 | PPFIA3 | bait_int | 0 | 0.996070196 | 5\|5\|10 | 6.67 | 3 | 0\|0\|0\|0\|0\|0\|0\|( | FALSE |
| PPP2R5D | Q02218 | OGDH | bait_int | 0 | 0.984994663 | 11\|9\|20 | 13.33 | 3 | 0\|0\|0\|0\|0\|0\|0\|( | FALSE |
| PPP2R5D | Q13724 | MOGS | bait_int | 0 | 0.980884656 | 7\|5\|10 | 7.33 | 3 | 0\|0\|0\|0\|0\|0\|0\|( | FALSE |
| SPAST | Q8NFH4 | NUP37 | bait_int | 0 | 0.993727561 | 6\|6\|8 | 6.67 | 3 | 0\|0\|0\|0\|0\|0\|0\|( | FALSE |
| SPAST | Q00765 | REEP5 | bait_int | 0 | 0.974646872 | 3\|5\|2 | 3.33 | 3 | 0\|0\|0\|0\|0\|0\|0\|( | FALSE |
| SPAST | Q92990 | GLMN | bait_int | 0.05 | 0.9885502 | 4\|3\|1 | 2.67 | 3 | 0\|0\|0\|0\|0\|0\|0\|( | FALSE |
| SPAST | P53990 | IST1 | bait_int | 0 | 0.985452101 | 3\|7\|7 | 5.67 | 3 | 0\|0\|0\|0\|0\|0\|0\|( | TRUE |
| SPAST | P52948 | NUP98 | bait_int | 0 | 0.99681873 | 25\|27\|33 | 28.33 | 3 | 0\|0\|0\|0\|0\|0\|0\|( | TRUE |
| SPAST | P40855 | PEX19 | bait_int | 0 | 0.992514659 | 20\|18\|15 | 17.67 | 3 | 0\|0\|0\|0\|0\|0\|0\|( | FALSE |
| SPAST | O43264 | ZW10 | bait_int | 0 | 0.985320414 | 3\|3\|3 | 3 | 3 | 0\|0\|0\|0\|0\|0\|0\|( | FALSE |
| SPAST | Q9P0L0 | VAPA | bait_int | 0.05 | 0.975783535 | 6\|3\|1 | 3.33 | 3 | 0\|0\|0\|0\|0\|0\|0\|( | FALSE |
| SPAST | Q92616 | GCN1 | bait_int | 0 | 0.990317573 | 8\|6\|3 | 5.67 | 3 | 0\|0\|0\|0\|0\|0\|0\|( | FALSE |
| SPAST | Q96T76 | MMS19 | bait_int | 0 | 0.98441247 | 5\|7\|6 | 6 | 3 | 0\|0\|0\|0\|0\|0\|0\|( | FALSE |
| SPAST | Q96ER3 | SAAL1 | bait_int | 0 | 0.979131146 | 5\|6\|8 | 6.33 | 3 | 0\|0\|0\|0\|0\|0\|0\|( | FALSE |
| SPAST | Q9BW27 | NUP85 | bait_int | 0 | 0.998911853 | 38\|43\|40 | 40.33 | 3 | 0\|0\|0\|0\|0\|0\|0\|( | FALSE |
| SPAST | Q9UIA9 | XPO7 | bait_int | 0.01 | 0.985861022 | 2\|2\|3 | 2.33 | 3 | 0\|0\|0\|0\|0\|0\|0\|( | FALSE |
| SPAST | Q86Y56 | DNAAF5 | bait_int | 0 | 0.990982936 | 7\|12\|7 | 8.67 | 3 | 0\|0\|0\|0\|0\|0\|0\|( | FALSE |
| SPAST | Q8TC07 | TBC1D15 | bait_int | 0.01 | 0.97952274 | 2\|2\|2 | 2 | 3 | 0\|0\|0\|0\|0\|0\|0\|( | FALSE |
| SPAST | Q96J02 | ITCH | bait_int | 0 | 0.979893542 | 3\|5\|3 | 3.67 | 3 | 0\|0\|0\|0\|0\|0\|0\|( | FALSE |
| SPAST | Q12769 | NUP160 | bait_int | 0 | 0.998925715 | 40\|48\|41 | 43 | 3 | 0\|0\|0\|0\|0\|0\|0\|( | FALSE |
| SPAST | Q9UI10 | EIF2B4 | bait_int | 0 | 0.992181977 | 4\|8\|8 | 6.67 | 3 | 0\|0\|0\|0\|0\|0\|0\|( | FALSE |
| SPAST | Q9HAV4 | XPO5 | bait_int | 0 | 0.984745152 | 12\|12\|11 | 11.67 | 3 | 0\|0\|0\|0\|0\|0\|0\|( | FALSE |
| SPAST | P56192 | MARS1 | bait_int | 0 | 0.971909178 | 16\|17\|19 | 17.33 | 3 | 0\|0\|0\|0\|0\|0\|0\|( | FALSE |
| SPAST | P57740 | NUP107 | bait_int | 0 | 0.987170957 | 23\|19\|20 | 20.67 | 3 | 0\|0\|0\|0\|0\|0\|0\|( | TRUE |
| SPAST | P78527 | PRKDC | bait_int | 0 | 0.977855866 | 49\|59\|65 | 57.67 | 3 | 3\|4\|2\|2\|3\|0\|3\|1 | FALSE |
| SPAST | O43592 | XPOT | bait_int | 0 | 0.977193967 | 21\|22\|24 | 22.33 | 3 | 0\|0\|0\|0\|0\|0\|0\|( | FALSE |
| SPAST | O95373 | IPO7 | bait_int | 0 | 0.974764697 | 11\|15\|13 | 13 | 3 | 0\|0\|0\|0\|0\|0\|0\|( | FALSE |
| SPAST | Q92973 | TNPO1 | bait_int | 0 | 0.981501504 | 14\|15\|15 | 14.67 | 3 | 0\|0\|0\|0\|0\|0\|0\|( | FALSE |
| SPAST | Q8TCT9 | HM13 | bait_int | 0 | 0.993228539 | 5\|9\|8 | 7.33 | 3 | 0\|0\|0\|0\|0\|0\|0\|( | FALSE |
| SPAST | Q8WUM0 | NUP133 | bait_int | 0 | 0.994205791 | 23\|29\|30 | 27.33 | 3 | 0\|0\|0\|0\|0\|0\|0\|( | TRUE |
| SPAST | Q8NFH3 | NUP43 | bait_int | 0 | 0.989596762 | 9\|12\|13 | 11.33 | 3 | 0\|0\|0\|0\|0\|0\|0\|( | TRUE |
| SPAST | Q93008 | USP9X | bait_int | 0 | 0.98393424 | 23\|37\|30 | 30 | 3 | 0\|0\|0\|0\|0\|0\|0\|( | FALSE |
| SPAST | O14980 | XPO1 | bait_int | 0 | 0.971139851 | 24\|25\|19 | 22.67 | 3 | 0\|0\|0\|0\|0\|0\|0\|( | FALSE |
| SPAST | Q7Z4Q2 | HEATR3 | bait_int | 0 | 0.974355775 | 5\|5\|3 | 4.33 | 3 | 0\|0\|0\|0\|0\|0\|0\|( | FALSE |
| SPAST | P55786 | NPEPPS | bait_int | 0 | 0.971486395 | 9\|6\|6 | 7 | 3 | 0\|0\|0\|0\|0\|0\|0\|( | FALSE |
| GNAI1 | Q07812 | BAX | bait_int | 0 | 0.983019365 | 3\|4\|3 | 3.33 | 3 | 0\|0\|0\|0\|0\|0\|0\|( | FALSE |
| GNAI1 | O14980 | XPO1 | bait_int | 0 | 0.971638874 | 22\|19\|29 | 23.33 | 3 | 0\|0\|0\|0\|0\|0\|0\|( | FALSE |
| GNAI1 | Q92973 | TNPO1 | bait_int | 0 | 0.985438239 | 20\|21\|29 | 23.33 | 3 | 0\|0\|0\|0\|0\|0\|0\|( | FALSE |
| GNAI1 | P80303 | NUCB2 | bait_int | 0.01 | 0.988473961 | 2\|2\|2 | 2 | 3 | 0\|0\|0\|0\|0\|0\|0\|( | FALSE |
| GNAI1 | Q9HAV4 | XPO5 | bait_int | 0.05 | 0.977239018 | 3\|1\|12 | 5.33 | 3 | 0\|0\|0\|0\|0\|0\|0\|( | FALSE |
| GNAI1 | P50150 | GNG4 | bait_int | 0 | 0.989437352 | 3\|2\|4 | 3 | 3 | 0\|0\|0\|0\|0\|0\|0\|( | TRUE |
| GNAI1 | O95379 | TNFAIP8 | bait_int | 0 | 0.994462234 | 9\|9\|8 | 8.67 | 3 | 0\|0\|0\|0\|0\|0\|0\|( | FALSE |
| GNAI1 | P47736 | RAP1GAP | bait_int | 0 | 0.994732538 | 4\|2\|8 | 4.67 | 3 | 0\|0\|0\|0\|0\|0\|0\|( | FALSE |
| GNAI1 | Q3V6T2 | CCDC88A | bait_int | 0 | 0.998593033 | 22\|19\|21 | 20.67 | 3 | 0\|0\|0\|0\|0\|0\|0\|( | FALSE |
| GNAI1 | Q5T2E6 | ARMH3 | bait_int | 0 | 0.99491274 | 4\|3\|3 | 3.33 | 3 | 0\|0\|0\|0\|0\|0\|0\|( | FALSE |
| GNAI1 | Q86YJ7 | ANKRD13B | bait_int | 0.04 | 0.994559266 | 4\|1\|4 | 3 | 3 | 0\|0\|0\|0\|0\|0\|0\|( | FALSE |
| GNAI1 | Q9NPQ8 | RIC8A | bait_int | 0 | 0.996777145 | 13\|22\|15 | 16.67 | 3 | 0\|0\|0\|0\|0\|0\|0\|( | TRUE |
| GNAI1 | Q9UIA9 | XPO7 | bait_int | 0 | 0.989271011 | 3\|2\|6 | 3.67 | 3 | 0\|0\|0\|0\|0\|0\|0\|( | FALSE |
| GNAI1 | Q86Y56 | DNAAF5 | bait_int | 0 | 0.990241333 | 8\|8\|7 | 7.67 | 3 | 0\|0\|0\|0\|0\|0\|0\|( | FALSE |
| GNAI1 | Q9Y5L0 | TNPO3 | bait_int | 0 | 0.982402517 | 3\|2\|9 | 4.67 | 3 | 0\|0\|0\|0\|0\|0\|0\|( | FALSE |
| GNAI1 | P53611 | RABGGTB | bait_int | 0 | 0.981882702 | 2\|3\|4 | 3 | 3 | 0\|0\|0\|0\|0\|0\|0\|( | FALSE |
| GNAI1 | Q92616 | GCN1 | bait_int | 0 | 0.995280077 | 7\|6\|35 | 16 | 3 | 0\|0\|0\|0\|0\|0\|0\|( | FALSE |
| GNAI1 | P53365 | ARFIP2 | bait_int | 0.01 | 0.987458588 | 4\|2\|2 | 2.67 | 3 | 0\|0\|0\|0\|0\|0\|0\|( | FALSE |
| GNAI1 | Q9NVI1 | FANCI | bait_int | 0.01 | 0.971015095 | 2\|2\|4 | 2.67 | 3 | 0\|0\|0\|0\|0\|0\|0\|( | FALSE |

| | | | | | | | | | |
|---|---|---|---|---|---|---|---|---|---|
| GNAI1 | Q5VYK3 | ECPAS | bait_int | 0 | 0.987843251 | 5\|4\|7 | 5.33 | 3 | 0\|0\|0\|0\|0\|0\|0\|( | FALSE |
| GNAI1 | Q9UI12 | ATP6V1H | bait_int | 0 | 0.972761675 | 6\|6\|8 | 6.67 | 3 | 0\|0\|0\|0\|0\|0\|0\|( | FALSE |
| GNAI1 | Q02818 | NUCB1 | bait_int | 0 | 0.993131506 | 6\|8\|5 | 6.33 | 3 | 0\|0\|0\|0\|0\|0\|0\|( | FALSE |
| GNAI1 | Q86YR5 | GPSM1 | bait_int | 0 | 0.99539097 | 14\|8\|15 | 12.33 | 3 | 0\|0\|0\|0\|0\|0\|0\|( | TRUE |
| GNAI1 | Q00765 | REEP5 | bait_int | 0 | 0.97168739 | 3\|3\|2 | 2.67 | 3 | 0\|0\|0\|0\|0\|0\|0\|( | FALSE |
| FOXP2 | Q6ZU65 | UBN2 | bait_int | 0 | 0.97264385 | 5\|5\|2 | 4 | 3 | 0\|0\|0\|0\|0\|0\|0\|( | FALSE |
| FOXP2 | Q5T5X7 | BEND3 | bait_int | 0 | 0.978389543 | 4\|11\|4 | 6.33 | 3 | 0\|0\|0\|0\|0\|0\|0\|( | FALSE |
| FOXP2 | P54198 | HIRA | bait_int | 0.05 | 0.975138964 | 3\|11\|1 | 5 | 3 | 0\|0\|0\|0\|0\|0\|0\|( | FALSE |
| FOXP2 | O14874 | BCKDK | bait_int | 0 | 0.990054199 | 9\|5\|11 | 8.33 | 3 | 0\|0\|0\|0\|0\|0\|0\|( | FALSE |
| FOXP2 | Q9H334 | FOXP1 | bait_bait | 0 | 0.997380131 | 7\|13\|18 | 12.67 | 3 | 0\|0\|0\|0\|0\|0\|0\|( | TRUE |
| FOXP2 | Q8IVH2 | FOXP4 | bait_int | 0 | 0.995481072 | 8\|9\|14 | 10.33 | 3 | 0\|0\|0\|0\|0\|0\|0\|( | TRUE |
| FOXP2 | P78527 | PRKDC | bait_int | 0 | 0.975606798 | 50\|51\|41 | 47.33 | 3 | 3\|4\|2\|2\|3\|0\|3\|1 | FALSE |
| FOXP2 | P49916 | LIG3 | bait_int | 0 | 0.983733245 | 24\|30\|19 | 24.33 | 3 | 0\|0\|0\|0\|0\|0\|0\|( | FALSE |
| LDB1 | Q10713 | PMPCA | bait_int | 0 | 0.979124215 | 8\|8\|12 | 9.33 | 3 | 0\|0\|0\|0\|0\|0\|0\|( | FALSE |
| LDB1 | Q7Z4Q2 | HEATR3 | bait_int | 0 | 0.972304238 | 3\|4\|4 | 3.67 | 3 | 0\|0\|0\|0\|0\|0\|0\|( | FALSE |
| LDB1 | Q93008 | USP9X | bait_int | 0 | 0.979443035 | 19\|20\|17 | 18.67 | 3 | 0\|0\|0\|0\|0\|0\|0\|( | FALSE |
| LDB1 | Q7Z6Z7 | HUWE1 | bait_int | 0 | 0.990303711 | 49\|25\|44 | 39.33 | 3 | 0\|0\|0\|0\|0\|0\|0\|( | FALSE |
| LDB1 | O95373 | IPO7 | bait_int | 0 | 0.973011186 | 11\|14\|9 | 11.33 | 3 | 0\|0\|0\|0\|0\|0\|0\|( | FALSE |
| LDB1 | P78527 | PRKDC | bait_int | 0 | 0.976195922 | 56\|48\|46 | 50 | 3 | 3\|4\|2\|2\|3\|0\|3\|1 | FALSE |
| LDB1 | Q70CQ2 | USP34 | bait_int | 0.01 | 0.985937262 | 2\|2\|4 | 2.67 | 3 | 0\|0\|0\|0\|0\|0\|0\|( | FALSE |
| LDB1 | Q9BWW4 | SSBP3 | bait_int | 0 | 0.996829126 | 8\|7\|5 | 6.67 | 3 | 0\|0\|0\|0\|0\|0\|0\|( | TRUE |
| LDB1 | Q96A47 | ISL2 | bait_int | 0 | 0.997879153 | 8\|9\|16 | 11 | 3 | 0\|0\|0\|0\|0\|0\|0\|( | TRUE |
| LDB1 | O60318 | MCM3AP | bait_int | 0 | 0.993034474 | 5\|3\|2 | 3.33 | 3 | 0\|0\|0\|0\|0\|0\|0\|( | FALSE |
| LDB1 | O60663 | LMX1B | bait_int | 0.05 | 0.99491274 | 1\|3\|6 | 3.33 | 3 | 0\|0\|0\|0\|0\|0\|0\|( | TRUE |
| LDB1 | P21359 | NF1 | bait_int | 0.05 | 0.990861646 | 3\|1\|4 | 2.67 | 3 | 0\|0\|0\|0\|0\|0\|0\|( | FALSE |
| LDB1 | P61968 | LMO4 | bait_int | 0 | 0.997879153 | 13\|11\|9 | 11 | 3 | 0\|0\|0\|0\|0\|0\|0\|( | TRUE |
| LDB1 | P81877 | SSBP2 | bait_int | 0.01 | 0.992732981 | 2\|2\|2 | 2 | 3 | 0\|0\|0\|0\|0\|0\|0\|( | TRUE |
| LDB1 | Q5TDH0 | DDI2 | bait_int | 0 | 0.996548426 | 7\|6\|5 | 6 | 3 | 0\|0\|0\|0\|0\|0\|0\|( | FALSE |
| LDB1 | Q9BWG4 | SSBP4 | bait_int | 0.01 | 0.993387949 | 2\|3\|2 | 2.33 | 3 | 0\|0\|0\|0\|0\|0\|0\|( | TRUE |
| LDB1 | Q13451 | FKBP5 | bait_int | 0 | 0.995425625 | 9\|11\|11 | 10.33 | 3 | 0\|0\|0\|0\|0\|0\|0\|( | FALSE |
| LDB1 | Q9UHI6 | DDX20 | bait_int | 0 | 0.98544517 | 5\|5\|4 | 4.67 | 3 | 0\|0\|0\|0\|0\|0\|0\|( | FALSE |
| LDB1 | Q9UJX3 | ANAPC7 | bait_int | 0 | 0.995515726 | 7\|8\|10 | 8.33 | 3 | 0\|0\|0\|0\|0\|0\|0\|( | FALSE |
| LDB1 | Q9UPU5 | USP24 | bait_int | 0 | 0.992431488 | 6\|5\|4 | 5 | 3 | 0\|0\|0\|0\|0\|0\|0\|( | FALSE |
| LDB1 | Q66LE6 | PPP2R2D | bait_int | 0 | 0.988023454 | 4\|2\|4 | 3.33 | 3 | 0\|0\|0\|0\|0\|0\|0\|( | FALSE |
| LDB1 | Q9BXW9 | FANCD2 | bait_int | 0 | 0.973330007 | 3\|5\|6 | 4.67 | 3 | 0\|0\|0\|0\|0\|0\|0\|( | FALSE |
| LDB1 | Q96T76 | MMS19 | bait_int | 0 | 0.978888565 | 3\|4\|3 | 3.33 | 3 | 0\|0\|0\|0\|0\|0\|0\|( | FALSE |
| LDB1 | O75439 | PMPCB | bait_int | 0 | 0.985666958 | 6\|10\|9 | 8.33 | 3 | 0\|0\|1\|0\|0\|0\|0\|( | FALSE |
| LDB1 | Q8TCG1 | CIP2A | bait_int | 0 | 0.995079081 | 32\|18\|29 | 26.33 | 3 | 0\|0\|0\|0\|0\|0\|0\|( | FALSE |
| KIAA0232 | Q709F0 | ACAD11 | bait_int | 0 | 0.985161004 | 18\|16\|22 | 18.67 | 3 | 0\|0\|0\|0\|0\|0\|0\|( | FALSE |
| KIAA0232 | Q5T9A4 | ATAD3B | bait_int | 0 | 0.973919131 | 26\|24\|26 | 25.33 | 3 | 0\|0\|0\|0\|0\|0\|0\|( | FALSE |
| KIAA0232 | Q6P1J9 | CDC73 | bait_int | 0.01 | 0.985420912 | 2\|2\|2 | 2 | 3 | 0\|0\|0\|0\|0\|0\|0\|( | FALSE |
| KIAA0232 | Q9UNF1 | MAGED2 | bait_int | 0 | 0.980856933 | 5\|5\|4 | 4.67 | 3 | 0\|0\|0\|0\|0\|0\|0\|( | FALSE |
| KIAA0232 | P61962 | DCAF7 | bait_int | 0 | 0.9895829 | 5\|8\|8 | 7 | 3 | 0\|1\|0\|0\|0\|0\|1\|1 | FALSE |
| KIAA0232 | Q15036 | SNX17 | bait_int | 0 | 0.973953785 | 2\|4\|6 | 4 | 3 | 0\|0\|0\|0\|0\|0\|0\|( | FALSE |
| KIAA0232 | Q96BR5 | COA7 | bait_int | 0 | 0.978493506 | 5\|7\|5 | 5.67 | 3 | 1\|0\|0\|0\|0\|0\|0\|1 | FALSE |
| KIAA0232 | Q5W0B1 | OBI1 | bait_int | 0 | 0.994108759 | 5\|6\|7 | 6 | 3 | 0\|0\|0\|0\|0\|0\|0\|( | FALSE |
| KIAA0232 | A5YKK6 | CNOT1 | bait_int | 0 | 0.998184112 | 28\|23\|29 | 26.67 | 3 | 0\|0\|0\|0\|0\|0\|0\|( | FALSE |
| KIAA0232 | Q92600 | CNOT9 | bait_int | 0 | 0.994327082 | 5\|3\|3 | 3.67 | 3 | 0\|0\|0\|0\|0\|0\|0\|( | FALSE |
| KIAA0232 | Q9H9A5 | CNOT10 | bait_int | 0 | 0.993866179 | 3\|6\|8 | 5.67 | 3 | 0\|0\|0\|0\|0\|0\|0\|( | FALSE |
| KIAA0232 | Q9UKZ1 | CNOT11 | bait_int | 0.04 | 0.990265591 | 4\|4\|1 | 3 | 3 | 0\|0\|0\|0\|0\|0\|0\|( | FALSE |
| KIAA0232 | Q9NZN8 | CNOT2 | bait_int | 0 | 0.992972096 | 4\|4\|7 | 5 | 3 | 0\|0\|0\|0\|0\|0\|0\|( | FALSE |
| KIAA0232 | Q7L590 | MCM10 | bait_int | 0 | 0.986609556 | 4\|6\|5 | 5 | 3 | 0\|0\|0\|0\|0\|0\|0\|( | FALSE |
| KIAA0232 | Q9C0C7 | AMBRA1 | bait_int | 0 | 0.991426512 | 3\|3\|2 | 2.67 | 3 | 0\|0\|0\|0\|0\|0\|0\|( | FALSE |
| KIAA0232 | Q99471 | PFDN5 | bait_int | 0.01 | 0.973056237 | 3\|2\|2 | 2.33 | 3 | 0\|0\|0\|0\|0\|0\|0\|( | FALSE |
| KIAA0232 | O95163 | ELP1 | bait_int | 0 | 0.988612578 | 3\|4\|4 | 3.67 | 3 | 0\|0\|0\|0\|0\|0\|0\|( | FALSE |
| KIAA0232 | Q13112 | CHAF1B | bait_int | 0 | 0.993041405 | 7\|3\|11 | 7 | 3 | 0\|0\|0\|0\|0\|0\|0\|( | FALSE |
| KIAA0232 | Q6PML9 | SLC30A9 | bait_int | 0 | 0.993859248 | 5\|3\|8 | 5.33 | 3 | 0\|0\|0\|0\|0\|0\|0\|( | FALSE |
| KIAA0232 | Q9Y2X9 | ZNF281 | bait_int | 0 | 0.989707656 | 4\|5\|3 | 4 | 3 | 0\|0\|0\|0\|0\|0\|0\|( | FALSE |
| KIAA0232 | Q15072 | ZNF146 | bait_int | 0 | 0.986949169 | 7\|7\|9 | 7.67 | 3 | 0\|0\|0\|0\|0\|0\|0\|( | FALSE |
| KIAA0232 | Q8IWC1 | MAP7D3 | bait_int | 0 | 0.98791256 | 8\|12\|11 | 10.33 | 3 | 0\|2\|0\|1\|0\|0\|0\|( | FALSE |
| KIAA0232 | Q9H0W5 | CCDC8 | bait_int | 0.01 | 0.988169002 | 2\|2\|3 | 2.33 | 3 | 0\|0\|0\|0\|0\|0\|0\|( | FALSE |
| KIAA0232 | Q6ZNB6 | NFXL1 | bait_int | 0 | 0.973932992 | 5\|3\|4 | 4 | 3 | 0\|0\|0\|0\|0\|0\|0\|( | FALSE |
| KIAA0232 | Q96TA2 | YME1L1 | bait_int | 0 | 0.971597289 | 4\|4\|6 | 4.67 | 3 | 0\|0\|0\|0\|0\|0\|0\|( | FALSE |
| KIAA0232 | O75175 | CNOT3 | bait_int | 0 | 0.99375875 | 2\|3\|5 | 3.33 | 3 | 0\|0\|0\|0\|0\|0\|0\|( | FALSE |
| KIAA0232 | Q13114 | TRAF3 | bait_int | 0 | 0.989257149 | 3\|4\|6 | 4.33 | 3 | 0\|0\|0\|0\|0\|0\|0\|( | FALSE |
| KIAA0232 | Q13627 | DYRK1A | bait_bait | 0 | 0.996375154 | 11\|9\|13 | 11 | 3 | 0\|0\|0\|0\|0\|0\|0\|( | FALSE |
| TRIM23 | Q8NCN4 | RNF169 | bait_int | 0 | 0.986346183 | 3\|3\|3 | 3 | 3 | 0\|0\|0\|0\|0\|0\|0\|( | FALSE |
| TRIM23 | Q9NYZ3 | GTSE1 | bait_int | 0.03 | 0.983518388 | 2\|2\|4 | 2.67 | 3 | 2\|1\|0\|0\|0\|0\|0\|( | FALSE |
| TRIM23 | Q9UPQ9 | TNRC6B | bait_int | 0.02 | 0.987004616 | 3\|3\|3 | 3 | 3 | 2\|0\|0\|1\|0\|0\|1\|( | FALSE |
| TRIM23 | Q9BY89 | KIAA1671 | bait_int | 0 | 0.988196726 | 10\|15\|8 | 11 | 3 | 0\|0\|0\|0\|0\|0\|0\|( | FALSE |
| TRIM23 | Q96S55 | WRNIP1 | bait_int | 0.05 | 0.974075075 | 3\|1\|8 | 4 | 3 | 0\|0\|0\|0\|0\|0\|0\|( | FALSE |
| TRIM23 | Q14008 | CKAP5 | bait_int | 0 | 0.974827075 | 8\|7\|8 | 7.67 | 3 | 1\|2\|0\|1\|1\|0\|2\|1 | FALSE |

| | | | | | | | | | | |
|---|---|---|---|---|---|---|---|---|---|---|
| TRIM23 | P49790 | NUP153 | bait_int | 0.02 | 0.994441441 | 11\|12\|3 | 8.67 | 3 | 1\|1\|0\|1\|0\|1\|1\|( | FALSE |
| TRIM23 | Q5JSZ5 | PRRC2B | bait_int | 0.01 | 0.990941351 | 7\|8\|6 | 7 | 3 | 3\|3\|0\|0\|0\|0\|1\|( | FALSE |
| TRIM23 | Q6Y7W6 | GIGYF2 | bait_int | 0 | 0.975388475 | 9\|12\|6 | 9 | 3 | 2\|1\|1\|2\|0\|1\|2\|( | FALSE |
| TRIM23 | Q9Y4E8 | USP15 | bait_int | 0 | 0.99538404 | 17\|9\|16 | 14 | 3 | 0\|0\|0\|0\|0\|0\|0\|( | FALSE |
| TRIM23 | Q9NZ52 | GGA3 | bait_int | 0.01 | 0.993387949 | 2\|2\|3 | 2.33 | 3 | 0\|0\|0\|0\|0\|0\|0\|( | FALSE |
| TRIM23 | Q96CP6 | GRAMD1A | bait_int | 0 | 0.98974231 | 3\|8\|2 | 4.33 | 3 | 0\|0\|0\|0\|0\|0\|0\|( | FALSE |
| TRIM23 | Q9C0C7 | AMBRA1 | bait_int | 0.01 | 0.989887858 | 2\|2\|2 | 2 | 3 | 0\|0\|0\|0\|0\|0\|0\|( | FALSE |
| TRIM23 | O60292 | SIPA1L3 | bait_int | 0 | 0.983123328 | 6\|6\|6 | 6 | 3 | 0\|0\|0\|0\|0\|0\|0\|( | FALSE |
| TRIM23 | Q01804 | OTUD4 | bait_int | 0.04 | 0.985992709 | 4\|1\|4 | 3 | 3 | 0\|0\|0\|0\|0\|0\|0\|( | FALSE |
| TRIM23 | Q8IWC1 | MAP7D3 | bait_int | 0 | 0.989915582 | 16\|13\|13 | 14 | 3 | 0\|2\|0\|1\|0\|0\|0\|( | FALSE |
| TRIM23 | Q9BRK4 | LZTS2 | bait_int | 0 | 0.987503639 | 2\|3\|4 | 3 | 3 | 0\|0\|0\|0\|0\|0\|0\|( | FALSE |
| TRIM23 | Q5VUA4 | ZNF318 | bait_int | 0 | 0.995203837 | 16\|17\|4 | 12.33 | 3 | 0\|0\|0\|0\|0\|0\|0\|( | FALSE |
| TRIM23 | Q14738 | PPP2R5D | bait_bait | 0 | 0.997269237 | 12\|13\|10 | 11.67 | 3 | 0\|0\|2\|0\|0\|0\|0\|( | FALSE |
| TRIM23 | O43166 | SIPA1L1 | bait_int | 0 | 0.987233335 | 5\|2\|5 | 4 | 3 | 0\|0\|0\|0\|0\|0\|0\|( | FALSE |
| TRAF7 | Q12959 | DLG1 | bait_int | 0 | 0.996347431 | 13\|18\|21 | 17.33 | 3 | 0\|0\|0\|0\|0\|0\|0\|( | FALSE |
| TRAF7 | Q9Y266 | NUDC | bait_int | 0 | 0.974293398 | 24\|35\|31 | 30 | 3 | 1\|0\|5\|1\|0\|1\|1\|( | FALSE |
| TRAF7 | Q7Z6Z7 | HUWE1 | bait_int | 0 | 0.977550907 | 3\|10\|12 | 8.33 | 3 | 0\|0\|0\|0\|0\|0\|0\|( | FALSE |
| TRAF7 | O60925 | PFDN1 | bait_int | 0 | 0.985867953 | 3\|9\|7 | 6.33 | 3 | 0\|0\|0\|0\|0\|0\|0\|( | FALSE |
| TRAF7 | Q14160 | SCRIB | bait_int | 0 | 0.995210768 | 60\|62\|79 | 67 | 3 | 5\|3\|0\|4\|1\|1\|4\|3 | FALSE |
| TRAF7 | Q5T2T1 | MPP7 | bait_int | 0 | 0.997317753 | 9\|10\|13 | 10.67 | 3 | 0\|0\|0\|0\|0\|0\|0\|( | FALSE |
| TRAF7 | Q9NQP4 | PFDN4 | bait_int | 0 | 0.984218406 | 3\|6\|3 | 4 | 3 | 0\|0\|0\|0\|0\|0\|0\|( | TRUE |
| TRAF7 | Q99471 | PFDN5 | bait_int | 0 | 0.986387768 | 8\|11\|8 | 9 | 3 | 0\|0\|0\|0\|0\|0\|0\|( | FALSE |
| TRAF7 | O15212 | PFDN6 | bait_int | 0 | 0.983996618 | 6\|7\|5 | 6 | 3 | 0\|0\|1\|0\|0\|0\|0\|( | FALSE |
| TRAF7 | O14936 | CASK | bait_int | 0 | 0.993006751 | 4\|7\|10 | 7 | 3 | 0\|0\|0\|0\|0\|0\|0\|( | FALSE |
| NR3C2 | P49916 | LIG3 | bait_int | 0 | 0.978136566 | 19\|13\|9 | 13.67 | 3 | 0\|0\|0\|0\|0\|0\|0\|( | FALSE |
| NR3C2 | Q6DN90 | IQSEC1 | bait_int | 0 | 0.994656298 | 12\|12\|20 | 14.67 | 3 | 0\|0\|0\|0\|0\|0\|0\|( | FALSE |
| NR3C2 | Q9UL15 | BAG5 | bait_int | 0 | 0.97486866 | 4\|6\|4 | 4.67 | 3 | 1\|1\|0\|0\|0\|0\|1\|( | FALSE |
| NR3C2 | P78527 | PRKDC | bait_int | 0 | 0.978666778 | 68\|55\|63 | 62 | 3 | 3\|4\|2\|2\|3\|0\|3\|1 | FALSE |
| NR3C2 | Q96E52 | OMA1 | bait_int | 0 | 0.976677617 | 5\|6\|9 | 6.67 | 3 | 1\|0\|0\|0\|0\|0\|0\|( | FALSE |
| NR3C2 | Q96BR5 | COA7 | bait_int | 0 | 0.980759901 | 8\|7\|6 | 7 | 3 | 1\|0\|0\|0\|0\|0\|0\|1 | FALSE |
| NR3C2 | Q13451 | FKBP5 | bait_int | 0 | 0.99514839 | 3\|12\|13 | 9.33 | 3 | 0\|0\|0\|0\|0\|0\|0\|( | FALSE |
| NR3C2 | Q8N4T8 | CBR4 | bait_int | 0 | 0.9932424 | 5\|7\|8 | 6.67 | 3 | 0\|0\|0\|0\|0\|0\|0\|( | FALSE |
| NR3C2 | O75688 | PPM1B | bait_int | 0 | 0.98496694 | 2\|3\|3 | 2.67 | 3 | 0\|0\|0\|0\|0\|0\|0\|( | FALSE |
| NR3C2 | Q8N4J0 | CARNMT1 | bait_int | 0 | 0.997802914 | 28\|30\|34 | 30.67 | 3 | 0\|0\|0\|0\|0\|0\|0\|( | FALSE |
| RAI1 | Q709F0 | ACAD11 | bait_int | 0 | 0.986644211 | 24\|22\|22 | 22.67 | 3 | 0\|0\|0\|0\|0\|0 | FALSE |
| RAI1 | P78406 | RAE1 | bait_int | 0 | 0.985958054 | 23\|8\|16 | 15.67 | 3 | 0\|0\|0\|0\|0\|0 | FALSE |
| RAI1 | P14373 | TRIM27 | bait_int | 0 | 0.98552834 | 14\|6\|14 | 11.33 | 3 | 0\|0\|0\|0\|0\|0 | FALSE |
| RAI1 | P63010 | AP2B1 | bait_int | 0.04 | 0.983414425 | 10\|1\|11 | 7.33 | 3 | 0\|0\|0\|0\|0\|0 | FALSE |
| RAI1 | Q96CW1 | AP2M1 | bait_int | 0.04 | 0.977703386 | 8\|1\|9 | 6 | 3 | 0\|0\|0\|0\|0\|0 | FALSE |
| RAI1 | Q00610 | CLTC | bait_int | 0 | 0.980524251 | 45\|17\|37 | 33 | 3 | 0\|0\|0\|0\|0\|0 | FALSE |
| RAI1 | O75821 | EIF3G | bait_bait | 0.03 | 0.98628034 | 2\|2\|3 | 2.33 | 3 | 1\|0\|1\|1\|0\|0 | FALSE |
| RAI1 | O00505 | KPNA3 | bait_int | 0 | 0.972740882 | 8\|6\|8 | 7.33 | 3 | 0\|0\|0\|0\|0\|0 | FALSE |
| RAI1 | Q9P1Y5 | CAMSAP3 | bait_int | 0.04 | 0.995688998 | 8\|1\|10 | 6.33 | 3 | 0\|0\|0\|0\|0\|0 | FALSE |
| RAI1 | Q8N9R8 | SCAI | bait_int | 0 | 0.996881108 | 9\|6\|8 | 7.67 | 3 | 0\|0\|0\|0\|0\|0 | FALSE |
| RAI1 | Q9Y2D8 | SSX2IP | bait_int | 0 | 0.990265591 | 4\|3\|2 | 3 | 3 | 0\|0\|0\|0\|0\|0 | FALSE |
| RAI1 | Q14145 | KEAP1 | bait_int | 0 | 0.98631846 | 5\|4\|5 | 4.67 | 3 | 0\|0\|0\|0\|0\|0 | FALSE |
| RAI1 | Q9P0L0 | VAPA | bait_int | 0.04 | 0.976767719 | 6\|1\|4 | 3.67 | 3 | 0\|0\|0\|0\|0\|0 | FALSE |
| RAI1 | O94973 | AP2A2 | bait_int | 0.04 | 0.974910245 | 22\|0\|18 | 13.33 | 3 | 0\|0\|0\|0\|0\|0 | FALSE |
| RAI1 | P85037 | FOXK1 | bait_int | 0 | 0.982950056 | 6\|3\|6 | 5 | 3 | 0\|0\|0\|0\|0\|0 | FALSE |
| RAI1 | Q13114 | TRAF3 | bait_int | 0 | 0.995848408 | 20\|23\|14 | 19 | 3 | 0\|0\|0\|0\|0\|0 | FALSE |
| RAI1 | O75592 | MYCBP2 | bait_int | 0 | 0.99864848 | 86\|29\|73 | 62.67 | 3 | 0\|0\|0\|0\|0\|0 | FALSE |
| RAI1 | Q12933 | TRAF2 | bait_int | 0 | 0.992168115 | 7\|3\|6 | 5.33 | 3 | 0\|0\|0\|0\|0\|0 | FALSE |
| RAI1 | Q2M1P5 | KIF7 | bait_int | 0 | 0.989541315 | 11\|4\|8 | 7.67 | 3 | 0\|0\|0\|0\|0\|0 | FALSE |
| MAP1A | Q9Y2S7 | POLDIP2 | bait_int | 0 | 0.974241416 | 8\|9\|6 | 7.67 | 3 | 0\|0\|0\|0\|0\|0 | FALSE |
| MAP1A | O75694 | NUP155 | bait_bait | 0 | 0.980565836 | 3\|4\|6 | 4.33 | 3 | 0\|1\|0\|0\|0\|0 | FALSE |
| TCF20 | Q2M1P5 | KIF7 | bait_int | 0 | 0.990802734 | 12\|7\|10 | 9.67 | 3 | 0\|0\|0\|0\|0\|0 | FALSE |
| TCF20 | Q8WV44 | TRIM41 | bait_int | 0 | 0.982381725 | 3\|3\|2 | 2.67 | 3 | 0\|0\|0\|0\|0\|0 | FALSE |
| TCF20 | Q8N960 | CEP120 | bait_int | 0 | 0.997310822 | 7\|24\|7 | 12.67 | 3 | 0\|0\|0\|0\|0\|0 | FALSE |
| TCF20 | Q8IU81 | IRF2BP1 | bait_int | 0 | 0.991454235 | 14\|18\|17 | 16.33 | 3 | 0\|0\|0\|1\|0\|0 | FALSE |
| TCF20 | Q9NWB7 | IFT57 | bait_int | 0 | 0.997192997 | 7\|17\|12 | 12 | 3 | 0\|0\|0\|0\|0\|0 | FALSE |
| TCF20 | Q9UG01 | IFT172 | bait_int | 0 | 0.999015816 | 39\|68\|44 | 50.33 | 3 | 0\|0\|0\|0\|0\|0 | FALSE |
| TCF20 | Q9P2H3 | IFT80 | bait_int | 0 | 0.998201439 | 7\|24\|11 | 14 | 3 | 0\|0\|0\|0\|0\|0 | FALSE |
| TCF20 | Q9BQG2 | NUDT12 | bait_int | 0 | 0.995044427 | 5\|4\|4 | 4.33 | 3 | 0\|0\|0\|0\|0\|0 | FALSE |
| TCF20 | Q96AJ1 | CLUAP1 | bait_int | 0 | 0.994327082 | 2\|7\|2 | 3.67 | 3 | 0\|0\|0\|0\|0\|0 | FALSE |
| TCF20 | Q8TDR0 | TRAF3IP1 | bait_int | 0 | 0.996548426 | 3\|11\|4 | 6 | 3 | 0\|0\|0\|0\|0\|0 | FALSE |
| TCF20 | Q86WT1 | TTC30A | bait_int | 0 | 0.997071707 | 5\|13\|4 | 7.33 | 3 | 0\|0\|0\|0\|0\|0 | FALSE |
| TCF20 | Q13099 | IFT88 | bait_int | 0 | 0.99725884 | 4\|14\|6 | 8 | 3 | 0\|0\|0\|0\|0\|0 | FALSE |
| TCF20 | A0AVF1 | TTC26 | bait_int | 0 | 0.971091335 | 3\|10\|3 | 5.33 | 3 | 0\|0\|0\|0\|0\|0 | FALSE |
| TCF20 | Q8WYA0 | IFT81 | bait_int | 0 | 0.974418153 | 12\|31\|16 | 19.67 | 3 | 0\|0\|0\|0\|0\|0 | FALSE |
| TCF20 | P61962 | DCAF7 | bait_int | 0 | 0.98604469 | 2\|5\|6 | 4.33 | 3 | 0\|0\|0\|0\|0\|0 | FALSE |
| TCF20 | P78527 | PRKDC | bait_int | 0 | 0.971940367 | 20\|53\|34 | 35.67 | 3 | 1\|1\|1\|0\|2\|1 | FALSE |
| TCF20 | O94776 | MTA2 | bait_int | 0 | 0.997387061 | 15\|25\|22 | 20.67 | 3 | 0\|0\|0\|0\|0\|0 | TRUE |

| | | | | | | | | | | |
|---|---|---|---|---|---|---|---|---|---|---|
| TCF20 | Q9HCE1 | MOV10 | bait_int | 0 | 0.971777491 | 7\|5\|9 | 7 | 3 | 0\|0\|0\|0\|0\|0 | FALSE |
| TCF20 | Q8NC56 | LEMD2 | bait_int | 0 | 0.974477066 | 7\|9\|10 | 8.67 | 3 | 0\|0\|0\|0\|0\|0 | FALSE |
| TCF20 | Q93008 | USP9X | bait_int | 0 | 0.987607602 | 40\|59\|48 | 49 | 3 | 0\|0\|0\|0\|0\|0 | FALSE |
| TCF20 | P14373 | TRIM27 | bait_int | 0 | 0.980080675 | 4\|4\|10 | 6 | 3 | 0\|0\|0\|0\|0\|0 | TRUE |
| TCF20 | Q92769 | HDAC2 | bait_int | 0 | 0.972609196 | 2\|4\|5 | 3.67 | 3 | 0\|0\|0\|0\|0\|0 | FALSE |
| TCF20 | Q9BW83 | IFT27 | bait_int | 0 | 0.978618261 | 3\|8\|3 | 4.67 | 3 | 0\|0\|0\|0\|0\|0 | FALSE |
| TCF20 | P51659 | HSD17B4 | bait_int | 0 | 0.995938509 | 32\|46\|30 | 36 | 3 | 0\|0\|0\|0\|0\|0 | FALSE |
| PTK7 | Q96RQ1 | ERGIC2 | bait_int | 0 | 0.994763727 | 5\|3\|4 | 4 | 3 | 0\|0\|0\|0\|0\|0 | FALSE |
| PTK7 | Q96N66 | MBOAT7 | bait_int | 0 | 0.979339072 | 2\|2\|2 | 2 | 3 | 0\|0\|0\|0\|0\|0 | FALSE |
| PTK7 | Q07065 | CKAP4 | bait_int | 0 | 0.973579518 | 8\|10\|4 | 7.33 | 3 | 0\|0\|0\|0\|0\|0 | FALSE |
| PTK7 | O96008 | TOMM40 | bait_int | 0 | 0.971354708 | 5\|4\|4 | 4.33 | 3 | 0\|0\|0\|0\|0\|0 | FALSE |
| TAOK1 | Q9H2K8 | TAOK3 | bait_int | 0 | 0.98942349 | 2\|3\|3 | 2.67 | 3 | 0\|0\|0\|0\|0\|0 | FALSE |
| TAOK1 | O94813 | SLIT2 | bait_int | 0 | 0.99419193 | 4\|6\|4 | 4.67 | 3 | 0\|0\|0\|0\|0\|0 | FALSE |
| TAOK1 | Q8N4N3 | KLHL36 | bait_int | 0 | 0.987794735 | 2\|2\|2 | 2 | 3 | 0\|0\|0\|0\|0\|0 | FALSE |
| TAOK1 | Q9UL54 | TAOK2 | bait_int | 0 | 0.994649367 | 13\|10\|7 | 10 | 3 | 0\|0\|0\|0\|0\|0 | FALSE |
| TAOK1 | Q96I51 | RCC1L | bait_int | 0 | 0.988162071 | 5\|4\|3 | 4 | 3 | 0\|0\|0\|0\|0\|0 | FALSE |
| GIGYF1 | Q9H7N4 | SCAF1 | bait_int | 0 | 0.978153893 | 3\|3\|2 | 2.67 | 3 | 0\|0\|0\|0\|0\|0 | FALSE |
| GIGYF1 | Q99569 | PKP4 | bait_int | 0 | 0.980766832 | 7\|4\|4 | 5 | 3 | 0\|0\|0\|0\|0\|0 | FALSE |
| GIGYF1 | Q8NHP8 | PLBD2 | bait_int | 0.04 | 0.994690952 | 5\|6\|1 | 4 | 3 | 0\|0\|0\|0\|0\|0 | FALSE |
| GIGYF1 | Q8N8D1 | PDCD7 | bait_int | 0 | 0.990969074 | 5\|3\|5 | 4.33 | 3 | 0\|0\|0\|0\|0\|0 | FALSE |
| GIGYF1 | Q9Y333 | LSM2 | bait_int | 0 | 0.980510389 | 5\|2\|5 | 4 | 3 | 0\|0\|0\|0\|0\|0 | FALSE |
| GIGYF1 | P62312 | LSM6 | bait_int | 0 | 0.982326278 | 3\|2\|4 | 3 | 3 | 0\|0\|0\|0\|0\|0 | FALSE |
| GIGYF1 | Q9UKV8 | AGO2 | bait_int | 0 | 0.978278649 | 3\|5\|4 | 4 | 3 | 0\|0\|0\|0\|0\|0 | FALSE |
| GIGYF1 | Q9UKF6 | CPSF3 | bait_int | 0 | 0.977821211 | 6\|8\|9 | 7.67 | 3 | 0\|0\|0\|0\|0\|0 | FALSE |
| GIGYF1 | O60568 | PLOD3 | bait_int | 0 | 0.984759014 | 11\|2\|10 | 7.67 | 3 | 0\|0\|0\|0\|0\|0 | FALSE |
| GIGYF1 | O95104 | SCAF4 | bait_int | 0 | 0.988127417 | 5\|8\|9 | 7.33 | 3 | 0\|0\|0\|0\|0\|0 | FALSE |
| GIGYF1 | Q14145 | KEAP1 | bait_int | 0 | 0.993984004 | 17\|18\|16 | 17 | 3 | 0\|0\|0\|0\|0\|0 | FALSE |
| GIGYF1 | O95639 | CPSF4 | bait_int | 0 | 0.987184819 | 4\|5\|2 | 3.67 | 3 | 0\|0\|0\|0\|0\|0 | FALSE |
| GIGYF1 | Q10570 | CPSF1 | bait_int | 0 | 0.997005864 | 33\|27\|22 | 27.33 | 3 | 0\|0\|0\|0\|0\|0 | FALSE |
| GIGYF1 | Q16637 | SMN2 | bait_int | 0.04 | 0.985583787 | 6\|1\|6 | 4.33 | 3 | 0\|0\|0\|0\|0\|0 | FALSE |
| GIGYF1 | P24928 | POLR2A | bait_int | 0 | 0.978507368 | 2\|3\|2 | 2.33 | 3 | 0\|0\|0\|0\|0\|0 | FALSE |
| GIGYF1 | Q96LT9 | RNPC3 | bait_int | 0 | 0.997165274 | 6\|7\|10 | 7.67 | 3 | 0\|0\|0\|0\|0\|0 | FALSE |
| GIGYF1 | Q9Y657 | SPIN1 | bait_int | 0 | 0.995581569 | 8\|4\|3 | 5 | 3 | 0\|0\|0\|0\|0\|0 | FALSE |
| GIGYF1 | Q9UHI6 | DDX20 | bait_int | 0.04 | 0.973939923 | 14\|0\|12 | 8.67 | 3 | 0\|0\|0\|0\|0\|0 | FALSE |
| GIGYF1 | Q9C0J8 | WDR33 | bait_int | 0 | 0.995654344 | 16\|14\|12 | 14 | 3 | 0\|0\|0\|0\|0\|0 | FALSE |
| GIGYF1 | Q8NBJ5 | COLGALT1 | bait_int | 0 | 0.989284873 | 3\|3\|4 | 3.33 | 3 | 0\|0\|0\|0\|0\|0 | FALSE |
| GIGYF1 | Q13617 | CUL2 | bait_int | 0 | 0.981293578 | 7\|4\|4 | 5 | 3 | 0\|0\|0\|0\|0\|0 | FALSE |
| GIGYF1 | O75643 | SNRNP200 | bait_int | 0 | 0.988730403 | 114\|60\|105 | 93 | 3 | 0\|0\|0\|0\|0\|0 | FALSE |
| GIGYF1 | Q8TEQ6 | GEMIN5 | bait_int | 0 | 0.977349912 | 6\|2\|3 | 3.67 | 3 | 0\|0\|0\|0\|0\|0 | FALSE |
| GIGYF1 | Q15029 | EFTUD2 | bait_int | 0 | 0.979214316 | 63\|30\|49 | 47.33 | 3 | 1\|0\|0\|0\|0\|0 | FALSE |
| GIGYF1 | P36776 | LONP1 | bait_int | 0 | 0.996527633 | 32\|56\|30 | 39.33 | 3 | 0\|0\|0\|0\|0\|0 | TRUE |
| GIGYF1 | P08621 | SNRNP70 | bait_int | 0 | 0.982832231 | 49\|55\|43 | 49 | 3 | 0\|0\|0\|0\|0\|0 | FALSE |
| GIGYF1 | P57678 | GEMIN4 | bait_int | 0.04 | 0.978639054 | 10\|1\|5 | 5.33 | 3 | 0\|0\|1\|0\|0\|0 | FALSE |
| GIGYF1 | O75821 | EIF3G | bait_bait | 0.02 | 0.988266035 | 3\|4\|2 | 3 | 3 | 1\|0\|1\|1\|0\|0 | FALSE |
| GIGYF1 | O43290 | SART1 | bait_int | 0 | 0.987683841 | 37\|12\|32 | 27 | 3 | 0\|0\|0\|0\|0\|0 | FALSE |
| GIGYF1 | Q6P2Q9 | PRPF8 | bait_int | 0 | 0.991502752 | 122\|64\|104 | 96.67 | 3 | 0\|0\|0\|1\|0\|0 | FALSE |
| GIGYF1 | Q9Y4Z0 | LSM4 | bait_int | 0 | 0.971278469 | 4\|3\|4 | 3.67 | 3 | 0\|0\|0\|0\|0\|0 | FALSE |
| GIGYF1 | Q92900 | UPF1 | bait_int | 0 | 0.974459739 | 9\|16\|9 | 11.33 | 3 | 0\|0\|0\|0\|0\|0 | FALSE |
| GIGYF1 | Q8WWY3 | PRPF31 | bait_int | 0 | 0.992847341 | 21\|8\|17 | 15.33 | 3 | 0\|0\|0\|0\|0\|0 | FALSE |
| GIGYF1 | Q9HCE1 | MOV10 | bait_int | 0 | 0.976906336 | 10\|14\|8 | 10.67 | 3 | 0\|0\|0\|0\|0\|0 | FALSE |
| GIGYF1 | Q9BUQ8 | DDX23 | bait_int | 0 | 0.996118712 | 34\|15\|28 | 25.67 | 3 | 0\|0\|0\|0\|0\|0 | FALSE |
| GIGYF1 | Q86VM9 | ZC3H18 | bait_int | 0 | 0.985431308 | 4\|3\|6 | 4.33 | 3 | 1\|0\|0\|0\|0\|0 | FALSE |
| GIGYF1 | O60573 | EIF4E2 | bait_int | 0 | 0.986075879 | 31\|28\|31 | 30 | 3 | 1\|1\|1\|0\|0\|0 | TRUE |
| GIGYF1 | Q09161 | NCBP1 | bait_int | 0 | 0.983747106 | 3\|5\|4 | 4 | 3 | 0\|0\|0\|0\|0\|0 | FALSE |
| GIGYF1 | Q9NTZ6 | RBM12 | bait_int | 0.05 | 0.971916109 | 5\|0\|3 | 2.67 | 3 | 0\|0\|0\|0\|0\|0 | FALSE |
| GIGYF1 | Q8N163 | CCAR2 | bait_int | 0 | 0.973829029 | 5\|14\|10 | 9.67 | 3 | 0\|0\|0\|0\|0\|0 | FALSE |
| GIGYF1 | Q9UPQ9 | TNRC6B | bait_int | 0 | 0.989194771 | 5\|5\|2 | 4 | 3 | 0\|0\|0\|0\|0\|0 | FALSE |
| GIGYF1 | O43395 | PRPF3 | bait_int | 0 | 0.996354362 | 28\|14\|24 | 22 | 3 | 0\|0\|0\|0\|0\|0 | FALSE |
| GIGYF1 | O43172 | PRPF4 | bait_int | 0 | 0.996631596 | 31\|21\|30 | 27.33 | 3 | 0\|0\|0\|0\|0\|0 | FALSE |
| GIGYF1 | Q9P2I0 | CPSF2 | bait_int | 0 | 0.992334456 | 13\|7\|11 | 10.33 | 3 | 0\|0\|0\|0\|0\|0 | FALSE |
| GIGYF1 | Q8NDV7 | TNRC6A | bait_int | 0 | 0.980281671 | 4\|5\|2 | 3.67 | 3 | 0\|0\|0\|0\|0\|0 | FALSE |
| GIGYF1 | P61981 | YWHAG | bait_int | 0 | 0.977096935 | 17\|21\|18 | 18.67 | 3 | 0\|0\|0\|0\|0\|0 | TRUE |
| GIGYF1 | Q53GS9 | USP39 | bait_int | 0 | 0.986249151 | 19\|10\|15 | 14.67 | 3 | 0\|0\|0\|0\|0\|0 | FALSE |
| GIGYF1 | O94906 | PRPF6 | bait_int | 0 | 0.994982049 | 43\|26\|33 | 34 | 3 | 0\|0\|0\|0\|0\|0 | FALSE |
| GIGYF1 | P09012 | SNRPA | bait_int | 0 | 0.979436104 | 15\|16\|16 | 15.67 | 3 | 0\|0\|0\|0\|0\|0 | FALSE |
| TLK2 | Q9P2N7 | KLHL13 | bait_int | 0 | 0.993602806 | 10\|8\|8 | 8.67 | 3 | 0\|0\|0\|0\|0\|0 | FALSE |
| TLK2 | Q9P2J3 | KLHL9 | bait_int | 0 | 0.992341387 | 8\|7\|7 | 7.33 | 3 | 0\|0\|0\|0\|0\|0 | FALSE |
| TLK2 | Q13618 | CUL3 | bait_int | 0 | 0.998073218 | 21\|12\|14 | 15.67 | 3 | 0\|0\|0\|0\|0\|0 | FALSE |
| TLK2 | Q9UKI8 | TLK1 | bait_int | 0 | 0.998690065 | 31\|15\|22 | 22.67 | 3 | 0\|0\|0\|0\|0\|0 | TRUE |
| MYT1L | P49916 | LIG3 | bait_int | 0 | 0.984045134 | 32\|19\|24 | 25 | 3 | 0\|0\|0\|0\|0\|0 | FALSE |
| MYT1L | P46013 | MKI67 | bait_int | 0 | 0.988460099 | 85\|14\|46 | 48.33 | 3 | 0\|0\|0\|0\|0\|0 | FALSE |
| MYT1L | P02545 | LMNA | bait_int | 0 | 0.975350356 | 14\|12\|7 | 11 | 3 | 0\|0\|0\|0\|0\|0 | FALSE |

| | | | | | | | | | | |
|---|---|---|---|---|---|---|---|---|---|---|
| MYT1L | P11388 | TOP2A | bait_int | 0 | 0.978216271 | 22\|7\|13 | 14 | 3 | 0\|0\|0\|0\|0\|0 | FALSE |
| MYT1L | Q709F0 | ACAD11 | bait_int | 0 | 0.978812326 | 12\|4\|12 | 9.33 | 3 | 0\|0\|0\|0\|0\|0 | FALSE |
| MYT1L | P78406 | RAE1 | bait_int | 0 | 0.989319527 | 20\|28\|25 | 24.33 | 3 | 0\|0\|0\|0\|0\|0 | FALSE |
| MYT1L | Q8NC56 | LEMD2 | bait_int | 0 | 0.981688638 | 18\|17\|15 | 16.67 | 3 | 0\|0\|0\|0\|0\|0 | FALSE |
| MYT1L | Q14676 | MDC1 | bait_int | 0 | 0.982374794 | 13\|4\|7 | 8 | 3 | 0\|0\|0\|0\|0\|0 | FALSE |
| MYT1L | Q9GZS1 | POLR1E | bait_int | 0.04 | 0.975942945 | 5\|1\|5 | 3.67 | 3 | 0\|0\|0\|0\|0\|0 | FALSE |
| MYT1L | Q9P2E9 | RRBP1 | bait_int | 0.04 | 0.972283445 | 30\|0\|11 | 13.67 | 3 | 0\|0\|0\|0\|0\|0 | FALSE |
| MYT1L | P78527 | PRKDC | bait_int | 0 | 0.977537046 | 72\|41\|55 | 56 | 3 | 1\|1\|1\|0\|2\|1 | FALSE |
| MYT1L | O60341 | KDM1A | bait_int | 0 | 0.991627507 | 10\|7\|2 | 6.33 | 3 | 0\|0\|0\|0\|0\|0 | FALSE |
| MYT1L | P17544 | ATF7 | bait_int | 0 | 0.989603693 | 5\|2\|2 | 3 | 3 | 0\|0\|0\|0\|0\|0 | FALSE |
| MYT1L | Q9Y2X0 | MED16 | bait_int | 0 | 0.991599784 | 5\|2\|5 | 4 | 3 | 0\|0\|0\|0\|0\|0 | FALSE |
| MYT1L | Q9ULK4 | MED23 | bait_int | 0 | 0.995775634 | 6\|2\|8 | 5.33 | 3 | 0\|0\|0\|0\|0\|0 | FALSE |
| MYT1L | Q9NX70 | MED29 | bait_int | 0.05 | 0.989201702 | 5\|1\|3 | 3 | 3 | 0\|0\|0\|0\|0\|0 | FALSE |
| MYT1L | O75446 | SAP30 | bait_int | 0 | 0.990740356 | 5\|2\|4 | 3.67 | 3 | 0\|0\|0\|0\|0\|0 | FALSE |
| MYT1L | Q9NVC6 | MED17 | bait_int | 0 | 0.991260171 | 10\|4\|6 | 6.67 | 3 | 0\|0\|0\|0\|0\|0 | FALSE |
| MYT1L | Q9NPJ6 | MED4 | bait_int | 0 | 0.992147323 | 4\|3\|5 | 4 | 3 | 0\|0\|0\|0\|0\|0 | FALSE |
| MYT1L | O60244 | MED14 | bait_int | 0 | 0.994233515 | 9\|2\|10 | 7 | 3 | 0\|0\|0\|0\|0\|0 | FALSE |
| MYT1L | O75448 | MED24 | bait_int | 0.04 | 0.992857737 | 6\|1\|6 | 4.33 | 3 | 0\|0\|0\|0\|0\|0 | FALSE |
| MYT1L | Q96RN5 | MED15 | bait_int | 0 | 0.990892835 | 9\|5\|7 | 7 | 3 | 0\|0\|0\|0\|0\|0 | FALSE |
| MYT1L | Q9H944 | MED20 | bait_int | 0 | 0.990012614 | 3\|2\|2 | 2.33 | 3 | 0\|0\|0\|0\|0\|0 | FALSE |
| MYT1L | O95402 | MED26 | bait_int | 0 | 0.996032076 | 9\|2\|4 | 5 | 3 | 0\|0\|0\|0\|0\|0 | FALSE |
| MYT1L | P0C2W1 | FBXO45 | bait_int | 0 | 0.996208813 | 7\|11\|8 | 8.67 | 3 | 0\|0\|0\|0\|0\|0 | FALSE |
| MYT1L | Q6IC98 | GRAMD4 | bait_int | 0.05 | 0.977062281 | 6\|0\|3 | 3 | 3 | 0\|0\|0\|0\|0\|0 | FALSE |
| MYT1L | Q8NCJ5 | SPRYD3 | bait_int | 0 | 0.995460279 | 5\|5\|2 | 4 | 3 | 0\|0\|0\|0\|0\|0 | FALSE |
| MYT1L | Q96BD5 | PHF21A | bait_bait | 0 | 0.99594544 | 7\|6\|4 | 5.67 | 3 | 0\|0\|0\|0\|0\|0 | FALSE |
| MYT1L | Q15648 | MED1 | bait_int | 0 | 0.99189088 | 10\|3\|6 | 6.33 | 3 | 0\|0\|0\|0\|0\|0 | FALSE |
| MYT1L | Q96ST3 | SIN3A | bait_bait | 0 | 0.998045494 | 45\|11\|29 | 28.33 | 3 | 0\|0\|0\|0\|0\|0 | FALSE |
| MYT1L | P22670 | RFX1 | bait_int | 0 | 0.973420108 | 4\|2\|5 | 3.67 | 3 | 0\|0\|0\|0\|0\|0 | FALSE |
| MYT1L | Q6SPF0 | SAMD1 | bait_int | 0 | 0.985576856 | 12\|7\|11 | 10 | 3 | 0\|0\|0\|0\|0\|0 | FALSE |
| MYT1L | A0JLT2 | MED19 | bait_int | 0 | 0.985549133 | 5\|2\|2 | 3 | 3 | 0\|0\|0\|0\|0\|0 | FALSE |
| MYT1L | A0A1W2PQ72 | LOC1004213 | bait_int | 0 | 0.982430241 | 9\|3\|5 | 5.67 | 3 | 0\|0\|0\|0\|0\|0 | FALSE |
| MYT1L | Q8IXQ5 | KLHL7 | bait_int | 0.04 | 0.992618622 | 11\|1\|8 | 6.67 | 3 | 0\|0\|0\|0\|0\|0 | FALSE |
| MYT1L | O75592 | MYCBP2 | bait_int | 0 | 0.999237604 | 202\|217\|17 | 196.33 | 3 | 0\|0\|0\|0\|0\|0 | FALSE |
| MYT1L | P39880 | CUX1 | bait_int | 0 | 0.98365354 | 9\|5\|6 | 6.67 | 3 | 0\|0\|0\|0\|0\|0 | FALSE |
| MYT1L | P54198 | HIRA | bait_int | 0 | 0.980891587 | 11\|5\|9 | 8.33 | 3 | 0\|0\|0\|0\|0\|0 | FALSE |
| MYT1L | Q5T5X7 | BEND3 | bait_int | 0 | 0.984980801 | 19\|8\|12 | 13 | 3 | 0\|0\|0\|0\|0\|0 | FALSE |
| MYT1L | Q6ZU65 | UBN2 | bait_int | 0 | 0.977606354 | 10\|3\|5 | 6 | 3 | 0\|0\|0\|0\|0\|0 | FALSE |
| MYT1L | Q02880 | TOP2B | bait_int | 0 | 0.978708363 | 10\|4\|5 | 6.33 | 3 | 0\|0\|0\|0\|0\|0 | FALSE |
| TCF7L2 | P35222 | CTNNB1 | bait_bait | 0 | 0.99006113 | 3\|2\|2 | 2.33 | 3 | 0\|0\|0\|0\|0\|0 | TRUE |
| TCF7L2 | P49916 | LIG3 | bait_int | 0 | 0.981979734 | 19\|23\|18 | 20 | 3 | 0\|0\|0\|0\|0\|0 | FALSE |
| TCF7L2 | P02545 | LMNA | bait_int | 0 | 0.972661177 | 8\|15\|4 | 9 | 3 | 0\|0\|0\|0\|0\|0 | FALSE |
| TCF7L2 | Q8NC56 | LEMD2 | bait_int | 0 | 0.98099555 | 11\|21\|14 | 15.33 | 3 | 0\|0\|0\|0\|0\|0 | FALSE |
| TCF7L2 | Q00610 | CLTC | bait_int | 0 | 0.986616487 | 65\|64\|77 | 68.67 | 3 | 0\|0\|0\|0\|0\|0 | FALSE |
| TCF7L2 | P78527 | PRKDC | bait_int | 0 | 0.975606798 | 39\|61\|42 | 47.33 | 3 | 1\|1\|1\|0\|2\|1 | FALSE |
| TCF7L2 | Q7L0Y3 | TRMT10C | bait_int | 0 | 0.982000527 | 21\|22\|21 | 21.33 | 3 | 0\|0\|0\|0\|0\|0 | FALSE |
| TCF7L2 | P22670 | RFX1 | bait_int | 0.05 | 0.972096311 | 1\|3\|6 | 3.33 | 3 | 0\|0\|0\|0\|0\|0 | FALSE |
| TCF7L2 | Q02447 | SP3 | bait_int | 0 | 0.972373546 | 4\|5\|3 | 4 | 3 | 0\|0\|0\|0\|0\|0 | FALSE |
| TCF7L2 | P09497 | CLTB | bait_int | 0 | 0.973808237 | 2\|2\|4 | 2.67 | 3 | 0\|0\|0\|0\|0\|0 | FALSE |
| TCF7L2 | A0A1W2PQ72 | LOC1004213 | bait_int | 0 | 0.978202409 | 3\|4\|4 | 3.67 | 3 | 0\|0\|0\|0\|0\|0 | FALSE |
| TCF7L2 | P39880 | CUX1 | bait_int | 0 | 0.982638167 | 4\|6\|8 | 6 | 3 | 0\|0\|0\|0\|0\|0 | FALSE |
| TCF7L2 | P54198 | HIRA | bait_int | 0 | 0.977169709 | 3\|7\|8 | 6 | 3 | 0\|0\|0\|0\|0\|0 | FALSE |
| TCF7L2 | Q5T5X7 | BEND3 | bait_int | 0 | 0.986235289 | 12\|18\|16 | 15.33 | 3 | 0\|0\|0\|0\|0\|0 | FALSE |
| TCF7L2 | Q6ZU65 | UBN2 | bait_int | 0 | 0.978770741 | 4\|6\|10 | 6.67 | 3 | 0\|0\|0\|0\|0\|0 | FALSE |
| TCF7L2 | Q02880 | TOP2B | bait_int | 0 | 0.973291887 | 4\|3\|5 | 4 | 3 | 0\|0\|0\|0\|0\|0 | FALSE |
| MED13L | Q93074 | MED12 | bait_int | 0 | 0.998454416 | 31\|18\|18 | 22.33 | 3 | 0\|0\|0\|0\|0\|0 | TRUE |
| MED13L | Q9Y4A5 | TRRAP | bait_int | 0 | 0.991960189 | 12\|4\|3 | 6.33 | 3 | 0\|0\|0\|0\|0\|0 | FALSE |
| MED13L | Q9NX70 | MED29 | bait_int | 0 | 0.988314551 | 3\|2\|3 | 2.67 | 3 | 0\|0\|0\|0\|0\|0 | TRUE |
| MED13L | Q9NVC6 | MED17 | bait_int | 0 | 0.986810552 | 5\|2\|3 | 3.33 | 3 | 0\|0\|0\|0\|0\|0 | TRUE |
| MED13L | Q96RN5 | MED15 | bait_int | 0 | 0.984259991 | 4\|2\|2 | 2.67 | 3 | 0\|0\|0\|0\|0\|0 | TRUE |
| MED13L | Q9H1B7 | IRF2BPL | bait_bait | 0 | 0.996714767 | 26\|14\|17 | 19 | 3 | 1\|1\|0\|1\|2\|1 | FALSE |
| MED13L | Q709F0 | ACAD11 | bait_int | 0 | 0.972020072 | 9\|2\|5 | 5.33 | 3 | 0\|0\|0\|0\|0\|0 | FALSE |
| ANKRD11 | P30414 | NKTR | bait_int | 0 | 0.991343342 | 8\|7\|4 | 6.33 | 3 | 0\|0\|0\|0\|0\|0 | FALSE |
| ANKRD11 | Q7KZF4 | SND1 | bait_int | 0 | 0.978347957 | 19\|4\|9 | 10.67 | 3 | 0\|0\|0\|0\|0\|0 | FALSE |
| ANKRD11 | P61981 | YWHAG | bait_int | 0 | 0.979671754 | 26\|20\|24 | 23.33 | 3 | 0\|0\|0\|0\|0\|0 | FALSE |
| ANKRD11 | P19784 | CSNK2A2 | bait_int | 0 | 0.980953965 | 29\|30\|31 | 30 | 3 | 0\|0\|0\|0\|0\|0 | FALSE |
| ANKRD11 | O60684 | KPNA6 | bait_int | 0 | 0.989406163 | 9\|8\|9 | 8.67 | 3 | 0\|0\|0\|0\|0\|0 | FALSE |
| ANKRD11 | O75147 | OBSL1 | bait_int | 0 | 0.980704454 | 2\|5\|4 | 3.67 | 3 | 0\|0\|0\|0\|0\|0 | FALSE |
| ANKRD11 | Q9UKX7 | NUP50 | bait_int | 0 | 0.985874884 | 4\|2\|3 | 3 | 3 | 0\|0\|0\|0\|0\|0 | FALSE |
| ANKRD11 | Q9Y5B9 | SUPT16H | bait_int | 0 | 0.982853024 | 77\|72\|72 | 73.67 | 3 | 0\|0\|0\|0\|0\|0 | FALSE |
| ANKRD11 | Q6P1J9 | CDC73 | bait_int | 0 | 0.996063265 | 22\|14\|11 | 15.67 | 3 | 0\|0\|0\|0\|0\|0 | FALSE |
| ANKRD11 | Q6PI48 | DARS2 | bait_int | 0 | 0.982284692 | 4\|2\|3 | 3 | 3 | 0\|0\|0\|0\|0\|0 | FALSE |
| ANKRD11 | Q14683 | SMC1A | bait_int | 0 | 0.993540428 | 15\|43\|31 | 29.67 | 3 | 0\|0\|0\|0\|0\|0 | FALSE |

<table>
<tr><td>ANKRD11</td><td>Q96ST2</td><td>IWS1</td><td>bait_int</td><td>0.01</td><td>0.984835253</td><td>9|3|5</td><td>5.67</td><td>3</td><td>0|0|1|0|1|1</td><td>FALSE</td></tr>
<tr><td>ANKRD11</td><td>Q9BZK7</td><td>TBL1XR1</td><td>bait_bait</td><td>0</td><td>0.99220277</td><td>7|9|9</td><td>8.33</td><td>3</td><td>0|0|0|0|0|0</td><td>FALSE</td></tr>
<tr><td>ANKRD11</td><td>Q9GZS3</td><td>WDR61</td><td>bait_int</td><td>0</td><td>0.984343161</td><td>9|5|8</td><td>7.33</td><td>3</td><td>0|0|0|0|0|0</td><td>FALSE</td></tr>
<tr><td>ANKRD11</td><td>Q8NDT2</td><td>RBM15B</td><td>bait_int</td><td>0</td><td>0.99610485</td><td>13|12|10</td><td>11.67</td><td>3</td><td>0|0|0|0|0|0</td><td>FALSE</td></tr>
<tr><td>ANKRD11</td><td>Q8WVC0</td><td>LEO1</td><td>bait_int</td><td>0</td><td>0.986373907</td><td>8|8|3</td><td>6.33</td><td>3</td><td>0|0|0|0|0|0</td><td>FALSE</td></tr>
<tr><td>ANKRD11</td><td>Q92759</td><td>GTF2H4</td><td>bait_int</td><td>0</td><td>0.991544337</td><td>6|6|5</td><td>5.67</td><td>3</td><td>0|0|0|0|0|0</td><td>FALSE</td></tr>
<tr><td>ANKRD11</td><td>Q9NTI5</td><td>PDS5B</td><td>bait_int</td><td>0</td><td>0.990414605</td><td>5|5|2</td><td>4</td><td>3</td><td>0|0|0|0|0|0</td><td>FALSE</td></tr>
<tr><td>ANKRD11</td><td>Q8N7H5</td><td>PAF1</td><td>bait_int</td><td>0</td><td>0.991031452</td><td>17|10|15</td><td>14</td><td>3</td><td>0|0|0|0|0|0</td><td>FALSE</td></tr>
<tr><td>ANKRD11</td><td>Q8N3U4</td><td>STAG2</td><td>bait_int</td><td>0</td><td>0.99658308</td><td>8|13|8</td><td>9.67</td><td>3</td><td>0|0|0|0|0|0</td><td>FALSE</td></tr>
<tr><td>ANKRD11</td><td>P19447</td><td>ERCC3</td><td>bait_int</td><td>0</td><td>0.997858361</td><td>21|24|24</td><td>23</td><td>3</td><td>0|0|0|0|0|0</td><td>FALSE</td></tr>
<tr><td>ANKRD11</td><td>P32780</td><td>GTF2H1</td><td>bait_int</td><td>0</td><td>0.996784076</td><td>10|10|13</td><td>11</td><td>3</td><td>0|0|0|0|0|0</td><td>FALSE</td></tr>
<tr><td>ANKRD11</td><td>O14646</td><td>CHD1</td><td>bait_int</td><td>0</td><td>0.995536519</td><td>19|12|11</td><td>14</td><td>3</td><td>0|0|0|0|0|0</td><td>FALSE</td></tr>
<tr><td>ANKRD11</td><td>O15131</td><td>KPNA5</td><td>bait_int</td><td>0</td><td>0.984142166</td><td>4|2|3</td><td>3</td><td>3</td><td>0|0|0|0|0|0</td><td>FALSE</td></tr>
<tr><td>ANKRD11</td><td>O15379</td><td>HDAC3</td><td>bait_int</td><td>0</td><td>0.980059883</td><td>4|3|2</td><td>3</td><td>3</td><td>0|0|0|0|0|0</td><td>TRUE</td></tr>
<tr><td>ANKRD11</td><td>O60216</td><td>RAD21</td><td>bait_int</td><td>0</td><td>0.994559266</td><td>3|4|2</td><td>3</td><td>3</td><td>0|0|0|0|0|0</td><td>FALSE</td></tr>
<tr><td>ANKRD11</td><td>Q6PD62</td><td>CTR9</td><td>bait_int</td><td>0</td><td>0.991357203</td><td>23|13|7</td><td>14.33</td><td>3</td><td>0|0|0|0|0|0</td><td>FALSE</td></tr>
<tr><td>ANKRD11</td><td>Q6ZYL4</td><td>GTF2H5</td><td>bait_int</td><td>0</td><td>0.990355692</td><td>2|2|2</td><td>2</td><td>3</td><td>0|0|0|0|0|0</td><td>FALSE</td></tr>
<tr><td>ANKRD11</td><td>Q13889</td><td>GTF2H3</td><td>bait_int</td><td>0</td><td>0.995581569</td><td>4|4|7</td><td>5</td><td>3</td><td>0|0|0|0|0|0</td><td>FALSE</td></tr>
<tr><td>ANKRD11</td><td>P57772</td><td>EEFSEC</td><td>bait_int</td><td>0</td><td>0.976642963</td><td>5|2|3</td><td>3.33</td><td>3</td><td>0|0|0|0|0|0</td><td>FALSE</td></tr>
<tr><td>ANKRD11</td><td>Q96T37</td><td>RBM15</td><td>bait_int</td><td>0</td><td>0.987621464</td><td>41|35|39</td><td>38.33</td><td>3</td><td>0|0|0|0|0|0</td><td>FALSE</td></tr>
<tr><td>ANKRD11</td><td>P30084</td><td>ECHS1</td><td>bait_int</td><td>0</td><td>0.99252159</td><td>17|10|19</td><td>15.33</td><td>3</td><td>0|0|0|0|0|0</td><td>FALSE</td></tr>
<tr><td>ANKRD11</td><td>Q9UQE7</td><td>SMC3</td><td>bait_int</td><td>0</td><td>0.994247377</td><td>25|34|23</td><td>27.33</td><td>3</td><td>0|0|0|0|0|0</td><td>FALSE</td></tr>
<tr><td>ANKRD11</td><td>O75821</td><td>EIF3G</td><td>bait_bait</td><td>0.01</td><td>0.990608669</td><td>7|2|4</td><td>4.33</td><td>3</td><td>1|0|1|1|0|0</td><td>FALSE</td></tr>
<tr><td>ANKRD11</td><td>P52292</td><td>KPNA2</td><td>bait_int</td><td>0</td><td>0.980787624</td><td>59|45|51</td><td>51.67</td><td>3</td><td>0|0|0|0|0|0</td><td>FALSE</td></tr>
<tr><td>ANKRD11</td><td>P52294</td><td>KPNA1</td><td>bait_int</td><td>0</td><td>0.988300689</td><td>16|11|18</td><td>15</td><td>3</td><td>0|0|0|0|0|0</td><td>FALSE</td></tr>
<tr><td>ANKRD11</td><td>P49790</td><td>NUP153</td><td>bait_int</td><td>0</td><td>0.991384927</td><td>3|6|5</td><td>4.67</td><td>3</td><td>0|0|0|0|0|0</td><td>FALSE</td></tr>
<tr><td>ANKRD11</td><td>Q14204</td><td>DYNC1H1</td><td>bait_bait</td><td>0.05</td><td>0.972619592</td><td>9|0|3</td><td>4</td><td>3</td><td>1|0|0|0|0|0</td><td>FALSE</td></tr>
<tr><td>ANKRD11</td><td>O00629</td><td>KPNA4</td><td>bait_int</td><td>0</td><td>0.981570813</td><td>22|19|18</td><td>19.67</td><td>3</td><td>0|0|0|0|0|0</td><td>FALSE</td></tr>
<tr><td>ANKRD11</td><td>O00505</td><td>KPNA3</td><td>bait_int</td><td>0</td><td>0.984232267</td><td>26|23|15</td><td>21.33</td><td>3</td><td>0|0|0|0|0|0</td><td>FALSE</td></tr>
<tr><td>IRF2BPL</td><td>Q4G0X4</td><td>KCTD21</td><td>bait_int</td><td>0</td><td>0.995834546</td><td>5|9|8</td><td>7.33</td><td>3</td><td>0|0|0|0|0|0</td><td>FALSE</td></tr>
<tr><td>IRF2BPL</td><td>Q9NQH7</td><td>XPNPEP3</td><td>bait_int</td><td>0</td><td>0.98814821</td><td>3|2|6</td><td>3.67</td><td>3</td><td>0|0|0|0|0|0</td><td>FALSE</td></tr>
<tr><td>IRF2BPL</td><td>Q8IU81</td><td>IRF2BP1</td><td>bait_int</td><td>0</td><td>0.990664116</td><td>14|21|7</td><td>14</td><td>3</td><td>0|0|0|1|0|0</td><td>TRUE</td></tr>
<tr><td>IRF2BPL</td><td>Q9UGU0</td><td>TCF20</td><td>bait_bait</td><td>0</td><td>0.996174159</td><td>2|6|16</td><td>8</td><td>3</td><td>0|0|0|0|0|0</td><td>FALSE</td></tr>
<tr><td>IRF2BPL</td><td>Q9NWH9</td><td>SLTM</td><td>bait_int</td><td>0</td><td>0.990109646</td><td>3|9|10</td><td>7.33</td><td>3</td><td>0|0|0|0|0|0</td><td>FALSE</td></tr>
<tr><td>IRF2BPL</td><td>Q14135</td><td>VGLL4</td><td>bait_int</td><td>0</td><td>0.994815708</td><td>11|16|13</td><td>13.33</td><td>3</td><td>0|0|0|0|1|1</td><td>TRUE</td></tr>
<tr><td>IRF2BPL</td><td>Q6NSI4</td><td>RADX</td><td>bait_int</td><td>0</td><td>0.998191043</td><td>35|48|44</td><td>42.33</td><td>3</td><td>0|0|0|0|0|0</td><td>FALSE</td></tr>
<tr><td>IRF2BPL</td><td>Q32P28</td><td>P3H1</td><td>bait_int</td><td>0</td><td>0.98369859</td><td>9|9|8</td><td>8.67</td><td>3</td><td>0|0|0|0|0|0</td><td>FALSE</td></tr>
<tr><td>IRF2BPL</td><td>O75718</td><td>CRTAP</td><td>bait_int</td><td>0</td><td>0.984246129</td><td>9|7|6</td><td>7.33</td><td>3</td><td>0|0|0|0|0|0</td><td>FALSE</td></tr>
<tr><td>IRF2BPL</td><td>Q13118</td><td>KLF10</td><td>bait_int</td><td>0</td><td>0.992476539</td><td>2|4|2</td><td>2.67</td><td>3</td><td>0|0|0|0|0|0</td><td>FALSE</td></tr>
<tr><td>IRF2BPL</td><td>Q01664</td><td>TFAP4</td><td>bait_int</td><td>0</td><td>0.998163319</td><td>14|18|14</td><td>15.33</td><td>3</td><td>0|0|0|0|0|0</td><td>FALSE</td></tr>
<tr><td>IRF2BPL</td><td>P17544</td><td>ATF7</td><td>bait_int</td><td>0</td><td>0.99205029</td><td>3|5|6</td><td>4.67</td><td>3</td><td>0|0|0|0|0|0</td><td>FALSE</td></tr>
<tr><td>IRF2BPL</td><td>Q14160</td><td>SCRIB</td><td>bait_int</td><td>0</td><td>0.977356843</td><td>2|6|7</td><td>5</td><td>3</td><td>0|0|0|0|0|0</td><td>TRUE</td></tr>
<tr><td>IRF2BPL</td><td>P50570</td><td>DNM2</td><td>bait_int</td><td>0</td><td>0.993027543</td><td>2|2|7</td><td>3.67</td><td>3</td><td>0|0|0|0|0|0</td><td>FALSE</td></tr>
<tr><td>IRF2BPL</td><td>P32519</td><td>ELF1</td><td>bait_int</td><td>0.04</td><td>0.984460986</td><td>0|5|14</td><td>6.33</td><td>3</td><td>0|0|0|0|0|0</td><td>FALSE</td></tr>
<tr><td>SATB1</td><td>Q5T5X7</td><td>BEND3</td><td>bait_int</td><td>0.04</td><td>0.981591605</td><td>16|1|9</td><td>8.67</td><td>3</td><td>0|0|0|0|0|0</td><td>FALSE</td></tr>
<tr><td>SATB1</td><td>Q32P28</td><td>P3H1</td><td>bait_int</td><td>0</td><td>0.9843293</td><td>11|9|8</td><td>9.33</td><td>3</td><td>0|0|0|0|0|0</td><td>FALSE</td></tr>
<tr><td>SATB1</td><td>Q96LD4</td><td>TRIM47</td><td>bait_int</td><td>0</td><td>0.997425181</td><td>6|13|7</td><td>8.67</td><td>3</td><td>0|0|0|0|0|0</td><td>TRUE</td></tr>
<tr><td>SATB1</td><td>O75718</td><td>CRTAP</td><td>bait_int</td><td>0</td><td>0.982804508</td><td>7|5|7</td><td>6.33</td><td>3</td><td>0|0|0|0|0|0</td><td>FALSE</td></tr>
<tr><td>SATB1</td><td>Q8NC56</td><td>LEMD2</td><td>bait_int</td><td>0</td><td>0.979505413</td><td>16|7|17</td><td>13.33</td><td>3</td><td>0|0|0|0|0|0</td><td>FALSE</td></tr>
<tr><td>SATB1</td><td>Q709F0</td><td>ACAD11</td><td>bait_int</td><td>0</td><td>0.980649007</td><td>10|14|9</td><td>11</td><td>3</td><td>0|0|0|0|0|0</td><td>FALSE</td></tr>
<tr><td>SATB1</td><td>P49916</td><td>LIG3</td><td>bait_int</td><td>0</td><td>0.98274213</td><td>26|12|28</td><td>22</td><td>3</td><td>0|0|0|0|0|0</td><td>FALSE</td></tr>
<tr><td>TEK</td><td>P40855</td><td>PEX19</td><td>bait_int</td><td>0</td><td>0.990837388</td><td>12|17|10</td><td>13</td><td>3</td><td>0|0|0|0|0|0</td><td>FALSE</td></tr>
<tr><td>TEK</td><td>Q8NE01</td><td>CNNM3</td><td>bait_int</td><td>0</td><td>0.981546555</td><td>4|4|3</td><td>3.67</td><td>3</td><td>0|0|0|0|0|0</td><td>FALSE</td></tr>
<tr><td>TEK</td><td>Q96ER3</td><td>SAAL1</td><td>bait_int</td><td>0</td><td>0.97120916</td><td>5|2|3</td><td>3.33</td><td>3</td><td>0|0|0|0|0|0</td><td>FALSE</td></tr>
<tr><td>TEK</td><td>Q9NQC3</td><td>RTN4</td><td>bait_int</td><td>0</td><td>0.986304598</td><td>8|7|5</td><td>6.67</td><td>3</td><td>0|0|0|0|0|0</td><td>FALSE</td></tr>
<tr><td>TEK</td><td>O95197</td><td>RTN3</td><td>bait_int</td><td>0</td><td>0.977842004</td><td>8|9|8</td><td>8.33</td><td>3</td><td>0|0|0|0|0|0</td><td>FALSE</td></tr>
<tr><td>TEK</td><td>Q7Z4Q2</td><td>HEATR3</td><td>bait_int</td><td>0.04</td><td>0.974355775</td><td>7|1|5</td><td>4.33</td><td>3</td><td>0|0|0|0|0|0</td><td>FALSE</td></tr>
<tr><td>TRIP12</td><td>Q9UKV8</td><td>AGO2</td><td>bait_int</td><td>0</td><td>0.979976712</td><td>5|4|5</td><td>4.67</td><td>3</td><td>0|0|0|0|0|0</td><td>FALSE</td></tr>
<tr><td>TRIP12</td><td>O00505</td><td>KPNA3</td><td>bait_int</td><td>0</td><td>0.972179482</td><td>8|5|8</td><td>7</td><td>3</td><td>0|0|0|0|0|0</td><td>FALSE</td></tr>
<tr><td>TRIP12</td><td>Q93009</td><td>USP7</td><td>bait_int</td><td>0</td><td>0.975402337</td><td>30|21|32</td><td>27.67</td><td>3</td><td>0|0|0|0|0|0</td><td>TRUE</td></tr>
<tr><td>TRIP12</td><td>Q6PD62</td><td>CTR9</td><td>bait_int</td><td>0.04</td><td>0.985597649</td><td>8|1|9</td><td>6</td><td>3</td><td>0|0|0|0|0|0</td><td>FALSE</td></tr>
<tr><td>TRIP12</td><td>Q8WVC0</td><td>LEO1</td><td>bait_int</td><td>0.04</td><td>0.981993596</td><td>6|1|4</td><td>3.67</td><td>3</td><td>0|0|0|0|0|0</td><td>FALSE</td></tr>
<tr><td>TRIP12</td><td>Q8N7H5</td><td>PAF1</td><td>bait_int</td><td>0.04</td><td>0.985341207</td><td>10|1|7</td><td>6</td><td>3</td><td>0|0|0|0|0|0</td><td>FALSE</td></tr>
<tr><td>TRIP12</td><td>Q6P1J9</td><td>CDC73</td><td>bait_int</td><td>0</td><td>0.995321662</td><td>20|3|12</td><td>11.67</td><td>3</td><td>0|0|0|0|0|0</td><td>FALSE</td></tr>
<tr><td>DIP2A</td><td>Q8IXT5</td><td>RBM12B</td><td>bait_int</td><td>0</td><td>0.995293938</td><td>11|4|7</td><td>7.33</td><td>3</td><td>0|0|0|0|0|0</td><td>FALSE</td></tr>
<tr><td>DIP2A</td><td>P51659</td><td>HSD17B4</td><td>bait_int</td><td>0.04</td><td>0.984682774</td><td>5|1|7</td><td>4.33</td><td>3</td><td>0|0|0|0|0|0</td><td>FALSE</td></tr>
<tr><td>DIP2A</td><td>P30414</td><td>NKTR</td><td>bait_int</td><td>0</td><td>0.9953078</td><td>21|12|12</td><td>15</td><td>3</td><td>0|0|0|0|0|0</td><td>FALSE</td></tr>
<tr><td>ASXL3</td><td>Q92560</td><td>BAP1</td><td>bait_int</td><td>0</td><td>0.993505773</td><td>9|11|6</td><td>8.67</td><td>3</td><td>0|0|0|0|0|0|0|(</td><td>TRUE</td></tr>
<tr><td>ASXL3</td><td>O60885</td><td>BRD4</td><td>bait_int</td><td>0</td><td>0.997047449</td><td>18|5|5</td><td>9.33</td><td>3</td><td>0|0|0|0|0|0|0|(</td><td>FALSE</td></tr>
<tr><td>ASXL3</td><td>P51610</td><td>HCFC1</td><td>bait_int</td><td>0.02</td><td>0.971479464</td><td>16|20|4</td><td>13.33</td><td>3</td><td>2|1|2|1|2|0|1|</td><td>FALSE</td></tr>
<tr><td>CTNNB1</td><td>P19022</td><td>CDH2</td><td>bait_int</td><td>0</td><td>0.972297307</td><td>9|11|8</td><td>9.33</td><td>3</td><td>0|0|0|0|0|0|0|(</td><td>TRUE</td></tr>
<tr><td>CTNNB1</td><td>Q709F0</td><td>ACAD11</td><td>bait_int</td><td>0</td><td>0.979221247</td><td>12|15|2</td><td>9.67</td><td>3</td><td>0|0|0|0|0|0|0|(</td><td>FALSE</td></tr>
</table>

| | | | | | | | | | | |
|---|---|---|---|---|---|---|---|---|---|---|
| CTNNB1 | Q9Y265 | RUVBL1 | bait_int | 0 | 0.971063612 | 80\|76\|31 | 62.33 | 3 | 0\|0\|0\|0\|1\|0\|0\|( | TRUE |
| CTNNB1 | P10619 | CTSA | bait_int | 0 | 0.987711565 | 9\|9\|7 | 8.33 | 3 | 0\|0\|0\|0\|0\|0\|0\|( | FALSE |
| CTNNB1 | P16278 | GLB1 | bait_int | 0 | 0.982049043 | 3\|12\|5 | 6.67 | 3 | 0\|0\|0\|0\|0\|0\|0\|( | FALSE |
| CTNNB1 | Q00610 | CLTC | bait_int | 0.01 | 0.978798464 | 44\|34\|7 | 28.33 | 3 | 0\|0\|0\|1\|8\|0\|1\|3 | FALSE |
| CTNNB1 | P35221 | CTNNA1 | bait_int | 0 | 0.99745637 | 78\|85\|64 | 75.67 | 3 | 0\|0\|0\|0\|1\|0\|0\|( | TRUE |
| CTNNB1 | Q9Y230 | RUVBL2 | bait_int | 0 | 0.979699477 | 69\|66\|36 | 57 | 3 | 0\|0\|0\|0\|1\|0\|0\|( | TRUE |
| CTNNB1 | Q9UJU2 | LEF1 | bait_int | 0 | 0.99757073 | 10\|13\|5 | 9.33 | 3 | 0\|0\|0\|0\|0\|0\|0\|( | TRUE |
| CTNNB1 | Q5JTC6 | AMER1 | bait_int | 0 | 0.994379063 | 12\|13\|4 | 9.67 | 3 | 0\|0\|0\|0\|0\|0\|0\|( | TRUE |
| CTNNB1 | Q9NQB0 | TCF7L2 | bait_bait | 0 | 0.993803801 | 3\|4\|2 | 3 | 3 | 0\|0\|0\|0\|0\|0\|0\|( | TRUE |
| CTNNB1 | Q9NSA3 | CTNNBIP1 | bait_int | 0 | 0.998007374 | 13\|11\|12 | 12 | 3 | 0\|0\|0\|0\|0\|0\|0\|( | TRUE |
| CTNNB1 | P25054 | APC | bait_int | 0 | 0.999251466 | 121\|129\|85 | 111.67 | 3 | 0\|0\|0\|0\|0\|0\|0\|( | TRUE |
| CTNNB1 | O15169 | AXIN1 | bait_int | 0 | 0.999050471 | 61\|57\|6 | 41.33 | 3 | 0\|0\|0\|0\|0\|0\|0\|( | TRUE |
| CHD8 | P25440 | BRD2 | bait_int | 0 | 0.992279009 | 29\|25\|7 | 20.33 | 3 | 0\|0\|0\|0\|0\|0\|0\|( | FALSE |
| CHD8 | P19784 | CSNK2A2 | bait_int | 0 | 0.977578631 | 29\|26\|11 | 22 | 3 | 0\|0\|0\|0\|0\|0\|0\|( | FALSE |
| CHD8 | O60684 | KPNA6 | bait_int | 0 | 0.993450326 | 21\|19\|16 | 18.67 | 3 | 0\|0\|0\|0\|0\|0\|0\|( | TRUE |
| CHD8 | O15131 | KPNA5 | bait_int | 0 | 0.986207566 | 3\|5\|4 | 4 | 3 | 0\|0\|0\|0\|0\|0\|0\|( | TRUE |
| CHD8 | P78527 | PRKDC | bait_int | 0 | 0.971271538 | 40\|22\|40 | 34 | 3 | 1\|1\|3\|3\|8\|5\|0\|2 | FALSE |
| CHD8 | P52292 | KPNA2 | bait_int | 0 | 0.980094537 | 51\|61\|33 | 48.33 | 3 | 0\|0\|0\|0\|0\|0\|0\|( | FALSE |
| CHD8 | P52294 | KPNA1 | bait_int | 0 | 0.99037302 | 22\|24\|17 | 21 | 3 | 0\|0\|0\|0\|0\|0\|0\|( | TRUE |
| CHD8 | P35580 | MYH10 | bait_int | 0.05 | 0.974432015 | 16\|61\|6 | 27.67 | 3 | 0\|0\|0\|7\|18\|0\|0 | FALSE |
| CHD8 | O94973 | AP2A2 | bait_int | 0 | 0.977128124 | 2\|2\|6 | 3.33 | 3 | 0\|0\|0\|0\|0\|0\|0\|( | FALSE |
| CHD8 | Q9Y697 | NFS1 | bait_int | 0 | 0.996707836 | 20\|13\|15 | 16 | 3 | 0\|0\|0\|0\|0\|0\|0\|( | FALSE |
| SHANK2 | Q9Y2A7 | NCKAP1 | bait_int | 0 | 0.996451394 | 5\|12\|7 | 8 | 3 | 0\|0\|0\|0\|0\|0\|0\|( | FALSE |
| SHANK2 | Q8IZP0 | ABI1 | bait_int | 0 | 0.980392565 | 3\|2\|2 | 2.33 | 3 | 0\|0\|0\|0\|0\|0\|0\|( | FALSE |
| SHANK2 | Q14683 | SMC1A | bait_int | 0 | 0.979359865 | 3\|2\|7 | 4 | 3 | 0\|0\|0\|0\|0\|0\|0\|( | FALSE |
| SHANK2 | O00299 | CLIC1 | bait_int | 0 | 0.99554345 | 11\|11\|7 | 9.67 | 3 | 0\|0\|0\|0\|0\|0\|0\|( | FALSE |
| SHANK2 | Q9BYB0 | SHANK3 | bait_bait | 0 | 0.992355249 | 2\|2\|2 | 2 | 3 | 0\|0\|0\|0\|0\|0\|0\|( | FALSE |
| SHANK2 | Q9Y566 | SHANK1 | bait_int | 0 | 0.992732981 | 2\|2\|2 | 2 | 3 | 0\|0\|0\|0\|0\|0\|0\|( | FALSE |
| SHANK2 | Q96F07 | CYFIP2 | bait_int | 0 | 0.985556064 | 2\|3\|4 | 3 | 3 | 0\|0\|0\|0\|0\|0\|0\|( | FALSE |
| SHANK2 | P98175 | RBM10 | bait_int | 0 | 0.993970142 | 10\|12\|8 | 10 | 3 | 0\|0\|0\|0\|0\|0\|0\|( | FALSE |
| SYNGAP1 | P61981 | YWHAG | bait_int | 0 | 0.982305485 | 38\|31\|24 | 31 | 3 | 0\|0\|0\|0\|0\|0\|0\|( | TRUE |
| SYNGAP1 | Q04917 | YWHAH | bait_int | 0 | 0.977939036 | 34\|28\|26 | 29.33 | 3 | 0\|0\|0\|0\|1\|0\|0\|( | FALSE |
| KDM5B | Q02880 | TOP2B | bait_int | 0 | 0.980815348 | 11\|6\|6 | 7.67 | 3 | 0\|0\|0\|0\|0\|0\|0\|( | FALSE |
| KDM5B | P17066 | HSPA6 | bait_int | 0 | 0.986263013 | 13\|13\|8 | 11.33 | 3 | 0\|0\|0\|0\|0\|0\|0\|( | FALSE |
| KDM5B | P13010 | XRCC5 | bait_int | 0 | 0.973045841 | 61\|71\|41 | 57.67 | 3 | 0\|0\|0\|0\|2\|2\|0\|( | FALSE |
| KDM5B | P78527 | PRKDC | bait_int | 0 | 0.985001594 | 135\|174\|63 | 124 | 3 | 1\|1\|3\|3\|8\|5\|0\|2 | FALSE |
| KDM5B | Q14676 | MDC1 | bait_int | 0 | 0.98353918 | 15\|4\|8 | 9 | 3 | 0\|0\|0\|0\|0\|0\|0\|( | FALSE |
| KDM5B | Q9Y5B9 | SUPT16H | bait_int | 0 | 0.973600311 | 36\|43\|14 | 31 | 3 | 0\|0\|0\|0\|0\|2\|0\|1 | FALSE |
| KDM5B | P11388 | TOP2A | bait_int | 0 | 0.974979554 | 16\|8\|8 | 10.67 | 3 | 0\|0\|0\|0\|0\|0\|0\|( | FALSE |
| CELF4 | Q96DH6 | MSI2 | bait_int | 0 | 0.989153186 | 2\|8\|3 | 4.33 | 3 | 0\|0\|0\|0\|0\|0\|0\|( | FALSE |
| CELF4 | Q9Y679 | AUP1 | bait_int | 0 | 0.97220374 | 5\|3\|2 | 3.33 | 3 | 0\|0\|0\|0\|0\|0\|0\|( | FALSE |
| CELF4 | Q8NF37 | LPCAT1 | bait_int | 0 | 0.971372035 | 6\|8\|3 | 5.67 | 3 | 0\|0\|0\|0\|0\|0\|0\|( | FALSE |
| PHF12 | P17066 | HSPA6 | bait_int | 0 | 0.984779806 | 9\|12\|7 | 9.33 | 3 | 0\|0\|0\|0\|0\|0\|0\|( | FALSE |
| PHF12 | Q8WUU5 | GATAD1 | bait_int | 0 | 0.996444463 | 12\|7\|12 | 10.33 | 3 | 0\|0\|0\|0\|0\|0\|0\|( | TRUE |
| PHF12 | Q96ST3 | SIN3A | bait_bait | 0 | 0.99618109 | 12\|14\|10 | 12 | 3 | 0\|0\|0\|0\|0\|0\|0\|( | FALSE |
| PHF12 | O75182 | SIN3B | bait_int | 0 | 0.997969255 | 27\|23\|26 | 25.33 | 3 | 0\|0\|0\|0\|0\|0\|0\|( | TRUE |
| PHF12 | Q9UBU8 | MORF4L1 | bait_int | 0 | 0.984828322 | 34\|30\|25 | 29.67 | 3 | 1\|0\|0\|4\|1\|0\|3\|1 | TRUE |
| PHF12 | Q13547 | HDAC1 | bait_int | 0 | 0.974542909 | 8\|8\|8 | 8 | 3 | 0\|0\|0\|0\|0\|0\|0\|( | TRUE |
| PHF12 | Q15014 | MORF4L2 | bait_int | 0 | 0.995959302 | 39\|37\|32 | 36 | 3 | 1\|0\|0\|3\|0\|1\|1\|( | TRUE |
| EIF3G | P46013 | MKI67 | bait_int | 0 | 0.982610443 | 29\|34\|6 | 23 | 3 | 0\|0\|0\|0\|0\|0\|0\|( | FALSE |
| EIF3G | Q07157 | TJP1 | bait_int | 0 | 0.975915222 | 7\|9\|7 | 7.67 | 3 | 0\|0\|0\|0\|0\|0\|0\|( | FALSE |
| EIF3G | P60228 | EIF3E | bait_int | 0 | 0.980350979 | 12\|11\|25 | 16 | 3 | 0\|0\|0\|0\|0\|0\|0\|( | TRUE |
| EIF3G | O00303 | EIF3F | bait_int | 0 | 0.979054906 | 8\|10\|18 | 12 | 3 | 0\|0\|0\|0\|0\|0\|0\|( | TRUE |
| EIF3G | Q9NTK5 | OLA1 | bait_int | 0 | 0.984266922 | 6\|2\|7 | 5 | 3 | 0\|0\|0\|0\|0\|0\|0\|( | FALSE |
| EIF3G | Q7L2H7 | EIF3M | bait_int | 0 | 0.976837027 | 3\|8\|16 | 9 | 3 | 0\|0\|0\|0\|0\|0\|0\|( | TRUE |
| EIF3G | Q9UBQ5 | EIF3K | bait_int | 0 | 0.971070542 | 6\|5\|11 | 7.33 | 3 | 0\|0\|0\|0\|0\|0\|0\|( | TRUE |
| EIF3G | Q13347 | EIF3I | bait_int | 0 | 0.991364134 | 74\|57\|56 | 62.33 | 3 | 0\|0\|0\|0\|0\|0\|0\|( | TRUE |
| EIF3G | Q9H2P0 | ADNP | bait_bait | 0 | 0.996791007 | 7\|14\|4 | 8.33 | 3 | 0\|0\|0\|0\|0\|0\|0\|( | FALSE |
| EIF3G | Q14152 | EIF3A | bait_int | 0 | 0.984911493 | 46\|54\|91 | 63.67 | 3 | 0\|0\|0\|0\|0\|0\|0\|( | TRUE |
| EIF3G | O60232 | ZNRD2 | bait_int | 0 | 0.990081923 | 5\|3\|4 | 4 | 3 | 0\|0\|0\|0\|0\|0\|0\|( | FALSE |
| EIF3G | Q9Y262 | EIF3L | bait_int | 0 | 0.97232503 | 20\|23\|32 | 25 | 3 | 0\|0\|0\|0\|0\|0\|0\|( | TRUE |
| EIF3G | Q7L2E3 | DHX30 | bait_int | 0 | 0.981459919 | 48\|55\|33 | 45.33 | 3 | 0\|0\|0\|0\|0\|0\|0\|( | FALSE |
| EIF3G | Q15020 | SART3 | bait_int | 0 | 0.99132948 | 6\|8\|3 | 5.67 | 3 | 0\|0\|0\|0\|0\|0\|0\|( | FALSE |
| EIF3G | O75648 | TRMU | bait_int | 0 | 0.991121554 | 4\|5\|3 | 4 | 3 | 0\|0\|0\|0\|0\|0\|0\|( | FALSE |
| TCF4 | P46013 | MKI67 | bait_int | 0 | 0.971652736 | 14\|2\|10 | 8.67 | 3 | 0\|0\|0\|0\|0\|0\|0\|( | FALSE |
| TCF4 | P49916 | LIG3 | bait_int | 0 | 0.98560458 | 32\|34\|27 | 31 | 3 | 0\|0\|0\|0\|0\|0\|0\|( | FALSE |
| TCF4 | P07197 | NEFM | bait_int | 0 | 0.973458228 | 15\|2\|7 | 8 | 3 | 0\|0\|0\|0\|0\|0\|0\|( | FALSE |
| TCF4 | P18887 | XRCC1 | bait_int | 0 | 0.977592492 | 13\|13\|11 | 12.33 | 3 | 0\|0\|0\|0\|0\|0\|0\|( | FALSE |
| TCF4 | P11388 | TOP2A | bait_int | 0 | 0.972380477 | 10\|5\|11 | 8.67 | 3 | 0\|0\|0\|0\|0\|0\|0\|( | TRUE |
| TCF4 | Q9Y5B9 | SUPT16H | bait_int | 0 | 0.972678505 | 33\|32\|22 | 29 | 3 | 0\|0\|0\|0\|0\|2\|0\|1 | FALSE |
| TCF4 | Q14676 | MDC1 | bait_int | 0 | 0.980877726 | 6\|3\|11 | 6.67 | 3 | 0\|0\|0\|0\|0\|0\|0\|( | FALSE |
| TCF4 | Q96ES7 | SGF29 | bait_int | 0 | 0.990643323 | 9\|7\|4 | 6.67 | 3 | 0\|0\|0\|0\|0\|0\|0\|( | FALSE |

| | | | | | | | | | | |
|---|---|---|---|---|---|---|---|---|---|---|
| TCF4 | Q9H7D7 | WDR26 | bait_int | 0 | 0.979318279 | 8\|9\|3 | 6.67 | 3 | 0\|0\|0\|0\|0\|0\|0\|( | FALSE |
| TCF4 | Q96SB3 | PPP1R9B | bait_bait | 0 | 0.993107248 | 15\|4\|2 | 7 | 3 | 0\|0\|0\|0\|0\|0\|0\|( | FALSE |
| TCF4 | Q9UNE7 | STUB1 | bait_int | 0 | 0.975658779 | 16\|11\|10 | 12.33 | 3 | 0\|0\|0\|0\|0\|0\|0\|( | FALSE |
| TCF4 | Q9UGN5 | PARP2 | bait_int | 0 | 0.97216562 | 12\|5\|3 | 6.67 | 3 | 0\|0\|0\|0\|0\|0\|0\|( | FALSE |
| TCF4 | Q00610 | CLTC | bait_int | 0 | 0.98433623 | 72\|70\|10 | 50.67 | 3 | 0\|0\|0\|1\|8\|0\|1\|3 | FALSE |
| TCF4 | P78527 | PRKDC | bait_int | 0 | 0.985119419 | 124\|163\|89 | 125.33 | 3 | 1\|1\|3\|3\|8\|5\|0\|2 | TRUE |
| TCF4 | Q01082 | SPTBN1 | bait_int | 0 | 0.980808417 | 88\|91\|23 | 67.33 | 3 | 0\|0\|0\|9\|26\|0\|0 | FALSE |
| TCF4 | P35580 | MYH10 | bait_int | 0.01 | 0.980735643 | 79\|54\|11 | 48 | 3 | 0\|0\|0\|7\|18\|0\|0 | FALSE |
| TCF4 | Q13813 | SPTAN1 | bait_int | 0 | 0.981452988 | 95\|103\|30 | 76 | 3 | 0\|0\|0\|13\|33\|0\| | FALSE |
| TCF4 | Q14839 | CHD4 | bait_int | 0 | 0.986935307 | 5\|2\|5 | 4 | 3 | 0\|0\|0\|0\|0\|0\|0\|( | FALSE |
| TCF4 | P04637 | TP53 | bait_int | 0 | 0.972117104 | 20\|11\|21 | 17.33 | 3 | 0\|0\|0\|1\|1\|7\|0\|( | TRUE |
| TCF4 | Q12800 | TFCP2 | bait_int | 0 | 0.981078721 | 7\|4\|4 | 5 | 3 | 0\|0\|0\|0\|0\|0\|0\|( | FALSE |
| TCF4 | Q92878 | RAD50 | bait_int | 0 | 0.984821391 | 3\|5\|21 | 9.67 | 3 | 0\|0\|0\|0\|0\|0\|0\|( | FALSE |
| TCF4 | Q99081 | TCF12 | bait_int | 0 | 0.99101066 | 10\|7\|2 | 6.33 | 3 | 0\|0\|0\|0\|0\|0\|0\|( | TRUE |
| TCF4 | Q96T60 | PNKP | bait_int | 0 | 0.974736973 | 3\|7\|4 | 4.67 | 3 | 0\|0\|0\|0\|0\|0\|0\|( | FALSE |
| TCF4 | Q96MH2 | HEXIM2 | bait_int | 0 | 0.997109827 | 12\|8\|5 | 8.33 | 3 | 0\|0\|0\|0\|0\|0\|0\|( | TRUE |
| TCF4 | Q02535 | ID3 | bait_int | 0 | 0.997501421 | 11\|7\|9 | 9 | 3 | 0\|0\|0\|0\|0\|0\|0\|( | TRUE |
| TCF4 | Q02363 | ID2 | bait_int | 0 | 0.998052425 | 18\|12\|7 | 12.33 | 3 | 0\|0\|0\|0\|0\|0\|0\|( | TRUE |
| TCF4 | P15923 | TCF3 | bait_int | 0 | 0.997012794 | 15\|10\|4 | 9.67 | 3 | 0\|0\|0\|0\|0\|0\|0\|( | TRUE |
| TCF4 | P47928 | ID4 | bait_int | 0 | 0.998239559 | 24\|16\|13 | 17.67 | 3 | 0\|0\|0\|0\|0\|0\|0\|( | TRUE |
| TCF4 | P62699 | YPEL5 | bait_int | 0 | 0.976750392 | 2\|2\|2 | 2 | 3 | 0\|0\|0\|0\|0\|0\|0\|( | FALSE |
| TCF4 | Q53H96 | PYCR3 | bait_int | 0 | 0.971534911 | 4\|3\|7 | 4.67 | 3 | 0\|0\|0\|0\|0\|0\|0\|( | FALSE |
| TCF4 | P41134 | ID1 | bait_int | 0 | 0.99848907 | 22\|17\|16 | 18.33 | 3 | 0\|0\|0\|0\|0\|0\|0\|( | TRUE |
| TCF4 | Q9UNY4 | TTF2 | bait_int | 0 | 0.991267102 | 15\|5\|14 | 11.33 | 3 | 0\|0\|0\|0\|0\|0\|0\|( | FALSE |
| TCF4 | P54198 | HIRA | bait_int | 0 | 0.981258923 | 11\|12\|3 | 8.67 | 3 | 0\|0\|0\|0\|0\|0\|0\|( | FALSE |
| TCF4 | Q6ZU65 | UBN2 | bait_int | 0 | 0.979352934 | 11\|4\|6 | 7 | 3 | 0\|0\|0\|0\|0\|0\|0\|( | FALSE |
| TCF4 | Q02880 | TOP2B | bait_int | 0 | 0.973291887 | 2\|3\|7 | 4 | 3 | 0\|0\|0\|0\|0\|0\|0\|( | FALSE |
| GRIA2 | Q07065 | CKAP4 | bait_int | 0 | 0.988564062 | 48\|38\|22 | 36 | 3 | 0\|0\|0\|0\|0\|0\|0\|( | FALSE |
| GRIA2 | Q5SWX8 | ODR4 | bait_int | 0 | 0.997920739 | 20\|16\|12 | 16 | 3 | 0\|0\|0\|0\|0\|0\|0\|( | FALSE |
| GRIA2 | Q9NUQ7 | UFSP2 | bait_int | 0 | 0.99841283 | 19\|19\|13 | 17 | 3 | 0\|0\|0\|0\|0\|0\|0\|( | FALSE |
| GRIA2 | P48058 | GRIA4 | bait_int | 0 | 0.993256262 | 2\|2\|3 | 2.33 | 3 | 0\|0\|0\|1\|0\|0\|0\|( | FALSE |
| CREBBP | P05412 | JUN | bait_int | 0 | 0.971756699 | 4\|3\|3 | 3.33 | 3 | 0\|0\|0\|0\|0\|0 | TRUE |
| CREBBP | Q13627 | DYRK1A | bait_bait | 0 | 0.990594807 | 2\|3\|3 | 2.67 | 3 | 0\|0\|0\|0\|0\|0 | TRUE |
| CREBBP | P06400 | RB1 | bait_int | 0 | 0.994268169 | 11\|15\|17 | 14.33 | 3 | 0\|0\|0\|0\|0\|0 | FALSE |
| CREBBP | Q09472 | EP300 | bait_int | 0 | 0.99013737 | 6\|4\|3 | 4.33 | 3 | 0\|0\|0\|0\|0\|0 | TRUE |
| CREBBP | Q32P28 | P3H1 | bait_int | 0 | 0.990650254 | 31\|21\|17 | 23 | 3 | 0\|0\|0\|0\|0\|0 | FALSE |
| CREBBP | P28749 | RBL1 | bait_int | 0 | 0.990234402 | 5\|16\|14 | 11.67 | 3 | 0\|0\|0\|0\|0\|0 | FALSE |
| CREBBP | Q15596 | NCOA2 | bait_int | 0 | 0.996368223 | 11\|16\|16 | 14.33 | 3 | 0\|0\|0\|0\|0\|0 | TRUE |
| CREBBP | Q9ULH7 | MRTFB | bait_int | 0 | 0.99491274 | 3\|2\|5 | 3.33 | 3 | 0\|0\|0\|0\|0\|0 | FALSE |
| CREBBP | Q04206 | RELA | bait_int | 0 | 0.998128665 | 19\|11\|9 | 13 | 3 | 0\|0\|0\|0\|0\|0 | TRUE |
| CREBBP | Q969V6 | MRTFA | bait_int | 0 | 0.997723209 | 6\|8\|19 | 11 | 3 | 0\|0\|0\|0\|0\|0 | FALSE |
| CREBBP | Q15788 | NCOA1 | bait_bait | 0.05 | 0.99411569 | 6\|3\|1 | 3.33 | 3 | 0\|0\|0\|0\|0\|0 | TRUE |
| CREBBP | O75718 | CRTAP | bait_int | 0 | 0.987940284 | 18\|13\|5 | 12 | 3 | 0\|0\|0\|0\|0\|0 | FALSE |
| CREBBP | P04637 | TP53 | bait_int | 0 | 0.981799531 | 47\|42\|32 | 40.33 | 3 | 0\|0\|0\|0\|0\|0 | TRUE |
| CREBBP | P61962 | DCAF7 | bait_int | 0 | 0.989309131 | 7\|8\|5 | 6.67 | 3 | 0\|0\|0\|0\|0\|0 | FALSE |
| CREBBP | P78527 | PRKDC | bait_int | 0 | 0.976195922 | 52\|54\|44 | 50 | 3 | 0\|0\|0\|0\|0\|0 | FALSE |
| CREBBP | Q9Y6Q9 | NCOA3 | bait_int | 0 | 0.976518207 | 8\|5\|5 | 6 | 3 | 0\|0\|0\|0\|0\|0 | TRUE |
| CREBBP | P49916 | LIG3 | bait_int | 0 | 0.978136566 | 20\|13\|8 | 13.67 | 3 | 0\|0\|0\|0\|0\|0 | FALSE |
| SRPRA | Q96JJ7 | TMX3 | bait_int | 0 | 0.990407674 | 11\|6\|5 | 7.33 | 3 | 0\|0\|0\|0\|0\|0 | FALSE |
| SRPRA | Q9Y5M8 | SRPRB | bait_int | 0 | 0.984987732 | 29\|33\|37 | 33 | 3 | 0\|0\|0\|0\|0\|0 | TRUE |
| SRPRA | Q9UBV2 | SEL1L | bait_int | 0 | 0.976975645 | 2\|3\|3 | 2.67 | 3 | 0\|0\|0\|0\|0\|0 | FALSE |
| SRPRA | P31948 | STIP1 | bait_int | 0 | 0.983206498 | 51\|35\|42 | 42.67 | 3 | 0\|0\|0\|0\|0\|0 | FALSE |
| TBL1XR1 | Q13227 | GPS2 | bait_int | 0 | 0.986755105 | 10\|5\|5 | 6.67 | 3 | 0\|0\|0\|0\|0\|0 | TRUE |
| TBL1XR1 | O75376 | NCOR1 | bait_int | 0 | 0.996409809 | 54\|21\|28 | 34.33 | 3 | 0\|0\|0\|0\|0\|0 | TRUE |
| TBL1XR1 | O60907 | TBL1X | bait_int | 0 | 0.992306733 | 19\|9\|5 | 11 | 3 | 0\|0\|0\|0\|0\|0 | TRUE |
| TBL1XR1 | O15379 | HDAC3 | bait_int | 0 | 0.987164026 | 9\|7\|5 | 7 | 3 | 0\|0\|0\|0\|0\|0 | TRUE |
| TBL1XR1 | Q9Y618 | NCOR2 | bait_int | 0 | 0.994254308 | 38\|11\|14 | 21 | 3 | 0\|0\|0\|0\|0\|0 | TRUE |
| DPYSL2 | Q9BPU6 | DPYSL5 | bait_int | 0 | 0.998288075 | 36\|29\|34 | 33 | 3 | 0\|0\|0\|0\|0\|0 | TRUE |
| DPYSL2 | O14531 | DPYSL4 | bait_int | 0 | 0.996077127 | 13\|11\|12 | 12 | 3 | 0\|0\|0\|0\|0\|0 | TRUE |
| SKI | Q13627 | DYRK1A | bait_bait | 0.05 | 0.991870088 | 1\|3\|6 | 3.33 | 3 | 0\|0\|0\|0\|0\|0 | FALSE |
| SKI | Q7L5Y9 | MAEA | bait_int | 0 | 0.995716721 | 16\|12\|10 | 12.67 | 3 | 0\|0\|0\|0\|0\|0 | TRUE |
| SKI | Q9NWU2 | GID8 | bait_int | 0 | 0.994850362 | 14\|12\|10 | 12 | 3 | 0\|0\|0\|0\|0\|0 | FALSE |
| SKI | Q13485 | SMAD4 | bait_int | 0 | 0.999036609 | 39\|45\|48 | 44 | 3 | 0\|0\|0\|0\|0\|0 | TRUE |
| SKI | Q9UL63 | MKLN1 | bait_int | 0 | 0.996194952 | 11\|5\|8 | 8 | 3 | 0\|0\|0\|0\|0\|0 | TRUE |
| SKI | Q8IUR7 | ARMC8 | bait_int | 0 | 0.994143414 | 5\|6\|8 | 6.33 | 3 | 0\|0\|0\|0\|0\|0 | TRUE |
| SKI | Q6VN20 | RANBP10 | bait_int | 0 | 0.996562288 | 12\|12\|12 | 12 | 3 | 0\|0\|0\|0\|0\|0 | FALSE |
| SKI | P62699 | YPEL5 | bait_int | 0 | 0.98448871 | 5\|4\|4 | 4.33 | 3 | 0\|0\|0\|0\|0\|0 | FALSE |
| SKI | Q9H871 | RMND5A | bait_int | 0 | 0.993623598 | 9\|7\|9 | 8.33 | 3 | 0\|0\|0\|0\|0\|0 | TRUE |
| SKI | Q15796 | SMAD2 | bait_int | 0 | 0.98401741 | 2\|3\|4 | 3 | 3 | 0\|0\|0\|0\|0\|0 | TRUE |
| SKI | Q96S59 | RANBP9 | bait_int | 0 | 0.996084058 | 19\|16\|15 | 16.67 | 3 | 0\|0\|0\|0\|0\|0 | FALSE |
| SKI | P61962 | DCAF7 | bait_int | 0 | 0.994455303 | 15\|19\|21 | 18.33 | 3 | 0\|0\|0\|0\|0\|0 | FALSE |
| SKI | P16278 | GLB1 | bait_int | 0 | 0.980621283 | 6\|7\|4 | 5.67 | 3 | 0\|0\|0\|0\|0\|0 | FALSE |

| | | | | | | | | | | |
|---|---|---|---|---|---|---|---|---|---|---|
| SKI | P10619 | CTSA | bait_int | 0 | 0.97602265 | 2\|3\|2 | 2.33 | 3 | 0\|0\|0\|0\|0\|0 | FALSE |
| SKI | Q9BZF1 | OSBPL8 | bait_int | 0 | 0.985167935 | 3\|5\|11 | 6.33 | 3 | 0\|0\|0\|0\|0\|0 | FALSE |
| SKI | Q9H7D7 | WDR26 | bait_int | 0 | 0.985459032 | 13\|11\|16 | 13.33 | 3 | 0\|0\|0\|0\|0\|0 | TRUE |
| SKI | Q93008 | USP9X | bait_int | 0 | 0.984481779 | 30\|26\|41 | 32.33 | 3 | 0\|0\|0\|0\|0\|0 | FALSE |
| SKI | Q13162 | PRDX4 | bait_int | 0 | 0.972290376 | 3\|5\|4 | 4 | 3 | 0\|0\|0\|0\|0\|0 | FALSE |
| PAX5 | Q02880 | TOP2B | bait_int | 0 | 0.987018478 | 23\|12\|14 | 16.33 | 3 | 0\|0\|0\|0\|0\|0 | FALSE |
| PAX5 | Q9Y697 | NFS1 | bait_int | 0 | 0.991475028 | 5\|3\|4 | 4 | 3 | 0\|0\|0\|0\|0\|0 | FALSE |
| PAX5 | Q6ZU65 | UBN2 | bait_int | 0 | 0.985327345 | 10\|16\|15 | 13.67 | 3 | 0\|0\|0\|0\|0\|0 | FALSE |
| PAX5 | Q5T5X7 | BEND3 | bait_int | 0 | 0.983726314 | 6\|13\|14 | 11 | 3 | 0\|0\|0\|0\|0\|0 | FALSE |
| PAX5 | P54198 | HIRA | bait_int | 0 | 0.988335343 | 17\|22\|24 | 21 | 3 | 0\|0\|0\|0\|0\|0 | FALSE |
| PAX5 | P39880 | CUX1 | bait_int | 0 | 0.983130259 | 2\|10\|7 | 6.33 | 3 | 0\|0\|0\|0\|0\|0 | FALSE |
| PAX5 | Q9C0D3 | ZYG11B | bait_int | 0 | 0.991156208 | 6\|4\|2 | 4 | 3 | 0\|0\|0\|0\|0\|0 | FALSE |
| PAX5 | Q02809 | PLOD1 | bait_int | 0 | 0.990310642 | 9\|6\|8 | 7.67 | 3 | 0\|0\|0\|0\|0\|0 | FALSE |
| PAX5 | Q9UNY4 | TTF2 | bait_int | 0 | 0.983435217 | 3\|4\|4 | 3.67 | 3 | 0\|0\|0\|0\|0\|0 | FALSE |
| PAX5 | O60568 | PLOD3 | bait_int | 0 | 0.986034294 | 9\|5\|13 | 9 | 3 | 0\|0\|0\|0\|0\|0 | FALSE |
| PAX5 | Q9Y4X5 | ARIH1 | bait_int | 0 | 0.994053312 | 3\|3\|2 | 2.67 | 3 | 0\|0\|0\|0\|0\|0 | FALSE |
| PAX5 | Q96T60 | PNKP | bait_int | 0 | 0.976254834 | 5\|7\|4 | 5.33 | 3 | 0\|0\|0\|0\|0\|0 | FALSE |
| PAX5 | P06746 | POLB | bait_int | 0 | 0.985770921 | 3\|6\|6 | 5 | 3 | 0\|0\|0\|0\|0\|0 | FALSE |
| PAX5 | Q9NPG3 | UBN1 | bait_int | 0.03 | 0.987600671 | 1\|7\|6 | 4.67 | 3 | 0\|0\|0\|0\|0\|0 | FALSE |
| PAX5 | Q9UK99 | FBXO3 | bait_int | 0.03 | 0.984405539 | 8\|7\|1 | 5.33 | 3 | 0\|0\|0\|0\|0\|0 | FALSE |
| PAX5 | Q92878 | RAD50 | bait_int | 0 | 0.981744085 | 10\|7\|3 | 6.67 | 3 | 0\|0\|0\|0\|0\|0 | FALSE |
| PAX5 | O60232 | ZNRD2 | bait_int | 0 | 0.997241513 | 20\|26\|30 | 25.33 | 3 | 0\|0\|0\|0\|0\|0 | FALSE |
| PAX5 | Q14839 | CHD4 | bait_int | 0 | 0.9875175 | 3\|5\|5 | 4.33 | 3 | 0\|0\|0\|0\|0\|0 | FALSE |
| PAX5 | P13010 | XRCC5 | bait_int | 0 | 0.977724179 | 96\|82\|78 | 85.33 | 3 | 0\|0\|0\|0\|0\|0 | FALSE |
| PAX5 | Q14527 | HLTF | bait_int | 0 | 0.979158869 | 16\|10\|9 | 11.67 | 3 | 0\|0\|0\|0\|0\|0 | FALSE |
| PAX5 | P78527 | PRKDC | bait_int | 0 | 0.990587877 | 304\|253\|2[illegible] | 271.33 | 3 | 0\|0\|0\|0\|0\|0 | FALSE |
| PAX5 | Q9UGN5 | PARP2 | bait_int | 0 | 0.98011533 | 17\|12\|10 | 13 | 3 | 0\|0\|0\|0\|0\|0 | FALSE |
| PAX5 | Q9Y2S7 | POLDIP2 | bait_int | 0 | 0.985854091 | 32\|26\|16 | 24.67 | 3 | 0\|0\|0\|0\|0\|0 | FALSE |
| PAX5 | Q9Y6J0 | CABIN1 | bait_int | 0 | 0.985396654 | 4\|11\|9 | 8 | 3 | 0\|0\|0\|0\|0\|0 | FALSE |
| PAX5 | Q14676 | MDC1 | bait_int | 0 | 0.989652209 | 31\|17\|13 | 20.33 | 3 | 0\|0\|0\|0\|0\|0 | FALSE |
| PAX5 | Q01831 | XPC | bait_int | 0 | 0.993325571 | 11\|8\|7 | 8.67 | 3 | 0\|0\|0\|0\|0\|0 | FALSE |
| PAX5 | Q9Y5B9 | SUPT16H | bait_int | 0 | 0.977259811 | 56\|36\|35 | 42.33 | 3 | 0\|0\|0\|0\|0\|0 | FALSE |
| PAX5 | Q9H3G5 | CPVL | bait_int | 0 | 0.974265674 | 26\|21\|22 | 23 | 3 | 0\|0\|0\|0\|0\|0 | FALSE |
| PAX5 | P11388 | TOP2A | bait_int | 0 | 0.9812312 | 21\|19\|16 | 18.67 | 3 | 0\|0\|0\|0\|0\|0 | FALSE |
| PAX5 | P13489 | RNH1 | bait_int | 0 | 0.975222134 | 4\|5\|3 | 4 | 3 | 0\|0\|0\|0\|0\|0 | FALSE |
| PAX5 | P18887 | XRCC1 | bait_int | 0 | 0.980448012 | 16\|16\|16 | 16 | 3 | 0\|0\|0\|0\|0\|0 | FALSE |
| PAX5 | O76031 | CLPX | bait_int | 0 | 0.978937082 | 28\|23\|13 | 21.33 | 3 | 0\|0\|0\|0\|0\|0 | FALSE |
| PAX5 | P49916 | LIG3 | bait_int | 0 | 0.987482846 | 36\|40\|43 | 39.67 | 3 | 0\|0\|0\|0\|0\|0 | FALSE |
| PAX5 | P35251 | RFC1 | bait_int | 0 | 0.981168822 | 17\|9\|6 | 10.67 | 3 | 0\|0\|0\|0\|0\|0 | FALSE |
| PAX5 | P35249 | RFC4 | bait_int | 0 | 0.978521229 | 10\|7\|8 | 8.33 | 3 | 0\|0\|0\|0\|0\|0 | FALSE |
| NRXN1 | P04083 | ANXA1 | bait_int | 0.05 | 0.979574722 | 13\|0\|12 | 8.33 | 3 | 0\|0\|0\|0\|0\|0 | FALSE |
| NRXN1 | Q6NZI2 | CAVIN1 | bait_int | 0.05 | 0.987371952 | 14\|0\|14 | 9.33 | 3 | 0\|0\|0\|0\|0\|0 | FALSE |
| NRXN1 | O60218 | AKR1B10 | bait_int | 0.05 | 0.985271898 | 12\|0\|11 | 7.67 | 3 | 0\|0\|0\|0\|0\|0 | FALSE |
| NRXN1 | Q15149 | PLEC | bait_int | 0 | 0.989465075 | 11\|25\|6 | 14 | 3 | 0\|0\|0\|0\|0\|0 | FALSE |
| NRXN1 | Q01813 | PFKP | bait_int | 0.05 | 0.985944193 | 15\|1\|13 | 9.67 | 3 | 0\|0\|0\|0\|0\|0 | FALSE |
| NRXN1 | P02545 | LMNA | bait_int | 0 | 0.972661177 | 11\|7\|9 | 9 | 3 | 0\|2\|0\|0\|0\|0 | FALSE |
| SIN3A | Q4LE39 | ARID4B | bait_int | 0 | 0.998696996 | 28\|28\|35 | 30.33 | 3 | 0\|0\|0\|0\|0\|0 | TRUE |
| SIN3A | O15294 | OGT | bait_int | 0 | 0.988214053 | 5\|5\|5 | 5 | 3 | 0\|0\|0\|0\|0\|0 | TRUE |
| SIN3A | Q9ULB1 | NRXN1 | bait_bait | 0 | 0.996915762 | 16\|13\|10 | 13 | 3 | 0\|0\|0\|0\|0\|0 | FALSE |
| SIN3A | P85037 | FOXK1 | bait_int | 0 | 0.989520522 | 11\|11\|13 | 11.67 | 3 | 0\|0\|0\|0\|0\|0 | FALSE |
| SIN3A | Q9H7L9 | SUDS3 | bait_int | 0 | 0.997449439 | 17\|20\|16 | 17.67 | 3 | 0\|0\|0\|0\|0\|0 | TRUE |
| SIN3A | Q9BUL5 | PHF23 | bait_int | 0 | 0.99757073 | 10\|9\|9 | 9.33 | 3 | 0\|0\|0\|0\|0\|0 | TRUE |
| SIN3A | P29374 | ARID4A | bait_int | 0 | 0.996690509 | 4\|8\|7 | 6.33 | 3 | 0\|0\|0\|0\|0\|0 | TRUE |
| SIN3A | O60381 | HBP1 | bait_int | 0 | 0.996829126 | 6\|4\|10 | 6.67 | 3 | 0\|0\|0\|0\|0\|0 | TRUE |
| SIN3A | Q8WY36 | BBX | bait_int | 0 | 0.999147503 | 45\|49\|55 | 49.67 | 3 | 0\|0\|0\|0\|0\|0 | FALSE |
| SIN3A | O15417 | TNRC18 | bait_int | 0 | 0.99467016 | 4\|5\|5 | 4.67 | 3 | 0\|0\|0\|0\|0\|0 | FALSE |
| SIN3A | Q9H160 | ING2 | bait_int | 0 | 0.995044427 | 4\|5\|4 | 4.33 | 3 | 0\|0\|0\|0\|0\|0 | TRUE |
| SIN3A | Q9NP50 | SINHCAF | bait_int | 0 | 0.998683134 | 29\|27\|22 | 26 | 3 | 0\|0\|0\|0\|0\|0 | FALSE |
| SIN3A | Q5PSV4 | BRMS1L | bait_int | 0 | 0.997324684 | 10\|11\|13 | 11.33 | 3 | 0\|0\|0\|0\|0\|0 | TRUE |
| SIN3A | Q9HAJ7 | SAP30L | bait_int | 0 | 0.997664296 | 11\|12\|12 | 11.67 | 3 | 0\|0\|0\|0\|0\|0 | TRUE |
| SIN3A | Q9HCU9 | BRMS1 | bait_int | 0 | 0.997206859 | 8\|9\|9 | 8.67 | 3 | 0\|0\|0\|0\|0\|0 | TRUE |
| SIN3A | Q9H0E3 | SAP130 | bait_int | 0 | 0.997123688 | 15\|15\|14 | 14.67 | 3 | 0\|0\|0\|0\|0\|0 | TRUE |
| SIN3A | O75446 | SAP30 | bait_int | 0 | 0.997102896 | 21\|22\|16 | 19.67 | 3 | 0\|0\|0\|0\|0\|0 | TRUE |
| SIN3A | Q13547 | HDAC1 | bait_int | 0 | 0.989160117 | 43\|42\|32 | 39 | 3 | 0\|1\|0\|0\|1\|0 | TRUE |
| SIN3A | Q16576 | RBBP7 | bait_int | 0 | 0.984467917 | 67\|58\|61 | 62 | 3 | 1\|3\|0\|0\|1\|1 | TRUE |
| SIN3A | Q09028 | RBBP4 | bait_int | 0 | 0.984107512 | 50\|46\|46 | 47.33 | 3 | 2\|4\|0\|1\|1\|2 | TRUE |
| SIN3A | Q92769 | HDAC2 | bait_int | 0 | 0.991190862 | 29\|33\|29 | 30.33 | 3 | 0\|0\|0\|0\|0\|0 | TRUE |
| KCNMA1 | Q9HDC5 | JPH1 | bait_int | 0 | 0.980101468 | 2\|3\|2 | 2.33 | 3 | 0\|0\|0\|0\|0\|0 | FALSE |
| KCNMA1 | Q9NVH0 | EXD2 | bait_int | 0 | 0.986214497 | 3\|2\|3 | 2.67 | 3 | 0\|0\|0\|0\|0\|0 | FALSE |
| KCNMA1 | Q9ULB1 | NRXN1 | bait_bait | 0 | 0.993665183 | 6\|3\|5 | 4.67 | 3 | 0\|0\|0\|0\|0\|0 | FALSE |
| KCNMA1 | Q9BVV7 | TIMM21 | bait_int | 0 | 0.971035888 | 2\|4\|4 | 3.33 | 3 | 0\|0\|0\|0\|0\|0 | FALSE |
| KCNMA1 | Q96A26 | FAM162A | bait_int | 0 | 0.989971029 | 8\|12\|11 | 10.33 | 3 | 0\|0\|0\|0\|0\|0 | FALSE |

| | | | | | | | | | | |
|---|---|---|---|---|---|---|---|---|---|---|
| KCNMA1 | P09132 | SRP19 | bait_int | 0 | 0.982908471 | 2\|3\|2 | 2.33 | 3 | 0\|0\|0\|0\|0\|0 | FALSE |
| KCNMA1 | Q9BWH2 | FUNDC2 | bait_int | 0 | 0.997941531 | 11\|10\|13 | 11.33 | 3 | 0\|0\|0\|0\|0\|0 | FALSE |
| KCNMA1 | Q9BSK0 | MARVELD1 | bait_int | 0 | 0.994559266 | 3\|3\|3 | 3 | 3 | 0\|0\|0\|0\|0\|0 | FALSE |
| KCNMA1 | Q96TC7 | RMDN3 | bait_int | 0 | 0.992629018 | 2\|4\|4 | 3.33 | 3 | 0\|0\|0\|0\|0\|0 | FALSE |
| KCNMA1 | Q8NBM4 | UBAC2 | bait_int | 0 | 0.984225336 | 4\|3\|4 | 3.67 | 3 | 0\|0\|0\|0\|0\|0 | FALSE |
| KCNMA1 | Q5HYI8 | RABL3 | bait_int | 0 | 0.97192304 | 2\|3\|3 | 2.67 | 3 | 0\|0\|0\|0\|0\|0 | FALSE |
| KCNMA1 | P78527 | PRKDC | bait_int | 0.01 | 0.972574542 | 43\|40\|29 | 37.33 | 3 | 8\|15\|15\|4\|4\|5 | FALSE |
| KCNMA1 | Q9P2E9 | RRBP1 | bait_int | 0 | 0.974140918 | 3\|3\|5 | 3.67 | 3 | 0\|0\|0\|0\|0\|0 | FALSE |
| KCNMA1 | P11279 | LAMP1 | bait_int | 0 | 0.972948809 | 4\|3\|5 | 4 | 3 | 0\|0\|0\|0\|0\|0 | FALSE |
| KCNMA1 | Q92973 | TNPO1 | bait_int | 0 | 0.976539 | 10\|10\|8 | 9.33 | 3 | 0\|0\|0\|0\|0\|0 | FALSE |
| SETD5 | O15294 | OGT | bait_int | 0 | 0.984370885 | 2\|3\|4 | 3 | 3 | 0\|0\|0\|0\|0\|0 | FALSE |
| SETD5 | Q9BQ70 | TCF25 | bait_int | 0 | 0.987316505 | 4\|8\|5 | 5.67 | 3 | 0\|0\|0\|0\|0\|0 | FALSE |
| SETD5 | Q96D09 | GPRASP2 | bait_int | 0.01 | 0.988737334 | 2\|2\|2 | 2 | 3 | 0\|0\|0\|0\|0\|0 | FALSE |
| SETD5 | Q13227 | GPS2 | bait_int | 0 | 0.988716541 | 6\|8\|12 | 8.67 | 3 | 0\|0\|0\|0\|0\|0 | FALSE |
| SETD5 | Q7KZI7 | MARK2 | bait_int | 0 | 0.992313664 | 3\|4\|8 | 5 | 3 | 0\|0\|0\|0\|0\|0 | FALSE |
| SETD5 | Q13643 | FHL3 | bait_int | 0 | 0.983088674 | 5\|6\|2 | 4.33 | 3 | 0\|0\|0\|0\|0\|0 | FALSE |
| SETD5 | Q6WCQ1 | MPRIP | bait_int | 0.04 | 0.978188547 | 54\|33\|0 | 29 | 3 | 0\|0\|0\|0\|1\|0 | FALSE |
| SETD5 | Q8WWI1 | LMO7 | bait_int | 0.04 | 0.973309214 | 42\|30\|0 | 24 | 3 | 0\|0\|0\|0\|0\|0 | FALSE |
| SETD5 | P35579 | MYH9 | bait_int | 0.04 | 0.987898698 | 193\|85\|11 | 96.33 | 3 | 17\|6\|3\|6\|11\|2 | FALSE |
| SETD5 | P21333 | FLNA | bait_int | 0.03 | 0.975339959 | 38\|34\|6 | 26 | 3 | 3\|1\|5\|1\|4\|2 | FALSE |
| SETD5 | P61962 | DCAF7 | bait_int | 0 | 0.989309131 | 8\|7\|5 | 6.67 | 3 | 0\|0\|0\|0\|0\|0 | FALSE |
| SETD5 | Q00610 | CLTC | bait_int | 0 | 0.972186413 | 18\|23\|8 | 16.33 | 3 | 0\|0\|0\|1\|3\|2 | FALSE |
| SETD5 | Q9Y4I1 | MYO5A | bait_int | 0.05 | 0.976705341 | 42\|11\|0 | 17.67 | 3 | 0\|0\|0\|0\|0\|0 | FALSE |
| SETD5 | Q9UQE7 | SMC3 | bait_int | 0 | 0.977398428 | 2\|3\|3 | 2.67 | 3 | 0\|0\|0\|0\|0\|0 | FALSE |
| SETD5 | Q9UPV0 | CEP164 | bait_int | 0 | 0.994822639 | 3\|6\|7 | 5.33 | 3 | 0\|0\|0\|0\|0\|0 | FALSE |
| SETD5 | Q9UNE7 | STUB1 | bait_int | 0 | 0.97208245 | 10\|11\|7 | 9.33 | 3 | 0\|0\|0\|0\|0\|0 | FALSE |
| SETD5 | Q9UM54 | MYO6 | bait_int | 0.04 | 0.976116217 | 63\|27\|0 | 30 | 3 | 1\|0\|0\|0\|0\|0 | FALSE |
| SETD5 | Q9P0K7 | RAI14 | bait_int | 0.04 | 0.981238131 | 56\|39\|0 | 31.67 | 3 | 0\|0\|0\|0\|0\|0 | FALSE |
| SETD5 | Q7Z406 | MYH14 | bait_int | 0.05 | 0.979235109 | 38\|9\|0 | 15.67 | 3 | 0\|0\|0\|0\|0\|0 | FALSE |
| SETD5 | Q96SB3 | PPP1R9B | bait_bait | 0.04 | 0.979013321 | 11\|12\|0 | 7.67 | 3 | 1\|0\|0\|0\|0\|0 | FALSE |
| SETD5 | Q6UB99 | ANKRD11 | bait_bait | 0 | 0.998232628 | 8\|21\|23 | 17.33 | 3 | 0\|0\|0\|0\|0\|0 | TRUE |
| SETD5 | Q6PD62 | CTR9 | bait_int | 0 | 0.995113736 | 27\|37\|33 | 32.33 | 3 | 0\|0\|0\|0\|0\|0 | FALSE |
| SETD5 | O75376 | NCOR1 | bait_int | 0 | 0.997865292 | 27\|73\|102 | 67.33 | 3 | 0\|0\|0\|0\|0\|0 | TRUE |
| SETD5 | O60907 | TBL1X | bait_int | 0 | 0.995980095 | 21\|36\|32 | 29.67 | 3 | 0\|0\|0\|0\|0\|0 | FALSE |
| SETD5 | O15379 | HDAC3 | bait_int | 0 | 0.991620576 | 8\|17\|18 | 14.33 | 3 | 0\|0\|0\|0\|0\|0 | FALSE |
| SETD5 | Q8N7H5 | PAF1 | bait_int | 0 | 0.992008705 | 12\|18\|19 | 16.33 | 3 | 0\|0\|0\|0\|0\|0 | FALSE |
| SETD5 | Q9Y618 | NCOR2 | bait_int | 0 | 0.99665932 | 19\|47\|73 | 46.33 | 3 | 0\|0\|0\|0\|0\|0 | FALSE |
| SETD5 | Q9Y2S7 | POLDIP2 | bait_int | 0 | 0.971022026 | 5\|8\|5 | 6 | 3 | 0\|0\|0\|0\|0\|0 | FALSE |
| SETD5 | Q8WVC0 | LEO1 | bait_int | 0 | 0.978791533 | 2\|3\|3 | 2.67 | 3 | 0\|0\|0\|0\|0\|0 | FALSE |
| SETD5 | Q9GZS3 | WDR61 | bait_int | 0 | 0.987392745 | 7\|14\|12 | 11 | 3 | 0\|0\|0\|0\|0\|0 | FALSE |
| SETD5 | Q9BZK7 | TBL1XR1 | bait_bait | 0 | 0.996867246 | 22\|35\|36 | 31 | 3 | 0\|0\|0\|0\|0\|0 | TRUE |
| SETD5 | Q6P1J9 | CDC73 | bait_int | 0 | 0.997518748 | 25\|32\|28 | 28.33 | 3 | 0\|0\|0\|0\|0\|0 | FALSE |
| SETD5 | Q93008 | USP9X | bait_int | 0 | 0.989236357 | 67\|57\|58 | 60.67 | 3 | 1\|0\|0\|0\|0\|0 | FALSE |
| SETD5 | Q8IZL8 | PELP1 | bait_int | 0 | 0.974501324 | 3\|6\|6 | 5 | 3 | 0\|0\|0\|0\|0\|0 | FALSE |
| SETD5 | Q69YQ0 | SPECC1L | bait_int | 0.04 | 0.978237064 | 60\|26\|0 | 28.67 | 3 | 0\|0\|0\|0\|0\|0 | FALSE |
| SETD5 | P14735 | IDE | bait_int | 0 | 0.997393992 | 41\|59\|48 | 49.33 | 3 | 0\|0\|0\|0\|0\|0 | FALSE |
| GABRB2 | Q9NYU2 | UGGT1 | bait_int | 0.01 | 0.982728268 | 9\|13\|8 | 10 | 3 | 1\|2\|6\|1\|0\|2 | FALSE |
| GABRB2 | Q7Z6Z7 | HUWE1 | bait_int | 0 | 0.980933173 | 10\|12\|12 | 11.33 | 3 | 0\|0\|0\|0\|0\|0 | FALSE |
| GABRB2 | Q9H078 | CLPB | bait_int | 0.03 | 0.975014208 | 5\|4\|5 | 4.67 | 3 | 1\|0\|0\|1\|4\|0 | FALSE |
| GABRB2 | Q14204 | DYNC1H1 | bait_bait | 0 | 0.994517681 | 8\|8\|8 | 8 | 3 | 4\|2\|2\|0\|1\|1 | FALSE |
| SHANK3 | Q8N567 | ZCCHC9 | bait_int | 0 | 0.984211475 | 5\|4\|2 | 3.67 | 3 | 0\|0\|0\|0\|0\|0 | FALSE |
| SHANK3 | P98175 | RBM10 | bait_int | 0 | 0.99721379 | 29\|30\|29 | 29.33 | 3 | 0\|0\|0\|0\|0\|0 | FALSE |
| SHANK3 | Q96DH6 | MSI2 | bait_int | 0 | 0.987094717 | 2\|4\|4 | 3.33 | 3 | 0\|0\|0\|0\|0\|0 | FALSE |
| SHANK3 | Q8TAF3 | WDR48 | bait_int | 0 | 0.981286647 | 2\|3\|4 | 3 | 3 | 0\|0\|0\|0\|1\|0 | FALSE |
| SHANK3 | Q9HBE1 | PATZ1 | bait_int | 0 | 0.992223562 | 3\|5\|6 | 4.67 | 3 | 0\|0\|0\|0\|0\|0 | FALSE |
| SHANK3 | Q99598 | TSNAX | bait_int | 0 | 0.981487642 | 3\|2\|2 | 2.33 | 3 | 0\|0\|0\|0\|0\|0 | FALSE |
| SHANK3 | Q9BVS4 | RIOK2 | bait_int | 0 | 0.987281851 | 4\|7\|5 | 5.33 | 3 | 0\|0\|0\|0\|0\|0 | FALSE |
| SHANK3 | P78332 | RBM6 | bait_int | 0 | 0.99761578 | 13\|13\|18 | 14.67 | 3 | 0\|0\|0\|0\|0\|0 | FALSE |
| SHANK3 | Q7L590 | MCM10 | bait_int | 0 | 0.989173979 | 7\|8\|6 | 7 | 3 | 0\|0\|0\|0\|0\|0 | FALSE |
| SHANK3 | Q9UII4 | HERC5 | bait_int | 0 | 0.987302643 | 4\|4\|3 | 3.67 | 3 | 0\|0\|0\|0\|0\|0 | FALSE |
| SHANK3 | Q9BXJ9 | NAA15 | bait_int | 0.02 | 0.995273146 | 8\|5\|8 | 7 | 3 | 3\|0\|0\|1\|4\|1 | FALSE |
| SHANK3 | Q9Y4L1 | HYOU1 | bait_int | 0 | 0.975436991 | 9\|7\|5 | 7 | 3 | 0\|0\|0\|0\|0\|0 | FALSE |
| SHANK3 | Q92900 | UPF1 | bait_int | 0 | 0.97900639 | 14\|15\|21 | 16.67 | 3 | 0\|0\|0\|0\|0\|0 | FALSE |
| SHANK3 | Q86VP6 | CAND1 | bait_int | 0 | 0.976078097 | 9\|4\|10 | 7.67 | 3 | 0\|0\|0\|0\|0\|0 | FALSE |
| SHANK3 | Q96S55 | WRNIP1 | bait_int | 0 | 0.974075075 | 2\|4\|6 | 4 | 3 | 0\|0\|0\|0\|0\|0 | FALSE |
| SHANK3 | Q9HCE1 | MOV10 | bait_int | 0 | 0.979540067 | 9\|12\|19 | 13.33 | 3 | 0\|0\|0\|0\|0\|0 | FALSE |
| SHANK3 | Q9H6S0 | YTHDC2 | bait_int | 0 | 0.97956086 | 7\|9\|8 | 8 | 3 | 0\|0\|0\|0\|0\|0 | FALSE |
| SHANK3 | Q9H5H4 | ZNF768 | bait_int | 0 | 0.975651848 | 4\|5\|5 | 4.67 | 3 | 0\|0\|0\|0\|0\|0 | FALSE |
| SHANK3 | Q9BRD0 | BUD13 | bait_int | 0 | 0.986415492 | 5\|3\|2 | 3.33 | 3 | 0\|0\|0\|0\|0\|0 | FALSE |
| SHANK3 | P30566 | ADSL | bait_int | 0 | 0.981605467 | 8\|6\|10 | 8 | 3 | 0\|0\|0\|0\|0\|0 | FALSE |
| SHANK3 | O43865 | AHCYL1 | bait_int | 0 | 0.974584494 | 5\|6\|8 | 6.33 | 3 | 0\|0\|0\|0\|0\|0 | FALSE |
| SHANK3 | Q9Y490 | TLN1 | bait_int | 0 | 0.976712272 | 9\|8\|12 | 9.67 | 3 | 0\|0\|0\|0\|0\|0 | FALSE |

| | | | | | | | | | | |
|---|---|---|---|---|---|---|---|---|---|---|
| SHANK3 | Q7L2E3 | DHX30 | bait_int | 0 | 0.98218766 | 50\|50\|49 | 49.67 | 3 | 0\|1\|0\|0\|0\|0 | FALSE |
| SHANK3 | P61201 | COPS2 | bait_int | 0 | 0.981175753 | 2\|2\|3 | 2.33 | 3 | 0\|0\|0\|0\|0\|0 | FALSE |
| SHANK3 | Q99733 | NAP1L4 | bait_int | 0.01 | 0.973264163 | 2\|2\|2 | 2 | 3 | 0\|0\|0\|0\|0\|0 | FALSE |
| SHANK3 | Q93008 | USP9X | bait_int | 0 | 0.972733952 | 3\|10\|19 | 10.67 | 3 | 1\|0\|0\|0\|0\|0 | FALSE |
| SHANK3 | O00566 | MPHOSPH1( | bait_int | 0 | 0.980032159 | 5\|3\|3 | 3.67 | 3 | 0\|0\|0\|0\|0\|0 | FALSE |
| SHANK3 | Q9Y2A7 | NCKAP1 | bait_int | 0 | 0.995910786 | 5\|6\|9 | 6.67 | 3 | 0\|0\|0\|0\|0\|0 | FALSE |
| SHANK3 | O43852 | CALU | bait_int | 0 | 0.978243994 | 10\|11\|14 | 11.67 | 3 | 0\|0\|1\|0\|1\|1 | FALSE |
| SHANK3 | Q13098 | GPS1 | bait_int | 0 | 0.977141986 | 3\|2\|2 | 2.33 | 3 | 0\|0\|0\|0\|0\|0 | FALSE |
| SHANK3 | Q01518 | CAP1 | bait_int | 0 | 0.972387408 | 7\|7\|8 | 7.33 | 3 | 0\|0\|0\|0\|0\|0 | FALSE |
| SHANK3 | Q14690 | PDCD11 | bait_int | 0 | 0.971812146 | 18\|18\|13 | 16.33 | 3 | 0\|0\|0\|0\|0\|0 | FALSE |
| SHANK3 | P14735 | IDE | bait_int | 0 | 0.989506661 | 4\|5\|10 | 6.33 | 3 | 0\|0\|0\|0\|0\|0 | FALSE |
| SHANK3 | P00387 | CYB5R3 | bait_int | 0 | 0.972921085 | 3\|3\|4 | 3.33 | 3 | 0\|0\|0\|0\|0\|0 | FALSE |
| VEZF1 | P82933 | MRPS9 | bait_int | 0 | 0.972837915 | 2\|3\|5 | 3.33 | 3 | 0\|0\|0\|0\|0\|0 | FALSE |
| VEZF1 | Q9H6S0 | YTHDC2 | bait_int | 0 | 0.979103422 | 10\|8\|5 | 7.67 | 3 | 0\|0\|0\|0\|0\|0 | FALSE |
| VEZF1 | Q8IWZ3 | ANKHD1 | bait_int | 0 | 0.993568151 | 31\|25\|30 | 28.67 | 3 | 0\|0\|0\|0\|0\|1 | FALSE |
| VEZF1 | O75179 | ANKRD17 | bait_int | 0 | 0.994975118 | 21\|16\|21 | 19.33 | 3 | 0\|0\|0\|0\|0\|0 | FALSE |
| VEZF1 | Q53H96 | PYCR3 | bait_int | 0 | 0.978632123 | 8\|11\|6 | 8.33 | 3 | 0\|0\|0\|0\|0\|0 | FALSE |
| VEZF1 | Q8N5P1 | ZC3H8 | bait_int | 0 | 0.975908291 | 5\|4\|5 | 4.67 | 3 | 1\|0\|0\|0\|1\|1 | FALSE |
| ADNP | O60341 | KDM1A | bait_int | 0 | 0.995196906 | 3\|20\|19 | 14 | 3 | 0\|0\|0\|0\|0\|0 | FALSE |
| ADNP | Q5VZL5 | ZMYM4 | bait_int | 0.04 | 0.996492979 | 1\|37\|39 | 25.67 | 3 | 0\|0\|0\|0\|0\|0 | FALSE |
| ADNP | Q14839 | CHD4 | bait_int | 0 | 0.998544517 | 78\|103\|11 | 97.33 | 3 | 0\|3\|1\|1\|1\|0 | FALSE |
| ADNP | Q14202 | ZMYM3 | bait_int | 0 | 0.999022747 | 16\|55\|52 | 41 | 3 | 0\|0\|0\|0\|0\|0 | FALSE |
| ADNP | P83916 | CBX1 | bait_int | 0 | 0.994684022 | 8\|15\|16 | 13 | 3 | 0\|1\|0\|0\|0\|0 | TRUE |
| ADNP | Q9UBW7 | ZMYM2 | bait_int | 0.04 | 0.988099694 | 0\|30\|37 | 22.33 | 3 | 0\|0\|0\|0\|0\|0 | TRUE |
| ADNP | O95785 | WIZ | bait_int | 0.05 | 0.976982576 | 0\|10\|14 | 8 | 3 | 0\|0\|0\|0\|0\|0 | FALSE |
| ADNP | Q9H582 | ZNF644 | bait_int | 0.05 | 0.983116397 | 0\|18\|14 | 10.67 | 3 | 0\|0\|0\|0\|0\|0 | FALSE |
| ADNP | Q15776 | ZKSCAN8 | bait_int | 0.05 | 0.984641189 | 0\|15\|11 | 8.67 | 3 | 0\|0\|0\|0\|0\|0 | FALSE |
| ADNP | Q14999 | CUL7 | bait_int | 0.05 | 0.976545931 | 0\|14\|11 | 8.33 | 3 | 0\|0\|0\|0\|0\|0 | FALSE |
| ANK2 | O75369 | FLNB | bait_int | 0 | 0.985521409 | 20\|60\|24 | 34.67 | 3 | 0\|0\|0\|0\|0\|0 | FALSE |
| ANK2 | O75083 | WDR1 | bait_int | 0 | 0.974393895 | 13\|25\|15 | 17.67 | 3 | 3\|0\|0\|0\|0\|0 | FALSE |
| ANK2 | O75643 | SNRNP200 | bait_int | 0.01 | 0.972172551 | 17\|16\|17 | 16.67 | 3 | 4\|8\|0\|1\|0\|0 | FALSE |
| ANK2 | O95425 | SVIL | bait_int | 0 | 0.978944012 | 4\|25\|8 | 12.33 | 3 | 0\|0\|0\|0\|0\|0 | FALSE |
| ANK2 | O95166 | GABARAP | bait_int | 0.02 | 0.975547885 | 3\|3\|3 | 3 | 3 | 2\|0\|0\|0\|1\|0 | FALSE |
| ANK2 | O94832 | MYO1D | bait_int | 0 | 0.975575609 | 44\|58\|34 | 45.33 | 3 | 2\|0\|0\|0\|0\|0 | FALSE |
| ANK2 | O00159 | MYO1C | bait_int | 0 | 0.977453875 | 49\|75\|45 | 56.33 | 3 | 3\|0\|0\|0\|0\|0 | FALSE |
| ANK2 | O43795 | MYO1B | bait_int | 0 | 0.974411222 | 59\|84\|53 | 65.33 | 3 | 7\|0\|0\|1\|8\|0 | FALSE |
| ANK2 | O43707 | ACTN4 | bait_int | 0 | 0.973094357 | 27\|53\|24 | 34.67 | 3 | 1\|0\|0\|0\|0\|0 | FALSE |
| ANK2 | P07237 | P4HB | bait_int | 0.02 | 0.971722044 | 3\|11\|8 | 7.33 | 3 | 2\|2\|1\|0\|1\|0 | FALSE |
| ANK2 | Q6WCQ1 | MPRIP | bait_int | 0 | 0.991669092 | 29\|77\|26 | 44 | 3 | 0\|0\|0\|0\|1\|0 | FALSE |
| ANK2 | Q14204 | DYNC1H1 | bait_bait | 0 | 0.997338545 | 12\|34\|20 | 22 | 3 | 4\|2\|2\|0\|1\|1 | FALSE |
| ANK2 | Q13813 | SPTAN1 | bait_int | 0 | 0.987011547 | 113\|199\|13 | 150.33 | 3 | 17\|0\|0\|4\|7\|0 | FALSE |
| ANK2 | Q16643 | DBN1 | bait_int | 0 | 0.972338892 | 69\|100\|82 | 83.67 | 3 | 27\|1\|0\|4\|15\|0 | FALSE |
| ANK2 | Q8WWI1 | LMO7 | bait_int | 0 | 0.989693794 | 33\|69\|23 | 41.67 | 3 | 0\|0\|0\|0\|0\|0 | FALSE |
| ANK2 | Q96N67 | DOCK7 | bait_int | 0 | 0.995682067 | 10\|43\|3 | 18.67 | 3 | 1\|0\|0\|0\|0\|1 | FALSE |
| ANK2 | P35221 | CTNNA1 | bait_int | 0 | 0.991946327 | 14\|18\|12 | 14.67 | 3 | 0\|0\|0\|0\|0\|0 | FALSE |
| ANK2 | P35579 | MYH9 | bait_int | 0 | 0.990809664 | 116\|252\|91 | 153 | 3 | 17\|6\|3\|6\|11\|2 | FALSE |
| ANK2 | P35580 | MYH10 | bait_int | 0 | 0.990393812 | 146\|234\|13 | 171 | 3 | 22\|0\|0\|6\|13\|1 | FALSE |
| ANK2 | P21333 | FLNA | bait_int | 0 | 0.975838982 | 17\|44\|21 | 27.33 | 3 | 3\|1\|5\|1\|4\|2 | FALSE |
| ANK2 | P63096 | GNAI1 | bait_bait | 0 | 0.984585742 | 5\|8\|8 | 7 | 3 | 0\|0\|0\|0\|0\|0 | FALSE |
| ANK2 | Q01082 | SPTBN1 | bait_int | 0 | 0.986741243 | 106\|179\|13 | 138.33 | 3 | 7\|0\|0\|0\|7\|0 | FALSE |
| ANK2 | P48634 | PRRC2A | bait_int | 0.01 | 0.985476359 | 2\|2\|4 | 2.67 | 3 | 0\|0\|0\|0\|0\|1 | FALSE |
| ANK2 | Q6P2Q9 | PRPF8 | bait_int | 0 | 0.971115593 | 10\|9\|11 | 10 | 3 | 3\|3\|1\|0\|0\|0 | FALSE |
| ANK2 | Q14160 | SCRIB | bait_int | 0 | 0.986602625 | 12\|16\|13 | 13.67 | 3 | 2\|2\|0\|0\|0\|0 | FALSE |
| ANK2 | Q14008 | CKAP5 | bait_int | 0 | 0.980205431 | 12\|16\|9 | 12.33 | 3 | 2\|0\|3\|0\|1\|1 | FALSE |
| ANK2 | Q00610 | CLTC | bait_int | 0 | 0.979193524 | 21\|41\|26 | 29.33 | 3 | 0\|0\|0\|1\|3\|2 | FALSE |
| ANK2 | Q9Y2D5 | AKAP2 | bait_int | 0 | 0.979664823 | 5\|23\|2 | 10 | 3 | 0\|0\|0\|0\|0\|0 | FALSE |
| ANK2 | Q9Y4I1 | MYO5A | bait_int | 0 | 0.991578991 | 13\|55\|5 | 24.33 | 3 | 0\|0\|0\|0\|0\|0 | FALSE |
| ANK2 | Q9Y5S2 | CDC42BPB | bait_int | 0 | 0.992133461 | 10\|38\|14 | 20.67 | 3 | 0\|0\|0\|0\|0\|0 | FALSE |
| ANK2 | Q9P0K7 | RAI14 | bait_int | 0 | 0.993685976 | 39\|84\|24 | 49 | 3 | 0\|0\|0\|0\|0\|0 | FALSE |
| ANK2 | Q9UM54 | MYO6 | bait_int | 0 | 0.990165093 | 32\|82\|22 | 45.33 | 3 | 1\|0\|0\|0\|0\|0 | FALSE |
| ANK2 | Q9ULV0 | MYO5B | bait_int | 0 | 0.989485868 | 2\|17\|5 | 8 | 3 | 0\|0\|0\|0\|0\|0 | FALSE |
| ANK2 | Q9ULJ8 | PPP1R9A | bait_int | 0 | 0.98473129 | 5\|4\|2 | 3.67 | 3 | 0\|0\|0\|0\|0\|0 | FALSE |
| ANK2 | Q92688 | ANP32B | bait_int | 0 | 0.981834186 | 2\|5\|4 | 3.67 | 3 | 0\|0\|0\|0\|0\|0 | FALSE |
| ANK2 | Q92614 | MYO18A | bait_int | 0 | 0.991090365 | 7\|35\|5 | 15.67 | 3 | 0\|0\|0\|0\|0\|0 | FALSE |
| ANK2 | Q86X29 | LSR | bait_int | 0 | 0.979942058 | 2\|8\|3 | 4.33 | 3 | 0\|0\|0\|0\|0\|0 | FALSE |
| ANK2 | Q7Z406 | MYH14 | bait_int | 0 | 0.993845386 | 16\|45\|9 | 23.33 | 3 | 0\|0\|0\|0\|0\|0 | FALSE |
| ANK2 | Q96SB3 | PPP1R9B | bait_bait | 0 | 0.994663229 | 10\|12\|8 | 10 | 3 | 1\|0\|0\|0\|0\|0 | FALSE |
| ANK2 | Q9H0H5 | RACGAP1 | bait_int | 0 | 0.984086719 | 2\|5\|3 | 3.33 | 3 | 0\|0\|0\|0\|0\|0 | FALSE |
| ANK2 | Q9NXR1 | NDE1 | bait_int | 0 | 0.991371065 | 6\|7\|9 | 7.33 | 3 | 0\|0\|0\|0\|0\|0 | FALSE |
| ANK2 | Q9NQX4 | MYO5C | bait_int | 0 | 0.991662162 | 11\|45\|9 | 21.67 | 3 | 0\|0\|0\|0\|0\|0 | FALSE |
| ANK2 | Q9BSC4 | NOL10 | bait_int | 0 | 0.980538113 | 3\|4\|3 | 3.33 | 3 | 0\|0\|0\|0\|0\|0 | FALSE |
| ANK2 | Q9BR76 | CORO1B | bait_int | 0 | 0.98250648 | 2\|6\|2 | 3.33 | 3 | 0\|0\|0\|0\|0\|0 | FALSE |

| | | | | | | | | | | |
|---|---|---|---|---|---|---|---|---|---|---|
| ANK2 | Q8IZL8 | PELP1 | bait_int | 0 | 0.975409268 | 7\|3\|6 | 5.33 | 3 | 0\|0\|0\|0\|0\|0 | FALSE |
| ANK2 | Q9Y5B9 | SUPT16H | bait_int | 0 | 0.974064679 | 40\|21\|36 | 32.33 | 3 | 0\|12\|0\|0\|0\|0 | FALSE |
| ANK2 | O43491 | EPB41L2 | bait_int | 0 | 0.972318099 | 7\|21\|9 | 12.33 | 3 | 0\|0\|0\|0\|0\|0 | FALSE |
| ANK2 | O15020 | SPTBN2 | bait_int | 0 | 0.990823526 | 32\|92\|51 | 58.33 | 3 | 0\|0\|0\|0\|0\|0 | FALSE |
| ANK2 | O14974 | PPP1R12A | bait_int | 0 | 0.987157095 | 18\|54\|15 | 29 | 3 | 0\|0\|0\|0\|0\|0 | FALSE |
| ANK2 | O14639 | ABLIM1 | bait_int | 0 | 0.984301576 | 2\|11\|2 | 5 | 3 | 0\|0\|0\|0\|0\|0 | FALSE |
| ANK2 | Q9UHB6 | LIMA1 | bait_int | 0 | 0.976483553 | 66\|90\|78 | 78 | 3 | 13\|0\|1\|2\|9\|0 | FALSE |
| ANK2 | Q7KZF4 | SND1 | bait_int | 0 | 0.983601558 | 16\|22\|17 | 18.33 | 3 | 0\|0\|0\|1\|0\|0 | FALSE |
| ANK2 | O75955 | FLOT1 | bait_int | 0 | 0.976940991 | 7\|9\|11 | 9 | 3 | 0\|0\|0\|0\|0\|0 | FALSE |
| ANK2 | Q5T9A4 | ATAD3B | bait_int | 0 | 0.973080495 | 20\|26\|25 | 23.67 | 3 | 1\|1\|0\|0\|0\|0 | FALSE |
| ANK2 | Q07157 | TJP1 | bait_int | 0 | 0.980967827 | 10\|18\|8 | 12 | 3 | 0\|0\|0\|0\|0\|0 | FALSE |
| ANK2 | Q13045 | FLII | bait_int | 0 | 0.98361542 | 4\|15\|6 | 8.33 | 3 | 0\|0\|0\|0\|0\|0 | FALSE |
| ANK2 | Q02218 | OGDH | bait_int | 0 | 0.979706408 | 8\|8\|6 | 7.33 | 3 | 0\|0\|0\|0\|0\|0 | FALSE |
| ANK2 | P60520 | GABARAPL2 | bait_int | 0 | 0.99753261 | 12\|12\|17 | 13.67 | 3 | 0\|0\|0\|0\|0\|0 | FALSE |
| ANK2 | P83111 | LACTB | bait_int | 0 | 0.975464715 | 8\|2\|4 | 4.67 | 3 | 0\|0\|0\|0\|0\|0 | FALSE |
| ANK2 | Q5M775 | SPECC1 | bait_int | 0 | 0.986380838 | 9\|27\|10 | 15.33 | 3 | 0\|0\|0\|0\|0\|0 | FALSE |
| ANK2 | Q5F1R6 | DNAJC21 | bait_int | 0.01 | 0.98727492 | 2\|2\|3 | 2.33 | 3 | 0\|0\|0\|0\|1\|0 | FALSE |
| ANK2 | Q5VT25 | CDC42BPA | bait_int | 0 | 0.986727381 | 38\|57\|39 | 44.67 | 3 | 0\|0\|0\|0\|0\|0 | FALSE |
| ANK2 | Q6P9B6 | MEAK7 | bait_int | 0 | 0.971410155 | 3\|5\|3 | 3.67 | 3 | 0\|0\|0\|0\|0\|0 | FALSE |
| ANK2 | Q69YQ0 | SPECC1L | bait_int | 0 | 0.992646345 | 39\|90\|25 | 51.33 | 3 | 0\|0\|0\|0\|0\|0 | FALSE |
| ANK2 | Q14690 | PDCD11 | bait_int | 0 | 0.981421799 | 36\|33\|43 | 37.33 | 3 | 0\|0\|0\|0\|0\|0 | FALSE |
| ANK2 | Q14573 | ITPR3 | bait_int | 0 | 0.988342274 | 7\|30\|10 | 15.67 | 3 | 0\|0\|0\|0\|0\|0 | FALSE |
| ANK2 | Q14571 | ITPR2 | bait_int | 0 | 0.984752083 | 6\|25\|5 | 12 | 3 | 0\|0\|0\|0\|0\|0 | FALSE |
| ANK2 | Q14254 | FLOT2 | bait_int | 0 | 0.976920198 | 9\|14\|12 | 11.67 | 3 | 1\|0\|0\|0\|0\|0 | FALSE |
| ANK2 | Q15691 | MAPRE1 | bait_int | 0 | 0.981875771 | 7\|9\|8 | 8 | 3 | 0\|0\|0\|0\|1\|0 | FALSE |
| ANK2 | Q14980 | NUMA1 | bait_int | 0 | 0.980794555 | 30\|32\|34 | 32 | 3 | 0\|0\|0\|0\|1\|0 | FALSE |
| ANK2 | P14735 | IDE | bait_int | 0 | 0.995418694 | 33\|16\|19 | 22.67 | 3 | 0\|0\|0\|0\|0\|0 | FALSE |
| ANK2 | P13861 | PRKAR2A | bait_int | 0 | 0.975194411 | 2\|8\|2 | 4 | 3 | 0\|0\|0\|0\|0\|0 | FALSE |
| ANK2 | P12814 | ACTN1 | bait_int | 0 | 0.977419221 | 10\|14\|7 | 10.33 | 3 | 0\|0\|0\|0\|0\|0 | FALSE |
| ANK2 | P25685 | DNAJB1 | bait_int | 0 | 0.976088493 | 11\|8\|6 | 8.33 | 3 | 0\|0\|0\|0\|0\|0 | FALSE |
| ANK2 | P04181 | OAT | bait_int | 0 | 0.99077501 | 2\|7\|6 | 5 | 3 | 0\|0\|0\|0\|0\|0 | FALSE |
| ANK2 | P00387 | CYB5R3 | bait_int | 0 | 0.971441344 | 2\|4\|3 | 3 | 3 | 0\|0\|0\|0\|0\|0 | FALSE |
| ANK2 | O94906 | PRPF6 | bait_int | 0 | 0.981515366 | 3\|3\|6 | 4 | 3 | 0\|0\|0\|0\|0\|0 | FALSE |
| ANK2 | P09493 | TPM1 | bait_int | 0 | 0.975125102 | 3\|13\|3 | 6.33 | 3 | 0\|0\|0\|0\|0\|0 | FALSE |
| ANK2 | P08133 | ANXA6 | bait_int | 0 | 0.979456897 | 3\|7\|5 | 5 | 3 | 0\|0\|0\|0\|0\|0 | FALSE |
| ANK2 | P06737 | PYGL | bait_int | 0 | 0.976951387 | 4\|13\|9 | 8.67 | 3 | 0\|0\|0\|0\|0\|0 | FALSE |
| ANK2 | P49773 | HINT1 | bait_int | 0 | 0.983421355 | 2\|6\|7 | 5 | 3 | 0\|0\|0\|0\|0\|0 | FALSE |
| ANK2 | P49591 | SARS1 | bait_int | 0 | 0.973031979 | 4\|6\|6 | 5.33 | 3 | 0\|0\|0\|0\|0\|0 | FALSE |
| ANK2 | P46940 | IQGAP1 | bait_int | 0 | 0.9828253 | 3\|20\|2 | 8.33 | 3 | 0\|0\|0\|0\|0\|0 | FALSE |
| ANK2 | P46013 | MKI67 | bait_int | 0 | 0.983764434 | 36\|13\|29 | 26 | 3 | 0\|2\|0\|0\|0\|0 | FALSE |
| ANK2 | P55196 | AFDN | bait_int | 0 | 0.986678865 | 2\|10\|6 | 6 | 3 | 0\|0\|0\|0\|0\|0 | FALSE |
| ANK2 | P53355 | DAPK1 | bait_int | 0 | 0.981432195 | 4\|16\|5 | 8.33 | 3 | 0\|0\|0\|0\|0\|0 | FALSE |
| ANK2 | P52209 | PGD | bait_int | 0 | 0.971285399 | 2\|5\|6 | 4.33 | 3 | 0\|0\|0\|0\|0\|0 | FALSE |
| ANK2 | P51659 | HSD17B4 | bait_int | 0 | 0.987614533 | 2\|11\|6 | 6.33 | 3 | 0\|0\|0\|0\|0\|0 | FALSE |
| ANK2 | P42285 | MTREX | bait_int | 0 | 0.974820144 | 7\|4\|5 | 5.33 | 3 | 0\|0\|0\|0\|0\|0 | FALSE |
| ANK2 | P31939 | ATIC | bait_int | 0 | 0.974813213 | 2\|10\|6 | 6 | 3 | 0\|0\|0\|0\|0\|0 | FALSE |
| ANK2 | P28290 | ITPRID2 | bait_int | 0 | 0.983629282 | 10\|29\|19 | 19.33 | 3 | 0\|0\|0\|0\|0\|0 | FALSE |
| NRXN1 | Q9P2S2 | NRXN2 | bait_int | 0.01 | 0.991606715 | 5\|4\|7 | 5.33 | 3 | 2\|0\|0\|0\|0\|0 | FALSE |
| NRXN1 | Q5VUJ6 | LRCH2 | bait_int | 0 | 0.984904562 | 6\|4\|3 | 4.33 | 3 | 0\|0\|0\|0\|0\|0 | FALSE |
| NRXN1 | Q92616 | GCN1 | bait_int | 0.05 | 0.974286467 | 9\|0\|10 | 6.33 | 3 | 1\|0\|0\|0\|0\|0 | FALSE |
| NRXN1 | O60292 | SIPA1L3 | bait_int | 0 | 0.98417682 | 13\|3\|4 | 6.67 | 3 | 0\|0\|0\|0\|0\|0 | FALSE |
| NRXN1 | Q8N3R9 | PALS1 | bait_int | 0 | 0.98313719 | 12\|2\|2 | 5.33 | 3 | 0\|0\|0\|0\|0\|0 | FALSE |
| NRXN1 | O75369 | FLNB | bait_int | 0 | 0.99069877 | 102\|42\|75 | 73 | 3 | 0\|0\|0\|0\|0\|0 | FALSE |
| NRXN1 | O75083 | WDR1 | bait_int | 0 | 0.977848935 | 26\|22\|23 | 23.67 | 3 | 3\|0\|0\|0\|0\|0 | FALSE |
| NRXN1 | O95425 | SVIL | bait_int | 0 | 0.98258272 | 21\|5\|28 | 18 | 3 | 0\|0\|0\|0\|0\|0 | FALSE |
| NRXN1 | O94832 | MYO1D | bait_int | 0 | 0.976726134 | 63\|24\|66 | 51 | 3 | 2\|0\|0\|0\|0\|0 | FALSE |
| NRXN1 | O43795 | MYO1B | bait_int | 0 | 0.976109286 | 85\|49\|92 | 75.33 | 3 | 7\|0\|0\|1\|8\|0 | FALSE |
| NRXN1 | O00159 | MYO1C | bait_int | 0 | 0.977627147 | 67\|33\|71 | 57 | 3 | 3\|0\|0\|0\|0\|0 | FALSE |
| NRXN1 | O43707 | ACTN4 | bait_int | 0 | 0.977363774 | 78\|24\|47 | 49.67 | 3 | 1\|0\|0\|0\|0\|0 | FALSE |
| NRXN1 | Q6WCQ1 | MPRIP | bait_int | 0 | 0.991897811 | 56\|16\|62 | 44.67 | 3 | 0\|0\|0\|0\|1\|0 | FALSE |
| NRXN1 | Q14204 | DYNC1H1 | bait_bait | 0 | 0.997359338 | 36\|9\|22 | 22.33 | 3 | 4\|2\|2\|0\|1\|1 | FALSE |
| NRXN1 | Q14126 | DSG2 | bait_int | 0 | 0.972727021 | 66\|50\|58 | 58 | 3 | 7\|0\|0\|0\|2\|0 | FALSE |
| NRXN1 | Q13813 | SPTAN1 | bait_int | 0 | 0.987053132 | 210\|76\|168 | 151.33 | 3 | 17\|0\|0\|4\|7\|0 | FALSE |
| NRXN1 | Q16643 | DBN1 | bait_int | 0 | 0.973316145 | 112\|46\|113 | 90.33 | 3 | 27\|1\|0\|4\|15\|0 | FALSE |
| NRXN1 | Q15149 | PLEC | bait_int | 0 | 0.993464188 | 57\|11\|22 | 30 | 3 | 0\|0\|0\|0\|0\|0 | FALSE |
| NRXN1 | Q9HAV4 | XPO5 | bait_int | 0 | 0.980337118 | 7\|2\|12 | 7 | 3 | 0\|0\|1\|0\|0\|0 | FALSE |
| NRXN1 | Q8WWI1 | LMO7 | bait_int | 0 | 0.988286827 | 53\|7\|42 | 34 | 3 | 0\|0\|0\|0\|0\|0 | FALSE |
| NRXN1 | Q96N67 | DOCK7 | bait_int | 0 | 0.996222675 | 39\|6\|24 | 23 | 3 | 1\|0\|0\|0\|0\|1 | FALSE |
| NRXN1 | P35221 | CTNNA1 | bait_int | 0 | 0.991177001 | 20\|6\|13 | 13 | 3 | 0\|0\|0\|0\|0\|0 | FALSE |
| NRXN1 | P35579 | MYH9 | bait_int | 0 | 0.989534384 | 154\|53\|15 | 121 | 3 | 17\|6\|3\|6\|11\|2 | FALSE |
| NRXN1 | P35580 | MYH10 | bait_int | 0 | 0.989936375 | 199\|71\|20 | 157.33 | 3 | 22\|0\|0\|6\|13\|1 | FALSE |
| NRXN1 | P21333 | FLNA | bait_int | 0 | 0.983386701 | 100\|27\|43 | 56.67 | 3 | 3\|1\|5\|1\|4\|2 | FALSE |

| | | | | | | | | | |
|---|---|---|---|---|---|---|---|---|---|
| NRXN1 | P63096 | GNAI1 | bait_bait | 0 | 0.985611511 | 9\|4\|11 | 8 | 3 | 0\|0\|0\|0\|0\|0 | FALSE |
| NRXN1 | Q01082 | SPTBN1 | bait_int | 0 | 0.987066994 | 186\|78\|17[illegible] | 145 | 3 | 7\|0\|0\|0\|7\|0 | FALSE |
| NRXN1 | Q14160 | SCRIB | bait_int | 0.01 | 0.988536339 | 26\|4\|23 | 17.67 | 3 | 2\|2\|0\|0\|0\|0 | FALSE |
| NRXN1 | Q00610 | CLTC | bait_int | 0 | 0.979872749 | 45\|16\|32 | 31 | 3 | 0\|0\|0\|1\|3\|2 | FALSE |
| NRXN1 | Q9Y2D5 | AKAP2 | bait_int | 0 | 0.985216451 | 31\|5\|20 | 18.67 | 3 | 0\|0\|0\|0\|0\|0 | FALSE |
| NRXN1 | Q9Y5S2 | CDC42BPB | bait_int | 0 | 0.995162252 | 52\|14\|59 | 41.67 | 3 | 0\|0\|0\|0\|0\|0 | FALSE |
| NRXN1 | Q9Y4I1 | MYO5A | bait_int | 0 | 0.993588944 | 55\|9\|45 | 36.33 | 3 | 0\|0\|0\|0\|0\|0 | FALSE |
| NRXN1 | Q9P0K7 | RAI14 | bait_int | 0 | 0.993914695 | 68\|20\|65 | 51 | 3 | 0\|0\|0\|0\|0\|0 | FALSE |
| NRXN1 | Q9NZR1 | TMOD2 | bait_int | 0.05 | 0.972782468 | 12\|1\|10 | 7.67 | 3 | 0\|0\|0\|0\|0\|0 | FALSE |
| NRXN1 | Q9UM54 | MYO6 | bait_int | 0 | 0.991925535 | 78\|35\|70 | 61 | 3 | 1\|0\|0\|0\|0\|0 | FALSE |
| NRXN1 | Q9ULV0 | MYO5B | bait_int | 0 | 0.992244355 | 18\|2\|20 | 13.33 | 3 | 0\|0\|0\|0\|0\|0 | FALSE |
| NRXN1 | Q9UJC5 | SH3BGRL2 | bait_int | 0 | 0.976234042 | 2\|5\|2 | 3 | 3 | 0\|0\|0\|0\|0\|0 | FALSE |
| NRXN1 | Q92614 | MYO18A | bait_int | 0 | 0.995363247 | 62\|18\|40 | 40 | 3 | 0\|0\|0\|0\|0\|0 | FALSE |
| NRXN1 | Q7Z406 | MYH14 | bait_int | 0 | 0.994219653 | 39\|6\|31 | 25.33 | 3 | 0\|0\|0\|0\|0\|0 | FALSE |
| NRXN1 | Q96PY5 | FMNL2 | bait_int | 0 | 0.988252173 | 20\|3\|19 | 14 | 3 | 0\|0\|0\|0\|0\|0 | FALSE |
| NRXN1 | Q9NQX4 | MYO5C | bait_int | 0 | 0.992320594 | 25\|9\|38 | 24 | 3 | 0\|0\|0\|0\|0\|0 | FALSE |
| NRXN1 | Q96SB3 | PPP1R9B | bait_bait | 0 | 0.995896924 | 22\|4\|20 | 15.33 | 3 | 1\|0\|0\|0\|0\|0 | FALSE |
| NRXN1 | Q9BY89 | KIAA1671 | bait_int | 0 | 0.99284041 | 41\|4\|27 | 24 | 3 | 0\|0\|0\|0\|0\|0 | FALSE |
| NRXN1 | Q9BY67 | CADM1 | bait_int | 0 | 0.97518748 | 5\|2\|7 | 4.67 | 3 | 0\|0\|0\|0\|0\|0 | FALSE |
| NRXN1 | Q7KZF4 | SND1 | bait_int | 0 | 0.972636919 | 12\|4\|4 | 6.67 | 3 | 0\|0\|0\|1\|0\|0 | FALSE |
| NRXN1 | Q92973 | TNPO1 | bait_int | 0 | 0.979553929 | 16\|3\|17 | 12 | 3 | 0\|0\|0\|0\|0\|0 | FALSE |
| NRXN1 | Q16891 | IMMT | bait_int | 0.01 | 0.973863684 | 20\|4\|26 | 16.67 | 3 | 2\|2\|0\|0\|0\|0 | FALSE |
| NRXN1 | O43491 | EPB41L2 | bait_int | 0 | 0.977585562 | 31\|4\|22 | 19 | 3 | 0\|0\|0\|0\|0\|0 | FALSE |
| NRXN1 | O15020 | SPTBN2 | bait_int | 0 | 0.991246309 | 89\|26\|76 | 63.67 | 3 | 0\|0\|0\|0\|0\|0 | FALSE |
| NRXN1 | O14974 | PPP1R12A | bait_int | 0 | 0.987330367 | 47\|8\|34 | 29.67 | 3 | 0\|0\|0\|0\|0\|0 | FALSE |
| NRXN1 | A1L390 | PLEKHG3 | bait_int | 0 | 0.975596401 | 11\|4\|12 | 9 | 3 | 0\|0\|0\|0\|0\|0 | FALSE |
| NRXN1 | Q9UHB6 | LIMA1 | bait_int | 0 | 0.97606077 | 93\|37\|94 | 74.67 | 3 | 13\|0\|1\|2\|9\|0 | FALSE |
| NRXN1 | Q0ZGT2 | NEXN | bait_int | 0 | 0.973877545 | 25\|11\|27 | 21 | 3 | 0\|0\|0\|0\|0\|0 | FALSE |
| NRXN1 | Q07157 | TJP1 | bait_int | 0 | 0.985625373 | 34\|7\|22 | 21 | 3 | 0\|0\|0\|0\|0\|0 | FALSE |
| NRXN1 | Q03252 | LMNB2 | bait_int | 0.01 | 0.971347777 | 19\|7\|29 | 18.33 | 3 | 0\|5\|0\|0\|0\|0 | FALSE |
| NRXN1 | Q12792 | TWF1 | bait_int | 0 | 0.980746039 | 9\|5\|9 | 7.67 | 3 | 0\|0\|0\|0\|0\|0 | FALSE |
| NRXN1 | Q13045 | FLII | bait_int | 0.05 | 0.986831344 | 21\|1\|16 | 12.67 | 3 | 0\|0\|0\|0\|0\|0 | FALSE |
| NRXN1 | Q12965 | MYO1E | bait_int | 0 | 0.988577924 | 14\|2\|6 | 7.33 | 3 | 0\|0\|0\|0\|0\|0 | FALSE |
| NRXN1 | Q12959 | DLG1 | bait_int | 0 | 0.987489777 | 3\|2\|3 | 2.67 | 3 | 0\|0\|0\|0\|0\|0 | FALSE |
| NRXN1 | P84095 | RHOG | bait_int | 0 | 0.972352754 | 3\|2\|6 | 3.67 | 3 | 0\|0\|0\|0\|0\|0 | FALSE |
| NRXN1 | Q5M775 | SPECC1 | bait_int | 0 | 0.98846703 | 26\|2\|32 | 20 | 3 | 0\|0\|0\|0\|0\|0 | FALSE |
| NRXN1 | Q6P9B6 | MEAK7 | bait_int | 0 | 0.976185526 | 7\|3\|6 | 5.33 | 3 | 0\|0\|0\|0\|0\|0 | FALSE |
| NRXN1 | Q69YQ0 | SPECC1L | bait_int | 0 | 0.991142346 | 59\|10\|47 | 38.67 | 3 | 0\|0\|0\|0\|0\|0 | FALSE |
| NRXN1 | Q5VT25 | CDC42BPA | bait_int | 0 | 0.989776965 | 81\|36\|89 | 68.67 | 3 | 0\|0\|0\|0\|0\|0 | FALSE |
| NRXN1 | Q14571 | ITPR2 | bait_int | 0 | 0.987905629 | 23\|4\|27 | 18 | 3 | 0\|0\|0\|0\|0\|0 | FALSE |
| NRXN1 | Q14254 | FLOT2 | bait_int | 0 | 0.979595514 | 15\|9\|20 | 14.67 | 3 | 1\|0\|0\|0\|0\|0 | FALSE |
| NRXN1 | Q14573 | ITPR3 | bait_int | 0 | 0.991280964 | 23\|12\|42 | 25.67 | 3 | 0\|0\|0\|0\|0\|0 | FALSE |
| NRXN1 | Q15691 | MAPRE1 | bait_int | 0 | 0.975312236 | 7\|2\|4 | 4.33 | 3 | 0\|0\|0\|0\|1\|0 | FALSE |
| NRXN1 | P55196 | AFDN | bait_int | 0 | 0.991294825 | 19\|7\|11 | 12.33 | 3 | 0\|0\|0\|0\|0\|0 | FALSE |
| NRXN1 | P13861 | PRKAR2A | bait_int | 0 | 0.979838095 | 10\|2\|6 | 6 | 3 | 0\|0\|0\|0\|0\|0 | FALSE |
| NRXN1 | P12814 | ACTN1 | bait_int | 0 | 0.985569926 | 40\|10\|24 | 24.67 | 3 | 0\|0\|0\|0\|0\|0 | FALSE |
| NRXN1 | P19022 | CDH2 | bait_int | 0 | 0.971146782 | 10\|7\|9 | 8.67 | 3 | 0\|0\|0\|0\|0\|0 | FALSE |
| NRXN1 | P02545 | LMNA | bait_int | 0 | 0.975350356 | 14\|4\|15 | 11 | 3 | 0\|2\|0\|0\|0\|0 | FALSE |
| NRXN1 | P04181 | OAT | bait_int | 0 | 0.986443215 | 4\|2\|2 | 2.67 | 3 | 0\|0\|0\|0\|0\|0 | FALSE |
| NRXN1 | P09493 | TPM1 | bait_int | 0 | 0.973704274 | 8\|3\|6 | 5.67 | 3 | 0\|0\|0\|0\|0\|0 | FALSE |
| NRXN1 | P07197 | NEFM | bait_int | 0 | 0.973458228 | 11\|4\|9 | 8 | 3 | 0\|0\|0\|0\|0\|0 | FALSE |
| NRXN1 | P06737 | PYGL | bait_int | 0 | 0.974529047 | 13\|3\|5 | 7 | 3 | 0\|0\|0\|0\|0\|0 | FALSE |
| NRXN1 | P06396 | GSN | bait_int | 0 | 0.972768606 | 9\|5\|4 | 6 | 3 | 0\|0\|0\|0\|0\|0 | FALSE |
| NRXN1 | P06241 | FYN | bait_int | 0 | 0.97145174 | 15\|3\|18 | 12 | 3 | 0\|0\|0\|0\|0\|0 | FALSE |
| NRXN1 | P49773 | HINT1 | bait_int | 0 | 0.978431128 | 5\|2\|2 | 3 | 3 | 0\|0\|0\|0\|0\|0 | FALSE |
| NRXN1 | P46940 | IQGAP1 | bait_int | 0 | 0.98910467 | 36\|10\|9 | 18.33 | 3 | 0\|0\|0\|0\|0\|0 | FALSE |
| NRXN1 | P54289 | CACNA2D1 | bait_int | 0 | 0.973219113 | 15\|2\|21 | 12.67 | 3 | 0\|0\|0\|0\|0\|0 | FALSE |
| NRXN1 | P53355 | DAPK1 | bait_int | 0 | 0.984204544 | 13\|3\|18 | 11.33 | 3 | 0\|0\|0\|0\|0\|0 | FALSE |
| NRXN1 | O75955 | FLOT1 | bait_int | 0 | 0.982645098 | 16\|8\|23 | 15.67 | 3 | 0\|0\|0\|0\|0\|0 | FALSE |
| NRXN1 | P31939 | ATIC | bait_int | 0 | 0.972477509 | 9\|2\|4 | 5 | 3 | 0\|0\|0\|0\|0\|0 | FALSE |
| NRXN1 | P28290 | ITPRID2 | bait_int | 0 | 0.986574902 | 33\|16\|36 | 28.33 | 3 | 0\|0\|0\|0\|0\|0 | FALSE |
| NRXN1 | P27797 | CALR | bait_int | 0 | 0.979851957 | 28\|19\|31 | 26 | 3 | 0\|0\|0\|0\|0\|0 | FALSE |
| SLC6A1 | Q16891 | IMMT | bait_int | 0 | 0.971555703 | 19\|11\|12 | 14 | 3 | 0\|0\|0\|1\|0\|0 | FALSE |
| SLC6A1 | P35579 | MYH9 | bait_int | 0.04 | 0.975374614 | 3\|29\|42 | 24.67 | 3 | 1\|6\|0\|4\|3\|11 | FALSE |
| SLC6A1 | O95197 | RTN3 | bait_int | 0 | 0.972456717 | 4\|5\|7 | 5.33 | 3 | 0\|0\|0\|0\|0\|0 | FALSE |
| SLC6A1 | Q5HYI8 | RABL3 | bait_int | 0 | 0.979512344 | 6\|5\|4 | 5 | 3 | 0\|0\|0\|0\|0\|0 | FALSE |
| SLC6A1 | Q9H0P0 | NT5C3A | bait_int | 0 | 0.989756172 | 2\|3\|2 | 2.33 | 3 | 0\|0\|0\|0\|0\|0 | FALSE |
| SLC6A1 | Q9Y639 | NPTN | bait_int | 0 | 0.995737514 | 7\|8\|5 | 6.67 | 3 | 0\|0\|0\|0\|0\|0 | FALSE |
| SLC6A1 | Q9Y679 | AUP1 | bait_int | 0 | 0.981695568 | 6\|7\|10 | 7.67 | 3 | 0\|0\|0\|0\|0\|0 | FALSE |
| SLC6A1 | Q15043 | SLC39A14 | bait_int | 0.01 | 0.972983463 | 2\|2\|2 | 2 | 3 | 0\|0\|0\|0\|0\|0 | FALSE |
| SLC6A1 | Q96G23 | CERS2 | bait_int | 0 | 0.977668732 | 5\|3\|4 | 4 | 3 | 0\|0\|0\|0\|0\|0 | FALSE |
| POGZ | Q9NQV6 | PRDM10 | bait_int | 0.04 | 0.989638347 | 1\|5\|3 | 3 | 3 | 0\|0\|0\|0\|0\|0 | FALSE |

| | | | | | | | | | | |
|---|---|---|---|---|---|---|---|---|---|---|
| POGZ | Q96JM3 | CHAMP1 | bait_int | 0 | 0.998537586 | 32\|37\|46 | 38.33 | 3 | 0\|0\|0\|0\|0\|0 | TRUE |
| POGZ | Q96B54 | ZNF428 | bait_int | 0 | 0.99375875 | 3\|5\|2 | 3.33 | 3 | 0\|0\|0\|0\|0\|0 | TRUE |
| POGZ | Q8IX15 | HOMEZ | bait_int | 0 | 0.998149457 | 14\|14\|12 | 13.33 | 3 | 0\|0\|0\|0\|0\|0 | FALSE |
| POGZ | P83916 | CBX1 | bait_int | 0 | 0.993076059 | 8\|10\|9 | 9 | 3 | 0\|0\|0\|0\|0\|0 | TRUE |
| POGZ | Q7Z4V5 | HDGFL2 | bait_int | 0 | 0.992272078 | 18\|22\|15 | 18.33 | 3 | 2\|0\|0\|2\|1\|0 | FALSE |
| POGZ | O95785 | WIZ | bait_int | 0 | 0.994843432 | 8\|18\|17 | 14.33 | 3 | 0\|0\|0\|0\|0\|0 | FALSE |
| POGZ | Q96KQ7 | EHMT2 | bait_int | 0 | 0.997442508 | 10\|31\|14 | 18.33 | 3 | 0\|0\|0\|0\|0\|0 | FALSE |
| POGZ | Q9H9B1 | EHMT1 | bait_int | 0 | 0.99689497 | 12\|19\|16 | 15.67 | 3 | 0\|0\|0\|0\|0\|0 | FALSE |
| POGZ | Q9H582 | ZNF644 | bait_int | 0 | 0.9973732 | 16\|34\|19 | 23 | 3 | 0\|0\|0\|0\|0\|0 | FALSE |
| POGZ | Q86T24 | ZBTB33 | bait_int | 0.04 | 0.978365285 | 0\|6\|4 | 3.33 | 3 | 0\|0\|0\|0\|0\|0 | FALSE |
| CHD2 | O14646 | CHD1 | bait_int | 0 | 0.989783895 | 4\|4\|4 | 4 | 3 | 0\|0\|0\|0\|0\|0 | FALSE |
| LRRC4C | Q8TCG1 | CIP2A | bait_int | 0 | 0.989527453 | 8\|10\|7 | 8.33 | 3 | 0\|0\|0\|0\|0\|0 | FALSE |
| LRRC4C | Q14574 | DSC3 | bait_int | 0 | 0.985258036 | 2\|4\|2 | 2.67 | 3 | 0\|0\|0\|0\|0\|0 | FALSE |
| LRRC4C | Q8IWV7 | UBR1 | bait_bait | 0 | 0.993124575 | 3\|2\|4 | 3 | 3 | 0\|0\|0\|0\|0\|0 | FALSE |
| LRRC4C | Q92616 | GCN1 | bait_int | 0.05 | 0.986800155 | 7\|2\|1 | 3.33 | 3 | 0\|0\|0\|0\|0\|0 | FALSE |
| LRRC4C | Q32P28 | P3H1 | bait_int | 0 | 0.979075699 | 5\|5\|6 | 5.33 | 3 | 0\|0\|0\|0\|0\|0 | FALSE |
| LRRC4C | Q7Z3U7 | MON2 | bait_int | 0 | 0.990400743 | 3\|4\|4 | 3.67 | 3 | 0\|0\|0\|0\|0\|0 | FALSE |
| LRRC4C | Q9C0E2 | XPO4 | bait_int | 0 | 0.987060063 | 4\|4\|6 | 4.67 | 3 | 0\|0\|0\|0\|0\|0 | FALSE |
| LRRC4C | O14787 | TNPO2 | bait_int | 0 | 0.981820324 | 2\|4\|3 | 3 | 3 | 0\|0\|0\|0\|0\|0 | FALSE |
| LRRC4C | Q9Y5L0 | TNPO3 | bait_int | 0 | 0.983047088 | 5\|6\|4 | 5 | 3 | 0\|0\|0\|0\|0\|0 | FALSE |
| LRRC4C | Q8N1S5 | SLC39A11 | bait_int | 0 | 0.994559266 | 3\|3\|3 | 3 | 3 | 0\|0\|0\|0\|0\|0 | TRUE |
| LRRC4C | O95260 | ATE1 | bait_int | 0 | 0.990996798 | 9\|3\|3 | 5 | 3 | 0\|0\|0\|0\|0\|0 | FALSE |
| LRRC4C | Q9H078 | CLPB | bait_int | 0.02 | 0.971867593 | 2\|5\|4 | 3.67 | 3 | 1\|1\|0\|1\|1\|0 | FALSE |
| LRRC4C | Q9HAV4 | XPO5 | bait_int | 0 | 0.983213429 | 7\|9\|13 | 9.67 | 3 | 0\|1\|0\|0\|0\|0 | FALSE |
| LRRC4C | P07237 | P4HB | bait_int | 0 | 0.975991461 | 9\|11\|11 | 10.33 | 3 | 0\|0\|0\|0\|0\|0 | FALSE |
| LRRC4C | Q07065 | CKAP4 | bait_int | 0 | 0.984308507 | 18\|17\|26 | 20.33 | 3 | 0\|0\|0\|0\|0\|0 | FALSE |
| LRRC4C | P78527 | PRKDC | bait_int | 0 | 0.979228178 | 76\|57\|62 | 65 | 3 | 7\|1\|6\|2\|0\|1 | FALSE |
| LRRC4C | P06280 | GLA | bait_int | 0 | 0.979657892 | 3\|2\|4 | 3 | 3 | 0\|0\|0\|0\|0\|0 | FALSE |
| LRRC4C | Q9UBV2 | SEL1L | bait_int | 0 | 0.986762035 | 8\|7\|8 | 7.67 | 3 | 0\|0\|0\|0\|0\|0 | FALSE |
| LRRC4C | Q96DZ1 | ERLEC1 | bait_int | 0 | 0.980940103 | 5\|5\|4 | 4.67 | 3 | 0\|0\|0\|0\|0\|0 | FALSE |
| LRRC4C | Q92973 | TNPO1 | bait_int | 0 | 0.986921446 | 25\|28\|32 | 28.33 | 3 | 0\|0\|0\|0\|0\|0 | FALSE |
| LRRC4C | Q9BS26 | ERP44 | bait_int | 0 | 0.994198861 | 11\|14\|12 | 12.33 | 3 | 0\|0\|0\|0\|0\|0 | FALSE |
| LRRC4C | Q96P70 | IPO9 | bait_int | 0.03 | 0.972844846 | 1\|9\|7 | 5.67 | 3 | 0\|0\|0\|0\|0\|0 | FALSE |
| LRRC4C | Q9NYU2 | UGGT1 | bait_int | 0 | 0.989548246 | 18\|28\|26 | 24 | 3 | 0\|0\|0\|1\|0\|0 | FALSE |
| LRRC4C | O60613 | SELENOF | bait_int | 0 | 0.976012254 | 2\|4\|4 | 3.33 | 3 | 0\|0\|0\|0\|0\|0 | FALSE |
| NSD1 | Q12923 | PTPN13 | bait_int | 0 | 0.996139505 | 25\|23\|22 | 23.33 | 3 | 0\|0\|0\|0\|0\|0 | FALSE |
| NSD1 | O95822 | MLYCD | bait_int | 0 | 0.998398969 | 22\|26\|21 | 23 | 3 | 0\|0\|0\|0\|0\|0 | FALSE |
| NSD1 | Q9H8H2 | DDX31 | bait_int | 0 | 0.977543976 | 2\|4\|3 | 3 | 3 | 0\|0\|0\|0\|0\|0 | FALSE |
| NSD1 | O75179 | ANKRD17 | bait_int | 0.04 | 0.984058996 | 1\|4\|4 | 3 | 3 | 0\|0\|0\|0\|0\|0 | FALSE |
| NSD1 | Q8N8A6 | DDX51 | bait_int | 0 | 0.979255902 | 3\|4\|4 | 3.67 | 3 | 0\|0\|0\|0\|0\|0 | FALSE |
| NSD1 | Q8IZL8 | PELP1 | bait_int | 0 | 0.97160422 | 3\|3\|6 | 4 | 3 | 0\|0\|0\|0\|0\|0 | FALSE |
| NSD1 | Q15397 | PUM3 | bait_int | 0 | 0.981972803 | 13\|14\|9 | 12 | 3 | 0\|0\|0\|0\|0\|0 | FALSE |
| NSD1 | Q14690 | PDCD11 | bait_int | 0 | 0.981421799 | 34\|36\|42 | 37.33 | 3 | 0\|0\|0\|0\|0\|0 | FALSE |
| NSD1 | O94906 | PRPF6 | bait_int | 0 | 0.975707295 | 3\|2\|2 | 2.33 | 3 | 0\|0\|0\|0\|0\|0 | FALSE |
| KMT2E | P49760 | CLK2 | bait_int | 0.05 | 0.973648827 | 3\|3\|1 | 2.33 | 3 | 0\|0\|0\|0\|0\|0 | FALSE |
| KMT2E | O15294 | OGT | bait_int | 0 | 0.995806823 | 19\|24\|30 | 24.33 | 3 | 0\|0\|0\|0\|0\|0 | TRUE |
| KMT2E | Q8IWV7 | UBR1 | bait_bait | 0 | 0.998745512 | 27\|33\|38 | 32.67 | 3 | 0\|0\|0\|0\|0\|0 | FALSE |
| KMT2E | Q13227 | GPS2 | bait_int | 0 | 0.981050997 | 4\|3\|3 | 3.33 | 3 | 0\|0\|0\|0\|0\|0 | FALSE |
| KMT2E | P85037 | FOXK1 | bait_int | 0 | 0.984516433 | 5\|5\|8 | 6 | 3 | 0\|0\|0\|0\|0\|0 | FALSE |
| KMT2E | Q10570 | CPSF1 | bait_int | 0 | 0.991398788 | 5\|7\|6 | 6 | 3 | 0\|0\|0\|0\|0\|0 | FALSE |
| KMT2E | Q32P28 | P3H1 | bait_int | 0 | 0.98894526 | 19\|12\|20 | 17 | 3 | 0\|0\|0\|0\|0\|0 | FALSE |
| KMT2E | O75718 | CRTAP | bait_int | 0 | 0.980087606 | 4\|4\|6 | 4.67 | 3 | 0\|0\|0\|0\|0\|0 | FALSE |
| KMT2E | O00308 | WWP2 | bait_int | 0 | 0.996680113 | 13\|13\|15 | 13.67 | 3 | 0\|0\|0\|0\|0\|0 | FALSE |
| KMT2E | P35579 | MYH9 | bait_int | 0.04 | 0.975977599 | 3\|28\|48 | 26.33 | 3 | 1\|6\|0\|4\|3\|11 | FALSE |
| KMT2E | P51610 | HCFC1 | bait_int | 0 | 0.98599964 | 53\|53\|57 | 54.33 | 3 | 1\|2\|3\|3\|2\|2 | FALSE |
| KMT2E | Q6UB99 | ANKRD11 | bait_bait | 0 | 0.998080149 | 24\|10\|13 | 15.67 | 3 | 0\|0\|0\|0\|0\|0 | FALSE |
| KMT2E | O75376 | NCOR1 | bait_int | 0 | 0.993595875 | 20\|10\|15 | 15 | 3 | 0\|0\|0\|0\|0\|0 | FALSE |
| KMT2E | O60907 | TBL1X | bait_int | 0 | 0.988931398 | 7\|6\|5 | 6 | 3 | 0\|0\|0\|0\|0\|0 | FALSE |
| KMT2E | O15379 | HDAC3 | bait_int | 0 | 0.981965872 | 4\|3\|4 | 3.67 | 3 | 0\|0\|0\|0\|0\|0 | FALSE |
| KMT2E | Q9Y618 | NCOR2 | bait_int | 0 | 0.989922513 | 12\|5\|10 | 9 | 3 | 0\|0\|0\|0\|0\|0 | FALSE |
| KMT2E | Q9BZK7 | TBL1XR1 | bait_bait | 0 | 0.994101828 | 15\|12\|10 | 12.33 | 3 | 0\|0\|0\|0\|0\|0 | FALSE |
| KMT2E | Q9P2I0 | CPSF2 | bait_int | 0.01 | 0.980461873 | 2\|2\|2 | 2 | 3 | 0\|0\|0\|0\|0\|0 | FALSE |
| UBR1 | P50213 | IDH3A | bait_int | 0 | 0.997095965 | 37\|30\|35 | 34 | 3 | 0\|0\|0\|0\|0\|0 | FALSE |
| UBR1 | P16949 | STMN1 | bait_int | 0 | 0.975256789 | 4\|3\|2 | 3 | 3 | 0\|0\|0\|0\|0\|0 | FALSE |
| UBR1 | Q4G0N4 | NADK2 | bait_int | 0 | 0.995917717 | 16\|32\|37 | 28.33 | 3 | 0\|0\|0\|0\|0\|2 | FALSE |
| UBR1 | Q13162 | PRDX4 | bait_int | 0 | 0.981508435 | 5\|8\|14 | 9 | 3 | 1\|0\|0\|0\|0\|0 | FALSE |
| UBR1 | Q9UPN9 | TRIM33 | bait_int | 0.01 | 0.982069835 | 4\|2\|2 | 2.67 | 3 | 0\|1\|0\|0\|0\|0 | FALSE |
| UBR1 | O00116 | AGPS | bait_int | 0 | 0.997400923 | 21\|31\|39 | 30.33 | 3 | 0\|0\|0\|0\|0\|0 | FALSE |
| UBR1 | A6NJ78 | METTL15 | bait_int | 0 | 0.99506522 | 23\|15\|14 | 17.33 | 3 | 1\|0\|1\|2\|0\|0 | FALSE |
| UBR1 | Q16891 | IMMT | bait_int | 0 | 0.98624222 | 48\|63\|66 | 59 | 3 | 0\|0\|0\|1\|0\|0 | FALSE |
| UBR1 | Q9BYD3 | MRPL4 | bait_int | 0 | 0.984668912 | 11\|9\|8 | 9.33 | 3 | 0\|0\|0\|0\|0\|0 | FALSE |
| UBR1 | Q9BU61 | NDUFAF3 | bait_int | 0 | 0.994184999 | 9\|8\|10 | 9 | 3 | 0\|0\|0\|0\|0\|0 | FALSE |

| | | | | | | | | | | |
|---|---|---|---|---|---|---|---|---|---|---|
| UBR1 | Q8NI60 | COQ8A | bait_int | 0 | 0.985909538 | 28\|21\|26 | 25 | 3 | 1\|2\|0\|0\|0\|0 | FALSE |
| UBR1 | Q9P032 | NDUFAF4 | bait_int | 0 | 0.986360045 | 12\|9\|7 | 9.33 | 3 | 0\|0\|0\|0\|0\|0 | FALSE |
| UBR1 | P82933 | MRPS9 | bait_int | 0 | 0.986824413 | 12\|13\|16 | 13.67 | 3 | 0\|0\|0\|0\|0\|0 | FALSE |
| UBR1 | Q5SW79 | CEP170 | bait_int | 0.01 | 0.986886791 | 4\|2\|2 | 2.67 | 3 | 0\|1\|0\|0\|0\|0 | FALSE |
| UBR1 | Q6P1J9 | CDC73 | bait_int | 0 | 0.98926408 | 3\|4\|3 | 3.33 | 3 | 0\|1\|0\|0\|0\|0 | FALSE |
| UBR1 | Q6PI48 | DARS2 | bait_int | 0 | 0.994295893 | 12\|21\|25 | 19.33 | 3 | 0\|0\|0\|0\|0\|0 | FALSE |
| UBR1 | Q9NQ50 | MRPL40 | bait_int | 0 | 0.981245062 | 8\|9\|9 | 8.67 | 3 | 1\|0\|0\|0\|0\|1 | FALSE |
| UBR1 | Q7Z7F7 | MRPL55 | bait_int | 0 | 0.990338365 | 7\|8\|7 | 7.33 | 3 | 0\|0\|0\|0\|0\|0 | FALSE |
| UBR1 | O95299 | NDUFA10 | bait_int | 0 | 0.979602445 | 9\|9\|9 | 9 | 3 | 0\|0\|0\|0\|0\|0 | FALSE |
| UBR1 | O43615 | TIMM44 | bait_int | 0 | 0.983081743 | 8\|7\|6 | 7 | 3 | 0\|0\|0\|0\|0\|0 | FALSE |
| UBR1 | O60763 | USO1 | bait_int | 0 | 0.980579698 | 3\|3\|5 | 3.67 | 3 | 0\|0\|0\|0\|0\|0 | FALSE |
| UBR1 | O75306 | NDUFS2 | bait_int | 0 | 0.975041932 | 12\|9\|8 | 9.67 | 3 | 0\|0\|0\|0\|0\|0 | FALSE |
| UBR1 | Q9Y399 | MRPS2 | bait_int | 0 | 0.98918091 | 6\|7\|11 | 8 | 3 | 0\|0\|0\|0\|0\|0 | FALSE |
| UBR1 | P54886 | ALDH18A1 | bait_int | 0 | 0.973135942 | 75\|70\|73 | 72.67 | 3 | 4\|6\|3\|3\|1\|3 | FALSE |
| UBR1 | O75521 | ECI2 | bait_int | 0 | 0.982139144 | 15\|25\|23 | 21 | 3 | 0\|0\|0\|0\|0\|0 | FALSE |
| UBR1 | P48634 | PRRC2A | bait_int | 0 | 0.989645278 | 7\|3\|4 | 4.67 | 3 | 0\|0\|0\|0\|0\|0 | FALSE |
| UBR1 | P22830 | FECH | bait_int | 0 | 0.983636212 | 18\|24\|28 | 23.33 | 3 | 0\|1\|0\|0\|0\|0 | FALSE |
| UBR1 | Q13011 | ECH1 | bait_int | 0 | 0.997227651 | 18\|22\|22 | 20.67 | 3 | 0\|0\|0\|0\|0\|0 | FALSE |
| UBR1 | P36776 | LONP1 | bait_int | 0 | 0.997470232 | 65\|50\|60 | 58.33 | 3 | 0\|0\|0\|0\|1\|0 | FALSE |
| UBR1 | Q99700 | ATXN2 | bait_int | 0.01 | 0.976656825 | 3\|2\|5 | 3.33 | 3 | 0\|1\|1\|0\|0\|0 | FALSE |
| UBR1 | Q9BW92 | TARS2 | bait_int | 0 | 0.995155321 | 32\|34\|48 | 38 | 3 | 0\|0\|0\|0\|0\|0 | FALSE |
| UBR1 | Q96SZ6 | CDK5RAP1 | bait_int | 0 | 0.989950236 | 22\|16\|17 | 18.33 | 3 | 0\|0\|0\|0\|0\|0 | FALSE |
| UBR1 | Q96N67 | DOCK7 | bait_int | 0 | 0.990781941 | 10\|4\|4 | 6 | 3 | 1\|1\|0\|0\|0\|0 | FALSE |
| UBR1 | Q9H078 | CLPB | bait_int | 0 | 0.993332502 | 46\|46\|55 | 49 | 3 | 1\|1\|0\|1\|1\|0 | FALSE |
| UBR1 | Q9H2W6 | MRPL46 | bait_int | 0 | 0.971902247 | 12\|13\|10 | 11.67 | 3 | 1\|1\|1\|1\|0\|0 | FALSE |
| UBR1 | Q96CB9 | NSUN4 | bait_int | 0 | 0.976275627 | 8\|7\|11 | 8.67 | 3 | 0\|1\|1\|0\|0\|0 | FALSE |
| UBR1 | O60313 | OPA1 | bait_int | 0 | 0.978555884 | 11\|2\|7 | 6.67 | 3 | 0\|0\|0\|0\|0\|1 | FALSE |
| UBR1 | O75323 | NIPSNAP2 | bait_int | 0 | 0.990913627 | 19\|19\|23 | 20.33 | 3 | 0\|1\|0\|2\|0\|0 | FALSE |
| UBR1 | O75431 | MTX2 | bait_int | 0 | 0.983469872 | 5\|7\|6 | 6 | 3 | 0\|0\|0\|0\|0\|0 | FALSE |
| UBR1 | Q9ULF5 | SLC39A10 | bait_int | 0 | 0.996215744 | 17\|5\|9 | 10.33 | 3 | 0\|0\|0\|0\|0\|0 | FALSE |
| UBR1 | Q5VT52 | RPRD2 | bait_int | 0 | 0.989718052 | 4\|3\|4 | 3.67 | 3 | 0\|0\|0\|0\|0\|0 | FALSE |
| UBR1 | Q9NP81 | SARS2 | bait_int | 0 | 0.998627687 | 30\|26\|27 | 27.67 | 3 | 0\|0\|0\|0\|0\|0 | FALSE |
| UBR1 | Q9BTY2 | FUCA2 | bait_int | 0.04 | 0.989298734 | 5\|1\|5 | 3.67 | 3 | 0\|0\|0\|0\|0\|0 | FALSE |
| UBR1 | Q9H173 | SIL1 | bait_int | 0 | 0.996319707 | 15\|7\|10 | 10.67 | 3 | 0\|0\|0\|0\|0\|0 | TRUE |
| UBR1 | Q8IYU8 | MICU2 | bait_int | 0 | 0.996520703 | 9\|6\|8 | 7.67 | 3 | 0\|0\|0\|1\|0\|0 | FALSE |
| UBR1 | Q9BRJ2 | MRPL45 | bait_int | 0 | 0.99157206 | 9\|10\|10 | 9.67 | 3 | 0\|0\|0\|0\|0\|0 | FALSE |
| UBR1 | Q8NBJ5 | COLGALT1 | bait_int | 0 | 0.997331614 | 16\|26\|35 | 25.67 | 3 | 0\|0\|1\|0\|0\|0 | FALSE |
| UBR1 | Q01968 | OCRL | bait_int | 0 | 0.994226584 | 13\|9\|7 | 9.67 | 3 | 0\|0\|0\|0\|0\|0 | FALSE |
| UBR1 | Q9NYY8 | FASTKD2 | bait_int | 0 | 0.997754398 | 15\|8\|13 | 12 | 3 | 0\|0\|0\|0\|0\|0 | FALSE |
| UBR1 | O75414 | NME6 | bait_int | 0 | 0.995137994 | 6\|4\|7 | 5.67 | 3 | 0\|0\|0\|0\|0\|0 | FALSE |
| UBR1 | Q5ST30 | VARS2 | bait_int | 0 | 0.998087079 | 23\|9\|12 | 14.67 | 3 | 0\|0\|0\|0\|0\|0 | FALSE |
| UBR1 | Q15942 | ZYX | bait_int | 0 | 0.99093442 | 6\|3\|4 | 4.33 | 3 | 0\|0\|0\|0\|0\|0 | FALSE |
| UBR1 | Q92888 | ARHGEF1 | bait_int | 0 | 0.995044427 | 8\|3\|2 | 4.33 | 3 | 0\|0\|0\|0\|0\|0 | FALSE |
| UBR1 | Q9H300 | PARL | bait_int | 0.04 | 0.993166161 | 5\|1\|3 | 3 | 3 | 0\|0\|0\|0\|0\|0 | FALSE |
| UBR1 | Q9H8M1 | COQ10B | bait_int | 0 | 0.995460279 | 4\|4\|4 | 4 | 3 | 0\|0\|0\|0\|0\|0 | FALSE |
| UBR1 | Q9NVS2 | MRPS18A | bait_int | 0.01 | 0.992732981 | 2\|2\|2 | 2 | 3 | 0\|0\|0\|0\|0\|0 | FALSE |
| UBR1 | Q9Y305 | ACOT9 | bait_int | 0 | 0.997983116 | 19\|15\|16 | 16.67 | 3 | 0\|0\|0\|0\|0\|0 | FALSE |
| UBR1 | Q8IWR0 | ZC3H7A | bait_int | 0.04 | 0.993582013 | 13\|1\|4 | 6 | 3 | 0\|0\|0\|0\|0\|0 | FALSE |
| UBR1 | Q86YH6 | PDSS2 | bait_int | 0 | 0.996936555 | 6\|6\|9 | 7 | 3 | 0\|0\|0\|0\|0\|0 | FALSE |
| UBR1 | O14786 | NRP1 | bait_int | 0 | 0.997071707 | 8\|5\|9 | 7.33 | 3 | 0\|0\|0\|0\|0\|0 | TRUE |
| UBR1 | O95363 | FARS2 | bait_int | 0 | 0.997941531 | 7\|15\|12 | 11.33 | 3 | 0\|0\|0\|0\|0\|0 | FALSE |
| UBR1 | O95490 | ADGRL2 | bait_int | 0 | 0.994763727 | 7\|2\|3 | 4 | 3 | 0\|0\|0\|0\|0\|0 | FALSE |
| UBR1 | P53370 | NUDT6 | bait_int | 0 | 0.995460279 | 8\|2\|2 | 4 | 3 | 0\|0\|0\|0\|0\|0 | FALSE |
| UBR1 | Q4G148 | GXYLT1 | bait_int | 0 | 0.993387949 | 3\|2\|2 | 2.33 | 3 | 0\|0\|0\|0\|0\|0 | FALSE |
| UBR1 | Q5T2R2 | PDSS1 | bait_int | 0 | 0.995235026 | 3\|4\|4 | 3.67 | 3 | 0\|0\|0\|0\|0\|0 | FALSE |
| UBR1 | Q15836 | VAMP3 | bait_int | 0.01 | 0.977038023 | 2\|2\|2 | 2 | 3 | 0\|0\|0\|0\|0\|0 | FALSE |
| UBR1 | P45954 | ACADSB | bait_int | 0 | 0.993249331 | 3\|4\|4 | 3.67 | 3 | 0\|0\|0\|0\|0\|0 | FALSE |
| UBR1 | Q13586 | STIM1 | bait_int | 0.04 | 0.993166161 | 1\|3\|5 | 3 | 3 | 0\|0\|0\|0\|0\|0 | FALSE |
| UBR1 | Q96ER9 | CCDC51 | bait_int | 0 | 0.995137994 | 6\|2\|9 | 5.67 | 3 | 0\|0\|0\|0\|0\|0 | FALSE |
| UBR1 | Q13433 | SLC39A6 | bait_int | 0.05 | 0.99117007 | 7\|1\|2 | 3.33 | 3 | 0\|0\|0\|0\|0\|0 | FALSE |
| UBR1 | Q96S66 | CLCC1 | bait_int | 0 | 0.998759374 | 28\|19\|26 | 24.33 | 3 | 0\|0\|0\|0\|0\|0 | TRUE |
| UBR1 | P42785 | PRCP | bait_int | 0 | 0.98703927 | 2\|2\|4 | 2.67 | 3 | 0\|0\|0\|0\|0\|0 | FALSE |
| UBR1 | P45877 | PPIC | bait_int | 0 | 0.994763727 | 5\|4\|3 | 4 | 3 | 0\|0\|0\|0\|0\|0 | FALSE |
| UBR1 | Q9NVH6 | TMLHE | bait_int | 0 | 0.976739995 | 3\|3\|4 | 3.33 | 3 | 0\|0\|0\|0\|0\|0 | FALSE |
| UBR1 | Q96I59 | NARS2 | bait_int | 0.01 | 0.981342094 | 2\|2\|2 | 2 | 3 | 0\|0\|0\|0\|0\|0 | FALSE |
| UBR1 | Q9BPX6 | MICU1 | bait_int | 0.03 | 0.990885904 | 6\|1\|8 | 5 | 3 | 0\|0\|0\|0\|0\|0 | FALSE |
| UBR1 | Q9UG56 | PISD | bait_int | 0 | 0.990199748 | 8\|6\|7 | 7 | 3 | 0\|0\|0\|0\|0\|0 | FALSE |
| UBR1 | P51553 | IDH3G | bait_int | 0 | 0.997511817 | 11\|14\|14 | 13 | 3 | 0\|0\|0\|1\|0\|0 | FALSE |
| UBR1 | Q96D53 | COQ8B | bait_int | 0 | 0.995903855 | 14\|7\|9 | 10 | 3 | 0\|0\|0\|0\|0\|0 | FALSE |
| UBR1 | Q9Y3D9 | MRPS23 | bait_int | 0 | 0.97248444 | 6\|6\|8 | 6.67 | 3 | 0\|0\|0\|1\|1\|0 | FALSE |
| UBR1 | Q8N5N7 | MRPL50 | bait_int | 0 | 0.992105737 | 8\|7\|5 | 6.67 | 3 | 0\|0\|0\|0\|0\|0 | FALSE |
| UBR1 | Q96DV4 | MRPL38 | bait_int | 0 | 0.991551268 | 17\|3\|4 | 8 | 3 | 0\|1\|0\|0\|0\|0 | FALSE |

| | | | | | | | | | | |
|---|---|---|---|---|---|---|---|---|---|---|
| UBR1 | Q8TB37 | NUBPL | bait_int | 0 | 0.996700905 | 12\|12\|12 | 12 | 3 | 0\|0\|0\|0\|0\|0 | FALSE |
| UBR1 | Q9H4F8 | SMOC1 | bait_int | 0 | 0.994763727 | 5\|4\|3 | 4 | 3 | 0\|0\|0\|0\|0\|0 | FALSE |
| UBR1 | Q9UGM6 | WARS2 | bait_int | 0 | 0.998863337 | 21\|33\|35 | 29.67 | 3 | 0\|0\|0\|0\|0\|0 | FALSE |
| UBR1 | P09001 | MRPL3 | bait_int | 0 | 0.981889633 | 2\|3\|4 | 3 | 3 | 0\|0\|0\|0\|0\|0 | FALSE |
| UBR1 | P10321 | HLA-C | bait_int | 0 | 0.972047795 | 9\|6\|8 | 7.67 | 3 | 0\|0\|0\|0\|0\|0 | FALSE |
| UBR1 | Q9NRA8 | EIF4ENIF1 | bait_int | 0 | 0.983362443 | 5\|3\|3 | 3.67 | 3 | 0\|0\|0\|0\|0\|0 | FALSE |
| UBR1 | Q9H5Q4 | TFB2M | bait_int | 0 | 0.982180729 | 6\|6\|6 | 6 | 3 | 0\|0\|0\|0\|0\|0 | FALSE |
| UBR1 | Q9NRX2 | MRPL17 | bait_int | 0 | 0.977273672 | 3\|5\|4 | 4 | 3 | 0\|0\|0\|0\|0\|0 | FALSE |
| UBR1 | Q96TA2 | YME1L1 | bait_int | 0 | 0.992542382 | 60\|48\|50 | 52.67 | 3 | 0\|0\|0\|0\|0\|0 | FALSE |
| UBR1 | Q9Y4W6 | AFG3L2 | bait_int | 0 | 0.992036429 | 40\|29\|29 | 32.67 | 3 | 0\|0\|0\|0\|0\|0 | FALSE |
| UBR1 | P09110 | ACAA1 | bait_int | 0 | 0.997733605 | 6\|13\|17 | 12 | 3 | 0\|0\|0\|0\|0\|0 | FALSE |
| UBR1 | Q6UB28 | METAP1D | bait_int | 0 | 0.993200815 | 2\|5\|4 | 3.67 | 3 | 0\|0\|0\|0\|0\|0 | FALSE |
| UBR1 | Q13405 | MRPL49 | bait_int | 0 | 0.992476539 | 2\|3\|3 | 2.67 | 3 | 0\|0\|0\|0\|0\|0 | FALSE |
| UBR1 | Q9BVS5 | TRMT61B | bait_int | 0 | 0.998516794 | 29\|14\|18 | 20.33 | 3 | 0\|0\|0\|0\|0\|0 | FALSE |
| UBR1 | Q9Y2Q9 | MRPS28 | bait_int | 0 | 0.986055087 | 5\|7\|7 | 6.33 | 3 | 0\|0\|0\|0\|0\|0 | FALSE |
| UBR1 | Q14738 | PPP2R5D | bait_bait | 0.05 | 0.992292871 | 1\|3\|4 | 2.67 | 3 | 0\|0\|0\|0\|0\|0 | FALSE |
| UBR1 | Q96I51 | RCC1L | bait_int | 0 | 0.994995911 | 12\|17\|16 | 15 | 3 | 0\|0\|0\|0\|0\|0 | FALSE |
| UBR1 | Q02252 | ALDH6A1 | bait_int | 0 | 0.999029678 | 46\|43\|47 | 45.33 | 3 | 0\|0\|0\|0\|0\|0 | FALSE |
| UBR1 | Q8TCG1 | CIP2A | bait_int | 0 | 0.991315618 | 10\|12\|13 | 11.67 | 3 | 0\|0\|0\|0\|0\|0 | FALSE |
| DYRK1A | P11274 | BCR | bait_int | 0 | 0.997782121 | 12\|19\|19 | 16.67 | 3 | 0\|0\|0\|0\|0\|0 | FALSE |
| DYRK1A | Q6P2E9 | EDC4 | bait_int | 0 | 0.999085125 | 74\|75\|64 | 71 | 3 | 1\|0\|0\|0\|0\|0 | FALSE |
| DYRK1A | P61981 | YWHAG | bait_int | 0 | 0.977287534 | 13\|22\|22 | 19 | 3 | 0\|0\|0\|0\|0\|0 | TRUE |
| DYRK1A | Q04917 | YWHAH | bait_int | 0 | 0.972498302 | 18\|19\|19 | 18.67 | 3 | 0\|0\|0\|0\|0\|0 | TRUE |
| DYRK1A | Q16531 | DDB1 | bait_int | 0 | 0.985486755 | 44\|52\|59 | 51.67 | 3 | 0\|0\|0\|1\|1\|0 | FALSE |
| DYRK1A | Q8NCN4 | RNF169 | bait_int | 0 | 0.996146435 | 21\|21\|22 | 21.33 | 3 | 0\|0\|0\|0\|0\|0 | TRUE |
| DYRK1A | O43852 | CALU | bait_int | 0 | 0.981113375 | 13\|10\|23 | 15.33 | 3 | 0\|0\|0\|1\|1\|0 | FALSE |
| DYRK1A | Q93008 | USP9X | bait_int | 0 | 0.975845913 | 7\|14\|20 | 13.67 | 3 | 0\|0\|0\|0\|0\|0 | FALSE |
| DYRK1A | P10644 | PRKAR1A | bait_int | 0 | 0.988411583 | 12\|10\|10 | 10.67 | 3 | 0\|0\|0\|0\|0\|0 | FALSE |
| DYRK1A | P49959 | MRE11 | bait_int | 0 | 0.986997685 | 10\|13\|16 | 13 | 3 | 0\|0\|0\|0\|0\|0 | FALSE |
| DYRK1A | Q8IZH2 | XRN1 | bait_int | 0 | 0.998031632 | 39\|29\|33 | 33.67 | 3 | 0\|0\|0\|0\|0\|0 | FALSE |
| DYRK1A | P82933 | MRPS9 | bait_int | 0.04 | 0.975267185 | 1\|7\|4 | 4 | 3 | 0\|0\|0\|0\|0\|0 | FALSE |
| DYRK1A | Q86VM9 | ZC3H18 | bait_int | 0 | 0.991565129 | 8\|9\|16 | 11 | 3 | 0\|0\|0\|0\|0\|0 | FALSE |
| DYRK1A | Q9GZS3 | WDR61 | bait_int | 0 | 0.979449966 | 3\|4\|6 | 4.33 | 3 | 0\|0\|0\|0\|0\|0 | FALSE |
| DYRK1A | Q9H6S0 | YTHDC2 | bait_int | 0 | 0.978583607 | 10\|2\|10 | 7.33 | 3 | 0\|0\|0\|0\|0\|0 | FALSE |
| DYRK1A | Q9HCE1 | MOV10 | bait_int | 0 | 0.979290556 | 18\|11\|10 | 13 | 3 | 0\|0\|0\|0\|0\|0 | TRUE |
| DYRK1A | Q7L2E3 | DHX30 | bait_int | 0 | 0.971500256 | 19\|21\|18 | 19.33 | 3 | 0\|0\|0\|0\|0\|0 | FALSE |
| DYRK1A | Q96II8 | LRCH3 | bait_int | 0 | 0.99379687 | 21\|34\|24 | 26.33 | 3 | 0\|0\|0\|0\|0\|0 | TRUE |
| DYRK1A | Q9UBU9 | NXF1 | bait_int | 0 | 0.980399495 | 2\|6\|6 | 4.67 | 3 | 0\|0\|0\|0\|0\|0 | FALSE |
| DYRK1A | P61962 | DCAF7 | bait_int | 0 | 0.995730583 | 25\|30\|27 | 27.33 | 3 | 4\|1\|2\|0\|0\|0 | TRUE |
| DYRK1A | P56545 | CTBP2 | bait_int | 0 | 0.975586005 | 11\|12\|11 | 11.33 | 3 | 0\|0\|0\|0\|0\|0 | FALSE |
| DYRK1A | Q96N67 | DOCK7 | bait_int | 0 | 0.998308867 | 54\|81\|63 | 66 | 3 | 1\|1\|0\|0\|0\|0 | FALSE |
| DYRK1A | Q96F86 | EDC3 | bait_int | 0 | 0.987254127 | 26\|23\|17 | 22 | 3 | 0\|0\|0\|0\|0\|0 | FALSE |
| DYRK1A | Q70CQ2 | USP34 | bait_int | 0 | 0.997477163 | 36\|30\|38 | 34.67 | 3 | 0\|0\|0\|1\|1\|1 | FALSE |
| DYRK1A | O15027 | SEC16A | bait_int | 0 | 0.986748174 | 6\|7\|4 | 5.67 | 3 | 0\|0\|0\|0\|0\|0 | FALSE |
| DYRK1A | P28749 | RBL1 | bait_int | 0 | 0.986855602 | 9\|4\|8 | 7 | 3 | 0\|0\|0\|0\|0\|0 | TRUE |
| DYRK1A | Q01804 | OTUD4 | bait_int | 0 | 0.995287007 | 16\|18\|14 | 16 | 3 | 0\|0\|0\|0\|0\|0 | FALSE |
| DYRK1A | Q12815 | TROAP | bait_int | 0 | 0.994094897 | 6\|8\|10 | 8 | 3 | 0\|1\|0\|0\|0\|0 | TRUE |
| DYRK1A | Q9H0W5 | CCDC8 | bait_int | 0 | 0.996797937 | 18\|13\|19 | 16.67 | 3 | 0\|0\|0\|0\|0\|0 | FALSE |
| DYRK1A | Q5VUJ6 | LRCH2 | bait_int | 0 | 0.99029678 | 6\|12\|10 | 9.33 | 3 | 0\|0\|0\|0\|0\|0 | FALSE |
| DYRK1A | Q9NQH7 | XPNPEP3 | bait_int | 0 | 0.989797757 | 3\|6\|5 | 4.67 | 3 | 0\|0\|0\|0\|0\|0 | FALSE |
| DYRK1A | Q9Y2L9 | LRCH1 | bait_int | 0 | 0.983670867 | 2\|3\|2 | 2.33 | 3 | 0\|0\|0\|0\|0\|0 | FALSE |
| DYRK1A | Q96HP0 | DOCK6 | bait_int | 0 | 0.993297847 | 4\|7\|7 | 6 | 3 | 0\|0\|0\|0\|0\|0 | FALSE |
| DYRK1A | O95071 | UBR5 | bait_int | 0 | 0.978417266 | 2\|3\|4 | 3 | 3 | 0\|0\|0\|0\|0\|0 | FALSE |
| DYRK1A | Q9BRK4 | LZTS2 | bait_int | 0 | 0.993935487 | 13\|8\|7 | 9.33 | 3 | 0\|0\|0\|0\|0\|0 | TRUE |
| DYRK1A | P49356 | FNTB | bait_int | 0 | 0.998017771 | 9\|14\|16 | 13 | 3 | 0\|0\|0\|0\|0\|0 | TRUE |
| DYRK1A | Q09472 | EP300 | bait_int | 0 | 0.995335523 | 15\|10\|14 | 13 | 3 | 0\|0\|0\|0\|0\|0 | TRUE |
| DYRK1A | Q13363 | CTBP1 | bait_int | 0 | 0.987080856 | 8\|5\|7 | 6.67 | 3 | 0\|0\|0\|0\|0\|0 | FALSE |
| DYRK1A | Q13490 | BIRC2 | bait_int | 0.04 | 0.993679045 | 9\|1\|3 | 4.33 | 3 | 0\|0\|0\|0\|0\|0 | FALSE |
| DYRK1A | Q14153 | FAM53B | bait_int | 0 | 0.995869201 | 3\|5\|6 | 4.67 | 3 | 0\|0\|0\|0\|0\|0 | FALSE |
| DYRK1A | Q15477 | SKIV2L | bait_int | 0 | 0.993997865 | 4\|6\|4 | 4.67 | 3 | 0\|0\|0\|0\|0\|0 | FALSE |
| DYRK1A | Q6P1L5 | FAM117B | bait_int | 0 | 0.997276168 | 17\|26\|23 | 22 | 3 | 0\|0\|0\|0\|0\|0 | TRUE |
| DYRK1A | Q6PGP7 | TTC37 | bait_int | 0 | 0.997640038 | 9\|11\|9 | 9.67 | 3 | 0\|0\|0\|0\|0\|0 | FALSE |
| DYRK1A | Q70EL1 | USP54 | bait_int | 0 | 0.995924648 | 18\|12\|13 | 14.33 | 3 | 0\|0\|0\|0\|0\|0 | TRUE |
| DYRK1A | P49354 | FNTA | bait_int | 0 | 0.998482139 | 15\|24\|26 | 21.67 | 3 | 0\|0\|0\|0\|0\|0 | TRUE |
| DYRK1A | O15116 | LSM1 | bait_int | 0.01 | 0.976927129 | 2\|2\|2 | 2 | 3 | 0\|0\|0\|0\|0\|0 | FALSE |
| DYRK1A | O94887 | FARP2 | bait_int | 0.01 | 0.985355069 | 2\|2\|2 | 2 | 3 | 0\|0\|0\|0\|0\|0 | FALSE |
| DYRK1A | P06400 | RB1 | bait_int | 0 | 0.993491912 | 14\|11\|11 | 12 | 3 | 0\|0\|0\|0\|0\|0 | TRUE |
| DYRK1A | P41743 | PRKCI | bait_int | 0 | 0.990227471 | 7\|6\|7 | 6.67 | 3 | 0\|0\|0\|0\|0\|0 | FALSE |
| DYRK1A | Q86TB9 | PATL1 | bait_int | 0 | 0.99507215 | 9\|7\|6 | 7.33 | 3 | 0\|0\|0\|0\|0\|0 | FALSE |
| DYRK1A | Q9NPI6 | DCP1A | bait_int | 0 | 0.995023634 | 18\|11\|8 | 12.33 | 3 | 0\|0\|0\|0\|0\|0 | FALSE |
| DYRK1A | Q9NWF9 | RNF216 | bait_int | 0 | 0.997553402 | 17\|13\|15 | 15 | 3 | 0\|0\|0\|0\|0\|0 | FALSE |
| DYRK1A | Q9NYF3 | FAM53C | bait_int | 0 | 0.996167228 | 8\|9\|9 | 8.67 | 3 | 0\|0\|0\|0\|0\|0 | TRUE |

| | | | | | | | | | | |
|---|---|---|---|---|---|---|---|---|---|---|
| DYRK1A | Q9UPT8 | ZC3H4 | bait_int | 0 | 0.995373643 | 2\|6\|10 | 6 | 3 | 0\|0\|0\|0\|0\|0 | FALSE |
| DYRK1A | Q9Y4B4 | RAD54L2 | bait_int | 0 | 0.99920295 | 65\|66\|61 | 64 | 3 | 0\|0\|0\|0\|0\|0 | TRUE |
| DYRK1A | Q9Y4B6 | DCAF1 | bait_int | 0 | 0.999001955 | 97\|90\|94 | 93.67 | 3 | 1\|0\|0\|0\|0\|0 | FALSE |
| DYRK1A | Q86VQ1 | GLCCI1 | bait_int | 0 | 0.995758307 | 11\|13\|13 | 12.33 | 3 | 0\|0\|0\|0\|0\|0 | FALSE |
| DYRK1A | Q8IU60 | DCP2 | bait_int | 0 | 0.997071707 | 11\|5\|6 | 7.33 | 3 | 0\|0\|0\|0\|0\|0 | FALSE |
| DYRK1A | Q8NG27 | PJA1 | bait_int | 0.04 | 0.993166161 | 1\|4\|4 | 3 | 3 | 0\|0\|0\|0\|0\|0 | FALSE |
| DYRK1A | Q8TEW0 | PARD3 | bait_int | 0 | 0.998322729 | 48\|37\|44 | 43 | 3 | 0\|0\|0\|0\|0\|0 | FALSE |
| DYRK1A | Q8WV44 | TRIM41 | bait_int | 0 | 0.986339252 | 3\|3\|7 | 4.33 | 3 | 0\|0\|0\|0\|0\|0 | FALSE |
| DYRK1A | Q92574 | TSC1 | bait_int | 0 | 0.996881108 | 8\|12\|3 | 7.67 | 3 | 0\|0\|0\|0\|0\|0 | FALSE |
| DYRK1A | Q92793 | CREBBP | bait_bait | 0 | 0.998059356 | 19\|16\|19 | 18 | 3 | 0\|0\|0\|0\|0\|0 | TRUE |
| DYRK1A | Q99417 | MYCBP | bait_int | 0 | 0.987857113 | 4\|5\|5 | 4.67 | 3 | 0\|0\|0\|0\|0\|0 | FALSE |
| DYRK1A | Q9BPZ3 | PAIP2 | bait_int | 0 | 0.980610887 | 3\|2\|3 | 2.67 | 3 | 0\|0\|0\|0\|0\|0 | FALSE |
| DYRK1A | Q9BQ70 | TCF25 | bait_int | 0 | 0.984100581 | 4\|3\|4 | 3.67 | 3 | 0\|0\|0\|0\|0\|0 | FALSE |
| DYRK1A | Q9BW61 | DDA1 | bait_int | 0 | 0.994302824 | 11\|9\|10 | 10 | 3 | 0\|0\|0\|0\|0\|0 | FALSE |
| DYRK1A | Q9C073 | FAM117A | bait_int | 0 | 0.997671227 | 12\|11\|12 | 11.67 | 3 | 0\|0\|0\|0\|0\|0 | FALSE |
| DYRK1A | Q6Q0C0 | TRAF7 | bait_bait | 0 | 0.996985071 | 12\|5\|10 | 9 | 3 | 0\|0\|0\|0\|0\|0 | FALSE |
| DYRK1A | Q8WUQ7 | CACTIN | bait_int | 0.02 | 0.979845026 | 3\|3\|4 | 3.33 | 3 | 1\|1\|1\|1\|0\|0 | FALSE |
| DYRK1A | P25054 | APC | bait_int | 0 | 0.997366269 | 20\|8\|10 | 12.67 | 3 | 0\|0\|0\|0\|0\|0 | FALSE |
| DYRK1A | Q12933 | TRAF2 | bait_int | 0 | 0.997740536 | 35\|28\|28 | 30.33 | 3 | 0\|0\|0\|0\|0\|0 | FALSE |
| DYRK1A | Q96CN4 | EVI5L | bait_int | 0 | 0.996569219 | 10\|9\|10 | 9.67 | 3 | 0\|0\|0\|0\|0\|0 | FALSE |
| GABRB3 | P51148 | RAB5C | bait_int | 0.02 | 0.994808777 | 6\|9\|3 | 6 | 3 | 0\|1\|0\|0\|0\|4 | FALSE |
| GABRB3 | Q9NYU2 | UGGT1 | bait_int | 0 | 0.984918424 | 14\|13\|12 | 13 | 3 | 0\|0\|0\|0\|0\|1 | FALSE |
| STXBP1 | O60568 | PLOD3 | bait_int | 0 | 0.990158162 | 14\|19\|16 | 16.33 | 3 | 0\|0\|0\|0\|0\|0 | FALSE |
| STXBP1 | O00255 | MEN1 | bait_int | 0.01 | 0.994053312 | 2\|4\|2 | 2.67 | 3 | 0\|0\|0\|0\|0\|0 | FALSE |
| STXBP1 | Q15833 | STXBP2 | bait_int | 0 | 0.989475472 | 2\|3\|4 | 3 | 3 | 0\|0\|0\|0\|0\|0 | FALSE |
| STXBP1 | Q9NV70 | EXOC1 | bait_int | 0.01 | 0.991516613 | 2\|2\|3 | 2.33 | 3 | 0\|0\|0\|0\|0\|0 | FALSE |
| STXBP1 | O60232 | ZNRD2 | bait_int | 0.01 | 0.986353114 | 2\|3\|2 | 2.33 | 3 | 0\|0\|0\|0\|0\|0 | FALSE |
| STXBP1 | O75643 | SNRNP200 | bait_int | 0 | 0.975152826 | 15\|22\|26 | 21 | 3 | 1\|1\|2\|2\|2\|1 | FALSE |
| STXBP1 | Q96N67 | DOCK7 | bait_int | 0 | 0.993308244 | 6\|10\|14 | 10 | 3 | 0\|0\|2\|1\|0\|0 | FALSE |
| STXBP1 | Q6P2Q9 | PRPF8 | bait_int | 0 | 0.983372839 | 24\|32\|33 | 29.67 | 3 | 2\|1\|3\|1\|1\|0 | FALSE |
| STXBP1 | Q9BUQ8 | DDX23 | bait_int | 0.04 | 0.98568775 | 1\|5\|4 | 3.33 | 3 | 0\|0\|0\|0\|0\|0 | FALSE |
| STXBP1 | O94906 | PRPF6 | bait_int | 0 | 0.985375861 | 4\|7\|8 | 6.33 | 3 | 1\|0\|0\|0\|0\|0 | FALSE |
| STXBP1 | P49773 | HINT1 | bait_int | 0 | 0.984454055 | 5\|8\|4 | 5.67 | 3 | 0\|0\|0\|0\|0\|0 | FALSE |
| KMT2C | Q16531 | DDB1 | bait_int | 0 | 0.978916289 | 30\|21\|23 | 24.67 | 3 | 0\|0\|0\|0\|0\|0 | FALSE |
| KMT2C | Q9C005 | DPY30 | bait_int | 0 | 0.973524071 | 7\|5\|10 | 7.33 | 3 | 0\|0\|0\|0\|0\|0 | TRUE |
| KMT2C | Q6ZW49 | PAXIP1 | bait_int | 0 | 0.996111781 | 12\|10\|13 | 11.67 | 3 | 0\|0\|0\|0\|0\|0 | TRUE |
| KMT2C | Q9UBL3 | ASH2L | bait_int | 0 | 0.998988093 | 32\|30\|37 | 33 | 3 | 0\|0\|0\|0\|0\|0 | TRUE |
| KMT2C | Q9Y4C2 | TCAF1 | bait_int | 0 | 0.992875064 | 17\|4\|6 | 9 | 3 | 0\|0\|0\|0\|0\|0 | FALSE |
| KMT2C | Q15291 | RBBP5 | bait_int | 0 | 0.997179135 | 22\|19\|22 | 21 | 3 | 1\|0\|1\|0\|0\|1 | TRUE |
| KMT2C | Q14686 | NCOA6 | bait_int | 0 | 0.991908208 | 2\|4\|3 | 3 | 3 | 0\|0\|0\|0\|0\|0 | TRUE |
| KMT2C | Q9Y4B6 | DCAF1 | bait_int | 0 | 0.998461347 | 58\|44\|50 | 50.67 | 3 | 0\|0\|0\|0\|0\|0 | FALSE |
| KMT2C | Q9BW61 | DDA1 | bait_int | 0 | 0.989686863 | 3\|4\|5 | 4 | 3 | 0\|0\|0\|0\|0\|0 | FALSE |
| DYNC1H1 | O43237 | DYNC1LI2 | bait_int | 0.05 | 0.996732094 | 6\|15\|1 | 7.33 | 3 | 0\|0\|0\|0\|0\|0 | TRUE |
| DYNC1H1 | Q9Y6G9 | DYNC1LI1 | bait_int | 0.05 | 0.996638527 | 20\|24\|1 | 15 | 3 | 0\|0\|0\|0\|0\|0 | TRUE |
| DYNC1H1 | Q13409 | DYNC1I2 | bait_int | 0 | 0.996596942 | 18\|21\|2 | 13.67 | 3 | 0\|0\|0\|0\|0\|0 | TRUE |
| DYNC1H1 | O43852 | CALU | bait_int | 0.02 | 0.971167575 | 12\|5\|3 | 6.67 | 3 | 0\|0\|2\|0\|0\|0 | FALSE |
| DYNC1H1 | P43034 | PAFAH1B1 | bait_int | 0.05 | 0.981952011 | 9\|10\|0 | 6.33 | 3 | 0\|0\|0\|0\|0\|0 | TRUE |
| CACNA2D: | Q96TA2 | YME1L1 | bait_int | 0 | 0.972539887 | 6\|5\|4 | 5 | 3 | 0\|0\|0\|0\|0\|0 | FALSE |
| CACNA2D: | P46379 | BAG6 | bait_int | 0 | 0.971822542 | 4\|5\|3 | 4 | 3 | 0\|0\|0\|0\|0\|0 | FALSE |
| DSCAM | Q13627 | DYRK1A | bait_bait | 0 | 0.996645458 | 10\|13\|14 | 12.33 | 3 | 0\|0\|0\|0\|0\|0 | FALSE |
| DSCAM | O60306 | AQR | bait_int | 0.01 | 0.987971473 | 2\|3\|3 | 2.67 | 3 | 0\|0\|0\|0\|0\|0 | FALSE |
| DSCAM | Q9NVI1 | FANCI | bait_int | 0 | 0.98377483 | 6\|9\|10 | 8.33 | 3 | 0\|0\|0\|0\|0\|0 | FALSE |
| DSCAM | P20020 | ATP2B1 | bait_int | 0.01 | 0.984135235 | 2\|3\|3 | 2.67 | 3 | 0\|0\|0\|0\|0\|0 | FALSE |
| DSCAM | P61962 | DCAF7 | bait_int | 0 | 0.990019545 | 11\|6\|6 | 7.67 | 3 | 0\|0\|0\|0\|0\|0 | FALSE |
| DSCAM | P78527 | PRKDC | bait_int | 0 | 0.972941878 | 25\|45\|45 | 38.33 | 3 | 5\|3\|4\|1\|4\|5 | FALSE |
| DSCAM | O43592 | XPOT | bait_int | 0 | 0.97177056 | 11\|18\|14 | 14.33 | 3 | 0\|0\|0\|0\|0\|0 | FALSE |
| GRIN2B | P17066 | HSPA6 | bait_int | 0 | 0.97646276 | 4\|3\|5 | 4 | 3 | 0\|0\|0\|0\|0\|0 | FALSE |
| GRIN2B | P46379 | BAG6 | bait_int | 0 | 0.978625192 | 4\|8\|9 | 7 | 3 | 0\|0\|0\|0\|0\|0 | FALSE |
| GRIN2B | Q8IWV8 | UBR2 | bait_int | 0 | 0.991932466 | 12\|13\|8 | 11 | 3 | 0\|0\|0\|0\|0\|0 | FALSE |
| GRIN2B | P82675 | MRPS5 | bait_int | 0 | 0.979408381 | 5\|2\|5 | 4 | 3 | 0\|0\|0\|0\|0\|0 | FALSE |
| GRIN2B | Q53H96 | PYCR3 | bait_int | 0 | 0.973378523 | 5\|8\|3 | 5.33 | 3 | 0\|0\|0\|0\|0\|0 | FALSE |
| GRIN2B | Q9UK99 | FBXO3 | bait_int | 0 | 0.979041045 | 3\|2\|4 | 3 | 3 | 0\|0\|0\|0\|0\|0 | FALSE |
| GRIN2B | Q6AZZ1 | TRIM68 | bait_int | 0 | 0.993096852 | 5\|5\|8 | 6 | 3 | 0\|0\|0\|0\|0\|0 | FALSE |
| GRIN2B | Q13131 | PRKAA1 | bait_int | 0.01 | 0.972709694 | 3\|2\|3 | 2.67 | 3 | 0\|0\|0\|0\|0\|0 | FALSE |
| GRIN2B | O75694 | NUP155 | bait_bait | 0.01 | 0.978677174 | 7\|2\|2 | 3.67 | 3 | 0\|0\|0\|0\|0\|0 | FALSE |
| GRIN2B | Q9Y399 | MRPS2 | bait_int | 0 | 0.981362887 | 4\|2\|3 | 3 | 3 | 0\|0\|0\|0\|0\|0 | FALSE |
| GRIN2B | O95714 | HERC2 | bait_int | 0 | 0.99395628 | 7\|10\|14 | 10.33 | 3 | 0\|0\|0\|0\|0\|0 | FALSE |
| GRIN2B | P82933 | MRPS9 | bait_int | 0 | 0.978535091 | 2\|7\|7 | 5.33 | 3 | 0\|0\|0\|0\|0\|0 | FALSE |
| GRIN2B | Q93008 | USP9X | bait_int | 0 | 0.989146255 | 68\|57\|54 | 59.67 | 3 | 0\|0\|0\|0\|0\|0 | FALSE |
| GRIN2B | P61981 | YWHAG | bait_int | 0 | 0.977096935 | 21\|16\|19 | 18.67 | 3 | 0\|0\|0\|0\|1\|0 | FALSE |
| GRIN2B | P14735 | IDE | bait_int | 0 | 0.998717789 | 138\|134\|1: | 132.67 | 3 | 0\|0\|0\|0\|0\|0 | FALSE |
| MBD5 | Q92560 | BAP1 | bait_int | 0 | 0.991308687 | 4\|6\|7 | 5.67 | 3 | 0\|0\|0\|0\|0\|0 | TRUE |

| | | | | | | | | | | |
|---|---|---|---|---|---|---|---|---|---|---|
| MBD5 | P09497 | CLTB | bait_int | 0 | 0.980302463 | 5\|5\|4 | 4.67 | 3 | 0\|0\|0\|0\|0\|0 | FALSE |
| MBD5 | Q9NSY1 | BMP2K | bait_int | 0 | 0.987385814 | 7\|5\|6 | 6 | 3 | 0\|0\|0\|0\|0\|0 | FALSE |
| MBD5 | O94973 | AP2A2 | bait_int | 0 | 0.987642256 | 12\|11\|9 | 10.67 | 3 | 0\|0\|0\|0\|0\|0 | FALSE |
| MBD5 | Q96T76 | MMS19 | bait_int | 0 | 0.993484981 | 30\|25\|21 | 25.33 | 3 | 0\|0\|0\|0\|0\|0 | FALSE |
| MBD5 | O76071 | CIAO1 | bait_int | 0 | 0.993907764 | 8\|6\|11 | 8.33 | 3 | 0\|0\|0\|0\|0\|0 | FALSE |
| MBD5 | Q00610 | CLTC | bait_int | 0 | 0.988529408 | 89\|90\|87 | 88.67 | 3 | 3\|0\|2\|3\|10\|14 | FALSE |
| MBD5 | Q9P0K7 | RAI14 | bait_int | 0.04 | 0.971583427 | 6\|2\|2 | 3.33 | 3 | 0\|0\|0\|0\|2\|0 | FALSE |
| MBD5 | Q9UM54 | MYO6 | bait_int | 0 | 0.987531362 | 27\|38\|26 | 30.33 | 3 | 1\|0\|1\|7\|10\|9 | FALSE |
| MBD5 | Q96CW1 | AP2M1 | bait_int | 0 | 0.981556951 | 8\|8\|10 | 8.67 | 3 | 0\|0\|0\|0\|0\|0 | FALSE |
| MBD5 | Q9BY89 | KIAA1671 | bait_int | 0.03 | 0.985105557 | 7\|9\|6 | 7.33 | 3 | 0\|0\|0\|1\|5\|1 | FALSE |
| MBD5 | P63010 | AP2B1 | bait_int | 0 | 0.982534204 | 5\|10\|5 | 6.67 | 3 | 0\|0\|0\|0\|0\|0 | FALSE |
| MBD5 | O95782 | AP2A1 | bait_int | 0 | 0.989631416 | 15\|14\|18 | 15.67 | 3 | 0\|0\|0\|0\|0\|0 | FALSE |
| MBD5 | P53680 | AP2S1 | bait_bait | 0 | 0.9906156 | 3\|4\|3 | 3.33 | 3 | 0\|0\|0\|0\|0\|0 | FALSE |
| MBD5 | O14974 | PPP1R12A | bait_int | 0.01 | 0.973898338 | 8\|10\|4 | 7.33 | 3 | 0\|0\|0\|0\|1\|2 | FALSE |
| MBD5 | Q86YT6 | MIB1 | bait_int | 0.01 | 0.978264787 | 2\|4\|2 | 2.67 | 3 | 0\|0\|0\|0\|0\|0 | FALSE |
| MBD5 | P09496 | CLTA | bait_int | 0 | 0.971784422 | 7\|6\|8 | 7 | 3 | 0\|0\|0\|0\|0\|0 | FALSE |
| NCOA1 | Q00610 | CLTC | bait_int | 0 | 0.98520952 | 58\|53\|60 | 57 | 3 | 3\|0\|2\|3\|10\|14 | FALSE |
| NCOA1 | Q96RN5 | MED15 | bait_int | 0 | 0.985043179 | 2\|4\|3 | 3 | 3 | 0\|0\|0\|0\|0\|0 | FALSE |
| NCOA1 | P09497 | CLTB | bait_int | 0 | 0.976407313 | 3\|2\|5 | 3.33 | 3 | 0\|0\|0\|0\|0\|0 | FALSE |
| PTEN | Q96L92 | SNX27 | bait_int | 0 | 0.972928016 | 4\|2\|3 | 3 | 3 | 0\|0\|0\|0\|0\|0 | TRUE |
| DEAF1 | Q14192 | FHL2 | bait_int | 0 | 0.988938329 | 9\|7\|3 | 6.33 | 3 | 0\|0\|0\|0\|0\|0 | TRUE |
| DEAF1 | Q8IY63 | AMOTL1 | bait_int | 0 | 0.971195298 | 4\|4\|2 | 3.33 | 3 | 0\|0\|0\|0\|0\|0 | FALSE |
| DEAF1 | Q8TAP6 | CEP76 | bait_int | 0 | 0.989229426 | 3\|4\|2 | 3 | 3 | 0\|0\|0\|0\|0\|0 | FALSE |
| DEAF1 | O75369 | FLNB | bait_int | 0 | 0.971389362 | 8\|12\|7 | 9 | 3 | 0\|0\|1\|0\|1\|2 | FALSE |
| DEAF1 | Q8WWI1 | LMO7 | bait_int | 0 | 0.976532069 | 10\|8\|10 | 9.33 | 3 | 0\|0\|0\|0\|0\|0 | FALSE |
| DEAF1 | P35580 | MYH10 | bait_int | 0 | 0.982194591 | 51\|62\|58 | 57 | 3 | 0\|0\|8\|1\|2\|8 | FALSE |
| DEAF1 | Q01082 | SPTBN1 | bait_int | 0.01 | 0.97940145 | 62\|61\|54 | 59 | 3 | 0\|0\|15\|8\|22\|3[illegible] | FALSE |
| DEAF1 | Q5VT25 | CDC42BPA | bait_int | 0.01 | 0.97781428 | 20\|18\|12 | 16.67 | 3 | 0\|0\|3\|0\|2\|10 | FALSE |
| SCN1A | Q5VT25 | CDC42BPA | bait_int | 0.04 | 0.978950943 | 10\|39\|6 | 18.33 | 3 | 0\|0\|3\|0\|2\|10 | FALSE |
| SCN1A | O95714 | HERC2 | bait_int | 0 | 0.998211835 | 59\|52\|54 | 55 | 3 | 0\|0\|0\|0\|0\|0 | FALSE |
| SCN1A | Q14204 | DYNC1H1 | bait_bait | 0.01 | 0.988765057 | 2\|3\|3 | 2.67 | 3 | 0\|0\|0\|1\|0\|1 | FALSE |
| SCN1A | Q5HYI8 | RABL3 | bait_int | 0 | 0.974847867 | 2\|5\|3 | 3.33 | 3 | 0\|0\|0\|0\|0\|0 | FALSE |
| SCN1A | Q96JN8 | NEURL4 | bait_int | 0 | 0.994968187 | 11\|13\|11 | 11.67 | 3 | 0\|0\|0\|0\|0\|0 | FALSE |
| SCN1A | Q99250 | SCN2A | bait_bait | 0 | 0.988328412 | 2\|3\|3 | 2.67 | 3 | 0\|0\|0\|0\|0\|0 | FALSE |
| SCN1A | Q8N766 | EMC1 | bait_int | 0 | 0.989617555 | 7\|10\|8 | 8.33 | 3 | 2\|0\|0\|3\|0\|1 | FALSE |
| SCN1A | Q92643 | PIGK | bait_int | 0 | 0.979630169 | 6\|5\|4 | 5 | 3 | 0\|0\|0\|0\|0\|0 | FALSE |
| SCN1A | Q9Y2G8 | DNAJC16 | bait_int | 0 | 0.995633551 | 24\|25\|24 | 24.33 | 3 | 0\|0\|0\|0\|0\|0 | FALSE |
| SCN1A | Q15392 | DHCR24 | bait_int | 0 | 0.976805839 | 9\|8\|9 | 8.67 | 3 | 0\|0\|0\|0\|0\|0 | FALSE |
| SCN1A | O15258 | RER1 | bait_int | 0 | 0.972432459 | 3\|3\|4 | 3.33 | 3 | 1\|0\|0\|0\|0\|0 | FALSE |
| SCN2A | Q5VT25 | CDC42BPA | bait_int | 0.01 | 0.98091238 | 20\|32\|14 | 22 | 3 | 0\|0\|3\|0\|2\|10 | FALSE |
| PRR12 | O00291 | HIP1 | bait_int | 0.05 | 0.980046021 | 1\|7\|5 | 4.33 | 3 | 0\|0\|0\|0\|0\|0 | FALSE |
| PRR12 | P09497 | CLTB | bait_int | 0 | 0.987656118 | 12\|12\|10 | 11.33 | 3 | 0\|0\|0\|0\|0\|0 | FALSE |
| PRR12 | P11717 | IGF2R | bait_int | 0 | 0.995598897 | 16\|17\|21 | 18 | 3 | 0\|0\|0\|0\|0\|0 | FALSE |
| PRR12 | Q96D71 | REPS1 | bait_int | 0 | 0.991301756 | 8\|11\|9 | 9.33 | 3 | 0\|0\|0\|0\|0\|0 | FALSE |
| PRR12 | Q9UEU0 | VTI1B | bait_int | 0 | 0.971084404 | 5\|3\|3 | 3.67 | 3 | 0\|0\|0\|0\|0\|0 | FALSE |
| PRR12 | Q9NSY1 | BMP2K | bait_int | 0 | 0.994483026 | 17\|27\|20 | 21.33 | 3 | 0\|0\|0\|0\|0\|0 | FALSE |
| PRR12 | Q10567 | AP1B1 | bait_int | 0 | 0.986900653 | 7\|6\|3 | 5.33 | 3 | 0\|0\|0\|0\|0\|0 | FALSE |
| PRR12 | O94973 | AP2A2 | bait_int | 0 | 0.992798825 | 22\|26\|27 | 25 | 3 | 0\|0\|0\|0\|0\|0 | FALSE |
| PRR12 | Q8IU81 | IRF2BP1 | bait_int | 0 | 0.992826548 | 22\|17\|23 | 20.67 | 3 | 0\|0\|0\|0\|0\|0 | FALSE |
| PRR12 | Q32P28 | P3H1 | bait_int | 0 | 0.98409365 | 10\|9\|8 | 9 | 3 | 0\|0\|0\|0\|0\|0 | FALSE |
| PRR12 | O14976 | GAK | bait_int | 0 | 0.994836501 | 8\|12\|9 | 9.67 | 3 | 0\|0\|0\|0\|0\|0 | FALSE |
| PRR12 | O15027 | SEC16A | bait_int | 0 | 0.995668205 | 31\|30\|35 | 32 | 3 | 0\|0\|0\|0\|0\|0 | FALSE |
| PRR12 | Q9UNK0 | STX8 | bait_int | 0 | 0.993166161 | 2\|3\|4 | 3 | 3 | 0\|0\|0\|0\|0\|0 | FALSE |
| PRR12 | O75718 | CRTAP | bait_int | 0 | 0.977280603 | 4\|3\|4 | 3.67 | 3 | 0\|0\|0\|0\|0\|0 | FALSE |
| PRR12 | Q2TAL8 | QRICH1 | bait_int | 0 | 0.998620757 | 24\|25\|19 | 22.67 | 3 | 0\|0\|0\|0\|0\|0 | FALSE |
| PRR12 | O43747 | AP1G1 | bait_int | 0 | 0.990324503 | 4\|5\|6 | 5 | 3 | 0\|0\|0\|0\|0\|0 | FALSE |
| PRR12 | Q9BXS5 | AP1M1 | bait_int | 0 | 0.983913447 | 4\|5\|5 | 4.67 | 3 | 0\|0\|0\|0\|0\|0 | FALSE |
| PRR12 | Q9H1B7 | IRF2BPL | bait_bait | 0.03 | 0.990629462 | 5\|5\|2 | 4 | 3 | 1\|0\|0\|3\|0\|1 | FALSE |
| PRR12 | Q5VZL5 | ZMYM4 | bait_int | 0 | 0.995952371 | 18\|17\|30 | 21.67 | 3 | 0\|0\|0\|0\|0\|0 | FALSE |
| PRR12 | P36776 | LONP1 | bait_int | 0 | 0.986713519 | 4\|3\|8 | 5 | 3 | 0\|0\|0\|0\|0\|0 | FALSE |
| PRR12 | P61962 | DCAF7 | bait_int | 0 | 0.987087786 | 4\|6\|5 | 5 | 3 | 0\|0\|0\|0\|0\|0 | FALSE |
| PRR12 | Q00610 | CLTC | bait_int | 0 | 0.991149277 | 129\|144\|1[illegible] | 138 | 3 | 0\|0\|0\|0\|0\|1 | FALSE |
| PRR12 | Q96CW1 | AP2M1 | bait_int | 0 | 0.986366976 | 16\|15\|16 | 15.67 | 3 | 0\|0\|0\|0\|0\|0 | FALSE |
| PRR12 | P63010 | AP2B1 | bait_int | 0 | 0.989610624 | 12\|20\|18 | 16.67 | 3 | 0\|0\|0\|0\|0\|0 | FALSE |
| PRR12 | P53680 | AP2S1 | bait_bait | 0 | 0.992819617 | 5\|6\|4 | 5 | 3 | 0\|0\|0\|0\|0\|0 | FALSE |
| PRR12 | O95782 | AP2A1 | bait_int | 0 | 0.991128484 | 20\|23\|19 | 20.67 | 3 | 0\|0\|0\|0\|0\|0 | FALSE |
| PRR12 | Q9UPQ9 | TNRC6B | bait_int | 0 | 0.991655231 | 6\|7\|6 | 6.33 | 3 | 1\|0\|0\|0\|0\|0 | FALSE |
| PRR12 | O00443 | PIK3C2A | bait_int | 0 | 0.9969019 | 47\|50\|50 | 49 | 3 | 0\|0\|0\|0\|0\|1 | FALSE |
| PRR12 | P14373 | TRIM27 | bait_int | 0 | 0.985070903 | 7\|14\|11 | 10.67 | 3 | 0\|0\|0\|0\|0\|0 | FALSE |
| PRR12 | Q9NYZ3 | GTSE1 | bait_int | 0 | 0.982256969 | 3\|2\|2 | 2.33 | 3 | 0\|0\|0\|0\|0\|0 | FALSE |
| PRR12 | Q8NDV7 | TNRC6A | bait_int | 0 | 0.976559793 | 3\|2\|3 | 2.67 | 3 | 0\|0\|0\|0\|0\|0 | FALSE |
| PRR12 | P14735 | IDE | bait_int | 0 | 0.992937442 | 15\|10\|11 | 12 | 3 | 0\|0\|0\|0\|0\|0 | FALSE |

| | | | | | | | | | | |
|---|---|---|---|---|---|---|---|---|---|---|
| PRR12 | P09496 | CLTA | bait_int | 0 | 0.976913267 | 9\|10\|13 | 10.67 | 3 | 0\|0\|0\|0\|0\|0 | FALSE |
| KDM6B | Q02809 | PLOD1 | bait_int | 0 | 0.994469165 | 15\|17\|20 | 17.33 | 3 | 0\|0\|0\|0\|0\|0 | FALSE |
| KDM6B | Q8IVL6 | P3H3 | bait_int | 0 | 0.998177181 | 20\|22\|21 | 21 | 3 | 0\|0\|0\|0\|0\|0 | FALSE |
| KDM6B | P09497 | CLTB | bait_int | 0 | 0.986124395 | 8\|10\|10 | 9.33 | 3 | 0\|0\|0\|0\|0\|0 | FALSE |
| KDM6B | Q9Y5X1 | SNX9 | bait_int | 0 | 0.994940464 | 27\|28\|24 | 26.33 | 3 | 0\|0\|0\|0\|0\|0 | FALSE |
| KDM6B | Q96D71 | REPS1 | bait_int | 0 | 0.986893722 | 2\|6\|6 | 4.67 | 3 | 0\|0\|0\|0\|0\|0 | FALSE |
| KDM6B | Q9NSY1 | BMP2K | bait_int | 0 | 0.99299982 | 13\|18\|15 | 15.33 | 3 | 0\|0\|0\|0\|0\|0 | FALSE |
| KDM6B | O94973 | AP2A2 | bait_int | 0 | 0.991218586 | 17\|23\|17 | 19 | 3 | 0\|0\|0\|0\|0\|0 | FALSE |
| KDM6B | Q32P28 | P3H1 | bait_int | 0 | 0.993020612 | 35\|35\|38 | 36 | 3 | 0\|0\|0\|0\|0\|0 | FALSE |
| KDM6B | O15027 | SEC16A | bait_int | 0 | 0.997290029 | 61\|56\|64 | 60.33 | 3 | 0\|0\|0\|0\|0\|0 | FALSE |
| KDM6B | P13674 | P4HA1 | bait_int | 0 | 0.980628214 | 3\|6\|3 | 4 | 3 | 0\|0\|0\|0\|0\|0 | FALSE |
| KDM6B | O75718 | CRTAP | bait_int | 0 | 0.990151232 | 18\|18\|15 | 17 | 3 | 0\|0\|0\|0\|0\|0 | FALSE |
| KDM6B | Q86UA1 | PRPF39 | bait_int | 0 | 0.989735379 | 6\|3\|4 | 4.33 | 3 | 0\|0\|0\|0\|0\|0 | FALSE |
| KDM6B | Q92791 | P3H4 | bait_int | 0 | 0.997601918 | 12\|13\|11 | 12 | 3 | 0\|0\|0\|0\|0\|0 | FALSE |
| KDM6B | P04637 | TP53 | bait_int | 0 | 0.980725246 | 34\|37\|37 | 36 | 3 | 0\|0\|0\|0\|0\|0 | FALSE |
| KDM6B | Q00610 | CLTC | bait_int | 0 | 0.990248264 | 117\|125\|1 | 118.67 | 3 | 0\|0\|0\|0\|0\|1 | FALSE |
| KDM6B | Q96CW1 | AP2M1 | bait_int | 0 | 0.982790646 | 8\|12\|10 | 10 | 3 | 0\|0\|0\|0\|0\|0 | FALSE |
| KDM6B | P63010 | AP2B1 | bait_int | 0 | 0.986256082 | 7\|15\|10 | 10.67 | 3 | 0\|0\|0\|0\|0\|0 | FALSE |
| KDM6B | O95782 | AP2A1 | bait_int | 0 | 0.989929444 | 14\|16\|20 | 16.67 | 3 | 0\|0\|0\|0\|0\|0 | FALSE |
| KDM6B | P53680 | AP2S1 | bait_bait | 0 | 0.99204336 | 5\|5\|3 | 4.33 | 3 | 0\|0\|0\|0\|0\|0 | FALSE |
| KDM6B | Q9UPQ9 | TNRC6B | bait_int | 0.01 | 0.983955033 | 2\|2\|2 | 2 | 3 | 1\|0\|0\|0\|0\|0 | FALSE |
| KDM6B | O43823 | AKAP8 | bait_int | 0 | 0.985334276 | 10\|11\|11 | 10.67 | 3 | 0\|0\|0\|0\|0\|0 | FALSE |
| KDM6B | O00443 | PIK3C2A | bait_int | 0 | 0.995647413 | 27\|38\|27 | 30.67 | 3 | 0\|0\|0\|0\|0\|1 | FALSE |
| KDM6B | Q8NDV7 | TNRC6A | bait_int | 0 | 0.975215204 | 2\|3\|2 | 2.33 | 3 | 0\|0\|0\|0\|0\|0 | FALSE |
| KDM6B | P09496 | CLTA | bait_int | 0 | 0.974556771 | 9\|9\|8 | 8.67 | 3 | 0\|0\|0\|0\|0\|0 | FALSE |
| ARID1B | Q92785 | DPF2 | bait_int | 0 | 0.996590011 | 13\|8\|16 | 12.33 | 3 | 0\|0\|0\|0\|0\|0 | TRUE |
| ARID1B | Q8TAQ2 | SMARCC2 | bait_bait | 0 | 0.996097919 | 5\|4\|8 | 5.67 | 3 | 0\|0\|0\|0\|0\|0 | TRUE |
| ARID1B | Q96TA2 | YME1L1 | bait_int | 0 | 0.982554996 | 15\|9\|13 | 12.33 | 3 | 0\|0\|0\|0\|0\|0 | FALSE |
| ARID1B | Q6IAN0 | DHRS7B | bait_int | 0 | 0.972699297 | 6\|4\|3 | 4.33 | 3 | 0\|0\|0\|0\|0\|0 | FALSE |
| ARID1B | Q9NVI1 | FANCI | bait_int | 0 | 0.972685435 | 3\|2\|4 | 3 | 3 | 0\|0\|0\|0\|0\|0 | FALSE |
| ARID1B | P51648 | ALDH3A2 | bait_int | 0 | 0.974931038 | 11\|9\|11 | 10.33 | 3 | 0\|0\|0\|0\|0\|0 | FALSE |
| ARID1B | Q9NX40 | OCIAD1 | bait_int | 0 | 0.986172911 | 6\|8\|6 | 6.67 | 3 | 0\|0\|0\|0\|0\|0 | FALSE |
| ARID1B | Q92925 | SMARCD2 | bait_int | 0 | 0.995494934 | 9\|9\|11 | 9.67 | 3 | 0\|0\|0\|0\|0\|0 | TRUE |
| ARID1B | Q12824 | SMARCB1 | bait_int | 0 | 0.99498898 | 8\|12\|9 | 9.67 | 3 | 0\|0\|0\|0\|0\|0 | TRUE |
| ARID1B | Q96GM5 | SMARCD1 | bait_int | 0 | 0.997345476 | 21\|18\|18 | 19 | 3 | 0\|0\|0\|0\|0\|0 | TRUE |
| ARID1B | Q01085 | TIAL1 | bait_int | 0 | 0.992476539 | 2\|3\|3 | 2.67 | 3 | 0\|0\|0\|0\|0\|0 | FALSE |
| ARID1B | Q15532 | SS18 | bait_int | 0 | 0.995235026 | 3\|5\|3 | 3.67 | 3 | 0\|0\|0\|0\|0\|0 | TRUE |
| ARID1B | P16278 | GLB1 | bait_int | 0 | 0.978500437 | 2\|5\|7 | 4.67 | 3 | 0\|0\|0\|0\|0\|0 | FALSE |
| ARID1B | P30084 | ECHS1 | bait_int | 0 | 0.976338005 | 2\|2\|2 | 2 | 3 | 0\|0\|0\|0\|0\|0 | FALSE |
| ARID1B | O96019 | ACTL6A | bait_int | 0 | 0.993270124 | 11\|10\|10 | 10.33 | 3 | 0\|0\|0\|0\|0\|0 | TRUE |
| ARID1B | Q92922 | SMARCC1 | bait_int | 0 | 0.998780167 | 41\|33\|39 | 37.67 | 3 | 0\|0\|0\|0\|0\|0 | TRUE |
| ARID1B | Q969G3 | SMARCE1 | bait_int | 0 | 0.996964278 | 18\|16\|20 | 18 | 3 | 0\|0\|0\|0\|0\|0 | TRUE |
| ARID1B | Q86VP6 | CAND1 | bait_int | 0 | 0.976490484 | 6\|8\|10 | 8 | 3 | 0\|0\|0\|0\|0\|0 | FALSE |
| ARID1B | Q9BT78 | COPS4 | bait_int | 0 | 0.978985598 | 2\|3\|2 | 2.33 | 3 | 0\|0\|0\|0\|0\|0 | FALSE |
| ARID1B | P51532 | SMARCA4 | bait_int | 0 | 0.998794028 | 42\|38\|42 | 40.67 | 3 | 0\|0\|0\|0\|0\|0 | TRUE |
| ARID1B | Q16891 | IMMT | bait_int | 0 | 0.974272605 | 19\|17\|16 | 17.33 | 3 | 0\|0\|0\|0\|0\|0 | FALSE |
| ARID1B | O00116 | AGPS | bait_int | 0 | 0.986685796 | 2\|4\|2 | 2.67 | 3 | 0\|0\|0\|0\|0\|0 | FALSE |
| ARID1B | O43852 | CALU | bait_int | 0 | 0.973187924 | 7\|6\|10 | 7.67 | 3 | 1\|0\|1\|1\|0\|1 | FALSE |
| ARID1B | Q01518 | CAP1 | bait_int | 0 | 0.97614394 | 11\|9\|10 | 10 | 3 | 0\|0\|0\|0\|0\|0 | FALSE |
| ARID1B | P31948 | STIP1 | bait_int | 0 | 0.978327165 | 16\|31\|30 | 25.67 | 3 | 0\|0\|0\|0\|0\|0 | FALSE |
| PHF2 | P42285 | MTREX | bait_int | 0 | 0.973125546 | 4\|5\|5 | 4.67 | 3 | 0\|0\|0\|0\|0\|0 | FALSE |
| PHF2 | P31939 | ATIC | bait_int | 0 | 0.97598453 | 5\|7\|8 | 6.67 | 3 | 0\|0\|0\|0\|0\|0 | FALSE |
| PHF2 | P14735 | IDE | bait_int | 0 | 0.988661094 | 8\|4\|5 | 5.67 | 3 | 0\|0\|0\|0\|0\|0 | FALSE |
| PHF2 | Q15691 | MAPRE1 | bait_int | 0 | 0.978978667 | 5\|8\|5 | 6 | 3 | 0\|0\|0\|0\|0\|0 | FALSE |
| PHF2 | Q15397 | PUM3 | bait_int | 0 | 0.978299441 | 11\|8\|6 | 8.33 | 3 | 0\|0\|0\|0\|0\|0 | FALSE |
| PHF2 | Q13868 | EXOSC2 | bait_int | 0 | 0.975291443 | 5\|4\|5 | 4.67 | 3 | 0\|0\|0\|0\|0\|0 | FALSE |
| PHF2 | Q6P158 | DHX57 | bait_int | 0 | 0.981681707 | 4\|4\|6 | 4.67 | 3 | 0\|0\|0\|0\|0\|0 | FALSE |
| PHF2 | P19784 | CSNK2A2 | bait_int | 0 | 0.982243107 | 37\|35\|33 | 35 | 3 | 2\|0\|0\|0\|0\|0 | FALSE |
| PHF2 | O60684 | KPNA6 | bait_int | 0 | 0.989406163 | 10\|7\|9 | 8.67 | 3 | 0\|0\|0\|0\|0\|0 | FALSE |
| PHF2 | Q8IZL8 | PELP1 | bait_int | 0 | 0.977876658 | 8\|6\|6 | 6.67 | 3 | 0\|0\|0\|0\|0\|0 | FALSE |
| PHF2 | Q6P1J9 | CDC73 | bait_int | 0 | 0.997546471 | 33\|24\|29 | 28.67 | 3 | 0\|0\|0\|0\|0\|0 | FALSE |
| PHF2 | Q14166 | TTLL12 | bait_int | 0.03 | 0.976733064 | 3\|1\|5 | 3 | 3 | 0\|0\|0\|0\|0\|0 | FALSE |
| PHF2 | Q14683 | SMC1A | bait_int | 0.03 | 0.975998392 | 5\|1\|3 | 3 | 3 | 0\|0\|0\|0\|0\|0 | FALSE |
| PHF2 | Q14692 | BMS1 | bait_int | 0 | 0.97812617 | 3\|4\|5 | 4 | 3 | 0\|0\|0\|0\|0\|0 | FALSE |
| PHF2 | Q96ST2 | IWS1 | bait_int | 0 | 0.993893902 | 23\|24\|27 | 24.67 | 3 | 0\|0\|0\|0\|0\|0 | FALSE |
| PHF2 | Q9BT78 | COPS4 | bait_int | 0.05 | 0.977135055 | 2\|1\|3 | 2 | 3 | 0\|0\|0\|0\|0\|0 | FALSE |
| PHF2 | Q8N7H5 | PAF1 | bait_int | 0 | 0.994871155 | 32\|32\|30 | 31.33 | 3 | 0\|0\|0\|0\|0\|0 | FALSE |
| PHF2 | Q8WVC0 | LEO1 | bait_int | 0 | 0.991274033 | 12\|13\|15 | 13.33 | 3 | 0\|0\|0\|0\|0\|0 | FALSE |
| PHF2 | Q9GZS3 | WDR61 | bait_int | 0 | 0.990331434 | 16\|16\|20 | 17.33 | 3 | 0\|0\|0\|0\|0\|0 | FALSE |
| PHF2 | Q9UBC1 | NFKBIL1 | bait_int | 0.04 | 0.983684729 | 4\|1\|2 | 2.33 | 3 | 0\|0\|0\|0\|0\|0 | FALSE |
| PHF2 | Q9UK58 | CCNL1 | bait_int | 0 | 0.991076503 | 4\|2\|2 | 2.67 | 3 | 0\|0\|0\|0\|0\|0 | FALSE |
| PHF2 | P30566 | ADSL | bait_int | 0 | 0.979900473 | 7\|7\|6 | 6.67 | 3 | 0\|0\|0\|0\|0\|0 | FALSE |

| | | | | | | | | | | |
|---|---|---|---|---|---|---|---|---|---|---|
| PHF2 | O15131 | KPNA5 | bait_int | 0 | 0.98687986 | 5\|6\|2 | 4.33 | 3 | 0\|0\|0\|0\|0\|0 | FALSE |
| PHF2 | P49588 | AARS1 | bait_int | 0 | 0.980947034 | 2\|4\|5 | 3.67 | 3 | 0\|0\|0\|0\|0\|0 | FALSE |
| PHF2 | Q6PD62 | CTR9 | bait_int | 0 | 0.994697883 | 30\|26\|30 | 28.67 | 3 | 0\|0\|0\|0\|0\|0 | FALSE |
| PHF2 | Q7L2E3 | DHX30 | bait_int | 0 | 0.979935127 | 41\|39\|36 | 38.67 | 3 | 0\|0\|0\|0\|0\|0 | FALSE |
| PHF2 | Q9BTT0 | ANP32E | bait_int | 0 | 0.978341027 | 5\|6\|4 | 5 | 3 | 0\|0\|0\|0\|0\|0 | FALSE |
| PHF2 | Q96T37 | RBM15 | bait_int | 0 | 0.981494573 | 19\|19\|16 | 18 | 3 | 0\|1\|0\|1\|1\|1 | FALSE |
| PHF2 | Q9NXF1 | TEX10 | bait_int | 0 | 0.986699658 | 5\|3\|4 | 4 | 3 | 0\|0\|0\|0\|0\|0 | FALSE |
| PHF2 | Q9H6S0 | YTHDC2 | bait_int | 0 | 0.97104975 | 3\|5\|4 | 4 | 3 | 0\|0\|0\|0\|0\|0 | FALSE |
| PHF2 | Q7Z4V5 | HDGFL2 | bait_int | 0 | 0.990671047 | 16\|15\|10 | 13.67 | 3 | 4\|0\|0\|1\|1\|0 | FALSE |
| PHF2 | Q86VP6 | CAND1 | bait_int | 0 | 0.975624125 | 8\|8\|6 | 7.33 | 3 | 0\|0\|0\|0\|0\|0 | FALSE |
| PHF2 | Q92688 | ANP32B | bait_int | 0 | 0.989444283 | 9\|10\|10 | 9.67 | 3 | 0\|0\|0\|0\|0\|0 | FALSE |
| PHF2 | Q9UBU9 | NXF1 | bait_int | 0 | 0.991322549 | 18\|19\|24 | 20.33 | 3 | 0\|0\|0\|0\|0\|0 | FALSE |
| PHF2 | Q9UQE7 | SMC3 | bait_int | 0.04 | 0.975890964 | 4\|2\|1 | 2.33 | 3 | 0\|0\|0\|0\|0\|0 | FALSE |
| PHF2 | P52292 | KPNA2 | bait_int | 0 | 0.974425084 | 27\|33\|29 | 29.67 | 3 | 1\|1\|0\|0\|0\|0 | FALSE |
| PHF2 | P52294 | KPNA1 | bait_int | 0 | 0.987406607 | 13\|17\|10 | 13.33 | 3 | 0\|0\|0\|0\|0\|0 | FALSE |
| PHF2 | O14617 | AP3D1 | bait_int | 0 | 0.9916483 | 5\|4\|5 | 4.67 | 3 | 2\|0\|0\|0\|0\|0 | FALSE |
| PHF2 | Q9NVC6 | MED17 | bait_int | 0 | 0.984162959 | 2\|2\|3 | 2.33 | 3 | 0\|0\|0\|0\|0\|0 | FALSE |
| PHF2 | Q9H4L4 | SENP3 | bait_int | 0.05 | 0.97952274 | 2\|3\|1 | 2 | 3 | 0\|0\|0\|0\|0\|0 | FALSE |
| PHF2 | P09132 | SRP19 | bait_int | 0 | 0.991558199 | 5\|11\|8 | 8 | 3 | 0\|0\|0\|0\|0\|0 | FALSE |
| PHF2 | Q96ST3 | SIN3A | bait_bait | 0.05 | 0.987718496 | 3\|1\|2 | 2 | 3 | 0\|0\|0\|0\|0\|0 | FALSE |
| PHF2 | O43818 | RRP9 | bait_int | 0 | 0.99626426 | 5\|7\|4 | 5.33 | 3 | 0\|0\|0\|0\|0\|0 | FALSE |
| PHF2 | Q7KZ85 | SUPT6H | bait_int | 0.03 | 0.974095868 | 4\|3\|0 | 2.33 | 3 | 0\|0\|0\|0\|0\|0 | FALSE |
| PHF2 | Q96S94 | CCNL2 | bait_int | 0 | 0.995869201 | 6\|5\|3 | 4.67 | 3 | 0\|0\|0\|0\|0\|0 | FALSE |
| PHF2 | Q96SY0 | INTS14 | bait_int | 0 | 0.994053312 | 3\|3\|2 | 2.67 | 3 | 0\|0\|0\|0\|0\|0 | FALSE |
| PHF2 | Q9UPN4 | CEP131 | bait_int | 0 | 0.995127597 | 11\|11\|9 | 10.33 | 3 | 0\|0\|0\|0\|0\|0 | FALSE |
| PHF2 | Q8NB90 | SPATA5 | bait_int | 0.05 | 0.980212362 | 3\|1\|2 | 2 | 3 | 0\|0\|0\|0\|0\|0 | FALSE |

**# Supplemental Table:  ASDmut-PPI: Differential interactions of hcASD mutants**
#
### Table shows results of AP-MS experiments using mutated hcASD as bait and a quantitative
### comparison of interactors with wild-type baits performed in parallel AP-MS experiments.
#
### Differential statistics (log2FC, SE, Tvalue, DF, pvalue) are as computed by MSstats for
### the comparison of mutant/WT. The number of replicates with observations for the prey in
### either the mutant or WT AP-MS are given in columns obsCountWT and obsCountMut. Column
### change describes the significance and direction of change, based on thresholds of
### pvalue < 0.05 and absolute(log2FC) > 1.
#
### Column issue describes issues around completely missing values in one or both conditions. Where
### a prey is completely missing in either mutant or WT bait, the log2FC is assigned value 8
### or -8 as a convenience.

| Bait | mutant | PreyGene | Protein | log2FC | SE | Tvalue | DF | pvalue | issue | obs_Mut | obs_WT | change |
|---|---|---|---|---|---|---|---|---|---|---|---|---|
| FOXP1 | L327P | SESTD1 | Q86VW0 | 8 | | | | | oneConditionMissi | 3 | 0 | up |
| FOXP1 | L327P | VPS33B | Q9H267 | 8 | | | | | oneConditionMissi | 3 | 0 | up |
| RORB | Y22D | GIGYF1 | O75420 | 8 | | | | | oneConditionMissi | 3 | 0 | up |
| STXBP1 | A251T | STXBP3 | O00186 | 8 | | | | | oneConditionMissi | 3 | 0 | up |
| STXBP1 | R551C | STXBP3 | O00186 | 8 | | | | | oneConditionMissi | 3 | 0 | up |
| TBR1 | K228E | RNGTT | O60942 | 8 | | | | | oneConditionMissi | 3 | 0 | up |
| TBR1 | N374H | RNGTT | O60942 | 8 | | | | | oneConditionMissi | 2 | 0 | up |
| TRAF7 | R593W | TM9SF4 | Q92544 | 8 | | | | | oneConditionMissi | 3 | 0 | up |
| KCNQ3 | R230C | ELOB | Q15370 | 4.294982981 | 0.2108906 | 20.365926 | 10 | 1.80E-09 | 0 | 3 | 3 | up |
| AP2S1 | R10W | EIF3G | O75821 | 3.475813887 | 0.5474906 | 6.3486276 | 1 | 0.0994596 | 0 | 2 | 1 | not sig |
| GNAI1 | I319T | TNPO1 | Q92973 | 2.974979678 | 0.4372931 | 6.8031715 | 4 | 0.0024389 | 0 | 3 | 3 | up |
| AP2S1 | G64D | EPS15L1 | Q9UBC2 | 2.785569169 | 0.5566434 | 5.004226 | 5 | 0.0040899 | 0 | 3 | 2 | up |
| TLK2 | D529G | ASF1B | Q9NVP2 | 2.743076058 | 0.273111 | 10.043812 | 4 | 0.0005526 | 0 | 3 | 3 | up |
| CHD8 | R1580W | FXR1 | P51114 | 2.727546455 | 0.2689068 | 10.143091 | 10 | 1.40E-06 | 0 | 3 | 2 | up |
| NSD1 | C1350R | PTPN13 | Q12923 | 2.692410072 | 0.9035147 | 2.9799295 | 2 | 0.0965753 | 0 | 1 | 2 | not sig |
| LRRC4C | I163N | PRDX4 | Q13162 | 2.682948905 | 0.2212856 | 12.124373 | 4 | 0.0002655 | 0 | 3 | 3 | up |
| SLC6A1 | G299V | YIF1B | Q5BJH7 | 2.588520223 | 0.3066319 | 8.4417826 | 11 | 3.90E-06 | 0 | 3 | 2 | up |
| AP2S1 | R10W | RHOT2 | Q8IXI1 | 2.54795136 | 0.7555514 | 3.372307 | 1 | 0.1835203 | 0 | 2 | 1 | not sig |
| AP2S1 | G64D | EPS15 | P42566 | 2.392714905 | 0.58484 | 4.0912296 | 5 | 0.0094355 | 0 | 3 | 2 | up |
| GNAI1 | I319T | RIC8A | Q9NPQ8 | 2.371597268 | 0.401742 | 5.9032839 | 5 | 0.0019852 | 0 | 3 | 3 | up |
| KCNMA1 | A1033P | RABL3 | Q5HYI8 | 2.228146108 | 0.8090118 | 2.7541577 | 8 | 0.0248985 | 0 | 3 | 3 | up |
| LRRC4C | I163N | ERP44 | Q9BS26 | 2.211553371 | 0.2380528 | 9.2901799 | 4 | 0.0007468 | 0 | 3 | 3 | up |
| AP2S1 | R10W | AFG3L2 | Q9Y4W6 | 2.200205356 | 0.6435382 | 3.4189195 | 2 | 0.0759343 | 0 | 2 | 2 | not sig |
| AP2S1 | G64D | AP1B1 | Q10567 | 2.16913984 | 0.6689385 | 3.2426597 | 5 | 0.0228816 | 0 | 3 | 2 | up |
| GNAI1 | I319T | GNG4 | P50150 | 2.133288998 | 0.6307652 | 3.3820653 | 4 | 0.0277325 | 0 | 3 | 3 | up |
| LRRC4C | I163N | ERLEC1 | Q96DZ1 | 2.067895127 | 0.2437998 | 8.4819382 | 4 | 0.0010592 | 0 | 3 | 3 | up |
| SLC6A1 | G299V | TM9SF3 | Q9HD45 | 2.043601793 | 0.2177489 | 9.3851309 | 14 | 2.04E-07 | 0 | 3 | 3 | up |
| LRRC4C | I163N | SEL1L | Q9UBV2 | 2.010770835 | 0.3419467 | 5.8803641 | 4 | 0.0041795 | 0 | 3 | 3 | up |
| AP2S1 | R10W | SLC25A13 | Q9UJS0 | 2.008399848 | 0.4081878 | 4.9202844 | 5 | 0.0043964 | 0 | 3 | 2 | up |
| KCNMA1 | A1033P | RRBP1 | Q9P2E9 | 1.985486655 | 1.2951377 | 1.5330313 | 6 | 0.1761587 | 0 | 3 | 3 | not sig |
| CREBBP | R413Q | EP300 | Q09472 | 1.982539453 | 1.0158077 | 1.9516878 | 4 | 0.1227192 | 0 | 3 | 3 | not sig |
| STXBP1 | R551C | STX2 | P32856 | 1.981946691 | 0.2055336 | 9.6429349 | 6 | 7.12E-05 | 0 | 3 | 3 | up |
| STXBP1 | R551C | STX3 | Q13277 | 1.953021158 | 0.5097043 | 3.8316751 | 5 | 0.0122268 | 0 | 3 | 2 | up |
| DPYSL2 | H333N | DPYSL5 | Q9BPU6 | 1.932861561 | 0.6607944 | 2.9250574 | 7 | 0.0221797 | 0 | 2 | 3 | up |
| RORB | Y22D | OAT | P04181 | 1.925013311 | 0.3590239 | 5.3617967 | 4 | 0.0058393 | 0 | 3 | 3 | up |
| STXBP1 | A251T | STX3 | Q13277 | 1.808343182 | 0.5097043 | 3.5478282 | 5 | 0.0164254 | 0 | 3 | 2 | up |
| SLC6A1 | G299V | SLC39A14 | Q15043 | 1.788861052 | 0.1388214 | 12.88606 | 13 | 8.85E-09 | 0 | 3 | 3 | up |
| TBR1 | N374H | NXN | Q6DKJ4 | 1.784741531 | 1.2053473 | 1.4806865 | 4 | 0.2128018 | 0 | 1 | 3 | not sig |
| KCNMA1 | A1033P | SRP19 | P09132 | 1.760638526 | 0.8182252 | 2.1517773 | 5 | 0.0840559 | 0 | 2 | 2 | not sig |
| STXBP1 | A251T | STX2 | P32856 | 1.746688055 | 0.2055336 | 8.4983109 | 6 | 0.0001453 | 0 | 3 | 3 | up |
| AP2S1 | R10W | YME1L1 | Q96TA2 | 1.745321027 | 0.5506532 | 3.1695466 | 3 | 0.0504983 | 0 | 3 | 2 | not sig |
| LRRC4C | I163N | SDF4 | Q9BRK5 | 1.732102124 | 0.4562301 | 3.7965534 | 4 | 0.0191604 | 0 | 3 | 3 | up |
| TLK2 | D529G | ASF1A | Q9Y294 | 1.729411613 | 0.3089315 | 5.598042 | 4 | 0.0049985 | 0 | 3 | 3 | up |
| SLC6A1 | G299V | YIPF3 | Q9GZM5 | 1.719016387 | 0.2375487 | 7.2364794 | 14 | 4.31E-06 | 0 | 3 | 3 | up |
| AP2S1 | R10W | UMPS | P11172 | 1.715337808 | 0.4433801 | 3.8687751 | 3 | 0.0305524 | 0 | 3 | 2 | up |
| FOXP1 | L327P | CTBP1 | Q13363 | 1.712871822 | 0.8129658 | 2.1069419 | 8 | 0.068197 | 0 | 3 | 3 | not sig |
| SLC6A1 | G299V | RFT1 | Q96AA3 | 1.679022814 | 0.1767227 | 9.5008906 | 14 | 1.75E-07 | 0 | 3 | 3 | up |
| AP2S1 | G64D | AP2B1 | P63010 | 1.651102632 | 0.3917739 | 4.214427 | 5 | 0.0083725 | 0 | 3 | 2 | up |
| FOXP1 | L327P | FHL2 | Q14192 | 1.630264358 | 0.399445 | 4.0813239 | 7 | 0.0046819 | 0 | 3 | 2 | up |
| RORB | Y22D | BCKDK | O14874 | 1.609098931 | 0.401867 | 4.004058 | 4 | 0.0160758 | 0 | 3 | 3 | up |
| SLC6A1 | G299V | CERS2 | Q96G23 | 1.60042018 | 0.2014027 | 7.9463694 | 14 | 1.48E-06 | 0 | 3 | 3 | up |
| LRRC4C | I163N | UGGT1 | Q9NYU2 | 1.594562337 | 0.1852262 | 8.6087301 | 4 | 0.0010007 | 0 | 3 | 3 | up |
| MKX | L89F | GPS2 | Q13227 | 1.563858082 | 0.9034417 | 1.7310005 | 4 | 0.158497 | 0 | 2 | 2 | not sig |
| AP2S1 | G64D | RNF41 | Q9H4P4 | 1.540867552 | 0.7126674 | 2.1621132 | 5 | 0.0829652 | 0 | 3 | 2 | not sig |
| LRRC4C | I163N | SELENOF | O60613 | 1.537182426 | 0.1988017 | 7.7322394 | 3 | 0.0044978 | 0 | 2 | 3 | up |
| SLC6A1 | G299V | CERS1 | P27544 | 1.531077185 | 0.2321979 | 6.593846 | 14 | 1.20E-05 | 0 | 3 | 3 | up |
| SLC6A1 | H198P | CDS2 | O95674 | 1.529618995 | 0.2076177 | 7.3674797 | 14 | 3.52E-06 | 0 | 3 | 3 | up |

| | | | | | | | | | | | | |
|---|---|---|---|---|---|---|---|---|---|---|---|---|
| SLC6A1 | G360S | SLC39A14 | Q15043 | 1.512945452 | 0.1388214 | 10.898502 | 13 | 6.57E-08 | 0 | 3 | 3 | up |
| FOXP1 | L327P | CTBP2 | P56545 | 1.509341269 | 0.8248123 | 1.829921 | 8 | 0.1046498 | 0 | 3 | 3 | not sig |
| AP2S1 | G64D | AP2M1 | Q96CW1 | 1.508933035 | 0.6884246 | 2.1918639 | 5 | 0.0799087 | 0 | 3 | 2 | not sig |
| SLC6A1 | G360S | YIPF3 | Q9GZM5 | 1.503938247 | 0.2375487 | 6.3310729 | 14 | 1.86E-05 | 0 | 3 | 3 | up |
| TBR1 | N374H | SEC16A | O15027 | 1.50138142 | 0.4500807 | 3.3358046 | 4 | 0.0289486 | 0 | 2 | 2 | up |
| SLC6A1 | G299V | CDS2 | O95674 | 1.499938603 | 0.2076177 | 7.2245228 | 14 | 4.39E-06 | 0 | 3 | 3 | up |
| TBR1 | K228E | AP2M1 | Q96CW1 | 1.488016576 | 1.5088881 | 0.9861676 | 6 | 0.3621336 | 0 | 3 | 3 | not sig |
| SLC6A1 | G360S | RTN3 | O95197 | 1.484988578 | 0.1649028 | 9.0052371 | 14 | 3.36E-07 | 0 | 3 | 3 | up |
| SLC6A1 | G360S | CERS1 | P27544 | 1.480093171 | 0.2321979 | 6.3742746 | 14 | 1.73E-05 | 0 | 3 | 3 | up |
| NSD1 | C1350R | DDX51 | Q8N8A6 | 1.447982494 | 0.5242906 | 2.7617935 | 6 | 0.0327762 | 0 | 3 | 3 | up |
| LRRC4C | I163N | P3H1 | Q32P28 | 1.445359643 | 0.1918961 | 7.5319895 | 4 | 0.0016639 | 0 | 3 | 3 | up |
| SLC6A1 | G299V | RTN3 | O95197 | 1.439667872 | 0.1649028 | 8.7304042 | 14 | 4.88E-07 | 0 | 3 | 3 | up |
| SLC6A1 | G360S | ILVBL | A1L0T0 | 1.423569721 | 0.1997308 | 7.127442 | 14 | 5.11E-06 | 0 | 3 | 3 | up |
| MKX | R93G | SIN3A | Q96ST3 | 1.412096367 | 0.2821105 | 5.0054727 | 5 | 0.0040855 | 0 | 3 | 2 | up |
| RORB | Y22D | CUL7 | Q14999 | 1.405327843 | 0.3975174 | 3.5352608 | 4 | 0.024116 | 0 | 3 | 3 | up |
| SLC6A1 | H198P | YIPF3 | Q9GZM5 | 1.401428445 | 0.2375487 | 5.8995412 | 14 | 3.87E-05 | 0 | 3 | 3 | up |
| SLC6A1 | G360S | RFT1 | Q96AA3 | 1.396108818 | 0.1767227 | 7.8999981 | 14 | 1.58E-06 | 0 | 3 | 3 | up |
| SLC6A1 | G299V | ILVBL | A1L0T0 | 1.380718448 | 0.1997308 | 6.9128969 | 14 | 7.18E-06 | 0 | 3 | 3 | up |
| SLC6A1 | G299V | DERL2 | Q9GZP9 | 1.371972395 | 0.1679615 | 8.1683734 | 14 | 1.07E-06 | 0 | 3 | 3 | up |
| SLC6A1 | G360S | CDS2 | O95674 | 1.368434414 | 0.2076177 | 6.5911268 | 14 | 1.21E-05 | 0 | 3 | 3 | up |
| MYT1L | H522Q | CTBP2 | P56545 | 1.368290826 | 0.7198017 | 1.9009276 | 6 | 0.1060334 | 0 | 3 | 3 | not sig |
| AP2S1 | G64D | AP2A1 | O95782 | 1.365966624 | 0.4793474 | 2.8496383 | 5 | 0.0358389 | 0 | 3 | 2 | up |
| AP2S1 | R10W | HUWE1 | Q7Z6Z7 | 1.364193183 | 1.8536786 | 0.7359384 | 2 | 0.538377 | 0 | 3 | 1 | not sig |
| SLC6A1 | G360S | CHP1 | Q99653 | 1.353461853 | 0.1798068 | 7.5273131 | 14 | 2.76E-06 | 0 | 3 | 3 | up |
| MKX | R93G | HDAC1 | Q13547 | 1.342774419 | 0.4320055 | 3.1082347 | 5 | 0.0266039 | 0 | 3 | 2 | up |
| SLC6A1 | H198P | RFT1 | Q96AA3 | 1.331920325 | 0.1767227 | 7.5367822 | 14 | 2.72E-06 | 0 | 3 | 3 | up |
| SLC6A1 | G360S | CERS2 | Q96G23 | 1.330535108 | 0.2014027 | 6.6063423 | 14 | 1.18E-05 | 0 | 3 | 3 | up |
| MKX | L89F | TBL1X | O60907 | 1.326960522 | 0.4131735 | 3.21163 | 5 | 0.0236856 | 0 | 3 | 2 | up |
| SLC6A1 | H198P | SLC39A14 | Q15043 | 1.321551613 | 0.1388214 | 9.5197964 | 13 | 3.17E-07 | 0 | 3 | 3 | up |
| SLC6A1 | H198P | YIF1B | Q5BJH7 | 1.320068822 | 0.3066319 | 4.3050597 | 11 | 0.0012454 | 0 | 3 | 2 | up |
| SLC6A1 | G299V | ESYT1 | Q9BSJ8 | 1.317247296 | 0.1185535 | 11.110995 | 14 | 2.50E-08 | 0 | 3 | 3 | up |
| SLC6A1 | G299V | RABL3 | Q5HYI8 | 1.314746959 | 0.1988697 | 6.6110962 | 14 | 1.17E-05 | 0 | 3 | 3 | up |
| LRRC4C | I163N | P4HB | P07237 | 1.314398539 | 0.2132675 | 6.1631454 | 4 | 0.0035182 | 0 | 3 | 3 | up |
| SLC6A1 | H198P | DERL2 | Q9GZP9 | 1.313003714 | 0.1679615 | 7.8172889 | 14 | 1.79E-06 | 0 | 3 | 3 | up |
| SLC6A1 | G360S | TM9SF3 | Q9HD45 | 1.311290851 | 0.2177489 | 6.0220325 | 14 | 3.13E-05 | 0 | 3 | 3 | up |
| MKX | L89F | SAP30 | O75446 | 1.286966391 | 0.3094788 | 4.158496 | 5 | 0.008837 | 0 | 3 | 2 | up |
| SLC6A1 | H198P | ILVBL | A1L0T0 | 1.286207882 | 0.1997308 | 6.4397072 | 14 | 1.55E-05 | 0 | 3 | 3 | up |
| MKX | L89F | SIN3A | Q96ST3 | 1.280939308 | 0.2821105 | 4.5405589 | 5 | 0.0061655 | 0 | 3 | 2 | up |
| AP2S1 | R10W | SRPRB | Q9Y5M8 | 1.27591833 | 0.4855003 | 2.6280484 | 5 | 0.0466428 | 0 | 3 | 2 | up |
| MKX | L89F | HDAC3 | O15379 | 1.268734195 | 0.3947937 | 3.2136641 | 5 | 0.023632 | 0 | 3 | 2 | up |
| SLC6A1 | H198P | TM9SF3 | Q9HD45 | 1.265703065 | 0.2177489 | 5.812673 | 14 | 4.50E-05 | 0 | 3 | 3 | up |
| SLC6A1 | A288V | YIPF3 | Q9GZM5 | 1.255981688 | 0.2375487 | 5.2872594 | 14 | 0.0001148 | 0 | 3 | 3 | up |
| SLC6A1 | H198P | CERS2 | Q96G23 | 1.249727093 | 0.2014027 | 6.2051162 | 14 | 2.29E-05 | 0 | 3 | 3 | up |
| LRRC4C | I163N | GLA | P06280 | 1.238619052 | 0.4418967 | 2.8029605 | 4 | 0.0486645 | 0 | 3 | 3 | up |
| SLC6A1 | H198P | CERS1 | P27544 | 1.238415384 | 0.2321979 | 5.3334478 | 14 | 0.0001056 | 0 | 3 | 3 | up |
| TLK2 | D529G | RIF1 | Q5UIP0 | 1.236359594 | 0.4279214 | 2.8892211 | 4 | 0.0445978 | 0 | 3 | 3 | up |
| SLC6A1 | A288V | TM9SF3 | Q9HD45 | 1.234946266 | 0.2177489 | 5.6714241 | 14 | 5.77E-05 | 0 | 3 | 3 | up |
| SLC6A1 | A288V | DERL2 | Q9GZP9 | 1.211559256 | 0.1679615 | 7.2133145 | 14 | 4.47E-06 | 0 | 3 | 3 | up |
| SLC6A1 | G299V | CHP1 | Q99653 | 1.204945571 | 0.1798068 | 6.7013359 | 14 | 1.01E-05 | 0 | 3 | 3 | up |
| SLC6A1 | G360S | AUP1 | Q9Y679 | 1.203558134 | 0.1326649 | 9.072169 | 14 | 3.08E-07 | 0 | 3 | 3 | up |
| MKX | R93G | SAP30 | O75446 | 1.193313676 | 0.3094788 | 3.8558817 | 5 | 0.0119297 | 0 | 3 | 2 | up |
| SLC6A1 | G299V | AUP1 | Q9Y679 | 1.187097959 | 0.1326649 | 8.9480957 | 14 | 3.63E-07 | 0 | 3 | 3 | up |
| MKX | L89F | NCOR2 | Q9Y618 | 1.181090135 | 0.4876855 | 2.4218275 | 5 | 0.059982 | 0 | 3 | 2 | not sig |
| MKX | R93G | TBL1X | O60907 | 1.179505215 | 0.4131735 | 2.8547453 | 5 | 0.0356251 | 0 | 3 | 2 | up |
| TBR1 | K228E | SGPL1 | O95470 | 1.176658628 | 2.5110797 | 0.4685867 | 3 | 0.671312 | 0 | 1 | 3 | not sig |
| MYT1L | C504R | CTBP2 | P56545 | 1.174358934 | 0.7198017 | 1.6315035 | 6 | 0.153905 | 0 | 3 | 3 | not sig |
| MKX | R93G | NCOR2 | Q9Y618 | 1.172186355 | 0.4876855 | 2.4035703 | 5 | 0.0613495 | 0 | 3 | 2 | not sig |
| LRRC4C | I163N | CKAP4 | Q07065 | 1.172133537 | 0.254043 | 4.6139181 | 4 | 0.0099264 | 0 | 3 | 3 | up |
| NSD1 | C1877R | PTPN13 | Q12923 | 1.168605806 | 0.7377166 | 1.5840849 | 2 | 0.2540277 | 0 | 2 | 2 | not sig |
| MKX | R93G | SUDS3 | Q9H7L9 | 1.159413671 | 0.1898482 | 6.1070556 | 5 | 0.0017054 | 0 | 3 | 2 | up |
| SLC6A1 | G360S | RABL3 | Q5HYI8 | 1.158923234 | 0.1988697 | 5.8275495 | 14 | 4.39E-05 | 0 | 3 | 3 | up |
| SLC6A1 | G360S | DERL2 | Q9GZP9 | 1.157175528 | 0.1679615 | 6.8895277 | 14 | 7.45E-06 | 0 | 3 | 3 | up |
| SLC6A1 | G299V | DHCR24 | Q15392 | 1.155729438 | 0.2596211 | 4.4516009 | 14 | 0.0005477 | 0 | 3 | 3 | up |
| SLC6A1 | A288V | ILVBL | A1L0T0 | 1.150993327 | 0.1997308 | 5.7627232 | 14 | 4.91E-05 | 0 | 3 | 3 | up |
| SLC6A1 | H198P | RTN3 | O95197 | 1.150800319 | 0.1649028 | 6.9786595 | 14 | 6.46E-06 | 0 | 3 | 3 | up |
| MKX | R93G | SAP130 | Q9H0E3 | 1.148573285 | 0.4624249 | 2.483805 | 5 | 0.0555813 | 0 | 3 | 2 | not sig |
| SLC6A1 | H198P | RHOT2 | Q8IXI1 | 1.143552621 | 0.1586902 | 7.2061941 | 14 | 4.52E-06 | 0 | 3 | 3 | up |
| NSD1 | C1350R | MKI67 | P46013 | 1.141751784 | 0.5628081 | 2.0286697 | 6 | 0.0888273 | 0 | 3 | 3 | not sig |
| SLC6A1 | G360S | YIF1B | Q5BJH7 | 1.137957329 | 0.3066319 | 3.7111506 | 11 | 0.0034343 | 0 | 3 | 2 | up |
| GNAI1 | I319T | TNFAIP8 | O95379 | 1.136683633 | 0.6398138 | 1.7765851 | 6 | 0.1259707 | 0 | 3 | 3 | not sig |
| SLC6A1 | A288V | RFT1 | Q96AA3 | 1.130073052 | 0.1767227 | 6.3946126 | 14 | 1.67E-05 | 0 | 3 | 3 | up |
| AP2S1 | R10W | AP2A1 | O95782 | 1.129272327 | 0.4793474 | 2.3558538 | 5 | 0.0650852 | 0 | 3 | 2 | not sig |

| | | | | | | | | | | | | |
|---|---|---|---|---|---|---|---|---|---|---|---|---|
| CHD8 | R1580W | NUP153 | P49790 | 1.12851559 | 0.5825294 | 1.937268 | 12 | 0.0766073 | 0 | 3 | 3 | not sig |
| SLC6A1 | H198P | SLC35B2 | Q8TB61 | 1.117990687 | 0.2255566 | 4.9565858 | 14 | 0.0002108 | 0 | 3 | 3 | up |
| NSD1 | C1350R | PDCD11 | Q14690 | 1.117616201 | 0.3621618 | 3.0859584 | 6 | 0.0214986 | 0 | 3 | 3 | up |
| SLC6A1 | G360S | DHCR24 | Q15392 | 1.113296842 | 0.2596211 | 4.2881604 | 14 | 0.0007505 | 0 | 3 | 3 | up |
| SLC6A1 | A288V | AUP1 | Q9Y679 | 1.106277472 | 0.1326649 | 8.3388878 | 14 | 8.42E-07 | 0 | 3 | 3 | up |
| NSD1 | C1350R | PELP1 | Q8IZL8 | 1.101786218 | 0.663122 | 1.6615136 | 6 | 0.1476747 | 0 | 3 | 3 | not sig |
| SLC6A1 | A288V | CDS2 | O95674 | 1.095520755 | 0.2076177 | 5.2766257 | 14 | 0.000117 | 0 | 3 | 3 | up |
| MKX | L89F | HDAC1 | Q13547 | 1.090026392 | 0.4320055 | 2.5231773 | 5 | 0.0529689 | 0 | 3 | 2 | not sig |
| FOXP1 | L327P | CEP55 | Q53EZ4 | 1.089635541 | 0.6835288 | 1.5941326 | 4 | 0.1861313 | 0 | 3 | 2 | not sig |
| SLC6A1 | A288V | CHP1 | Q99653 | 1.079191559 | 0.1798068 | 6.0019518 | 14 | 3.24E-05 | 0 | 3 | 3 | up |
| AP2S1 | G64D | AAGAB | Q6PD74 | 1.077013853 | 0.719704 | 1.4964679 | 5 | 0.1947852 | 0 | 3 | 2 | not sig |
| SLC6A1 | G299V | RHOT2 | Q8IXI1 | 1.07062392 | 0.1586902 | 6.7466277 | 14 | 9.37E-06 | 0 | 3 | 3 | up |
| SLC6A1 | H198P | POR | P16435 | 1.06610584 | 0.1728276 | 6.16861 | 14 | 2.44E-05 | 0 | 3 | 3 | up |
| AP2S1 | R10W | EPS15L1 | Q9UBC2 | 1.064777724 | 0.5566434 | 1.9128545 | 5 | 0.1139682 | 0 | 3 | 2 | not sig |
| SLC6A1 | H198P | AUP1 | Q9Y679 | 1.061601888 | 0.1326649 | 8.0021326 | 14 | 1.37E-06 | 0 | 3 | 3 | up |
| SLC6A1 | G299V | POR | P16435 | 1.058167813 | 0.1728276 | 6.1226797 | 14 | 2.64E-05 | 0 | 3 | 3 | up |
| TBR1 | N374H | AP2M1 | Q96CW1 | 1.055518139 | 1.5088881 | 0.6995337 | 6 | 0.5104146 | 0 | 3 | 3 | not sig |
| SLC6A1 | A288V | RTN3 | O95197 | 1.053898162 | 0.1649028 | 6.3910275 | 14 | 1.68E-05 | 0 | 3 | 3 | up |
| NSD1 | C1350R | NUMA1 | Q14980 | 1.052924256 | 0.5229604 | 2.013392 | 6 | 0.0907272 | 0 | 3 | 3 | not sig |
| SLC6A1 | H198P | RABL3 | Q5HYI8 | 1.040986659 | 0.1988697 | 5.2345152 | 14 | 0.0001264 | 0 | 3 | 3 | up |
| MKX | L89F | SAP130 | Q9H0E3 | 1.035377633 | 0.4624249 | 2.2390179 | 5 | 0.0753074 | 0 | 3 | 2 | not sig |
| SLC6A1 | G360S | RHOT2 | Q8IXI1 | 1.019801383 | 0.1586902 | 6.4263651 | 14 | 1.58E-05 | 0 | 3 | 3 | up |
| MKX | R93G | HDAC2 | Q92769 | 1.005964825 | 0.2917663 | 3.4478442 | 5 | 0.0182801 | 0 | 3 | 2 | up |
| SLC6A1 | A288V | RABL3 | Q5HYI8 | 0.98803911 | 0.1988697 | 4.9682728 | 14 | 0.0002063 | 0 | 3 | 3 | not sig |
| SLC6A1 | H198P | CHP1 | Q99653 | 0.976039126 | 0.1798068 | 5.4282668 | 14 | 8.90E-05 | 0 | 3 | 3 | not sig |
| MKX | L89F | SUDS3 | Q9H7L9 | 0.971581804 | 0.1898482 | 5.1176765 | 5 | 0.0037144 | 0 | 3 | 2 | not sig |
| SLC6A1 | A288V | POR | P16435 | 0.961148383 | 0.1728276 | 5.5613142 | 14 | 7.01E-05 | 0 | 3 | 3 | not sig |
| MKX | R93G | HDAC3 | O15379 | 0.958892202 | 0.3947937 | 2.428844 | 5 | 0.0594653 | 0 | 3 | 2 | not sig |
| SLC6A1 | A288V | DHCR24 | Q15392 | 0.951813625 | 0.2596211 | 3.6661647 | 14 | 0.0025421 | 0 | 3 | 3 | not sig |
| MKX | L89F | MRPS9 | P82933 | 0.945814246 | 0.241319 | 3.9193531 | 5 | 0.0111893 | 0 | 3 | 2 | not sig |
| AP2S1 | R10W | GALK1 | P51570 | 0.944402695 | 0.6091835 | 1.5502763 | 1 | 0.3647099 | 0 | 2 | 1 | not sig |
| LRRC4C | I163N | GCN1 | Q92616 | 0.940011 | 0.182393 | 5.1537676 | 1 | 0.122009 | 0 | 1 | 2 | not sig |
| MKX | L89F | MAGED2 | Q9UNF1 | 0.931490816 | 0.1258191 | 7.4034144 | 5 | 0.0007076 | 0 | 3 | 2 | not sig |
| MKX | L89F | TBL1XR1 | Q9BZK7 | 0.929923681 | 0.2527546 | 3.6791561 | 5 | 0.0143062 | 0 | 3 | 2 | not sig |
| SLC6A1 | H198P | DHCR24 | Q15392 | 0.92833622 | 0.2596211 | 3.5757352 | 14 | 0.0030415 | 0 | 3 | 3 | not sig |
| MKX | R93G | IRF2BPL | Q9H1B7 | 0.92806834 | 0.3446328 | 2.6929195 | 5 | 0.0431477 | 0 | 3 | 2 | not sig |
| MKX | R93G | SAP30L | Q9HAJ7 | 0.91964892 | 0.4449706 | 2.0667631 | 5 | 0.0936242 | 0 | 3 | 2 | not sig |
| SLC6A1 | A288V | YIF1B | Q5BJH7 | 0.917015976 | 0.3066319 | 2.9906081 | 11 | 0.0122843 | 0 | 3 | 2 | not sig |
| CREBBP | R413Q | NCOA3 | Q9Y6Q9 | 0.912980445 | 0.504127 | 1.8110129 | 4 | 0.1443772 | 0 | 3 | 3 | not sig |
| SLC6A1 | G360S | POR | P16435 | 0.912700443 | 0.1728276 | 5.2809889 | 14 | 0.0001161 | 0 | 3 | 3 | not sig |
| AP2S1 | R10W | EPS15 | P42566 | 0.908092057 | 0.58484 | 1.5527187 | 5 | 0.1811971 | 0 | 3 | 2 | not sig |
| MYT1L | C504R | RORB | Q92753 | 0.904936054 | 0.6094434 | 1.4848566 | 6 | 0.1881202 | 0 | 3 | 3 | not sig |
| CHD8 | R1580W | BRD3 | Q15059 | 0.903232432 | 1.1163786 | 0.8090736 | 12 | 0.4342219 | 0 | 3 | 3 | not sig |
| RORB | Y22D | CCDC8 | Q9H0W5 | 0.901302845 | 0.3232037 | 2.7886523 | 4 | 0.04938 | 0 | 3 | 3 | not sig |
| SLC6A1 | A288V | RHOT2 | Q8IXI1 | 0.897980113 | 0.1586902 | 5.6586981 | 14 | 5.90E-05 | 0 | 3 | 3 | not sig |
| TRAF7 | R593W | BAG6 | P46379 | 0.895007654 | 0.1378852 | 6.4909624 | 6 | 0.0006359 | 0 | 3 | 3 | not sig |
| TBR1 | N374H | NAA15 | Q9BXJ9 | 0.892549971 | 0.3023794 | 2.9517549 | 5 | 0.0318207 | 0 | 3 | 3 | not sig |
| SLC6A1 | A288V | SLC39A14 | Q15043 | 0.890396911 | 0.1388214 | 6.4139737 | 13 | 2.29E-05 | 0 | 3 | 3 | not sig |
| MKX | R93G | TBL1XR1 | Q9BZK7 | 0.888318341 | 0.2527546 | 3.5145484 | 5 | 0.0170178 | 0 | 3 | 2 | not sig |
| SLC6A1 | G360S | SLC35B2 | Q8TB61 | 0.885869824 | 0.2255566 | 3.9274833 | 14 | 0.0015175 | 0 | 3 | 3 | not sig |
| CHD8 | L834P | COPS3 | Q9UNS2 | 0.882470711 | 0.3250456 | 2.7149135 | 6 | 0.0348776 | 0 | 2 | 3 | not sig |
| AP2S1 | R10W | AAGAB | Q6PD74 | 0.861141883 | 0.719704 | 1.1965224 | 5 | 0.2851291 | 0 | 3 | 2 | not sig |
| AP2S1 | R10W | CCDC32 | Q9BV29 | 0.859205599 | 0.5214971 | 1.6475752 | 5 | 0.1603562 | 0 | 3 | 2 | not sig |
| SLC6A1 | G299V | NUP210 | Q8TEM1 | 0.856801401 | 0.221156 | 3.8741943 | 14 | 0.0016852 | 0 | 3 | 3 | not sig |
| SLC6A1 | A288V | SLC35B2 | Q8TB61 | 0.853956943 | 0.2255566 | 3.7859983 | 14 | 0.0020055 | 0 | 3 | 3 | not sig |
| SLC6A1 | L54F | NPTN | Q9Y639 | 0.853607262 | 0.2023386 | 4.2187077 | 6 | 0.0055692 | 0 | 3 | 3 | not sig |
| MKX | L89F | MRPS5 | P82675 | 0.853244633 | 0.1867723 | 4.5683684 | 5 | 0.0060108 | 0 | 3 | 2 | not sig |
| MKX | R93G | MRPS5 | P82675 | 0.850941617 | 0.1867723 | 4.5560378 | 5 | 0.0060789 | 0 | 3 | 2 | not sig |
| MKX | L89F | HDAC2 | Q92769 | 0.847094975 | 0.2917663 | 2.9033337 | 5 | 0.0336598 | 0 | 3 | 2 | not sig |
| MKX | L89F | SAP30L | Q9HAJ7 | 0.844708319 | 0.4449706 | 1.8983462 | 5 | 0.1161095 | 0 | 3 | 2 | not sig |
| SLC6A1 | H198P | NUP155 | O75694 | 0.841532808 | 0.1733885 | 4.8534519 | 14 | 0.0002556 | 0 | 3 | 3 | not sig |
| SLC6A1 | A288V | CERS1 | P27544 | 0.838486222 | 0.2321979 | 3.6110844 | 14 | 0.0028354 | 0 | 3 | 3 | not sig |
| CHD8 | L834P | AP2M1 | Q96CW1 | 0.833676608 | 0.9235584 | 0.9026788 | 12 | 0.384458 | 0 | 3 | 3 | not sig |
| MKX | R93G | MRPS9 | P82933 | 0.832975129 | 0.241319 | 3.4517598 | 5 | 0.0182031 | 0 | 3 | 2 | not sig |
| SLC6A1 | G299V | MYH9 | P35579 | 0.821505992 | 0.3279728 | 2.5047988 | 14 | 0.0252316 | 0 | 3 | 3 | not sig |
| MYT1L | H522Q | PHF21A | Q96BD5 | 0.814333081 | 0.6022643 | 1.3521191 | 6 | 0.2250763 | 0 | 3 | 3 | not sig |
| SLC6A1 | G299V | UBAC2 | Q8NBM4 | 0.810395369 | 0.1337856 | 6.057417 | 14 | 2.95E-05 | 0 | 3 | 3 | not sig |
| AP2S1 | R10W | AP2M1 | Q96CW1 | 0.809731574 | 0.6884246 | 1.1762095 | 5 | 0.2924572 | 0 | 3 | 2 | not sig |
| MKX | L89F | IRF2BPL | Q9H1B7 | 0.806725379 | 0.3446328 | 2.340826 | 5 | 0.066312 | 0 | 3 | 2 | not sig |
| FOXP1 | R513H | FHL2 | Q14192 | 0.805840182 | 0.399445 | 2.0173997 | 7 | 0.0834499 | 0 | 3 | 2 | not sig |
| SLC6A1 | A288V | ESYT1 | Q9BSJ8 | 0.800600934 | 0.1185535 | 6.7530772 | 14 | 9.27E-06 | 0 | 3 | 3 | not sig |
| SLC6A1 | A288V | CERS2 | Q96G23 | 0.799225664 | 0.2014027 | 3.9682969 | 14 | 0.0014005 | 0 | 3 | 3 | not sig |

| | | | | | | | | | | | | |
|---|---|---|---|---|---|---|---|---|---|---|---|---|
| CHD8 | C1095Y | AP2M1 | Q96CW1 | 0.774521934 | 0.9235584 | 0.838628 | 12 | 0.4180706 | 0 | 3 | 3 | not sig |
| MKX | R93G | MAGED2 | Q9UNF1 | 0.766011522 | 0.1258191 | 6.0881982 | 5 | 0.0017293 | 0 | 3 | 2 | not sig |
| CHD8 | L834P | AP2B1 | P63010 | 0.755295505 | 0.9105977 | 0.8294503 | 12 | 0.4230429 | 0 | 3 | 3 | not sig |
| SLC6A1 | G299V | LMBR1 | Q8WVP7 | 0.753527208 | 0.5228505 | 1.4411905 | 11 | 0.1773859 | 0 | 3 | 2 | not sig |
| MKX | R93G | GPS2 | Q13227 | 0.749243877 | 0.8247257 | 0.9084765 | 4 | 0.4150121 | 0 | 3 | 2 | not sig |
| SLC6A1 | A288V | NUP210 | Q8TEM1 | 0.737730195 | 0.221156 | 3.3357906 | 14 | 0.0049006 | 0 | 3 | 3 | not sig |
| TRAF7 | R593W | PDCD5 | O14737 | 0.73710632 | 0.1941998 | 3.7956082 | 6 | 0.009014 | 0 | 3 | 3 | not sig |
| CHD8 | L834P | BRD3 | Q15059 | 0.734369009 | 1.1163786 | 0.6578136 | 12 | 0.5230768 | 0 | 3 | 3 | not sig |
| DPYSL2 | R496C | DPYSL5 | Q9BPU6 | 0.732631207 | 0.5910325 | 1.2395786 | 7 | 0.255074 | 0 | 3 | 3 | not sig |
| SLC6A1 | G299V | NUP155 | O75694 | 0.727764926 | 0.1733885 | 4.1973077 | 14 | 0.0008952 | 0 | 3 | 3 | not sig |
| RORB | Y22D | OBSL1 | O75147 | 0.72465525 | 0.2054413 | 3.5273112 | 4 | 0.0242893 | 0 | 3 | 3 | not sig |
| AP2S1 | R10W | AP2A2 | O94973 | 0.706341951 | 0.3422877 | 2.0635913 | 5 | 0.0940027 | 0 | 3 | 2 | not sig |
| AP2S1 | R10W | AP2B1 | P63010 | 0.700343931 | 0.3917739 | 1.7876226 | 5 | 0.1338742 | 0 | 3 | 2 | not sig |
| NSD1 | C1350R | PUM3 | Q15397 | 0.69120883 | 0.2386531 | 2.8962912 | 5 | 0.033937 | 0 | 2 | 3 | not sig |
| CHD8 | R1580W | AP2M1 | Q96CW1 | 0.685103615 | 0.9235584 | 0.7418086 | 12 | 0.4724729 | 0 | 3 | 3 | not sig |
| NSD1 | C1350R | ANKRD17 | O75179 | 0.683681585 | 0.4643586 | 1.4723139 | 5 | 0.200918 | 0 | 2 | 3 | not sig |
| SLC6A1 | H198P | IMMT | Q16891 | 0.678511395 | 0.1809264 | 3.7502068 | 14 | 0.0021525 | 0 | 3 | 3 | not sig |
| TBR1 | K228E | IDE | P14735 | 0.674409076 | 1.0208507 | 0.6606344 | 6 | 0.5333751 | 0 | 3 | 3 | not sig |
| LRRC4C | I163N | CLPB | Q9H078 | 0.674000787 | 0.6095471 | 1.1057404 | 3 | 0.3495469 | 0 | 2 | 3 | not sig |
| MYT1L | C504R | CTBP1 | Q13363 | 0.67345744 | 0.7264448 | 0.9270594 | 4 | 0.4063685 | 0 | 3 | 1 | not sig |
| TRAF7 | R593W | DLG1 | Q12959 | 0.671693663 | 0.3643008 | 1.8437885 | 6 | 0.114774 | 0 | 3 | 3 | not sig |
| KCNQ3 | R230H | NAA15 | Q9BXJ9 | 0.665384701 | 0.2555836 | 2.6033934 | 9 | 0.0285787 | 0 | 3 | 3 | not sig |
| SLC6A1 | G299V | SLC35B2 | Q8TB61 | 0.661951312 | 0.2255566 | 2.9347458 | 14 | 0.0108691 | 0 | 3 | 3 | not sig |
| SLC6A1 | G360S | IMMT | Q16891 | 0.660135095 | 0.1809264 | 3.648639 | 14 | 0.0026319 | 0 | 3 | 3 | not sig |
| SLC6A1 | A288V | NUP155 | O75694 | 0.656502954 | 0.1733885 | 3.7863117 | 14 | 0.0020043 | 0 | 3 | 3 | not sig |
| SLC6A1 | G360S | NUP155 | O75694 | 0.650815857 | 0.1733885 | 3.753512 | 14 | 0.0021385 | 0 | 3 | 3 | not sig |
| SLC6A1 | H198P | ESYT1 | Q9BSJ8 | 0.650143238 | 0.1185535 | 5.483965 | 14 | 8.05E-05 | 0 | 3 | 3 | not sig |
| RORB | Y22D | RANBP2 | P49792 | 0.6441488 | 0.2093103 | 3.0774827 | 4 | 0.0370222 | 0 | 3 | 3 | not sig |
| CHD8 | C1095Y | CLTC | Q00610 | 0.636460271 | 0.8366249 | 0.7607474 | 12 | 0.4614954 | 0 | 3 | 3 | not sig |
| CHD8 | C1095Y | BRD3 | Q15059 | 0.630996373 | 1.1163786 | 0.5652172 | 12 | 0.5823395 | 0 | 3 | 3 | not sig |
| CHD8 | Q696K | BRD3 | Q15059 | 0.624700783 | 1.1163786 | 0.5595779 | 12 | 0.5860607 | 0 | 3 | 3 | not sig |
| MKX | L89F | NCOR1 | O75376 | 0.620447882 | 0.4242599 | 1.462424 | 5 | 0.2034824 | 0 | 3 | 2 | not sig |
| CHD8 | L834P | KPNA6 | O60684 | 0.619637481 | 0.1163106 | 5.3274384 | 12 | 0.0001801 | 0 | 3 | 3 | not sig |
| TRAF7 | R593W | PFDN6 | O15212 | 0.619294986 | 0.1515763 | 4.0856981 | 6 | 0.0064605 | 0 | 3 | 3 | not sig |
| TRAF7 | R593W | HUWE1 | Q7Z6Z7 | 0.615640194 | 0.0839166 | 7.3363312 | 6 | 0.0003279 | 0 | 3 | 3 | not sig |
| CHD8 | L834P | AP2A1 | O95782 | 0.61510811 | 0.9726299 | 0.6324174 | 12 | 0.5389777 | 0 | 3 | 3 | not sig |
| AP2S1 | G64D | AP2A2 | O94973 | 0.608439271 | 0.3422877 | 1.7775668 | 5 | 0.1356193 | 0 | 3 | 2 | not sig |
| LRRC4C | I163N | PRKDC | P78527 | 0.60808943 | 0.3676931 | 1.653796 | 4 | 0.1735121 | 0 | 3 | 3 | not sig |
| CHD8 | L834P | AP2A2 | O94973 | 0.597473988 | 0.9054971 | 0.6598299 | 12 | 0.5218261 | 0 | 3 | 3 | not sig |
| CHD8 | C1095Y | AP2B1 | P63010 | 0.593666915 | 0.9105977 | 0.651953 | 12 | 0.5267219 | 0 | 3 | 3 | not sig |
| SLC6A1 | A288V | UBAC2 | Q8NBM4 | 0.592566365 | 0.1337856 | 4.4292227 | 14 | 0.0005718 | 0 | 3 | 3 | not sig |
| KCNQ3 | R230H | ARL10 | Q8N8L6 | 0.592525456 | 0.2217619 | 2.6718989 | 10 | 0.0234161 | 0 | 3 | 3 | not sig |
| TBR1 | N374H | SGPL1 | O95470 | 0.584586729 | 1.9851828 | 0.294475 | 3 | 0.7875957 | 0 | 2 | 3 | not sig |
| TRAF7 | R593W | PFDN4 | Q9NQP4 | 0.577627021 | 0.2261186 | 2.5545316 | 6 | 0.0432259 | 0 | 3 | 3 | not sig |
| TBR1 | N374H | PIGT | Q969N2 | 0.574564034 | 0.3520113 | 1.6322316 | 5 | 0.1635592 | 0 | 3 | 3 | not sig |
| MKX | R93G | SIN3B | O75182 | 0.571528452 | 0.2935017 | 1.9472744 | 5 | 0.1090501 | 0 | 3 | 2 | not sig |
| SLC6A1 | G360S | NUP210 | Q8TEM1 | 0.570965615 | 0.221156 | 2.5817321 | 14 | 0.0217361 | 0 | 3 | 3 | not sig |
| CHD8 | L834P | NUMB | P49757 | 0.561130514 | 0.8504484 | 0.6598054 | 12 | 0.5218412 | 0 | 3 | 3 | not sig |
| SLC6A1 | A288V | IMMT | Q16891 | 0.561114634 | 0.1809264 | 3.1013421 | 14 | 0.0078115 | 0 | 3 | 3 | not sig |
| CHD8 | C1095Y | AP2A1 | O95782 | 0.556726851 | 0.9726299 | 0.5723933 | 12 | 0.5776222 | 0 | 3 | 3 | not sig |
| SLC6A1 | G360S | UBAC2 | Q8NBM4 | 0.552439715 | 0.1337856 | 4.1292903 | 14 | 0.001022 | 0 | 3 | 3 | not sig |
| SLC6A1 | G299V | IMMT | Q16891 | 0.546976157 | 0.1809264 | 3.0231972 | 14 | 0.0091222 | 0 | 3 | 3 | not sig |
| CHD8 | R1580W | CLTC | Q00610 | 0.54663655 | 0.8366249 | 0.653383 | 12 | 0.5258311 | 0 | 3 | 3 | not sig |
| MYT1L | H522Q | SCOC | Q9UIL1 | 0.54603853 | 0.3381821 | 1.6146286 | 6 | 0.1575183 | 0 | 3 | 3 | not sig |
| KCNQ3 | R230H | NUP155 | O75694 | 0.544334594 | 0.1741688 | 3.1253276 | 10 | 0.0107758 | 0 | 3 | 3 | not sig |
| MKX | R93G | PRPF31 | Q8WWY3 | 0.544156481 | 0.2444547 | 2.2260014 | 1 | 0.2687926 | 0 | 1 | 1 | not sig |
| CHD8 | L834P | KPNA1 | P52294 | 0.541150287 | 0.1018711 | 5.3121106 | 12 | 0.0001847 | 0 | 3 | 3 | not sig |
| CHD8 | C1095Y | AP2A2 | O94973 | 0.540426572 | 0.9054971 | 0.5968286 | 12 | 0.5617133 | 0 | 3 | 3 | not sig |
| PPP2R5D | E198K | MOGS | Q13724 | 0.540219297 | 0.2914516 | 1.853547 | 3 | 0.1608569 | 0 | 3 | 2 | not sig |
| NSD1 | C1877R | MLYCD | O95822 | 0.535340736 | 0.8078051 | 0.6627103 | 4 | 0.5437533 | 0 | 3 | 3 | not sig |
| TBR1 | N374H | KIF7 | Q2M1P5 | 0.528607882 | 0.6496592 | 0.8136695 | 6 | 0.446914 | 0 | 3 | 3 | not sig |
| SLC6A1 | H198P | NUP210 | Q8TEM1 | 0.525292563 | 0.221156 | 2.3752125 | 14 | 0.032368 | 0 | 3 | 3 | not sig |
| MYT1L | H522Q | CTBP1 | Q13363 | 0.522734659 | 0.7264448 | 0.7195793 | 4 | 0.5115843 | 0 | 3 | 1 | not sig |
| MYT1L | C504R | ATAD3B | Q5T9A4 | 0.51581258 | 0.2814149 | 1.8329259 | 6 | 0.1165148 | 0 | 3 | 3 | not sig |
| CHD8 | L834P | TAOK2 | Q9UL54 | 0.514950905 | 0.2536063 | 2.030513 | 11 | 0.0671916 | 0 | 3 | 3 | not sig |
| MYT1L | H522Q | CEP55 | Q53EZ4 | 0.509887932 | 0.3648333 | 1.3975914 | 6 | 0.211728 | 0 | 3 | 3 | not sig |
| CHD8 | R1580W | BRD4 | O60885 | 0.506481873 | 0.704989 | 0.7184252 | 11 | 0.4874747 | 0 | 3 | 3 | not sig |
| CHD8 | L834P | AP2S1 | P53680 | 0.504423769 | 0.9506887 | 0.5305878 | 12 | 0.6053848 | 0 | 3 | 3 | not sig |
| NSD1 | C1350R | MLYCD | O95822 | 0.497416652 | 1.1424089 | 0.4354104 | 4 | 0.6857292 | 0 | 1 | 3 | not sig |
| SLC6A1 | G360S | ESYT1 | Q9BSJ8 | 0.496937065 | 0.1185535 | 4.1916693 | 14 | 0.0009051 | 0 | 3 | 3 | not sig |
| TBR1 | N374H | AP1B1 | Q10567 | 0.494172121 | 0.8127064 | 0.6080574 | 2 | 0.6050021 | 0 | 1 | 3 | not sig |
| CHD8 | L834P | BRD4 | O60885 | 0.489083989 | 0.704989 | 0.693747 | 11 | 0.502228 | 0 | 3 | 3 | not sig |

| | | | | | | | | | | | | |
|---|---|---|---|---|---|---|---|---|---|---|---|---|
| CHD8 | L834P | TRIM27 | P14373 | 0.488266383 | 0.1765667 | 2.765337 | 12 | 0.0171091 | 0 | 3 | 3 | not sig |
| SLC6A1 | G307R | SLC39A14 | Q15043 | 0.487351999 | 0.1388214 | 3.5106399 | 13 | 0.003835 | 0 | 3 | 3 | not sig |
| TRAF7 | R593W | SCRIB | Q14160 | 0.484637549 | 0.1830177 | 2.6480364 | 6 | 0.0381285 | 0 | 3 | 3 | not sig |
| CHD8 | R1580W | GDI2 | P50395 | 0.481670657 | 0.4272981 | 1.1272474 | 10 | 0.2859592 | 0 | 3 | 3 | not sig |
| TRAF7 | R593W | PFDN1 | O60925 | 0.477773444 | 0.2118824 | 2.2548995 | 6 | 0.0650021 | 0 | 3 | 3 | not sig |
| CHD8 | L834P | FXR1 | P51114 | 0.477337982 | 0.2689068 | 1.7751055 | 10 | 0.1062683 | 0 | 3 | 2 | not sig |
| KCNQ3 | R230L | ELOB | Q15370 | 0.474637855 | 0.2108906 | 2.2506351 | 10 | 0.0481277 | 0 | 3 | 3 | not sig |
| CHD8 | C1095Y | NUMB | P49757 | 0.469784705 | 0.8504484 | 0.5523965 | 12 | 0.5908175 | 0 | 3 | 3 | not sig |
| CHD8 | R1580W | KPNA6 | O60684 | 0.467747551 | 0.1163106 | 4.021539 | 12 | 0.0016948 | 0 | 3 | 3 | not sig |
| CHD8 | L834P | CLTC | Q00610 | 0.467245091 | 0.8366249 | 0.5584881 | 12 | 0.5867812 | 0 | 3 | 3 | not sig |
| CREBBP | R413Q | ZNRD2 | O60232 | 0.465905067 | 0.1964087 | 2.3721205 | 3 | 0.0983153 | 0 | 3 | 2 | not sig |
| AP2S1 | R10W | RDH13 | Q8NBN7 | 0.46393415 | 0.440108 | 1.0541371 | 5 | 0.3400715 | 0 | 3 | 2 | not sig |
| MYT1L | H522Q | ATAD3B | Q5T9A4 | 0.462817963 | 0.2814149 | 1.6446109 | 6 | 0.1511533 | 0 | 3 | 3 | not sig |
| VEZF1 | Q209P | PYCR3 | Q53H96 | 0.462350945 | 0.6959114 | 0.6643819 | 4 | 0.5427877 | 0 | 3 | 3 | not sig |
| PPP1R9B | R753H | ALDH2 | P05091 | 0.460947984 | 0.5039458 | 0.9146777 | 2 | 0.4569169 | 0 | 1 | 3 | not sig |
| CHD8 | R1580W | NUMB | P49757 | 0.460698132 | 0.8504484 | 0.541712 | 12 | 0.5979315 | 0 | 3 | 3 | not sig |
| CHD8 | Q696K | AP2M1 | Q96CW1 | 0.458261507 | 0.9235584 | 0.4961911 | 12 | 0.6287232 | 0 | 3 | 3 | not sig |
| CHD8 | R1580W | AP2A1 | O95782 | 0.454117697 | 0.9726299 | 0.4668967 | 12 | 0.648937 | 0 | 3 | 3 | not sig |
| DPYSL2 | R496C | SIRT2 | Q8IXJ6 | 0.453848302 | 0.3589376 | 1.2644212 | 8 | 0.2416708 | 0 | 3 | 3 | not sig |
| CHD8 | L834P | NUP153 | P49790 | 0.45361489 | 0.5825294 | 0.7786987 | 12 | 0.45124 | 0 | 3 | 3 | not sig |
| KCNQ3 | R230H | ELOB | Q15370 | 0.451649993 | 0.2108906 | 2.1416314 | 10 | 0.0578756 | 0 | 3 | 3 | not sig |
| CHD8 | L834P | NDUFAB1 | O14561 | 0.449498514 | 0.4637705 | 0.9692263 | 12 | 0.3515625 | 0 | 3 | 3 | not sig |
| PPP1R9B | R753H | SNX9 | Q9Y5X1 | 0.446931948 | 0.3161491 | 1.4136744 | 4 | 0.2303464 | 0 | 3 | 3 | not sig |
| CHD8 | Q696K | ZBTB11 | O95625 | 0.446826644 | 0.3572072 | 1.2508892 | 8 | 0.2463179 | 0 | 2 | 2 | not sig |
| PPP1R9B | R753H | MAP7D3 | Q8IWC1 | 0.445263997 | 0.2672132 | 1.6663247 | 4 | 0.1709774 | 0 | 3 | 3 | not sig |
| CHD8 | R1580W | AP2B1 | P63010 | 0.445143981 | 0.9105977 | 0.4888482 | 12 | 0.6337615 | 0 | 3 | 3 | not sig |
| CHD8 | Q696K | AP2A1 | O95782 | 0.433170135 | 0.9726299 | 0.4453597 | 12 | 0.6639877 | 0 | 3 | 3 | not sig |
| AP2S1 | R10W | AP1B1 | Q10567 | 0.430581989 | 0.6689385 | 0.6436795 | 5 | 0.5481405 | 0 | 3 | 2 | not sig |
| STXBP1 | R551C | STXBP2 | Q15833 | 0.425441563 | 1.2317404 | 0.3453987 | 6 | 0.7415819 | 0 | 3 | 3 | not sig |
| MKX | R93G | NCOR1 | O75376 | 0.424521141 | 0.4242599 | 1.0006157 | 5 | 0.362947 | 0 | 3 | 2 | not sig |
| AP2S1 | R10W | RNF41 | Q9H4P4 | 0.42396224 | 0.7126674 | 0.594895 | 5 | 0.5778118 | 0 | 3 | 2 | not sig |
| CHD8 | Q696K | BRD4 | O60885 | 0.423701285 | 0.704989 | 0.6010041 | 11 | 0.5600177 | 0 | 3 | 3 | not sig |
| PPP1R9B | R753H | BEX3 | Q00994 | 0.422333051 | 0.1491462 | 2.8316709 | 4 | 0.0472649 | 0 | 3 | 3 | not sig |
| SLC6A1 | L54F | YIPF3 | Q9GZM5 | 0.421657633 | 0.2375487 | 1.7750365 | 14 | 0.0976275 | 0 | 3 | 3 | not sig |
| MKX | L89F | NAP1L4 | Q99733 | 0.418516239 | 0.3031999 | 1.3803312 | 5 | 0.2260063 | 0 | 3 | 2 | not sig |
| SLC6A1 | L54F | DHCR24 | Q15392 | 0.414717337 | 0.2596211 | 1.5973947 | 14 | 0.1324963 | 0 | 3 | 3 | not sig |
| MYT1L | H522Q | PCM1 | Q15154 | 0.410933126 | 0.0893653 | 4.5983529 | 6 | 0.0036984 | 0 | 3 | 3 | not sig |
| SLC6A1 | H198P | UBAC2 | Q8NBM4 | 0.41069225 | 0.1337856 | 3.0697784 | 14 | 0.0083168 | 0 | 3 | 3 | not sig |
| CHD8 | M904I | AP2M1 | Q96CW1 | 0.407837855 | 0.9235584 | 0.441594 | 12 | 0.6666353 | 0 | 3 | 3 | not sig |
| RFX3 | A508E | IRF2BPL | Q9H1B7 | 0.407690379 | 0.3069056 | 1.32839 | 4 | 0.2547709 | 0 | 3 | 3 | not sig |
| TRAF7 | R593W | NUDC | Q9Y266 | 0.406118869 | 0.1898883 | 2.1387255 | 6 | 0.0762845 | 0 | 3 | 3 | not sig |
| CHD8 | C1095Y | CRLF3 | Q8IUI8 | 0.404567899 | 0.199416 | 2.028763 | 11 | 0.067394 | 0 | 3 | 2 | not sig |
| CHD8 | R1580W | KPNA1 | P52294 | 0.403723609 | 0.1018711 | 3.9630847 | 12 | 0.0018828 | 0 | 3 | 3 | not sig |
| CHD8 | R1580W | AP2A2 | O94973 | 0.403615758 | 0.9054971 | 0.4457395 | 12 | 0.663721 | 0 | 3 | 3 | not sig |
| CHD8 | R1580W | SNRNP200 | O75643 | 0.40307316 | 0.1812241 | 2.2241692 | 12 | 0.0460917 | 0 | 3 | 3 | not sig |
| CHD8 | C1095Y | TAOK2 | Q9UL54 | 0.401371431 | 0.2536063 | 1.5826556 | 11 | 0.1418064 | 0 | 3 | 3 | not sig |
| AP2S1 | G64D | YME1L1 | Q96TA2 | 0.400163274 | 0.7387788 | 0.5416551 | 3 | 0.625696 | 0 | 1 | 2 | not sig |
| SLC6A1 | G307R | POR | P16435 | 0.399945457 | 0.1728276 | 2.31413 | 14 | 0.0363621 | 0 | 3 | 3 | not sig |
| SLC6A1 | L54F | CERS2 | Q96G23 | 0.396515021 | 0.2014027 | 1.9687672 | 14 | 0.0691033 | 0 | 3 | 3 | not sig |
| SLC6A1 | L54F | IMMT | Q16891 | 0.39577027 | 0.1809264 | 2.1874656 | 14 | 0.0461716 | 0 | 3 | 3 | not sig |
| TBR1 | K228E | FBXO3 | Q9UK99 | 0.393843617 | 0.238537 | 1.6510795 | 6 | 0.1498129 | 0 | 3 | 3 | not sig |
| GRIA2 | E776D | CKAP4 | Q07065 | 0.392220558 | 0.2393151 | 1.6389292 | 4 | 0.1765708 | 0 | 3 | 3 | not sig |
| CHD8 | M904I | CRLF3 | Q8IUI8 | 0.391602036 | 0.199416 | 1.9637438 | 11 | 0.0753318 | 0 | 3 | 2 | not sig |
| CHD8 | Q696K | NUMB | P49757 | 0.391024571 | 0.8504484 | 0.4597863 | 12 | 0.6538885 | 0 | 3 | 3 | not sig |
| CHD8 | Q696K | AP2B1 | P63010 | 0.388010457 | 0.9105977 | 0.4261053 | 12 | 0.6775744 | 0 | 3 | 3 | not sig |
| CHD8 | Q696K | COPS3 | Q9UNS2 | 0.387685638 | 0.3250456 | 1.1927115 | 6 | 0.2780058 | 0 | 2 | 3 | not sig |
| PPP1R9B | R753H | CKAP5 | Q14008 | 0.38652198 | 0.1702695 | 2.2700606 | 4 | 0.0857219 | 0 | 3 | 3 | not sig |
| SLC6A1 | L54F | RHOT2 | Q8IXI1 | 0.376853847 | 0.1586902 | 2.3747766 | 14 | 0.0323949 | 0 | 3 | 3 | not sig |
| SLC6A1 | H198P | MYH9 | P35579 | 0.375198945 | 0.3279728 | 1.1439939 | 14 | 0.27181 | 0 | 3 | 3 | not sig |
| SLC6A1 | L54F | AUP1 | Q9Y679 | 0.373281087 | 0.1326649 | 2.8137146 | 14 | 0.0138026 | 0 | 3 | 3 | not sig |
| PPP1R9B | R753H | MCM10 | Q7L590 | 0.372464547 | 0.3081578 | 1.2086813 | 4 | 0.2933473 | 0 | 3 | 3 | not sig |
| CHD8 | Q696K | BRD2 | P25440 | 0.372205707 | 0.5976791 | 0.6227518 | 12 | 0.5451008 | 0 | 3 | 3 | not sig |
| CHD8 | R1580W | AP2S1 | P53680 | 0.372190348 | 0.9506887 | 0.3914955 | 12 | 0.702294 | 0 | 3 | 3 | not sig |
| CHD8 | Q696K | CRLF3 | Q8IUI8 | 0.370041188 | 0.199416 | 1.8556239 | 11 | 0.090478 | 0 | 3 | 2 | not sig |
| MKX | R93G | IRF2BP1 | Q8IU81 | 0.366782668 | 0.4881738 | 0.7513362 | 5 | 0.4862878 | 0 | 3 | 2 | not sig |
| CHD8 | R1580W | NDUFAB1 | O14561 | 0.366492487 | 0.4637705 | 0.7902454 | 12 | 0.4447209 | 0 | 3 | 3 | not sig |
| CHD8 | C1095Y | AP2S1 | P53680 | 0.360976408 | 0.9506887 | 0.3796999 | 12 | 0.7108026 | 0 | 3 | 3 | not sig |
| TRAF7 | R593W | PFDN5 | Q99471 | 0.356652064 | 0.1218812 | 2.9262263 | 6 | 0.0264164 | 0 | 3 | 3 | not sig |
| PTK7 | R562Q | ERGIC2 | Q96RQ1 | 0.355423435 | 0.4351674 | 0.8167511 | 4 | 0.4599209 | 0 | 3 | 3 | not sig |
| TBR1 | N374H | ELP1 | O95163 | 0.351241513 | 0 | 0 | 0 | 0 | 0 | 1 | 1 | not sig |
| KCNQ3 | R230H | CAND2 | O75155 | 0.351225833 | 0.1872798 | 1.8754067 | 10 | 0.0902027 | 0 | 3 | 3 | not sig |
| CHD8 | C1095Y | FXR1 | P51114 | 0.351222639 | 0.2689068 | 1.3061128 | 10 | 0.220758 | 0 | 3 | 2 | not sig |

| | | | | | | | | | | | | |
|---|---|---|---|---|---|---|---|---|---|---|---|---|
| SLC6A1 | L54F | CERS1 | P27544 | 0.350673948 | 0.2321979 | 1.5102374 | 14 | 0.15322 | 0 | 3 | 3 | not sig |
| CHD8 | C1095Y | BRD4 | O60885 | 0.350020109 | 0.704989 | 0.4964902 | 11 | 0.6293207 | 0 | 3 | 3 | not sig |
| CHD8 | L834P | ZBTB11 | O95625 | 0.346742139 | 0.3572072 | 0.9707031 | 8 | 0.3601257 | 0 | 2 | 2 | not sig |
| CHD8 | R1580W | HUWE1 | Q7Z6Z7 | 0.345272684 | 0.2091506 | 1.6508331 | 12 | 0.1246812 | 0 | 3 | 3 | not sig |
| MKX | L89F | SIN3B | O75182 | 0.344323115 | 0.2935017 | 1.1731552 | 5 | 0.2935734 | 0 | 3 | 2 | not sig |
| SLC6A1 | G307R | YIF1B | Q5BJH7 | 0.33935149 | 0.3358985 | 1.01028 | 11 | 0.334068 | 0 | 2 | 2 | not sig |
| CREBBP | R413Q | RB1 | P06400 | 0.338081621 | 0.4036431 | 0.8375757 | 3 | 0.4637458 | 0 | 3 | 2 | not sig |
| CHD8 | Q696K | AP2A2 | O94973 | 0.337819792 | 0.9054971 | 0.3730766 | 12 | 0.7155982 | 0 | 3 | 3 | not sig |
| CHD8 | R1580W | KPNA3 | O00505 | 0.328911292 | 0.2281738 | 1.4414947 | 12 | 0.1750287 | 0 | 3 | 3 | not sig |
| RFX3 | A508E | ARL6IP5 | O75915 | 0.328376816 | 0.1676996 | 1.9581248 | 4 | 0.1218163 | 0 | 3 | 3 | not sig |
| MYT1L | C504R | PCM1 | Q15154 | 0.322205612 | 0.0893653 | 3.6054896 | 6 | 0.0112919 | 0 | 3 | 3 | not sig |
| CHD8 | L834P | SNRNP200 | O75643 | 0.322069943 | 0.1812241 | 1.7771912 | 12 | 0.1008688 | 0 | 3 | 3 | not sig |
| SLC6A1 | L54F | TM9SF3 | Q9HD45 | 0.321403527 | 0.2177489 | 1.4760283 | 14 | 0.162072 | 0 | 3 | 3 | not sig |
| SLC6A1 | L54F | RFT1 | Q96AA3 | 0.320277601 | 0.1767227 | 1.8123175 | 14 | 0.0914337 | 0 | 3 | 3 | not sig |
| RORB | Y22D | MATR3 | P43243 | 0.319503371 | 0.0720532 | 4.4342718 | 4 | 0.0113843 | 0 | 3 | 3 | not sig |
| TBR1 | N374H | IDE | P14735 | 0.318034273 | 1.0208507 | 0.3115385 | 6 | 0.7659323 | 0 | 3 | 3 | not sig |
| TBL1XR1 | L282P | NCOR2 | Q9Y618 | 0.3148516 | 0.2065602 | 1.5242604 | 4 | 0.2021238 | 0 | 3 | 3 | not sig |
| CHD8 | L834P | GDI2 | P50395 | 0.309361074 | 0.6042908 | 0.5119408 | 10 | 0.6198073 | 0 | 1 | 3 | not sig |
| TBR1 | K228E | ELP1 | O95163 | 0.309019913 | 0 | 0 | 0 | 0 | 0 | 1 | 1 | not sig |
| CHD8 | C1095Y | KPNA6 | O60684 | 0.306374131 | 0.1163106 | 2.6341036 | 12 | 0.0218099 | 0 | 3 | 3 | not sig |
| SLC6A1 | G360S | MYH9 | P35579 | 0.301398189 | 0.3279728 | 0.918973 | 14 | 0.373672 | 0 | 3 | 3 | not sig |
| CHD8 | M904I | AP2B1 | P63010 | 0.300315616 | 0.9105977 | 0.3298006 | 12 | 0.7472365 | 0 | 3 | 3 | not sig |
| TBL1XR1 | L282P | NCOR1 | O75376 | 0.296277107 | 0.1947069 | 1.5216572 | 4 | 0.2027462 | 0 | 3 | 3 | not sig |
| CHD8 | M904I | NDUFAB1 | O14561 | 0.29244395 | 0.4637705 | 0.6305791 | 12 | 0.5401393 | 0 | 3 | 3 | not sig |
| FOXP1 | R513H | CEP55 | Q53EZ4 | 0.290127192 | 0.9170501 | 0.3163701 | 4 | 0.7675436 | 0 | 1 | 2 | not sig |
| SLC6A1 | L54F | CHP1 | Q99653 | 0.290117756 | 0.1798068 | 1.6134974 | 14 | 0.1289427 | 0 | 3 | 3 | not sig |
| CHD8 | L834P | CRLF3 | Q8IUI8 | 0.287245608 | 0.199416 | 1.4404338 | 11 | 0.1775951 | 0 | 3 | 2 | not sig |
| TBR1 | N374H | ATAD3B | Q5T9A4 | 0.286234244 | 0.8334873 | 0.3434177 | 6 | 0.7429979 | 0 | 3 | 3 | not sig |
| TBR1 | N374H | BAIAP2 | Q9UQB8 | 0.28526063 | 1.295009 | 0.2202769 | 1 | 0.8619716 | 0 | 1 | 2 | not sig |
| KCNQ3 | R230H | CLPTM1 | O96005 | 0.282887472 | 0.1968038 | 1.4374084 | 6 | 0.2006369 | 0 | 3 | 3 | not sig |
| CHD8 | Q696K | AP2S1 | P53680 | 0.2819477 | 0.9506887 | 0.2965721 | 12 | 0.7718637 | 0 | 3 | 3 | not sig |
| MKX | R93G | HSPA6 | P17066 | 0.281159365 | 0.4979344 | 0.5646515 | 5 | 0.5967011 | 0 | 3 | 2 | not sig |
| CHD8 | C1095Y | NDUFAB1 | O14561 | 0.279834832 | 0.4637705 | 0.6033908 | 12 | 0.557482 | 0 | 3 | 3 | not sig |
| CHD8 | C1095Y | LYRM4 | Q9HD34 | 0.278914229 | 0.5579203 | 0.4999177 | 12 | 0.6261737 | 0 | 3 | 3 | not sig |
| FOXP2 | R570C | CTBP2 | P56545 | 0.275365686 | 0.6569635 | 0.4191491 | 4 | 0.6966374 | 0 | 3 | 3 | not sig |
| CHD8 | Q696K | FXR1 | P51114 | 0.266211077 | 0.2689068 | 0.9899752 | 10 | 0.345535 | 0 | 3 | 2 | not sig |
| TBR1 | N374H | ZMYM2 | Q9UBW7 | 0.26331231 | 0.709013 | 0.3713787 | 6 | 0.7231166 | 0 | 3 | 3 | not sig |
| CHD8 | R1580W | KPNA5 | O15131 | 0.260180357 | 0.1222152 | 2.1288703 | 12 | 0.0546648 | 0 | 3 | 3 | not sig |
| FOXP1 | R513C | FHL2 | Q14192 | 0.253257706 | 0.399445 | 0.634024 | 7 | 0.5462174 | 0 | 3 | 2 | not sig |
| MKX | L89F | IRF2BP1 | Q8IU81 | 0.252719 | 0.4881738 | 0.5176824 | 5 | 0.6267611 | 0 | 3 | 2 | not sig |
| NSD1 | C1877R | NUMA1 | Q14980 | 0.251146533 | 0.5229604 | 0.4802401 | 6 | 0.6480637 | 0 | 3 | 3 | not sig |
| CHD8 | L834P | HUWE1 | Q7Z6Z7 | 0.250055781 | 0.2091506 | 1.1955778 | 12 | 0.2549533 | 0 | 3 | 3 | not sig |
| RFX3 | A508E | RAE1 | P78406 | 0.247696816 | 0.4940426 | 0.5013674 | 4 | 0.642449 | 0 | 3 | 3 | not sig |
| CHD8 | L834P | KPNA5 | O15131 | 0.247612596 | 0.1222152 | 2.0260373 | 12 | 0.0655844 | 0 | 3 | 3 | not sig |
| SLC6A1 | L54F | ESYT1 | Q9BSJ8 | 0.24565482 | 0.1185535 | 2.072101 | 14 | 0.0572054 | 0 | 3 | 3 | not sig |
| NSD1 | C1877R | PDCD11 | Q14690 | 0.243217772 | 0.3621618 | 0.6715722 | 6 | 0.5268526 | 0 | 3 | 3 | not sig |
| KCNQ3 | R230L | ARL10 | Q8N8L6 | 0.238892436 | 0.2217619 | 1.0772473 | 10 | 0.3066703 | 0 | 3 | 3 | not sig |
| SLC6A1 | L54F | UBAC2 | Q8NBM4 | 0.236671151 | 0.1337856 | 1.7690326 | 14 | 0.098659 | 0 | 3 | 3 | not sig |
| AP2S1 | R10W | GPX8 | Q8TED1 | 0.234866654 | 0.4494857 | 0.5225231 | 3 | 0.6374389 | 0 | 3 | 2 | not sig |
| GRIA2 | E776D | UFSP2 | Q9NUQ7 | 0.234088851 | 0.3580234 | 0.6538367 | 4 | 0.5488995 | 0 | 3 | 3 | not sig |
| TBR1 | N374H | COP1 | Q8NHY2 | 0.231297855 | 0.5002169 | 0.4623952 | 6 | 0.6600912 | 0 | 3 | 3 | not sig |
| CHD8 | R1580W | XPO5 | Q9HAV4 | 0.230610998 | 0.2355528 | 0.9790205 | 12 | 0.3468955 | 0 | 3 | 3 | not sig |
| TRAF7 | R655Q | PFDN1 | O60925 | 0.230606225 | 0.2118824 | 1.0883691 | 6 | 0.3182058 | 0 | 3 | 3 | not sig |
| MYT1L | H522Q | KDM1A | O60341 | 0.229066072 | 0.3785711 | 0.6050807 | 6 | 0.56729 | 0 | 3 | 3 | not sig |
| CHD8 | C1095Y | KPNA1 | P52294 | 0.224338723 | 0.1018711 | 2.2021832 | 12 | 0.0479486 | 0 | 3 | 3 | not sig |
| CHD8 | M904I | AP2A1 | O95782 | 0.219569705 | 0.9726299 | 0.2257485 | 12 | 0.8251962 | 0 | 3 | 3 | not sig |
| CHD8 | M904I | TAOK2 | Q9UL54 | 0.218948385 | 0.2536063 | 0.8633397 | 11 | 0.4063814 | 0 | 3 | 3 | not sig |
| CHD8 | M904I | FXR1 | P51114 | 0.218262416 | 0.2945727 | 0.7409459 | 10 | 0.4757651 | 0 | 2 | 2 | not sig |
| CHD8 | M904I | CLTC | Q00610 | 0.217419028 | 0.8366249 | 0.2598763 | 12 | 0.7993643 | 0 | 3 | 3 | not sig |
| CHD8 | L834P | KEAP1 | Q14145 | 0.21619903 | 0.2821492 | 0.7662578 | 12 | 0.4583318 | 0 | 3 | 3 | not sig |
| CHD8 | Q696K | CLTC | Q00610 | 0.214685458 | 0.8366249 | 0.256609 | 12 | 0.8018273 | 0 | 3 | 3 | not sig |
| AP2S1 | G64D | HSPA6 | P17066 | 0.211651784 | 0.5566592 | 0.3802179 | 5 | 0.7193998 | 0 | 3 | 2 | not sig |
| SLC6A1 | L54F | POR | P16435 | 0.210840648 | 0.1728276 | 1.219948 | 14 | 0.2426374 | 0 | 3 | 3 | not sig |
| NSD1 | C1877R | PRPF6 | O94906 | 0.210521545 | 0.5438891 | 0.387067 | 1 | 0.7648914 | 0 | 2 | 1 | not sig |
| CHD8 | R1580W | OAT | P04181 | 0.210388961 | 0.3848641 | 0.5466578 | 11 | 0.5955286 | 0 | 3 | 3 | not sig |
| DPYSL2 | H333N | SIRT2 | Q8IXJ6 | 0.207446083 | 0.3589376 | 0.5779447 | 8 | 0.5792015 | 0 | 3 | 3 | not sig |
| SLC6A1 | G307R | CDS2 | O95674 | 0.20673991 | 0.2076177 | 0.9957722 | 14 | 0.3362612 | 0 | 3 | 3 | not sig |
| TBR1 | K228E | COP1 | Q8NHY2 | 0.206436317 | 0.5002169 | 0.4126936 | 6 | 0.6941725 | 0 | 3 | 3 | not sig |
| KCNMA1 | A1033P | EXD2 | Q9NVH0 | 0.205978485 | 0.2682314 | 0.7679135 | 8 | 0.464599 | 0 | 3 | 3 | not sig |
| KCNQ3 | R230H | CLPB | Q9H078 | 0.20426257 | 0.1558742 | 1.3104322 | 10 | 0.2193482 | 0 | 3 | 3 | not sig |
| SLC6A1 | L54F | MYH9 | P35579 | 0.19989053 | 0.3279728 | 0.6094728 | 14 | 0.5519746 | 0 | 3 | 3 | not sig |
| CREBBP | R413Q | CRTAP | O75718 | 0.197617664 | 0.4590256 | 0.4305156 | 3 | 0.695876 | 0 | 3 | 2 | not sig |

| | | | | | | | | | | | | |
|---|---|---|---|---|---|---|---|---|---|---|---|---|
| AP2S1 | R10W | CKAP4 | Q07065 | 0.195425087 | 0.2890255 | 0.6761517 | 3 | 0.5474045 | 0 | 3 | 2 | not sig |
| RFX3 | A508E | SRPRA | P08240 | 0.191598128 | 0.198009 | 0.9676235 | 4 | 0.388027 | 0 | 3 | 3 | not sig |
| CHD8 | M904I | NUMB | P49757 | 0.190578443 | 0.8504484 | 0.2240917 | 12 | 0.8264559 | 0 | 3 | 3 | not sig |
| MYT1L | H522Q | GRAMD4 | Q6IC98 | 0.183702101 | 0.5287773 | 0.3474092 | 5 | 0.7424298 | 0 | 3 | 2 | not sig |
| MYT1L | C504R | CEP55 | Q53EZ4 | 0.18139298 | 0.3648333 | 0.4971941 | 6 | 0.6367433 | 0 | 3 | 3 | not sig |
| CHD8 | M904I | BRD3 | Q15059 | 0.180769233 | 1.1163786 | 0.1619247 | 12 | 0.8740597 | 0 | 3 | 3 | not sig |
| MKX | L89F | PRPF31 | Q8WWY3 | 0.17731186 | 0.211704 | 0.8375462 | 1 | 0.5561363 | 0 | 2 | 1 | not sig |
| CHD8 | L834P | NFS1 | Q9Y697 | 0.174303211 | 0.4975548 | 0.3503196 | 12 | 0.7321711 | 0 | 3 | 3 | not sig |
| SLC6A1 | L54F | NUP155 | O75694 | 0.172216832 | 0.1733885 | 0.9932425 | 14 | 0.3374495 | 0 | 3 | 3 | not sig |
| TBR1 | N374H | TCAF1 | Q9Y4C2 | 0.171791117 | 0.242007 | 0.7098601 | 1 | 0.6070064 | 0 | 1 | 2 | not sig |
| CHD8 | R1580W | CRLF3 | Q8IUI8 | 0.171202754 | 0.199416 | 0.8585204 | 11 | 0.4089232 | 0 | 3 | 2 | not sig |
| CHD8 | M904I | NUP153 | P49790 | 0.164023712 | 0.5825294 | 0.2815715 | 12 | 0.7830688 | 0 | 3 | 3 | not sig |
| TBR1 | N374H | PITRM1 | Q5JRX3 | 0.16262225 | 2.116517 | 0.0768348 | 6 | 0.941253 | 0 | 3 | 3 | not sig |
| MKX | L89F | HSPA6 | P17066 | 0.15959638 | 0.4979344 | 0.3205169 | 5 | 0.7615387 | 0 | 3 | 2 | not sig |
| GRIA2 | E776D | GRIA4 | P48058 | 0.158427188 | 0.3485584 | 0.4545212 | 4 | 0.6730269 | 0 | 3 | 3 | not sig |
| CHD8 | L834P | KPNA3 | O00505 | 0.157890336 | 0.2281738 | 0.691974 | 12 | 0.5021236 | 0 | 3 | 3 | not sig |
| PTK7 | R562Q | MBOAT7 | Q96N66 | 0.15780969 | 0.1817721 | 0.8681732 | 3 | 0.4491754 | 0 | 3 | 2 | not sig |
| PPP1R9B | R753H | ACTN1 | P12814 | 0.157682074 | 0.5316574 | 0.2965859 | 4 | 0.781545 | 0 | 3 | 3 | not sig |
| CHD8 | C1095Y | BRD2 | P25440 | 0.155544379 | 0.5976791 | 0.2602473 | 12 | 0.7990848 | 0 | 3 | 3 | not sig |
| CHD8 | L834P | EPS15L1 | Q9UBC2 | 0.154088174 | 1.4746707 | 0.1044899 | 8 | 0.9193528 | 0 | 2 | 1 | not sig |
| KCNQ3 | R230H | FANCI | Q9NVI1 | 0.1537367 | 0.4168111 | 0.3688402 | 7 | 0.7231499 | 0 | 1 | 3 | not sig |
| KCNMA1 | A1033P | FUNDC2 | Q9BWH2 | 0.153548677 | 0.2665762 | 0.576003 | 8 | 0.5804515 | 0 | 3 | 3 | not sig |
| SLC6A1 | L54F | MOGS | Q13724 | 0.152907761 | 0.2829625 | 0.5403817 | 14 | 0.5974263 | 0 | 3 | 3 | not sig |
| TBR1 | N374H | FBXW11 | Q9UKB1 | 0.145192832 | 1.1222771 | 0.1293734 | 6 | 0.9012902 | 0 | 3 | 3 | not sig |
| SLC6A1 | G307R | NUP155 | O75694 | 0.144285946 | 0.1733885 | 0.832154 | 14 | 0.4192908 | 0 | 3 | 3 | not sig |
| SLC6A1 | G307R | RABL3 | Q5HYI8 | 0.139976784 | 0.1988697 | 0.7038617 | 14 | 0.4930606 | 0 | 3 | 3 | not sig |
| CHD8 | M904I | KPNA3 | O00505 | 0.138774692 | 0.2281738 | 0.6081974 | 12 | 0.5543938 | 0 | 3 | 3 | not sig |
| CHD8 | M904I | AP2A2 | O94973 | 0.138723397 | 0.9054971 | 0.1532014 | 12 | 0.8807855 | 0 | 3 | 3 | not sig |
| CHD8 | R1580W | ZBTB11 | O95625 | 0.138373667 | 0.3260841 | 0.4243497 | 8 | 0.6824921 | 0 | 3 | 2 | not sig |
| TBR1 | N374H | FBXO3 | Q9UK99 | 0.136882926 | 0.238537 | 0.5738435 | 6 | 0.5869243 | 0 | 3 | 3 | not sig |
| SLC6A1 | A288V | MYH9 | P35579 | 0.136864191 | 0.3279728 | 0.4173034 | 14 | 0.6827852 | 0 | 3 | 3 | not sig |
| SLC6A1 | G360S | LMBR1 | Q8WVP7 | 0.135577645 | 0.5228505 | 0.2593048 | 11 | 0.8001888 | 0 | 3 | 2 | not sig |
| PPP1R9B | R753H | SNX33 | Q8WV41 | 0.135236886 | 0.3911273 | 0.3457619 | 4 | 0.746941 | 0 | 3 | 3 | not sig |
| CHD8 | R1580W | PTCD1 | O75127 | 0.133967104 | 0.3824447 | 0.3502914 | 10 | 0.7333862 | 0 | 3 | 3 | not sig |
| PPP1R9B | R753H | TRAF4 | Q9BUZ4 | 0.133957124 | 0.1225751 | 1.0928574 | 4 | 0.3358609 | 0 | 3 | 3 | not sig |
| SLC6A1 | G307R | YIPF3 | Q9GZM5 | 0.13354626 | 0.2375487 | 0.5621847 | 14 | 0.58288 | 0 | 3 | 3 | not sig |
| CHD8 | M904I | GDI2 | P50395 | 0.128005157 | 0.4272981 | 0.2995688 | 10 | 0.7706401 | 0 | 3 | 3 | not sig |
| TBL1XR1 | L282P | HDAC3 | O15379 | 0.126968027 | 0.2001026 | 0.6345146 | 4 | 0.560222 | 0 | 3 | 3 | not sig |
| KCNQ3 | R230L | NAA15 | Q9BXJ9 | 0.122759744 | 0.2555836 | 0.4803115 | 9 | 0.6424651 | 0 | 3 | 3 | not sig |
| CHD8 | Q696K | COPS6 | Q7L5N1 | 0.122272179 | 0.1984548 | 0.6161209 | 11 | 0.5503515 | 0 | 3 | 3 | not sig |
| CHD8 | Q696K | KPNA6 | O60684 | 0.121298573 | 0.1163106 | 1.0428851 | 12 | 0.3175567 | 0 | 3 | 3 | not sig |
| PPP1R9B | R753H | ACTN4 | O43707 | 0.120866265 | 0.4900888 | 0.2466211 | 4 | 0.8173411 | 0 | 3 | 3 | not sig |
| CHD8 | C1095Y | NUP153 | P49790 | 0.117351671 | 0.5825294 | 0.2014519 | 12 | 0.8437197 | 0 | 3 | 3 | not sig |
| DNMT3A | L508P | KPNA4 | O00629 | 0.115781506 | 0.3804885 | 0.304297 | 4 | 0.7760755 | 0 | 3 | 3 | not sig |
| TBR1 | N374H | WDR6 | Q9NNW5 | 0.113147314 | 0.4649536 | 0.2433518 | 6 | 0.8158393 | 0 | 3 | 3 | not sig |
| CHD8 | R1580W | KEAP1 | Q14145 | 0.11263179 | 0.2821492 | 0.3991923 | 12 | 0.6967645 | 0 | 3 | 3 | not sig |
| CHD8 | C1095Y | NFS1 | Q9Y697 | 0.110968486 | 0.4975548 | 0.2230277 | 12 | 0.8272652 | 0 | 3 | 3 | not sig |
| NSD1 | C1877R | PUM3 | Q15397 | 0.110443475 | 0.2134578 | 0.5174019 | 5 | 0.6269432 | 0 | 3 | 3 | not sig |
| CHD8 | M904I | AP2S1 | P53680 | 0.109264345 | 0.9506887 | 0.1149318 | 12 | 0.9103999 | 0 | 3 | 3 | not sig |
| CHD8 | M904I | COPS6 | Q7L5N1 | 0.109129955 | 0.1984548 | 0.5498982 | 11 | 0.5933787 | 0 | 3 | 3 | not sig |
| CREBBP | R413Q | P3H1 | Q32P28 | 0.107567861 | 0.210537 | 0.5109213 | 4 | 0.6363136 | 0 | 3 | 3 | not sig |
| GRIA2 | E776D | ODR4 | Q5SWX8 | 0.107288294 | 0.2050188 | 0.5233095 | 4 | 0.6284104 | 0 | 3 | 3 | not sig |
| MYT1L | C504R | PHF21A | Q96BD5 | 0.105262098 | 0.6022643 | 0.1747772 | 6 | 0.8670033 | 0 | 3 | 3 | not sig |
| CHD8 | M904I | SNRNP200 | O75643 | 0.100818635 | 0.1812241 | 0.5563201 | 12 | 0.5882161 | 0 | 3 | 3 | not sig |
| RORB | Y22D | NCOA2 | Q15596 | 0.100489658 | 0.2852718 | 0.3522594 | 4 | 0.74242 | 0 | 3 | 3 | not sig |
| SLC6A1 | G307R | TM9SF3 | Q9HD45 | 0.099604828 | 0.2177489 | 0.4574298 | 14 | 0.6543802 | 0 | 3 | 3 | not sig |
| CHD8 | R1580W | BRD2 | P25440 | 0.09703888 | 0.5976791 | 0.1623595 | 12 | 0.8737246 | 0 | 3 | 3 | not sig |
| CHD8 | L834P | CSNK2A2 | P19784 | 0.095662546 | 0.2201047 | 0.434623 | 12 | 0.671549 | 0 | 3 | 3 | not sig |
| CHD8 | M904I | KPNA6 | O60684 | 0.094895303 | 0.1163106 | 0.8158785 | 12 | 0.4304673 | 0 | 3 | 3 | not sig |
| CHD8 | L834P | BRD2 | P25440 | 0.094341899 | 0.5976791 | 0.1578471 | 12 | 0.8772023 | 0 | 3 | 3 | not sig |
| CHD8 | Q696K | KPNA1 | P52294 | 0.093912756 | 0.1018711 | 0.9218787 | 12 | 0.3747548 | 0 | 3 | 3 | not sig |
| TBR1 | K228E | OCRL | Q01968 | 0.089967604 | 0.3690435 | 0.2437859 | 5 | 0.8170835 | 0 | 2 | 3 | not sig |
| TRAF7 | R655Q | BAG6 | P46379 | 0.089907523 | 0.1378852 | 0.6520462 | 6 | 0.5385325 | 0 | 3 | 3 | not sig |
| CHD8 | C1095Y | KEAP1 | Q14145 | 0.089331336 | 0.2821492 | 0.3166103 | 12 | 0.7569794 | 0 | 3 | 3 | not sig |
| NSD1 | C1877R | DDX51 | Q8N8A6 | 0.088015741 | 0.5242906 | 0.1678759 | 6 | 0.8721966 | 0 | 3 | 3 | not sig |
| AP2S1 | R10W | HSPA6 | P17066 | 0.087239031 | 0.5566592 | 0.1567189 | 5 | 0.8815979 | 0 | 3 | 2 | not sig |
| SLC6A1 | G307R | RHOT2 | Q8IXI1 | 0.086687741 | 0.1586902 | 0.5462702 | 14 | 0.5934796 | 0 | 3 | 3 | not sig |
| CHD8 | C1095Y | KPNA3 | O00505 | 0.086227598 | 0.2281738 | 0.3779032 | 12 | 0.7121022 | 0 | 3 | 3 | not sig |
| CHD8 | M904I | CWC27 | Q6UX04 | 0.084762565 | 0.8586502 | 0.0987161 | 11 | 0.9231397 | 0 | 3 | 3 | not sig |
| SLC6A1 | G307R | RFT1 | Q96AA3 | 0.08382312 | 0.1767227 | 0.4743201 | 14 | 0.6425847 | 0 | 3 | 3 | not sig |
| TBR1 | N374H | OCRL | Q01968 | 0.083811044 | 0.3300826 | 0.2539093 | 5 | 0.8096763 | 0 | 3 | 3 | not sig |
| TBL1XR1 | L282P | TBL1X | O60907 | 0.082557495 | 0.2584684 | 0.3194105 | 4 | 0.7654012 | 0 | 3 | 3 | not sig |

| | | | | | | | | | | | | |
|---|---|---|---|---|---|---|---|---|---|---|---|---|
| SLC6A1 | L54F | RABL3 | Q5HYI8 | 0.079414195 | 0.1988697 | 0.3993277 | 14 | 0.695677 | 0 | 3 | 3 | not sig |
| CHD8 | M904I | KPNA1 | P52294 | 0.075146453 | 0.1018711 | 0.7376625 | 12 | 0.4748977 | 0 | 3 | 3 | not sig |
| RFX3 | A508E | NUP155 | O75694 | 0.074006166 | 0.0889182 | 0.8322948 | 3 | 0.4663018 | 0 | 2 | 3 | not sig |
| NSD1 | C1877R | PELP1 | Q8IZL8 | 0.072866905 | 0.663122 | 0.1098846 | 6 | 0.9160841 | 0 | 3 | 3 | not sig |
| TBR1 | K228E | TCAF1 | Q9Y4C2 | 0.071130104 | 0.242007 | 0.2939175 | 1 | 0.8180112 | 0 | 1 | 2 | not sig |
| LRRC4C | I163N | IPO9 | Q96P70 | 0.069974899 | 0.2142381 | 0.3266222 | 4 | 0.7603296 | 0 | 3 | 3 | not sig |
| CHD8 | Q696K | KPNA3 | O00505 | 0.068467312 | 0.2281738 | 0.3000665 | 12 | 0.769261 | 0 | 3 | 3 | not sig |
| PPP1R9B | R753H | ADD1 | P35611 | 0.06843536 | 0.7244524 | 0.094465 | 4 | 0.9292827 | 0 | 3 | 3 | not sig |
| KCNQ3 | R230C | PIP4P2 | Q8N4L2 | 0.06701451 | 0.3841589 | 0.1744448 | 3 | 0.8726241 | 0 | 3 | 1 | not sig |
| SLC6A1 | L54F | CDS2 | O95674 | 0.066632534 | 0.2076177 | 0.3209386 | 14 | 0.7529976 | 0 | 3 | 3 | not sig |
| CHD8 | M904I | PTCD1 | O75127 | 0.065244017 | 0.3824447 | 0.1705972 | 10 | 0.8679427 | 0 | 3 | 3 | not sig |
| TBR1 | N374H | CASK | O14936 | 0.063993126 | 0.2660503 | 0.2405302 | 6 | 0.817927 | 0 | 3 | 3 | not sig |
| KCNQ3 | R236C | CAND2 | O75155 | 0.062687709 | 0.1872798 | 0.3347275 | 10 | 0.7447456 | 0 | 3 | 3 | not sig |
| CHD8 | Q696K | LYRM4 | Q9HD34 | 0.06170127 | 0.5579203 | 0.1105915 | 12 | 0.9137684 | 0 | 3 | 3 | not sig |
| TRAF7 | R593W | CASK | O14936 | 0.06128035 | 0.330842 | 0.1852254 | 1 | 0.8834033 | 0 | 2 | 1 | not sig |
| CHD8 | L834P | XPO5 | Q9HAV4 | 0.059370935 | 0.2355528 | 0.2520494 | 12 | 0.805268 | 0 | 3 | 3 | not sig |
| CHD8 | M904I | KEAP1 | Q14145 | 0.059228415 | 0.2821492 | 0.2099188 | 12 | 0.8372529 | 0 | 3 | 3 | not sig |
| KCNMA1 | A1033P | LAMP1 | P11279 | 0.051404395 | 0.3442553 | 0.1493205 | 8 | 0.8849964 | 0 | 3 | 3 | not sig |
| TBR1 | K228E | ZMYM4 | Q5VZL5 | 0.051030319 | 0.460916 | 0.110715 | 6 | 0.9154529 | 0 | 3 | 3 | not sig |
| MKX | L89F | CHD4 | Q14839 | 0.050394934 | 0 | 0 | 0 | 0 | 0 | 1 | 1 | not sig |
| TRAF7 | R593W | BAG5 | Q9UL15 | 0.049479664 | 0.4828624 | 0.1024716 | 6 | 0.9217213 | 0 | 3 | 3 | not sig |
| TBR1 | N374H | COA7 | Q96BR5 | 0.048541884 | 1.4427159 | 0.0336462 | 6 | 0.9742507 | 0 | 3 | 3 | not sig |
| KCNQ3 | R230H | TNPO1 | Q92973 | 0.047480498 | 0.2814741 | 0.1686851 | 10 | 0.8694074 | 0 | 3 | 3 | not sig |
| PPP1R9B | R753H | MYH10 | P35580 | 0.04656999 | 0.3044433 | 0.1529677 | 4 | 0.8858301 | 0 | 3 | 3 | not sig |
| SLC6A1 | G307R | ILVBL | A1L0T0 | 0.045370395 | 0.1997308 | 0.2271577 | 14 | 0.8235847 | 0 | 3 | 3 | not sig |
| KCNQ3 | R230H | YME1L1 | Q96TA2 | 0.045308615 | 0.1636193 | 0.2769148 | 10 | 0.7874851 | 0 | 3 | 3 | not sig |
| CHD8 | C1095Y | CWC27 | Q6UX04 | 0.045121258 | 0.8586502 | 0.0525491 | 11 | 0.9590334 | 0 | 3 | 3 | not sig |
| SYNGAP1 | C233Y | GDAP1 | Q8TB36 | 0.04482507 | 0.2380166 | 0.1883275 | 6 | 0.8568281 | 0 | 3 | 3 | not sig |
| CHD8 | C1095Y | SNRNP200 | O75643 | 0.044506974 | 0.1812241 | 0.2455908 | 12 | 0.8101491 | 0 | 3 | 3 | not sig |
| RFX3 | A508E | RFX1 | P22670 | 0.04448422 | 0.2088565 | 0.2129894 | 4 | 0.8417499 | 0 | 3 | 3 | not sig |
| PPP1R9B | R753H | WDR48 | Q8TAF3 | 0.043619354 | 0.5019786 | 0.0868948 | 4 | 0.9349312 | 0 | 3 | 3 | not sig |
| KCNMA1 | A1033P | FAM162A | Q96A26 | 0.042224436 | 0.1918833 | 0.2200527 | 8 | 0.8313416 | 0 | 3 | 3 | not sig |
| RFX3 | A508E | OCIAD1 | Q9NX40 | 0.041316035 | 0.3219492 | 0.1283309 | 4 | 0.9040806 | 0 | 3 | 3 | not sig |
| TRAF7 | R655Q | PFDN6 | O15212 | 0.039338983 | 0.1515763 | 0.2595326 | 6 | 0.8038998 | 0 | 3 | 3 | not sig |
| DPYSL2 | R238H | SIRT2 | Q8IXJ6 | 0.037601029 | 0.3589376 | 0.1047565 | 8 | 0.9191479 | 0 | 3 | 3 | not sig |
| SLC6A1 | L54F | DERL2 | Q9GZP9 | 0.0367442 | 0.1679615 | 0.2187656 | 14 | 0.8299897 | 0 | 3 | 3 | not sig |
| SLC6A1 | G299V | MOGS | Q13724 | 0.035566708 | 0.2829625 | 0.1256941 | 14 | 0.9017613 | 0 | 3 | 3 | not sig |
| SLC6A1 | L54F | ILVBL | A1L0T0 | 0.033279881 | 0.1997308 | 0.1666237 | 14 | 0.8700487 | 0 | 3 | 3 | not sig |
| CHD8 | C1095Y | TRIM27 | P14373 | 0.032734014 | 0.1765667 | 0.1853918 | 12 | 0.8560184 | 0 | 3 | 3 | not sig |
| CHD8 | Q696K | XPO5 | Q9HAV4 | 0.030574265 | 0.2355528 | 0.1297979 | 12 | 0.8988766 | 0 | 3 | 3 | not sig |
| TBR1 | K228E | TIMM9 | Q9Y5J7 | 0.029271399 | 0.2815837 | 0.1039528 | 6 | 0.9205945 | 0 | 3 | 3 | not sig |
| TRAF7 | R655Q | NUDC | Q9Y266 | 0.028971319 | 0.1898883 | 0.1525704 | 6 | 0.8837385 | 0 | 3 | 3 | not sig |
| SLC6A1 | G307R | DERL2 | Q9GZP9 | 0.022272292 | 0.1679615 | 0.1326035 | 14 | 0.896394 | 0 | 3 | 3 | not sig |
| MYT1L | H522Q | SAP130 | Q9H0E3 | 0.021754748 | 0.4308345 | 0.0504944 | 6 | 0.9613674 | 0 | 3 | 3 | not sig |
| TBR1 | N374H | ZMYM4 | Q5VZL5 | 0.02156456 | 0.460916 | 0.0467863 | 6 | 0.9642019 | 0 | 3 | 3 | not sig |
| NSD1 | C1877R | ANKRD17 | O75179 | 0.021342736 | 0.4153349 | 0.0513868 | 5 | 0.961007 | 0 | 3 | 3 | not sig |
| PPP1R9B | R753H | TMOD1 | P28289 | 0.018766215 | 0.2550148 | 0.0735887 | 4 | 0.9448706 | 0 | 3 | 3 | not sig |
| RORB | Y22D | EIF4ENIF1 | Q9NRA8 | 0.013566304 | 0.3132368 | 0.0433101 | 4 | 0.9675301 | 0 | 3 | 3 | not sig |
| CHD8 | M904I | XPO5 | Q9HAV4 | 0.012919674 | 0.2355528 | 0.0548483 | 12 | 0.9571619 | 0 | 3 | 3 | not sig |
| CHD8 | C1095Y | CSNK2A2 | P19784 | 0.007447393 | 0.2201047 | 0.0338357 | 12 | 0.9735645 | 0 | 3 | 3 | not sig |
| PPP1R9B | R753H | IQGAP1 | P46940 | 0.006264694 | 0.569778 | 0.010995 | 3 | 0.9919178 | 0 | 2 | 3 | not sig |
| CHD8 | Q696K | GDI2 | P50395 | 0.005527741 | 0.4272981 | 0.0129365 | 10 | 0.9899329 | 0 | 3 | 3 | not sig |
| CHD8 | M904I | BRD2 | P25440 | 0.004655835 | 0.5976791 | 0.0077899 | 12 | 0.9939127 | 0 | 3 | 3 | not sig |
| MYT1L | H522Q | BRMS1L | Q5PSV4 | 0.003927966 | 0.2974702 | 0.0132046 | 6 | 0.9898927 | 0 | 3 | 3 | not sig |
| CHD8 | Q696K | SNRNP200 | O75643 | 0.003899586 | 0.1812241 | 0.021518 | 12 | 0.9831861 | 0 | 3 | 3 | not sig |
| TBR1 | N374H | USP9X | Q93008 | 0.002816778 | 0.2352943 | 0.0119713 | 6 | 0.9908366 | 0 | 3 | 3 | not sig |
| CHD8 | Q696K | TAOK2 | Q9UL54 | 0.002326705 | 0.2536063 | 0.0091745 | 11 | 0.9928442 | 0 | 3 | 3 | not sig |
| CREBBP | R413Q | MRTFB | Q9ULH7 | 0.001428752 | 0.1807483 | 0.0079046 | 1 | 0.9949678 | 0 | 2 | 1 | not sig |
| FOXP1 | R513C | VPS33B | Q9H267 | 0 | 0 | 0 | 0 | 0 | completeMissing | 0 | 0 | not sig |
| FOXP1 | R513C | SESTD1 | Q86VW0 | 0 | 0 | 0 | 0 | 0 | completeMissing | 0 | 0 | not sig |
| FOXP1 | R513H | VPS33B | Q9H267 | 0 | 0 | 0 | 0 | 0 | completeMissing | 0 | 0 | not sig |
| FOXP1 | R513H | SESTD1 | Q86VW0 | 0 | 0 | 0 | 0 | 0 | completeMissing | 0 | 0 | not sig |
| TRAF7 | R655Q | TM9SF4 | Q92544 | 0 | 0 | 0 | 0 | 0 | completeMissing | 0 | 0 | not sig |
| TBL1XR1 | L282P | GPS2 | Q13227 | -8.76E-06 | 0.253021 | -3.46E-05 | 4 | 0.999974 | 0 | 3 | 3 | not sig |
| CHD8 | M904I | CSNK2A2 | P19784 | -0.00053055 | 0.2201047 | -0.00241 | 12 | 0.9981163 | 0 | 3 | 3 | not sig |
| KCNQ3 | R230L | NUP155 | O75694 | -0.0005633 | 0.1741688 | -0.003234 | 10 | 0.9974831 | 0 | 3 | 3 | not sig |
| CHD8 | L834P | LYRM4 | Q9HD34 | -0.00108197 | 0.5579203 | -0.001939 | 12 | 0.9984845 | 0 | 3 | 3 | not sig |
| TBR1 | K228E | ATAD3B | Q5T9A4 | -0.00183736 | 0.8334873 | -0.002204 | 6 | 0.9983126 | 0 | 3 | 3 | not sig |
| CHD8 | C1095Y | HUWE1 | Q7Z6Z7 | -0.0022538 | 0.2091506 | -0.010776 | 12 | 0.9915793 | 0 | 3 | 3 | not sig |
| SLC6A1 | G307R | AUP1 | Q9Y679 | -0.00367231 | 0.1326649 | -0.027681 | 14 | 0.9783072 | 0 | 3 | 3 | not sig |
| KCNQ3 | R236C | YME1L1 | Q96TA2 | -0.00458277 | 0.1636193 | -0.028009 | 10 | 0.9782063 | 0 | 3 | 3 | not sig |
| CHD8 | C1095Y | KPNA5 | O15131 | -0.00517288 | 0.1222152 | -0.042326 | 12 | 0.966935 | 0 | 3 | 3 | not sig |

| | | | | | | | | | | | | |
|---|---|---|---|---|---|---|---|---|---|---|---|---|
| RFX3 | A508E | VPS45 | Q9NRW7 | -0.00786933 | 0.6428317 | -0.012242 | 3 | 0.9910014 | 0 | 3 | 2 | not sig |
| CHD8 | R1580W | NFS1 | Q9Y697 | -0.01317754 | 0.4975548 | -0.026485 | 12 | 0.9793062 | 0 | 3 | 3 | not sig |
| PPP1R9B | R753H | CLPP | Q16740 | -0.01404622 | 0.2984526 | -0.047063 | 4 | 0.9647187 | 0 | 3 | 3 | not sig |
| TBR1 | N374H | PIP4K2C | Q8TBX8 | -0.01490619 | 0.1454686 | -0.10247 | 6 | 0.9217224 | 0 | 3 | 3 | not sig |
| KCNQ3 | R230L | ECD | O95905 | -0.01859459 | 0.2080645 | -0.089369 | 10 | 0.9305529 | 0 | 3 | 3 | not sig |
| SLC6A1 | G307R | IMMT | Q16891 | -0.01922681 | 0.1809264 | -0.106269 | 14 | 0.9168769 | 0 | 3 | 3 | not sig |
| MKX | R93G | LEMD2 | Q8NC56 | -0.02190731 | 0.5157581 | -0.042476 | 5 | 0.9677633 | 0 | 3 | 2 | not sig |
| PPP1R9B | R753H | SPTBN1 | Q01082 | -0.02554741 | 0.7248794 | -0.035244 | 4 | 0.9735741 | 0 | 3 | 3 | not sig |
| PPP1R9B | R753H | CAND1 | Q86VP6 | -0.02560115 | 0.0770866 | -0.332109 | 4 | 0.7564809 | 0 | 3 | 3 | not sig |
| CHD8 | R1580W | CWC27 | Q6UX04 | -0.02611636 | 0.9600001 | -0.027205 | 11 | 0.9787839 | 0 | 2 | 3 | not sig |
| SLC6A1 | L54F | SLC39A14 | Q15043 | -0.02737979 | 0.1552071 | -0.176408 | 13 | 0.8626918 | 0 | 2 | 3 | not sig |
| CHD8 | M904I | NFS1 | Q9Y697 | -0.02795597 | 0.4975548 | -0.056187 | 12 | 0.9561177 | 0 | 3 | 3 | not sig |
| CHD8 | Q696K | NUP153 | P49790 | -0.02949768 | 0.5825294 | -0.050637 | 12 | 0.9604477 | 0 | 3 | 3 | not sig |
| KCNQ3 | R230L | TNPO1 | Q92973 | -0.02950388 | 0.2814741 | -0.104819 | 10 | 0.9185919 | 0 | 3 | 3 | not sig |
| TBR1 | K228E | ZMYM2 | Q9UBW7 | -0.03442988 | 0.709013 | -0.04856 | 6 | 0.9628458 | 0 | 3 | 3 | not sig |
| CHD8 | Q696K | NDUFAB1 | O14561 | -0.03558119 | 0.4637705 | -0.076722 | 12 | 0.9401094 | 0 | 3 | 3 | not sig |
| SLC6A1 | L54F | YIF1B | Q5BJH7 | -0.03725101 | 0.3358985 | -0.1109 | 11 | 0.9136935 | 0 | 2 | 2 | not sig |
| NSD1 | C1877R | MKI67 | P46013 | -0.03731854 | 0.5628081 | -0.066308 | 6 | 0.9492871 | 0 | 3 | 3 | not sig |
| CHD8 | M904I | TRIM27 | P14373 | -0.03732608 | 0.1765667 | -0.211399 | 12 | 0.8361233 | 0 | 3 | 3 | not sig |
| MYT1L | C504R | KDM1A | O60341 | -0.03747883 | 0.3785711 | -0.099001 | 6 | 0.9243624 | 0 | 3 | 3 | not sig |
| TRAF7 | R655Q | HUWE1 | Q7Z6Z7 | -0.03824658 | 0.0839166 | -0.455769 | 6 | 0.6645861 | 0 | 3 | 3 | not sig |
| TBR1 | N374H | POLDIP2 | Q9Y2S7 | -0.03970753 | 0.1041988 | -0.381075 | 6 | 0.7162764 | 0 | 3 | 3 | not sig |
| MKX | R93G | IDE | P14735 | -0.0397557 | 0.3523154 | -0.112841 | 5 | 0.9145469 | 0 | 3 | 2 | not sig |
| CREBBP | R413Q | MRTFA | Q969V6 | -0.04024373 | 0.1588996 | -0.253265 | 4 | 0.8125475 | 0 | 3 | 3 | not sig |
| SLC6A1 | L54F | NUP210 | Q8TEM1 | -0.04034679 | 0.221156 | -0.182436 | 14 | 0.857856 | 0 | 3 | 3 | not sig |
| TBR1 | K228E | NAA15 | Q9BXJ9 | -0.04275425 | 0.3380705 | -0.126465 | 5 | 0.9042917 | 0 | 2 | 3 | not sig |
| TRAF7 | R655Q | PFDN5 | Q99471 | -0.0428285 | 0.1218812 | -0.351395 | 6 | 0.7373024 | 0 | 3 | 3 | not sig |
| MKX | R93G | CHD4 | Q14839 | -0.04521779 | 0 | 0 | 0 | 0 | 0 | 1 | 1 | not sig |
| CREBBP | R413Q | NCOA2 | Q15596 | -0.04697551 | 0.1853994 | -0.253375 | 4 | 0.8124685 | 0 | 3 | 3 | not sig |
| KCNQ3 | R230C | NAA15 | Q9BXJ9 | -0.04717081 | 0.2555836 | -0.184561 | 9 | 0.857665 | 0 | 3 | 3 | not sig |
| TRAF7 | R655Q | PDCD5 | O14737 | -0.04778784 | 0.1941998 | -0.246076 | 6 | 0.8138256 | 0 | 3 | 3 | not sig |
| CHD8 | M904I | BRD4 | O60885 | -0.04865833 | 0.7882016 | -0.061733 | 11 | 0.9518826 | 0 | 2 | 3 | not sig |
| SLC6A1 | G307R | ESYT1 | Q9BSJ8 | -0.04866083 | 0.1185535 | -0.410455 | 14 | 0.6876851 | 0 | 3 | 3 | not sig |
| CHD8 | M904I | HUWE1 | Q7Z6Z7 | -0.04891695 | 0.2091506 | -0.233884 | 12 | 0.8190176 | 0 | 3 | 3 | not sig |
| PPP1R9B | R753H | TLN1 | Q9Y490 | -0.05027823 | 0.1898231 | -0.264869 | 4 | 0.8041993 | 0 | 3 | 3 | not sig |
| CHD8 | Q696K | CWC27 | Q6UX04 | -0.05259155 | 0.8586502 | -0.061249 | 11 | 0.9522595 | 0 | 3 | 3 | not sig |
| CHD8 | Q696K | NFS1 | Q9Y697 | -0.05285402 | 0.4975548 | -0.106228 | 12 | 0.917157 | 0 | 3 | 3 | not sig |
| TBR1 | N374H | DLG1 | Q12959 | -0.05377566 | 0.2011385 | -0.267356 | 6 | 0.7981473 | 0 | 3 | 3 | not sig |
| TBR1 | K228E | CLPB | Q9H078 | -0.05608936 | 0.7005874 | -0.08006 | 6 | 0.9387927 | 0 | 3 | 3 | not sig |
| TBR1 | N374H | DARS2 | Q6PI48 | -0.05738878 | 0 | 0 | 0 | 0 | 0 | 1 | 1 | not sig |
| KCNQ3 | R236C | IPO9 | Q96P70 | -0.05880435 | 0.1425257 | -0.412588 | 10 | 0.688615 | 0 | 3 | 3 | not sig |
| CHD8 | R1580W | MYH9 | P35579 | -0.0671576 | 0.350281 | -0.191725 | 12 | 0.8511636 | 0 | 3 | 3 | not sig |
| SLC6A1 | G307R | CERS1 | P27544 | -0.067845 | 0.2321979 | -0.292186 | 14 | 0.7744296 | 0 | 3 | 3 | not sig |
| SLC6A1 | G307R | CERS2 | Q96G23 | -0.06891771 | 0.2014027 | -0.342189 | 14 | 0.7372916 | 0 | 3 | 3 | not sig |
| SLC6A1 | L54F | RTN3 | O95197 | -0.06897981 | 0.1649028 | -0.418306 | 14 | 0.6820692 | 0 | 3 | 3 | not sig |
| MYT1L | H522Q | SIN3A | Q96ST3 | -0.07068211 | 0.4172319 | -0.169407 | 6 | 0.8710436 | 0 | 3 | 3 | not sig |
| SLC6A1 | G307R | MYH9 | P35579 | -0.07093231 | 0.3279728 | -0.216275 | 14 | 0.831893 | 0 | 3 | 3 | not sig |
| SLC6A1 | G307R | UBAC2 | Q8NBM4 | -0.07524376 | 0.1337856 | -0.56242 | 14 | 0.5827238 | 0 | 3 | 3 | not sig |
| KCNQ3 | R236C | TNPO1 | Q92973 | -0.07879464 | 0.2814741 | -0.279936 | 10 | 0.785232 | 0 | 3 | 3 | not sig |
| KCNMA1 | A1033P | TIMM21 | Q9BVV7 | -0.08401176 | 0.2810451 | -0.298926 | 8 | 0.7726143 | 0 | 3 | 3 | not sig |
| VEZF1 | Q209P | MRPS9 | P82933 | -0.08461451 | 0.4066281 | -0.208088 | 4 | 0.8453259 | 0 | 3 | 3 | not sig |
| CHD8 | L834P | CWC27 | Q6UX04 | -0.08627298 | 0.8586502 | -0.100475 | 11 | 0.921775 | 0 | 3 | 3 | not sig |
| MYT1L | C504R | GRAMD4 | Q6IC98 | -0.08786039 | 0.5287773 | -0.166158 | 5 | 0.8745429 | 0 | 3 | 2 | not sig |
| SLC6A1 | G307R | DHCR24 | Q15392 | -0.08999353 | 0.2596211 | -0.346634 | 14 | 0.734021 | 0 | 3 | 3 | not sig |
| RFX3 | A508E | CKAP4 | Q07065 | -0.09480758 | 0.1767265 | -0.536465 | 4 | 0.6200829 | 0 | 3 | 3 | not sig |
| TBR1 | K228E | COA7 | Q96BR5 | -0.10017716 | 1.4427159 | -0.069437 | 6 | 0.9468985 | 0 | 3 | 3 | not sig |
| KCNQ3 | R230L | CLPB | Q9H078 | -0.10194908 | 0.1558742 | -0.654047 | 10 | 0.5278352 | 0 | 3 | 3 | not sig |
| FOXP2 | R570C | CTBP1 | Q13363 | -0.10207167 | 0.3848451 | -0.265228 | 3 | 0.8080145 | 0 | 3 | 2 | not sig |
| SLC6A1 | G307R | NUP210 | Q8TEM1 | -0.10618593 | 0.221156 | -0.48014 | 14 | 0.638543 | 0 | 3 | 3 | not sig |
| DPYSL2 | H333N | DPYSL4 | O14531 | -0.10672068 | 1.502358 | -0.071035 | 5 | 0.9461233 | 0 | 2 | 3 | not sig |
| SYNGAP1 | C233Y | YWHAG | P61981 | -0.11288629 | 0.1599776 | -0.705638 | 6 | 0.5068715 | 0 | 3 | 3 | not sig |
| SLC6A1 | G307R | RTN3 | O95197 | -0.11398547 | 0.1649028 | -0.691228 | 14 | 0.5007241 | 0 | 3 | 3 | not sig |
| KCNQ3 | R236C | ELOB | Q15370 | -0.11880015 | 0.2108906 | -0.563326 | 10 | 0.5856218 | 0 | 3 | 3 | not sig |
| CHD8 | C1095Y | XPO5 | Q9HAV4 | -0.12358442 | 0.2355528 | -0.524657 | 12 | 0.6093775 | 0 | 3 | 3 | not sig |
| CHD8 | M904I | COPS3 | Q9UNS2 | -0.1245869 | 0.3250456 | -0.383291 | 6 | 0.7147171 | 0 | 2 | 3 | not sig |
| CHD8 | R1580W | TRIM27 | P14373 | -0.12594892 | 0.1765667 | -0.713322 | 12 | 0.4892872 | 0 | 3 | 3 | not sig |
| TBR1 | N374H | CLPB | Q9H078 | -0.12639196 | 0.7005874 | -0.180409 | 6 | 0.8627711 | 0 | 3 | 3 | not sig |
| CHD8 | Q696K | CSNK2A2 | P19784 | -0.12927847 | 0.2201047 | -0.58735 | 12 | 0.5678561 | 0 | 3 | 3 | not sig |
| KCNQ3 | R236C | ARL10 | Q8N8L6 | -0.13376467 | 0.2217619 | -0.60319 | 10 | 0.5598133 | 0 | 3 | 3 | not sig |
| MYT1L | H522Q | SUDS3 | Q9H7L9 | -0.13428684 | 0.3884724 | -0.345679 | 6 | 0.7413814 | 0 | 3 | 3 | not sig |
| TRAF7 | R655Q | DLG1 | Q12959 | -0.13439125 | 0.3643008 | -0.368902 | 6 | 0.7248684 | 0 | 3 | 3 | not sig |
| TBR1 | K228E | CASK | O14936 | -0.13545997 | 0.2660503 | -0.509152 | 6 | 0.628823 | 0 | 3 | 3 | not sig |

| | | | | | | | | | | | | |
|---|---|---|---|---|---|---|---|---|---|---|---|---|
| SLC6A1 | G307R | CHP1 | Q99653 | -0.13773962 | 0.1798068 | -0.766042 | 14 | 0.4563734 | 0 | 3 | 3 | not sig |
| TBR1 | K228E | BAG3 | O95817 | -0.13939562 | 0.4058449 | -0.34347 | 6 | 0.7429604 | 0 | 3 | 3 | not sig |
| PPP1R9B | R753H | HYOU1 | Q9Y4L1 | -0.1431094 | 0.1831844 | -0.781231 | 4 | 0.4783186 | 0 | 3 | 3 | not sig |
| TBR1 | N374H | BAG3 | O95817 | -0.14526227 | 0.4058449 | -0.357926 | 6 | 0.7326539 | 0 | 3 | 3 | not sig |
| SLC6A1 | H198P | LMBR1 | Q8WVP7 | -0.14705643 | 0.572754 | -0.256753 | 11 | 0.8021082 | 0 | 2 | 2 | not sig |
| MKX | L89F | IDE | P14735 | -0.14857289 | 0.3523154 | -0.421704 | 5 | 0.6907575 | 0 | 3 | 2 | not sig |
| TBR1 | K228E | FBXW11 | Q9UKB1 | -0.15231241 | 1.1222771 | -0.135717 | 6 | 0.8964836 | 0 | 3 | 3 | not sig |
| CHD8 | M904I | KPNA5 | O15131 | -0.15240392 | 0.1222152 | -1.247013 | 12 | 0.2361833 | 0 | 3 | 3 | not sig |
| KCNMA1 | A1033P | UBAC2 | Q8NBM4 | -0.15467535 | 0.2089699 | -0.74018 | 8 | 0.4803467 | 0 | 3 | 3 | not sig |
| PPP1R9B | R753H | DYNLT1 | P63172 | -0.15764265 | 0.1194895 | -1.319301 | 4 | 0.2575193 | 0 | 3 | 3 | not sig |
| TBR1 | K228E | USP9X | Q93008 | -0.15900923 | 0.2352943 | -0.675789 | 6 | 0.524352 | 0 | 3 | 3 | not sig |
| CHD8 | C1095Y | COPS3 | Q9UNS2 | -0.15950776 | 0.3250456 | -0.490724 | 6 | 0.6410509 | 0 | 2 | 3 | not sig |
| ASH1L | F1944C | MORF4L2 | Q15014 | -0.16380775 | 0.136868 | -1.19683 | 4 | 0.2974554 | 0 | 3 | 3 | not sig |
| TBR1 | K228E | HSD17B4 | P51659 | -0.16428171 | 0.8405961 | -0.195435 | 1 | 0.877131 | 0 | 2 | 1 | not sig |
| KCNQ3 | R230H | PRKDC | P78527 | -0.16651141 | 0.1166369 | -1.427605 | 10 | 0.1838796 | 0 | 3 | 3 | not sig |
| MYT1L | H522Q | SAP30 | O75446 | -0.16949002 | 0.3767999 | -0.449814 | 6 | 0.6686383 | 0 | 3 | 3 | not sig |
| TBR1 | K228E | DLG1 | Q12959 | -0.17154799 | 0.2011385 | -0.852885 | 6 | 0.4264529 | 0 | 3 | 3 | not sig |
| KCNQ3 | R236C | FANCI | Q9NVI1 | -0.17294832 | 0.29473 | -0.586803 | 7 | 0.5757639 | 0 | 3 | 3 | not sig |
| CREBBP | R413Q | TP53 | P04637 | -0.17477961 | 0.311547 | -0.561006 | 4 | 0.6047319 | 0 | 3 | 3 | not sig |
| TBR1 | N374H | TMEM177 | Q53S58 | -0.18395678 | 0.948553 | -0.193934 | 6 | 0.8526266 | 0 | 3 | 3 | not sig |
| CHD8 | R1580W | COPS6 | Q7L5N1 | -0.18400383 | 0.1984548 | -0.927182 | 11 | 0.3737282 | 0 | 3 | 3 | not sig |
| CHD8 | M904I | OAT | P04181 | -0.18638272 | 0.3848641 | -0.484282 | 11 | 0.6376841 | 0 | 3 | 3 | not sig |
| SYNGAP1 | C233Y | YWHAH | Q04917 | -0.19196225 | 0.1455771 | -1.318629 | 6 | 0.235388 | 0 | 3 | 3 | not sig |
| SYNGAP1 | S140F | YWHAG | P61981 | -0.19435714 | 0.1599776 | -1.214902 | 6 | 0.2700395 | 0 | 3 | 3 | not sig |
| CHD8 | C1095Y | PTCD1 | O75127 | -0.19582816 | 0.3824447 | -0.512043 | 10 | 0.6197383 | 0 | 3 | 3 | not sig |
| TBR1 | N374H | TIMM9 | Q9Y5J7 | -0.20073812 | 0.2815837 | -0.71289 | 6 | 0.5026837 | 0 | 3 | 3 | not sig |
| KCNQ3 | R230H | FANCD2 | Q9BXW9 | -0.20273215 | 0.1665422 | -1.217302 | 8 | 0.2581761 | 0 | 2 | 3 | not sig |
| KCNQ3 | R230L | CAND2 | O75155 | -0.20326755 | 0.1872798 | -1.085368 | 10 | 0.3032302 | 0 | 3 | 3 | not sig |
| CHD8 | C1095Y | COPS6 | Q7L5N1 | -0.20376934 | 0.2218793 | -0.918379 | 11 | 0.3781178 | 0 | 2 | 3 | not sig |
| KCNQ3 | R230L | PPM1G | O15355 | -0.20698735 | 0.2028349 | -1.020472 | 10 | 0.3315576 | 0 | 3 | 3 | not sig |
| KCNQ3 | R230H | PPM1G | O15355 | -0.20834894 | 0.2028349 | -1.027185 | 10 | 0.3285384 | 0 | 3 | 3 | not sig |
| MKX | R93G | NAP1L4 | Q99733 | -0.21050831 | 0.3031999 | -0.694289 | 5 | 0.5184316 | 0 | 3 | 2 | not sig |
| TBR1 | K228E | PIGT | Q969N2 | -0.21429476 | 0.3935606 | -0.544503 | 5 | 0.6094902 | 0 | 2 | 3 | not sig |
| KCNMA1 | A1033P | TNPO1 | Q92973 | -0.21573875 | 0.8592198 | -0.251087 | 7 | 0.8089581 | 0 | 3 | 3 | not sig |
| CHD8 | Q696K | PTCD1 | O75127 | -0.2175156 | 0.3824447 | -0.56875 | 10 | 0.5820726 | 0 | 3 | 3 | not sig |
| TBR1 | N374H | AP2A2 | O94973 | -0.21770368 | 0.7278634 | -0.2991 | 3 | 0.7843873 | 0 | 1 | 3 | not sig |
| TBR1 | K228E | AP2S1 | P53680 | -0.21860219 | 1.0971996 | -0.199236 | 3 | 0.854817 | 0 | 1 | 3 | not sig |
| TRAF7 | R655Q | SCRIB | Q14160 | -0.22080076 | 0.1830177 | -1.206445 | 6 | 0.273052 | 0 | 3 | 3 | not sig |
| SLC6A1 | A288V | MOGS | Q13724 | -0.22456297 | 0.2829625 | -0.793614 | 14 | 0.4406645 | 0 | 3 | 3 | not sig |
| TBR1 | K228E | BAIAP2 | Q9UQB8 | -0.22515655 | 1.295009 | -0.173865 | 1 | 0.8904097 | 0 | 1 | 2 | not sig |
| SYNGAP1 | S140F | YWHAE | P62258 | -0.22750258 | 0.0839505 | -2.70996 | 6 | 0.035108 | 0 | 3 | 3 | not sig |
| RORB | Y22D | GLB1 | P16278 | -0.22846158 | 0.4218801 | -0.541532 | 4 | 0.6168937 | 0 | 3 | 3 | not sig |
| TBR1 | N374H | OCLN | Q16625 | -0.22967214 | 0.2380271 | -0.964899 | 6 | 0.3718584 | 0 | 3 | 3 | not sig |
| CHD8 | Q696K | HUWE1 | Q7Z6Z7 | -0.23330616 | 0.2091506 | -1.115494 | 12 | 0.2864796 | 0 | 3 | 3 | not sig |
| MYT1L | C504R | SCOC | Q9UIL1 | -0.23640348 | 0.3381821 | -0.699042 | 6 | 0.5107008 | 0 | 3 | 3 | not sig |
| KCNQ3 | R230H | IPO9 | Q96P70 | -0.23877813 | 0.1425257 | -1.675334 | 10 | 0.1248064 | 0 | 3 | 3 | not sig |
| KCNQ3 | R236C | CLPB | Q9H078 | -0.24318449 | 0.1558742 | -1.560133 | 10 | 0.1497892 | 0 | 3 | 3 | not sig |
| KCNMA1 | A1033P | RMDN3 | Q96TC7 | -0.2457325 | 0.4973211 | -0.494112 | 7 | 0.6363532 | 0 | 2 | 3 | not sig |
| SLC6A1 | H198P | MOGS | Q13724 | -0.24581946 | 0.2829625 | -0.868735 | 14 | 0.3996403 | 0 | 3 | 3 | not sig |
| RFX3 | A508E | MOGS | Q13724 | -0.24972314 | 0.1210621 | -2.06277 | 4 | 0.1081124 | 0 | 3 | 3 | not sig |
| CHD8 | L834P | COPS6 | Q7L5N1 | -0.24989828 | 0.1984548 | -1.25922 | 11 | 0.2340158 | 0 | 3 | 3 | not sig |
| CHD8 | Q696K | KPNA5 | O15131 | -0.25072632 | 0.1222152 | -2.051515 | 12 | 0.0627038 | 0 | 3 | 3 | not sig |
| RORB | Y22D | NCOA3 | Q9Y6Q9 | -0.25208809 | 0.3310554 | -0.761468 | 4 | 0.4887986 | 0 | 3 | 3 | not sig |
| CHD8 | R1580W | KPNA2 | P52292 | -0.25315356 | 0.1379634 | -1.834933 | 12 | 0.0914121 | 0 | 3 | 3 | not sig |
| KCNQ3 | R230C | CLPB | Q9H078 | -0.25402991 | 0.1558742 | -1.629711 | 10 | 0.1342173 | 0 | 3 | 3 | not sig |
| TBR1 | N374H | POLD1 | P28340 | -0.25426065 | 0.2855381 | -0.890461 | 6 | 0.4075032 | 0 | 3 | 3 | not sig |
| CHD8 | R1580W | LYRM4 | Q9HD34 | -0.25469552 | 0.5579203 | -0.456509 | 12 | 0.6561767 | 0 | 3 | 3 | not sig |
| CHD8 | M904I | KPNA2 | P52292 | -0.25504172 | 0.1379634 | -1.848619 | 12 | 0.0892917 | 0 | 3 | 3 | not sig |
| CHD8 | M904I | LYRM4 | Q9HD34 | -0.25862598 | 0.5579203 | -0.463554 | 12 | 0.6512629 | 0 | 3 | 3 | not sig |
| KCNQ3 | R236C | PRKDC | P78527 | -0.25923687 | 0.1166369 | -2.222598 | 10 | 0.0504718 | 0 | 3 | 3 | not sig |
| TBR1 | N374H | MCAT | Q8IVS2 | -0.26562382 | 0.8757478 | -0.303311 | 4 | 0.7767741 | 0 | 2 | 3 | not sig |
| CHD8 | R1580W | CSNK2A2 | P19784 | -0.27158434 | 0.2201047 | -1.233887 | 12 | 0.2408654 | 0 | 3 | 3 | not sig |
| KCNQ3 | R230L | FANCI | Q9NVI1 | -0.27188821 | 0.3295181 | -0.825109 | 7 | 0.4365217 | 0 | 2 | 3 | not sig |
| PPP1R9B | R753H | ZC3H18 | Q86VM9 | -0.27220874 | 0.1726246 | -1.576883 | 4 | 0.1899544 | 0 | 3 | 3 | not sig |
| TBR1 | N374H | PKP4 | Q99569 | -0.27230884 | 0.6358089 | -0.428287 | 4 | 0.6904963 | 0 | 3 | 2 | not sig |
| TBR1 | N374H | WDR11 | Q9BZH6 | -0.28012286 | 0.6649928 | -0.421242 | 5 | 0.6910736 | 0 | 3 | 2 | not sig |
| TBR1 | K228E | TMEM177 | Q53S58 | -0.28144942 | 0.948553 | -0.296714 | 6 | 0.7766878 | 0 | 3 | 3 | not sig |
| KCNQ3 | R230H | NAP1L4 | Q99733 | -0.28196215 | 0.215446 | -1.308737 | 9 | 0.2230535 | 0 | 3 | 3 | not sig |
| KCNQ3 | R230L | PIP4P2 | Q8N4L2 | -0.28358436 | 0.407462 | -0.695977 | 3 | 0.5365155 | 0 | 2 | 1 | not sig |
| TRAF7 | R655Q | PFDN4 | Q9NQP4 | -0.28489883 | 0.2261186 | -1.259953 | 6 | 0.2544741 | 0 | 3 | 3 | not sig |
| TBR1 | K228E | POLDIP2 | Q9Y2S7 | -0.28663615 | 0.1041988 | -2.750857 | 6 | 0.033254 | 0 | 3 | 3 | not sig |
| KCNQ3 | R230C | ARL10 | Q8N8L6 | -0.28731453 | 0.2217619 | -1.295599 | 10 | 0.2242208 | 0 | 3 | 3 | not sig |

| | | | | | | | | | | | | |
|---|---|---|---|---|---|---|---|---|---|---|---|---|
| SYNGAP1 | C233Y | YWHAE | P62258 | -0.28739275 | 0.0839505 | -3.423358 | 6 | 0.0140855 | 0 | 3 | 3 | not sig |
| TBR1 | N374H | TARS2 | Q9BW92 | -0.28781556 | 0.7110081 | -0.404799 | 6 | 0.6996615 | 0 | 3 | 3 | not sig |
| TBR1 | K228E | POLD1 | P28340 | -0.29017298 | 0.2855381 | -1.016232 | 6 | 0.3487322 | 0 | 3 | 3 | not sig |
| PTK7 | R562Q | TOMM40 | O96008 | -0.29080382 | 0.0935504 | -3.108524 | 4 | 0.0359232 | 0 | 3 | 3 | not sig |
| KCNQ3 | R230L | IPO9 | Q96P70 | -0.2915844 | 0.1425257 | -2.045837 | 10 | 0.0679802 | 0 | 3 | 3 | not sig |
| TBR1 | K228E | PITRM1 | Q5JRX3 | -0.29485841 | 2.116517 | -0.139313 | 6 | 0.8937614 | 0 | 3 | 3 | not sig |
| SLC6A1 | A288V | LMBR1 | Q8WVP7 | -0.29612256 | 0.5228505 | -0.566362 | 11 | 0.5825184 | 0 | 3 | 2 | not sig |
| TBR1 | N374H | FHL3 | Q13643 | -0.2994419 | 0.7232408 | -0.414028 | 6 | 0.6932467 | 0 | 3 | 3 | not sig |
| FOXP1 | R513C | CEP55 | Q53EZ4 | -0.30363249 | 0.7487683 | -0.405509 | 4 | 0.7058562 | 0 | 2 | 2 | not sig |
| RFX3 | A508E | PKP2 | Q99959 | -0.30732716 | 0.3838382 | -0.800669 | 3 | 0.4818652 | 0 | 3 | 2 | not sig |
| SYNGAP1 | S140F | YWHAH | Q04917 | -0.30818927 | 0.1455771 | -2.117017 | 6 | 0.0786068 | 0 | 3 | 3 | not sig |
| MYT1L | H522Q | ACAD11 | Q709F0 | -0.31009651 | 0.6172821 | -0.502358 | 6 | 0.6333165 | 0 | 3 | 3 | not sig |
| TBR1 | K228E | BTRC | Q9Y297 | -0.31215285 | 0.9975799 | -0.31291 | 4 | 0.7699847 | 0 | 2 | 2 | not sig |
| RORB | Y22D | NCOA1 | Q15788 | -0.31791949 | 0.154785 | -2.053943 | 4 | 0.1092004 | 0 | 3 | 3 | not sig |
| CHD8 | R1580W | EPS15L1 | Q9UBC2 | -0.32143307 | 1.3903328 | -0.231191 | 8 | 0.8229694 | 0 | 3 | 1 | not sig |
| TBR1 | K228E | MCAT | Q8IVS2 | -0.32161153 | 0.8757478 | -0.367242 | 4 | 0.7320428 | 0 | 2 | 3 | not sig |
| KCNQ3 | R236C | NAP1L4 | Q99733 | -0.32410942 | 0.215446 | -1.504365 | 9 | 0.1667438 | 0 | 3 | 3 | not sig |
| KCNMA1 | A1033P | MARVELD1 | Q9BSK0 | -0.3266772 | 0.3001061 | -1.088539 | 8 | 0.3080597 | 0 | 3 | 3 | not sig |
| CHD8 | L834P | MYH9 | P35579 | -0.33047299 | 0.350281 | -0.943451 | 12 | 0.3640582 | 0 | 3 | 3 | not sig |
| TBR1 | K228E | SEC16A | O15027 | -0.33359869 | 0.4108656 | -0.811941 | 4 | 0.4623793 | 0 | 3 | 2 | not sig |
| CHD8 | Q696K | KPNA2 | P52292 | -0.33646314 | 0.1379634 | -2.438786 | 12 | 0.0312271 | 0 | 3 | 3 | not sig |
| RFX3 | A508E | ANKHD1 | Q8IWZ3 | -0.33682773 | 0.1932241 | -1.743198 | 4 | 0.156253 | 0 | 3 | 3 | not sig |
| PPP1R9B | R753H | CFAP97 | Q9P2B7 | -0.3388257 | 0.2414109 | -1.403523 | 4 | 0.2331276 | 0 | 3 | 3 | not sig |
| SLC6A1 | L54F | LMBR1 | Q8WVP7 | -0.34572408 | 0.5228505 | -0.661229 | 11 | 0.522074 | 0 | 3 | 2 | not sig |
| KCNQ3 | R236C | FANCD2 | Q9BXW9 | -0.34620649 | 0.1489599 | -2.324159 | 8 | 0.0486028 | 0 | 3 | 3 | not sig |
| TBR1 | N374H | AP2S1 | P53680 | -0.34622597 | 0.8674125 | -0.399148 | 3 | 0.7164998 | 0 | 2 | 3 | not sig |
| SYNGAP1 | S140F | GDAP1 | Q8TB36 | -0.3488248 | 0.2380166 | -1.465548 | 6 | 0.1931237 | 0 | 3 | 3 | not sig |
| CHD8 | Q696K | MYH9 | P35579 | -0.34937751 | 0.350281 | -0.997421 | 12 | 0.3382486 | 0 | 3 | 3 | not sig |
| KCNQ3 | R236C | NUP155 | O75694 | -0.34983571 | 0.1741688 | -2.008601 | 10 | 0.0723432 | 0 | 3 | 3 | not sig |
| CHD8 | C1095Y | MYH9 | P35579 | -0.35645702 | 0.350281 | -1.017632 | 12 | 0.328932 | 0 | 3 | 3 | not sig |
| TBR1 | K228E | FHL3 | Q13643 | -0.36184782 | 0.7232408 | -0.500314 | 6 | 0.6346714 | 0 | 3 | 3 | not sig |
| TRAF7 | R655Q | BAG5 | Q9UL15 | -0.36263605 | 0.4828624 | -0.751013 | 6 | 0.4810513 | 0 | 3 | 3 | not sig |
| FOXP2 | R570C | BCKDK | O14874 | -0.37047277 | 0.8519911 | -0.434832 | 2 | 0.7061061 | 0 | 3 | 1 | not sig |
| KCNQ3 | R236C | SAAL1 | Q96ER3 | -0.37922456 | 0.3112131 | -1.218537 | 10 | 0.2509865 | 0 | 3 | 3 | not sig |
| KCNQ3 | R230L | HEATR3 | Q7Z4Q2 | -0.38027386 | 0.3223961 | -1.179524 | 8 | 0.2720749 | 0 | 2 | 3 | not sig |
| CHD8 | M904I | MYH9 | P35579 | -0.38131349 | 0.350281 | -1.088593 | 12 | 0.2977132 | 0 | 3 | 3 | not sig |
| KCNMA1 | A1033P | EMC1 | Q8N766 | -0.38139171 | 0.3649036 | -1.045185 | 8 | 0.3264876 | 0 | 3 | 3 | not sig |
| PTK7 | R562Q | CKAP4 | Q07065 | -0.38192239 | 0.0809766 | -4.716452 | 4 | 0.0091957 | 0 | 3 | 3 | not sig |
| LRRC4C | I163N | XPO5 | Q9HAV4 | -0.38240105 | 0.3901289 | -0.980192 | 3 | 0.3992743 | 0 | 2 | 3 | not sig |
| CHD8 | M904I | MYH10 | P35580 | -0.38336634 | 0.3615007 | -1.060486 | 12 | 0.3098023 | 0 | 3 | 3 | not sig |
| RORB | Y22D | ZNF629 | Q9UEG4 | -0.38337508 | 0.2615423 | -1.465825 | 4 | 0.2165732 | 0 | 3 | 3 | not sig |
| STXBP1 | A251T | STXBP2 | Q15833 | -0.38686036 | 1.2317404 | -0.314076 | 6 | 0.7640966 | 0 | 3 | 3 | not sig |
| FOXP1 | L327P | NUP155 | O75694 | -0.39013406 | 0.2814166 | -1.386322 | 8 | 0.2030596 | 0 | 3 | 3 | not sig |
| KCNQ3 | R236C | ECD | O95905 | -0.39065312 | 0.2080645 | -1.877558 | 10 | 0.0898841 | 0 | 3 | 3 | not sig |
| CHD8 | R1580W | MYH10 | P35580 | -0.39765808 | 0.3615007 | -1.10002 | 12 | 0.2929012 | 0 | 3 | 3 | not sig |
| KCNQ3 | R230C | CAND2 | O75155 | -0.40209004 | 0.1872798 | -2.147001 | 10 | 0.0573538 | 0 | 3 | 3 | not sig |
| KCNQ3 | R230H | HEATR3 | Q7Z4Q2 | -0.40277323 | 0.2883598 | -1.396773 | 8 | 0.2000123 | 0 | 3 | 3 | not sig |
| TBR1 | K228E | WDR6 | Q9NNW5 | -0.41024619 | 0.4649536 | -0.882338 | 6 | 0.4115454 | 0 | 3 | 3 | not sig |
| CHD8 | Q696K | TRIM27 | P14373 | -0.41975567 | 0.1765667 | -2.377321 | 12 | 0.0349328 | 0 | 3 | 3 | not sig |
| TBR1 | N374H | ERCC3 | P19447 | -0.42003887 | 0.3828032 | -1.097271 | 4 | 0.3341422 | 0 | 3 | 2 | not sig |
| TBR1 | K228E | TARS2 | Q9BW92 | -0.4216626 | 0.7110081 | -0.593049 | 6 | 0.5748054 | 0 | 3 | 3 | not sig |
| MKX | R93G | AMPD2 | Q01433 | -0.42431244 | 0.1970635 | -2.153176 | 5 | 0.0839075 | 0 | 3 | 2 | not sig |
| KCNQ3 | R230C | NUP155 | O75694 | -0.43022033 | 0.1741688 | -2.470134 | 10 | 0.0330973 | 0 | 3 | 3 | not sig |
| KCNQ3 | R230L | FANCD2 | Q9BXW9 | -0.43151533 | 0.1665422 | -2.591027 | 8 | 0.0320619 | 0 | 2 | 3 | not sig |
| CHD8 | C1095Y | KPNA2 | P52292 | -0.43254382 | 0.1379634 | -3.135208 | 12 | 0.0086082 | 0 | 3 | 3 | not sig |
| SLC6A1 | G360S | MOGS | Q13724 | -0.43607833 | 0.2829625 | -1.541117 | 14 | 0.1455832 | 0 | 3 | 3 | not sig |
| CHD8 | C1095Y | MYH10 | P35580 | -0.43707669 | 0.3615007 | -1.209062 | 12 | 0.2499225 | 0 | 3 | 3 | not sig |
| KCNQ3 | R230C | YME1L1 | Q96TA2 | -0.44206007 | 0.1636193 | -2.70176 | 10 | 0.0222461 | 0 | 3 | 3 | not sig |
| MYT1L | C504R | SAP130 | Q9H0E3 | -0.44473103 | 0.4308345 | -1.032255 | 6 | 0.3417543 | 0 | 3 | 3 | not sig |
| CHD8 | M904I | EPS15L1 | Q9UBC2 | -0.44838036 | 1.4746707 | -0.304055 | 8 | 0.7688447 | 0 | 2 | 1 | not sig |
| AP2S1 | G64D | CCDC32 | Q9BV29 | -0.4530251 | 0.5214971 | -0.868701 | 5 | 0.4247369 | 0 | 3 | 2 | not sig |
| CHD8 | Q696K | KEAP1 | Q14145 | -0.45477864 | 0.2821492 | -1.611837 | 12 | 0.1329707 | 0 | 3 | 3 | not sig |
| KCNQ3 | R230H | ECD | O95905 | -0.45668079 | 0.2080645 | -2.1949 | 10 | 0.0528956 | 0 | 3 | 3 | not sig |
| MYT1L | C504R | SUDS3 | Q9H7L9 | -0.45676553 | 0.3884724 | -1.175799 | 6 | 0.2842122 | 0 | 3 | 3 | not sig |
| KCNMA1 | A1033P | PRKDC | P78527 | -0.45772433 | 0.159534 | -2.869133 | 8 | 0.0208567 | 0 | 3 | 3 | not sig |
| KCNQ3 | R230L | PRKDC | P78527 | -0.45837904 | 0.1166369 | -3.929967 | 10 | 0.0028204 | 0 | 3 | 3 | not sig |
| KCNQ3 | R230H | IPO4 | Q8TEX9 | -0.45860073 | 0.2797404 | -1.63938 | 10 | 0.1321714 | 0 | 3 | 3 | not sig |
| SLC6A1 | L54F | SLC35B2 | Q8TB61 | -0.45879699 | 0.2255566 | -2.034066 | 14 | 0.0613469 | 0 | 3 | 3 | not sig |
| CHD8 | L834P | KPNA2 | P52292 | -0.46020141 | 0.1379634 | -3.335678 | 12 | 0.0059354 | 0 | 3 | 3 | not sig |
| CHD8 | L834P | MYH10 | P35580 | -0.46225984 | 0.3615007 | -1.278725 | 12 | 0.2251708 | 0 | 3 | 3 | not sig |
| TBR1 | K228E | BCKDK | O14874 | -0.46501669 | 1.5246204 | -0.305005 | 3 | 0.7802988 | 0 | 2 | 1 | not sig |
| STXBP1 | A251T | PLOD3 | O60568 | -0.46681836 | 0.8288402 | -0.563219 | 6 | 0.5936927 | 0 | 3 | 3 | not sig |

| | | | | | | | | | | | | |
|---|---|---|---|---|---|---|---|---|---|---|---|---|
| FOXP1 | R513H | NUP155 | O75694 | -0.47370328 | 0.2814166 | -1.683281 | 8 | 0.1308186 | 0 | 3 | 3 | not sig |
| KCNQ3 | R230L | IPO4 | Q8TEX9 | -0.4844514 | 0.2797404 | -1.731789 | 10 | 0.1139855 | 0 | 3 | 3 | not sig |
| TBR1 | K228E | PIP4K2C | Q8TBX8 | -0.49101462 | 0.1454686 | -3.3754 | 6 | 0.0149424 | 0 | 3 | 3 | not sig |
| PPP1R9B | R753H | ADSL | P30566 | -0.4952339 | 0.1064796 | -4.650974 | 4 | 0.0096544 | 0 | 3 | 3 | not sig |
| RORB | Y22D | MED1 | Q15648 | -0.49714196 | 0.2005035 | -2.479468 | 4 | 0.06825 | 0 | 3 | 3 | not sig |
| TBR1 | N374H | ACAD11 | Q709F0 | -0.4995051 | 0.2181514 | -2.289718 | 2 | 0.1491981 | 0 | 2 | 2 | not sig |
| PPP1R9B | R753H | LAP3 | P28838 | -0.49984008 | 0.0067741 | -73.78681 | 1 | 0.0086273 | 0 | 2 | 1 | not sig |
| MYT1L | C504R | SIN3A | Q96ST3 | -0.50191148 | 0.4172319 | -1.202956 | 6 | 0.2743033 | 0 | 3 | 3 | not sig |
| TBR1 | K228E | PKP4 | Q99569 | -0.50218359 | 0.6964937 | -0.721017 | 4 | 0.5107896 | 0 | 2 | 2 | not sig |
| SLC6A1 | G307R | SLC35B2 | Q8TB61 | -0.50697822 | 0.2255566 | -2.247676 | 14 | 0.0412342 | 0 | 3 | 3 | not sig |
| CHD8 | C1095Y | GDI2 | P50395 | -0.516447 | 0.4272981 | -1.208634 | 10 | 0.2546062 | 0 | 3 | 3 | not sig |
| KCNQ3 | R230L | YME1L1 | Q96TA2 | -0.51645885 | 0.1636193 | -3.156466 | 10 | 0.01022 | 0 | 3 | 3 | not sig |
| CHD8 | C1095Y | ZBTB11 | O95625 | -0.51758964 | 0.3572072 | -1.44899 | 8 | 0.1853779 | 0 | 2 | 2 | not sig |
| TBR1 | K228E | WDR11 | Q9BZH6 | -0.51867824 | 0.6649928 | -0.779976 | 5 | 0.4706966 | 0 | 3 | 2 | not sig |
| STXBP1 | A251T | DDX23 | Q9BUQ8 | -0.52020769 | 1.4721821 | -0.353358 | 2 | 0.7575904 | 0 | 1 | 3 | not sig |
| KCNQ3 | R236C | CLPTM1 | O96005 | -0.52548372 | 0.1968038 | -2.670089 | 6 | 0.0370222 | 0 | 3 | 3 | not sig |
| KCNQ3 | R230L | NAP1L4 | Q99733 | -0.53665501 | 0.2408759 | -2.227931 | 9 | 0.0528751 | 0 | 2 | 3 | not sig |
| RFX3 | A508E | VAPA | Q9P0L0 | -0.53963837 | 0.6220594 | -0.867503 | 4 | 0.4346103 | 0 | 3 | 3 | not sig |
| RFX3 | A508E | UBAP2L | Q14157 | -0.54354813 | 0.3187843 | -1.705065 | 4 | 0.1633824 | 0 | 3 | 3 | not sig |
| PPP1R9B | R753H | CEP131 | Q9UPN4 | -0.54506786 | 0.4794915 | -1.136762 | 3 | 0.3382223 | 0 | 3 | 2 | not sig |
| STXBP1 | R551C | PLOD3 | O60568 | -0.54518603 | 0.8288402 | -0.65777 | 6 | 0.5350919 | 0 | 3 | 3 | not sig |
| KCNMA1 | A1033P | JPH1 | Q9HDC5 | -0.54728511 | 0.176691 | -3.097414 | 8 | 0.0147233 | 0 | 3 | 3 | not sig |
| KCNQ3 | R230C | TNPO1 | Q92973 | -0.54744984 | 0.2814741 | -1.944939 | 10 | 0.0804202 | 0 | 3 | 3 | not sig |
| TBR1 | K228E | OCLN | Q16625 | -0.55311854 | 0.2380271 | -2.323763 | 6 | 0.0591435 | 0 | 3 | 3 | not sig |
| MYT1L | C504R | BRMS1L | Q5PSV4 | -0.55717665 | 0.2974702 | -1.873051 | 6 | 0.1102117 | 0 | 3 | 3 | not sig |
| MKX | R93G | PRKDC | P78527 | -0.56592838 | 0.617964 | -0.915795 | 5 | 0.4017842 | 0 | 3 | 2 | not sig |
| KCNQ3 | R236C | NAA15 | Q9BXJ9 | -0.56668658 | 0.2857512 | -1.983147 | 9 | 0.07866 | 0 | 2 | 3 | not sig |
| AP2S1 | G64D | RDH13 | Q8NBN7 | -0.57117309 | 0.440108 | -1.297802 | 5 | 0.2509981 | 0 | 3 | 2 | not sig |
| TBR1 | K228E | KIF7 | Q2M1P5 | -0.57168197 | 0.6496592 | -0.879972 | 6 | 0.4127283 | 0 | 3 | 3 | not sig |
| KCNQ3 | R236C | HEATR3 | Q7Z4Q2 | -0.58111929 | 0.2883598 | -2.015257 | 8 | 0.0786347 | 0 | 3 | 3 | not sig |
| MYT1L | C504R | SAP30 | O75446 | -0.58406708 | 0.3767999 | -1.550072 | 6 | 0.1721012 | 0 | 3 | 3 | not sig |
| KCNQ3 | R230C | FANCI | Q9NVI1 | -0.58423875 | 0.29473 | -1.982285 | 7 | 0.0878853 | 0 | 3 | 3 | not sig |
| SLC6A1 | G307R | MOGS | Q13724 | -0.60465885 | 0.2829625 | -2.136887 | 14 | 0.0507402 | 0 | 3 | 3 | not sig |
| KCNQ3 | R230H | IPO7 | O95373 | -0.61234575 | 0.2169668 | -2.822302 | 10 | 0.0180891 | 0 | 3 | 3 | not sig |
| KCNQ3 | R236C | PPM1G | O15355 | -0.61401516 | 0.2028349 | -3.027167 | 10 | 0.0127386 | 0 | 3 | 3 | not sig |
| FOXP1 | R513H | FOXP2 | O15409 | -0.62602588 | 0.1172029 | -5.341383 | 8 | 0.000693 | 0 | 3 | 3 | not sig |
| CHD8 | Q696K | MYH10 | P35580 | -0.62810279 | 0.3615007 | -1.737487 | 12 | 0.1078723 | 0 | 3 | 3 | not sig |
| MYT1L | H522Q | RORB | Q92753 | -0.63788882 | 0.6094434 | -1.046674 | 6 | 0.3355714 | 0 | 3 | 3 | not sig |
| MKX | L89F | XRCC1 | P18887 | -0.64236327 | 1.3438066 | -0.478018 | 2 | 0.6797881 | 0 | 1 | 2 | not sig |
| FOXP1 | R513C | FOXP2 | O15409 | -0.64747126 | 0.1172029 | -5.52436 | 8 | 0.0005575 | 0 | 3 | 3 | not sig |
| VEZF1 | Q209P | YTHDC2 | Q9H6S0 | -0.65251134 | 0.3440904 | -1.896337 | 4 | 0.1307904 | 0 | 3 | 3 | not sig |
| STXBP1 | A251T | EXOC1 | Q9NV70 | -0.65471145 | 0.378191 | -1.731166 | 3 | 0.1818528 | 0 | 1 | 2 | not sig |
| RORB | Y22D | TP53 | P04637 | -0.65489605 | 0.3088046 | -2.120746 | 4 | 0.101257 | 0 | 3 | 3 | not sig |
| FOXP1 | L327P | FOXP2 | O15409 | -0.65518778 | 0.1172029 | -5.590199 | 8 | 0.0005161 | 0 | 3 | 3 | not sig |
| KCNQ3 | R230L | IPO7 | O95373 | -0.65676732 | 0.2169668 | -3.027041 | 10 | 0.0127413 | 0 | 3 | 3 | not sig |
| TBR1 | K228E | DARS2 | Q6PI48 | -0.66286698 | 0 | 0 | 0 | 0 | 0 | 1 | 1 | not sig |
| MKX | L89F | LEMD2 | Q8NC56 | -0.66420585 | 0.5157581 | -1.287824 | 5 | 0.2541872 | 0 | 3 | 2 | not sig |
| CHD8 | C1095Y | EPS15L1 | Q9UBC2 | -0.6794902 | 1.3903328 | -0.488725 | 8 | 0.6381519 | 0 | 3 | 1 | not sig |
| IRFBPL | F30L | VGLL4 | Q14135 | -0.68435709 | 0.2768399 | -2.472032 | 3 | 0.0899023 | 0 | 2 | 3 | not sig |
| FOXP1 | R513C | NUP155 | O75694 | -0.68657043 | 0.2814166 | -2.439694 | 8 | 0.0405838 | 0 | 3 | 3 | not sig |
| FOXP2 | R570C | PRKDC | P78527 | -0.68727185 | 0.0971751 | -7.07251 | 4 | 0.002109 | 0 | 3 | 3 | not sig |
| CHD8 | Q696K | EPS15L1 | Q9UBC2 | -0.68882758 | 1.3903328 | -0.495441 | 8 | 0.63361 | 0 | 3 | 1 | not sig |
| DNMT3A | L508P | KPNA3 | O00505 | -0.69024538 | 0.3554095 | -1.942113 | 4 | 0.1240758 | 0 | 3 | 3 | not sig |
| CHD8 | M904I | ZBTB11 | O95625 | -0.69108133 | 0.3260841 | -2.119335 | 8 | 0.0668944 | 0 | 3 | 2 | not sig |
| KCNQ3 | R230C | PRKDC | P78527 | -0.6976776 | 0.1166369 | -5.981622 | 10 | 0.0001354 | 0 | 3 | 3 | not sig |
| STXBP1 | R551C | EXOC1 | Q9NV70 | -0.70765419 | 0.2818869 | -2.510419 | 3 | 0.0869056 | 0 | 3 | 2 | not sig |
| MYT1L | C504R | ACAD11 | Q709F0 | -0.71505636 | 0.6172821 | -1.158395 | 6 | 0.2907226 | 0 | 3 | 3 | not sig |
| CHD8 | Q696K | OAT | P04181 | -0.71957574 | 0.3848641 | -1.869688 | 11 | 0.0883605 | 0 | 3 | 3 | not sig |
| KCNQ3 | R230C | FANCD2 | Q9BXW9 | -0.72599868 | 0.1489599 | -4.873786 | 8 | 0.001234 | 0 | 3 | 3 | not sig |
| MYT1L | H522Q | HIRA | P54198 | -0.73227867 | 0.1886339 | -3.88201 | 6 | 0.0081517 | 0 | 3 | 3 | not sig |
| TBR1 | N374H | RBM15 | Q96T37 | -0.73350773 | 0.9227374 | -0.794926 | 6 | 0.4569404 | 0 | 3 | 3 | not sig |
| FOXP1 | R513H | CTBP2 | P56545 | -0.73484275 | 0.8248123 | -0.890921 | 8 | 0.3989725 | 0 | 3 | 3 | not sig |
| KCNQ3 | R236C | IPO4 | Q8TEX9 | -0.73659242 | 0.2797404 | -2.633129 | 10 | 0.0250274 | 0 | 3 | 3 | not sig |
| RFX3 | A508E | KPNA1 | P52294 | -0.73775839 | 0.4059972 | -1.817151 | 4 | 0.1433508 | 0 | 3 | 3 | not sig |
| RFX3 | A508E | KPNA6 | O60684 | -0.74116668 | 0.1209368 | -6.128543 | 4 | 0.0035918 | 0 | 3 | 3 | not sig |
| MYT1L | H522Q | MED23 | Q9ULK4 | -0.74607376 | 0.2391815 | -3.119279 | 6 | 0.0206038 | 0 | 3 | 3 | not sig |
| TBR1 | N374H | BTRC | Q9Y297 | -0.74718608 | 0.9106616 | -0.820487 | 4 | 0.4580186 | 0 | 3 | 2 | not sig |
| TBR1 | K228E | ERCC3 | P19447 | -0.7482932 | 0.4193399 | -1.784455 | 4 | 0.1489093 | 0 | 2 | 2 | not sig |
| KCNQ3 | R236C | IPO7 | O95373 | -0.75021718 | 0.2169668 | -3.457752 | 10 | 0.0061459 | 0 | 3 | 3 | not sig |
| MYT1L | H522Q | XRCC1 | P18887 | -0.75239439 | 0.4084842 | -1.841918 | 6 | 0.1150718 | 0 | 3 | 3 | not sig |
| SLC6A1 | G307R | LMBR1 | Q8WVP7 | -0.76083378 | 0.572754 | -1.328378 | 11 | 0.2109544 | 0 | 2 | 2 | not sig |
| MYT1L | C504R | MED23 | Q9ULK4 | -0.7726291 | 0.2391815 | -3.230305 | 6 | 0.0179036 | 0 | 3 | 3 | not sig |

| | | | | | | | | | | | | |
|---|---|---|---|---|---|---|---|---|---|---|---|---|
| TBR1 | N374H | AP2B1 | P63010 | -0.77481362 | 0.5409652 | -1.43228 | 4 | 0.2253346 | 0 | 2 | 3 | not sig |
| MKX | L89F | ERCC3 | P19447 | -0.77715342 | 0.4060955 | -1.913721 | 5 | 0.1138416 | 0 | 3 | 2 | not sig |
| MYT1L | H522Q | MED15 | Q96RN5 | -0.78616564 | 0.1728793 | -4.547484 | 6 | 0.0039022 | 0 | 3 | 3 | not sig |
| RORB | Y22D | SHKBP1 | Q8TBC3 | -0.79891266 | 0.5283311 | -1.512144 | 3 | 0.2276891 | 0 | 2 | 3 | not sig |
| KCNQ3 | R230C | HEATR3 | Q7Z4Q2 | -0.79971174 | 0.3223961 | -2.480525 | 8 | 0.0380805 | 0 | 2 | 3 | not sig |
| TBR1 | K228E | NXN | Q6DKJ4 | -0.80237815 | 0.8523093 | -0.941417 | 4 | 0.3997944 | 0 | 3 | 3 | not sig |
| KCNQ3 | R230C | ECD | O95905 | -0.80580148 | 0.2080645 | -3.872845 | 10 | 0.0030948 | 0 | 3 | 3 | not sig |
| DNMT3A | L508P | MKI67 | P46013 | -0.81058149 | 0.2544663 | -3.185417 | 2 | 0.0860259 | 0 | 1 | 3 | not sig |
| MKX | R93G | ERCC3 | P19447 | -0.81628009 | 0.4060955 | -2.010069 | 5 | 0.1006374 | 0 | 3 | 2 | not sig |
| MYT1L | C504R | KLHL7 | Q8IXQ5 | -0.82302833 | 0.2160346 | -3.809707 | 6 | 0.0088666 | 0 | 3 | 3 | not sig |
| MKX | L89F | AMPD2 | Q01433 | -0.82636122 | 0.1970635 | -4.193375 | 5 | 0.0085439 | 0 | 3 | 2 | not sig |
| RFX3 | A508E | ANP32E | Q9BTT0 | -0.83855995 | 0.5265776 | -1.592472 | 4 | 0.1864959 | 0 | 3 | 3 | not sig |
| MYT1L | C504R | MED10 | Q9BTT4 | -0.85100993 | 0.1843173 | -4.617093 | 6 | 0.0036264 | 0 | 3 | 3 | not sig |
| MYT1L | H522Q | MED17 | Q9NVC6 | -0.8519542 | 0.2635543 | -3.232557 | 6 | 0.017853 | 0 | 3 | 3 | not sig |
| MYT1L | C504R | MED20 | Q9H944 | -0.8584215 | 0.1564413 | -5.48718 | 6 | 0.0015331 | 0 | 3 | 3 | not sig |
| MYT1L | C504R | MED14 | O60244 | -0.86340221 | 0.1661007 | -5.198064 | 6 | 0.0020186 | 0 | 3 | 3 | not sig |
| RFX3 | A508E | ANP32B | Q92688 | -0.88394088 | 0.6070549 | -1.456114 | 4 | 0.2190737 | 0 | 3 | 3 | not sig |
| TBR1 | N374H | JPH1 | Q9HDC5 | -0.88528913 | 0.0773246 | -11.449 | 5 | 8.90E-05 | 0 | 3 | 2 | not sig |
| MYT1L | H522Q | MED14 | O60244 | -0.89076697 | 0.1661007 | -5.362811 | 6 | 0.0017235 | 0 | 3 | 3 | not sig |
| RORB | Y22D | CTSA | P10619 | -0.89565766 | 0.1152492 | -7.771484 | 4 | 0.0014779 | 0 | 3 | 3 | not sig |
| MYT1L | H522Q | LIG3 | P49916 | -0.90625854 | 0.4707499 | -1.925138 | 6 | 0.1025331 | 0 | 3 | 3 | not sig |
| MKX | R93G | LMNA | P02545 | -0.90776085 | 0.3674232 | -2.470614 | 5 | 0.0564876 | 0 | 3 | 2 | not sig |
| AP2S1 | G64D | SLC25A13 | Q9UJS0 | -0.91514018 | 0.4081878 | -2.241959 | 5 | 0.07503 | 0 | 3 | 2 | not sig |
| MYT1L | C504R | MED28 | Q9H204 | -0.91624862 | 0.288506 | -3.175839 | 6 | 0.0191765 | 0 | 3 | 3 | not sig |
| TBR1 | K228E | RBM15 | Q96T37 | -0.91740228 | 0.9227374 | -0.994218 | 6 | 0.3585055 | 0 | 3 | 3 | not sig |
| CHD8 | L834P | OAT | P04181 | -0.92122538 | 0.3848641 | -2.393638 | 11 | 0.0356314 | 0 | 3 | 3 | not sig |
| STXBP1 | A251T | DOCK7 | Q96N67 | -0.92681853 | 0.5709571 | -1.623272 | 4 | 0.1798531 | 0 | 2 | 3 | not sig |
| MYT1L | H522Q | MED10 | Q9BTT4 | -0.9292804 | 0.1843173 | -5.041744 | 6 | 0.0023528 | 0 | 3 | 3 | not sig |
| MKX | L89F | LIG3 | P49916 | -0.93601341 | 1.0844769 | -0.863101 | 3 | 0.4515624 | 0 | 1 | 2 | not sig |
| MYT1L | C504R | MED24 | O75448 | -0.94300854 | 0.2267059 | -4.159612 | 6 | 0.005947 | 0 | 3 | 3 | not sig |
| KCNQ3 | R236C | PEX19 | P40855 | -0.94511374 | 0.2184886 | -4.32569 | 10 | 0.0014997 | 0 | 3 | 3 | not sig |
| TBR1 | K228E | AP2B1 | P63010 | -0.94893301 | 0.5409652 | -1.754148 | 4 | 0.1542671 | 0 | 2 | 3 | not sig |
| MYT1L | H522Q | MED24 | O75448 | -0.95070356 | 0.2267059 | -4.193555 | 6 | 0.0057266 | 0 | 3 | 3 | not sig |
| MYT1L | H522Q | PRKDC | P78527 | -0.95763293 | 0.3387457 | -2.826997 | 6 | 0.0300762 | 0 | 3 | 3 | not sig |
| TBR1 | K228E | AP2A2 | O94973 | -0.95991921 | 0.5754265 | -1.668187 | 3 | 0.1938703 | 0 | 2 | 3 | not sig |
| MKX | L89F | BEND3 | Q5T5X7 | -0.9693517 | 1.3848244 | -0.699982 | 1 | 0.6112076 | 0 | 1 | 2 | not sig |
| MYT1L | C504R | MED22 | Q15528 | -0.97345357 | 0.2280643 | -4.268329 | 6 | 0.0052727 | 0 | 3 | 3 | not sig |
| MYT1L | H522Q | BEND3 | Q5T5X7 | -0.97425489 | 0.2236642 | -4.355882 | 5 | 0.0073186 | 0 | 3 | 2 | not sig |
| MYT1L | C504R | MED17 | Q9NVC6 | -0.97970285 | 0.2635543 | -3.717271 | 6 | 0.0098842 | 0 | 3 | 3 | not sig |
| KCNQ3 | R230H | SAAL1 | Q96ER3 | -0.98114285 | 0.3112131 | -3.15264 | 10 | 0.0102867 | 0 | 3 | 3 | not sig |
| TLK2 | D529G | KLHL9 | Q9P2J3 | -0.98183656 | 0.2946901 | -3.33176 | 4 | 0.0290579 | 0 | 3 | 3 | not sig |
| MYT1L | C504R | MED6 | O75586 | -0.9887725 | 0.2080578 | -4.752392 | 6 | 0.003151 | 0 | 3 | 3 | not sig |
| MYT1L | C504R | MED1 | Q15648 | -0.98926011 | 0.1271158 | -7.782356 | 6 | 0.000237 | 0 | 3 | 3 | not sig |
| MYT1L | H522Q | MED19 | A0JLT2 | -0.9900535 | 0.3287322 | -3.011732 | 6 | 0.0236477 | 0 | 3 | 3 | not sig |
| MYT1L | C504R | MED26 | O95402 | -0.99334336 | 0.176248 | -5.636054 | 6 | 0.0013359 | 0 | 3 | 3 | not sig |
| MYT1L | H522Q | MED22 | Q15528 | -0.99371178 | 0.2280643 | -4.357156 | 6 | 0.0047852 | 0 | 3 | 3 | not sig |
| MYT1L | C504R | MED15 | Q96RN5 | -0.99916762 | 0.1728793 | -5.779569 | 6 | 0.0011728 | 0 | 3 | 3 | not sig |
| TLK2 | D529G | KLHL13 | Q9P2N7 | -1.00405902 | 0.244052 | -4.114119 | 4 | 0.0146827 | 0 | 3 | 3 | down |
| FOXP1 | L327P | PRKDC | P78527 | -1.02044301 | 0.2547319 | -4.005949 | 8 | 0.0039171 | 0 | 3 | 3 | down |
| MYT1L | C504R | MED29 | Q9NX70 | -1.02665826 | 0.1564319 | -6.562971 | 6 | 0.0005995 | 0 | 3 | 3 | down |
| STXBP1 | R551C | DOCK7 | Q96N67 | -1.02836395 | 0.5709571 | -1.801123 | 4 | 0.1460474 | 0 | 2 | 3 | not sig |
| FOXP1 | R513C | CTBP1 | Q13363 | -1.03672947 | 0.8129658 | -1.275244 | 8 | 0.2380075 | 0 | 3 | 3 | not sig |
| MYT1L | C504R | MED16 | Q9Y2X0 | -1.03927512 | 0.2271772 | -4.574734 | 6 | 0.0037915 | 0 | 3 | 3 | down |
| CHD8 | C1095Y | OAT | P04181 | -1.0456979 | 0.4302912 | -2.43021 | 11 | 0.0333973 | 0 | 2 | 3 | down |
| IRFBPL | F30L | DNM2 | P50570 | -1.04686158 | 0.5804656 | -1.803486 | 3 | 0.1690888 | 0 | 2 | 3 | not sig |
| FOXP1 | R513H | CTBP1 | Q13363 | -1.04953695 | 0.8129658 | -1.290998 | 8 | 0.2327587 | 0 | 3 | 3 | not sig |
| MYT1L | H522Q | KLHL7 | Q8IXQ5 | -1.05010472 | 0.2160346 | -4.860818 | 6 | 0.0028205 | 0 | 3 | 3 | down |
| KCNQ3 | R230C | IPO9 | Q96P70 | -1.0506604 | 0.1425257 | -7.371726 | 10 | 2.39E-05 | 0 | 3 | 3 | down |
| RORB | Y22D | MRE11 | P49959 | -1.05203129 | 0.4859655 | -2.164827 | 4 | 0.0963666 | 0 | 3 | 3 | not sig |
| MYT1L | H522Q | MED29 | Q9NX70 | -1.05293615 | 0.1564319 | -6.730954 | 6 | 0.0005234 | 0 | 3 | 3 | down |
| STXBP1 | A251T | PRPF6 | O94906 | -1.05314951 | 0.7379515 | -1.427126 | 3 | 0.2488233 | 0 | 1 | 3 | not sig |
| MYT1L | C504R | MED4 | Q9NPJ6 | -1.05611856 | 0.2013324 | -5.245647 | 6 | 0.0019279 | 0 | 3 | 3 | down |
| TLK2 | D529G | CUL3 | Q13618 | -1.06236479 | 0.218131 | -4.870307 | 4 | 0.0082179 | 0 | 3 | 3 | down |
| MYT1L | C504R | MED19 | A0JLT2 | -1.07123254 | 0.3287322 | -3.258678 | 6 | 0.0172774 | 0 | 3 | 3 | down |
| MKX | L89F | PRKDC | P78527 | -1.07341366 | 0.617964 | -1.737017 | 5 | 0.1428933 | 0 | 3 | 2 | not sig |
| MYT1L | H522Q | TOP2A | P11388 | -1.07457371 | 0.4420684 | -2.430786 | 6 | 0.0511078 | 0 | 3 | 3 | not sig |
| KCNQ3 | R230L | CLPTM1 | O96005 | -1.07489897 | 0.2783226 | -3.862061 | 6 | 0.0083423 | 0 | 1 | 3 | down |
| MYT1L | C504R | RRBP1 | Q9P2E9 | -1.07910639 | 0.4379766 | -2.463845 | 6 | 0.0488639 | 0 | 3 | 3 | down |
| MYT1L | C504R | TOP2A | P11388 | -1.08767275 | 0.4420684 | -2.460417 | 6 | 0.0490917 | 0 | 3 | 3 | down |
| MYT1L | H522Q | MED6 | O75586 | -1.08819936 | 0.2080578 | -5.230273 | 6 | 0.0019567 | 0 | 3 | 3 | down |
| VEZF1 | Q209P | ZC3H8 | Q8N5P1 | -1.08824853 | 0.4178899 | -2.604151 | 4 | 0.0597864 | 0 | 3 | 3 | not sig |
| MYT1L | H522Q | MED1 | Q15648 | -1.09274502 | 0.1271158 | -8.596455 | 6 | 0.0001362 | 0 | 3 | 3 | down |

| | | | | | | | | | | | | |
|---|---|---|---|---|---|---|---|---|---|---|---|---|
| MYT1L | H522Q | LMNA | P02545 | -1.0936485 | 0.5128562 | -2.132466 | 6 | 0.0769468 | 0 | 3 | 3 | not sig |
| MYT1L | C504R | MED9 | Q9NWA0 | -1.09497064 | 0.2448122 | -4.472697 | 6 | 0.0042252 | 0 | 3 | 3 | down |
| FOXP1 | R513C | CTBP2 | P56545 | -1.09537216 | 0.8248123 | -1.328026 | 8 | 0.2208076 | 0 | 3 | 3 | not sig |
| MYT1L | H522Q | MED20 | Q9H944 | -1.09559968 | 0.1564413 | -7.003264 | 6 | 0.0004224 | 0 | 3 | 3 | down |
| KCNQ3 | R230L | SAAL1 | Q96ER3 | -1.09859024 | 0.3112131 | -3.530026 | 10 | 0.0054466 | 0 | 3 | 3 | down |
| PPP2R5D | E198K | PPFIA1 | Q13136 | -1.10176691 | 0.3419908 | -3.221628 | 3 | 0.048524 | 0 | 3 | 2 | down |
| MYT1L | H522Q | MED26 | O95402 | -1.12307449 | 0.176248 | -6.372125 | 6 | 0.0007018 | 0 | 3 | 3 | down |
| IRFBPL | F30L | TFAP4 | Q01664 | -1.12482331 | 0.1128742 | -9.965279 | 4 | 0.0005696 | 0 | 3 | 3 | down |
| KCNQ3 | R230C | PPM1G | O15355 | -1.12603863 | 0.2028349 | -5.551503 | 10 | 0.0002436 | 0 | 3 | 3 | down |
| TBR1 | N374H | STK3 | Q13188 | -1.12797893 | 0.03089 | -36.51596 | 2 | 0.0007491 | 0 | 2 | 2 | down |
| MYT1L | C504R | LMNA | P02545 | -1.12853651 | 0.5128562 | -2.200493 | 6 | 0.0700546 | 0 | 3 | 3 | not sig |
| TBR1 | K228E | JPH1 | Q9HDC5 | -1.1303619 | 0.0773246 | -14.61841 | 5 | 2.71E-05 | 0 | 3 | 2 | down |
| MYT1L | C504R | HIRA | P54198 | -1.13528374 | 0.1886339 | -6.01845 | 6 | 0.0009491 | 0 | 3 | 3 | down |
| MYT1L | C504R | POLR1G | O15446 | -1.14623485 | 0.1761679 | -6.506492 | 6 | 0.0006279 | 0 | 3 | 3 | down |
| KCNQ3 | R236C | PIP4P2 | Q8N4L2 | -1.16125535 | 0.4704966 | -2.468148 | 3 | 0.0902125 | 0 | 1 | 1 | not sig |
| KCNQ3 | R230C | NAP1L4 | Q99733 | -1.17464573 | 0.215446 | -5.452159 | 9 | 0.0004045 | 0 | 3 | 3 | down |
| KCNQ3 | R230C | IPO4 | Q8TEX9 | -1.17786999 | 0.2797404 | -4.210583 | 10 | 0.0017983 | 0 | 3 | 3 | down |
| TLK2 | D529G | TLK1 | Q9UKI8 | -1.18856105 | 0.1682294 | -7.065119 | 4 | 0.0021173 | 0 | 3 | 3 | down |
| TBR1 | K228E | STK3 | Q13188 | -1.18954223 | 0.0378324 | -31.44242 | 2 | 0.00101 | 0 | 1 | 2 | down |
| MYT1L | H522Q | LEMD2 | Q8NC56 | -1.19232863 | 0.7585737 | -1.571803 | 6 | 0.1670549 | 0 | 3 | 3 | not sig |
| TBR1 | N374H | BCKDK | O14874 | -1.19924948 | 1.4374259 | -0.834303 | 3 | 0.4653281 | 0 | 3 | 1 | not sig |
| RORB | Y22D | RAD50 | Q92878 | -1.20290541 | 0.4933942 | -2.438021 | 4 | 0.0713606 | 0 | 3 | 3 | not sig |
| MYT1L | H522Q | SAMD1 | Q6SPF0 | -1.20729483 | 0.3942694 | -3.062106 | 6 | 0.0221651 | 0 | 3 | 3 | down |
| MKX | L89F | SAMD1 | Q6SPF0 | -1.2076603 | 1.3347331 | -0.904795 | 1 | 0.5317928 | 0 | 1 | 2 | not sig |
| RORB | Y22D | NRIP1 | P48552 | -1.21776762 | 0.2427559 | -5.016428 | 4 | 0.007404 | 0 | 3 | 3 | down |
| STXBP1 | A251T | PRPF8 | Q6P2Q9 | -1.22345074 | 0.5189705 | -2.357457 | 6 | 0.0564799 | 0 | 3 | 3 | not sig |
| KCNQ3 | R230H | PEX19 | P40855 | -1.23011687 | 0.2184886 | -5.63012 | 10 | 0.0002184 | 0 | 3 | 3 | down |
| DPYSL2 | R238H | DPYSL4 | O14531 | -1.24175784 | 1.502358 | -0.826539 | 5 | 0.4461341 | 0 | 2 | 3 | not sig |
| MYT1L | H522Q | MED4 | Q9NPJ6 | -1.24643017 | 0.2013324 | -6.190907 | 6 | 0.0008178 | 0 | 3 | 3 | down |
| MYT1L | H522Q | MED16 | Q9Y2X0 | -1.25062697 | 0.2271772 | -5.505073 | 6 | 0.0015077 | 0 | 3 | 3 | down |
| MYT1L | C504R | BEND3 | Q5T5X7 | -1.26753279 | 0.2236642 | -5.667124 | 5 | 0.0023797 | 0 | 3 | 2 | down |
| SLC6A1 | A288V | NPTN | Q9Y639 | -1.27703985 | 0.2023386 | -6.311401 | 6 | 0.0007384 | 0 | 3 | 3 | down |
| MYT1L | C504R | PRKDC | P78527 | -1.27728286 | 0.3387457 | -3.770625 | 6 | 0.0092819 | 0 | 3 | 3 | down |
| IRFBPL | F30L | XPNPEP3 | Q9NQH7 | -1.2805633 | 0.4414521 | -2.900798 | 2 | 0.1011327 | 0 | 2 | 2 | not sig |
| MYT1L | H522Q | MED28 | Q9H204 | -1.3078177 | 0.288506 | -4.53307 | 6 | 0.0039622 | 0 | 3 | 3 | down |
| MYT1L | H522Q | MED9 | Q9NWA0 | -1.31083547 | 0.2448122 | -5.354454 | 6 | 0.0017372 | 0 | 3 | 3 | down |
| FOXP2 | R570C | FOXP1 | Q9H334 | -1.31996534 | 0.4303465 | -3.067215 | 4 | 0.0373943 | 0 | 3 | 3 | down |
| KCNQ3 | R230C | SAAL1 | Q96ER3 | -1.32746538 | 0.3112131 | -4.265455 | 10 | 0.0016488 | 0 | 3 | 3 | down |
| MYT1L | C504R | POLR1E | Q9GZS1 | -1.34414392 | 0.1781697 | -7.544177 | 6 | 0.0002813 | 0 | 3 | 3 | down |
| MYT1L | C504R | MDC1 | Q14676 | -1.34432022 | 0.2238776 | -6.004711 | 6 | 0.0009606 | 0 | 3 | 3 | down |
| KCNQ3 | R230C | CLPTM1 | O96005 | -1.36008919 | 0.2783226 | -4.886736 | 6 | 0.0027474 | 0 | 1 | 3 | down |
| MYT1L | H522Q | UBN2 | Q6ZU65 | -1.36519302 | 0.2722348 | -5.014763 | 5 | 0.0040532 | 0 | 3 | 3 | down |
| CHD8 | R1580W | TAOK2 | Q9UL54 | -1.37419254 | 0.2835405 | -4.846548 | 11 | 0.0005136 | 0 | 2 | 3 | down |
| MYT1L | H522Q | RAE1 | P78406 | -1.37511098 | 0.6507933 | -2.112977 | 6 | 0.0790469 | 0 | 3 | 3 | not sig |
| RORB | Y22D | MIDEAS | Q6PJG2 | -1.37833366 | 0.2781079 | -4.956111 | 4 | 0.0077275 | 0 | 3 | 3 | down |
| MKX | R93G | XRCC1 | P18887 | -1.37953581 | 1.0972135 | -1.257308 | 2 | 0.3355683 | 0 | 2 | 2 | not sig |
| TBR1 | N374H | SAV1 | Q9H4B6 | -1.3922055 | 0.6524569 | -2.133789 | 4 | 0.0997819 | 0 | 2 | 2 | not sig |
| RORB | Y22D | DNTTIP1 | Q9H147 | -1.4001624 | 0.3337346 | -4.195436 | 4 | 0.0137462 | 0 | 3 | 3 | down |
| MYT1L | H522Q | RRBP1 | Q9P2E9 | -1.40036483 | 0.4379766 | -3.197351 | 6 | 0.0186623 | 0 | 3 | 3 | down |
| LRRC4C | I163N | TNPO1 | Q92973 | -1.40036954 | 0.2180415 | -6.422492 | 4 | 0.0030214 | 0 | 3 | 3 | down |
| MYT1L | C504R | LIG3 | P49916 | -1.40331393 | 0.4707499 | -2.981018 | 6 | 0.0246042 | 0 | 3 | 3 | down |
| MYT1L | H522Q | MDC1 | Q14676 | -1.41921167 | 0.2238776 | -6.339231 | 6 | 0.0007214 | 0 | 3 | 3 | down |
| KCNQ3 | R230C | IPO7 | O95373 | -1.43458494 | 0.2169668 | -6.612003 | 10 | 5.99E-05 | 0 | 3 | 3 | down |
| LRRC4C | I163N | TNPO3 | Q9Y5L0 | -1.4387926 | 0.4576923 | -3.143581 | 1 | 0.1960702 | 0 | 1 | 2 | not sig |
| TBR1 | K228E | POMGNT2 | Q8NAT1 | -1.44656529 | 0.2047812 | -7.063956 | 1 | 0.0895274 | 0 | 1 | 2 | not sig |
| MYT1L | C504R | SAMD1 | Q6SPF0 | -1.44946077 | 0.3942694 | -3.676321 | 6 | 0.010376 | 0 | 3 | 3 | down |
| TBR1 | N374H | AP2A1 | O95782 | -1.45013675 | 0.9354734 | -1.550164 | 2 | 0.2612409 | 0 | 2 | 2 | not sig |
| MKX | R93G | XRCC5 | P13010 | -1.46675958 | 0.9967617 | -1.471525 | 5 | 0.2011215 | 0 | 3 | 2 | not sig |
| MYT1L | H522Q | MYCBP2 | O75592 | -1.49470818 | 0.5672311 | -2.635096 | 6 | 0.0387941 | 0 | 3 | 3 | down |
| MYT1L | C504R | XRCC1 | P18887 | -1.49790343 | 0.4084842 | -3.666981 | 6 | 0.0104919 | 0 | 3 | 3 | down |
| MYT1L | C504R | MKI67 | P46013 | -1.52652454 | 0.1433092 | -10.65197 | 6 | 4.04E-05 | 0 | 3 | 3 | down |
| MYT1L | C504R | POLR1A | O95602 | -1.52961866 | 0.2122881 | -7.205391 | 6 | 0.0003618 | 0 | 3 | 3 | down |
| MKX | L89F | LMNA | P02545 | -1.53644045 | 0.3674232 | -4.181665 | 5 | 0.008641 | 0 | 3 | 2 | down |
| MYT1L | H522Q | FBXO45 | P0C2W1 | -1.57847496 | 0.5400383 | -2.922894 | 6 | 0.0265312 | 0 | 3 | 3 | down |
| MYT1L | C504R | UBN2 | Q6ZU65 | -1.59468935 | 0.3043678 | -5.23935 | 5 | 0.0033556 | 0 | 2 | 3 | down |
| MYT1L | H522Q | POLR1G | O15446 | -1.59757084 | 0.1761679 | -9.068457 | 6 | 0.0001009 | 0 | 3 | 3 | down |
| FOXP1 | L327P | FOXP4 | Q8IVH2 | -1.62163929 | 0.1243622 | -13.03964 | 8 | 1.14E-06 | 0 | 3 | 3 | down |
| KCNQ3 | R230L | PEX19 | P40855 | -1.62498354 | 0.2184886 | -7.437385 | 10 | 2.22E-05 | 0 | 3 | 3 | down |
| AP2S1 | G64D | GALK1 | P51570 | -1.63869178 | 0.7034245 | -2.329592 | 1 | 0.2581322 | 0 | 1 | 1 | not sig |
| RFX3 | A508E | FOXJ3 | Q9UPW0 | -1.64600737 | 0.4942256 | -3.330478 | 4 | 0.0290927 | 0 | 3 | 3 | down |
| MYT1L | H522Q | MKI67 | P46013 | -1.66145253 | 0.1433092 | -11.59348 | 6 | 2.48E-05 | 0 | 3 | 3 | down |
| ASH1L | F1944C | NUP155 | O75694 | -1.66814001 | 0.3238246 | -5.151368 | 4 | 0.0067381 | 0 | 3 | 3 | down |

| | | | | | | | | | | | | |
|---|---|---|---|---|---|---|---|---|---|---|---|---|
| AP2S1 | G64D | SRPRB | Q9Y5M8 | -1.68239374 | 0.4855003 | -3.465278 | 5 | 0.0179402 | 0 | 3 | 2 | down |
| FOXP1 | R513H | PRKDC | P78527 | -1.69638315 | 0.2547319 | -6.659485 | 8 | 0.0001592 | 0 | 3 | 3 | down |
| MYT1L | C504R | LEMD2 | Q8NC56 | -1.70800202 | 0.7585737 | -2.251597 | 6 | 0.0652978 | 0 | 3 | 3 | not sig |
| STXBP1 | A251T | SNRNP200 | O75643 | -1.72354904 | 0.6047479 | -2.850029 | 6 | 0.0291805 | 0 | 3 | 3 | down |
| MKX | R93G | LIG3 | P49916 | -1.72844059 | 0.8083214 | -2.138309 | 3 | 0.122048 | 0 | 3 | 2 | not sig |
| MYT1L | H522Q | SPRYD3 | Q8NCJ5 | -1.75334264 | 0.5414258 | -3.23838 | 6 | 0.0177229 | 0 | 3 | 3 | down |
| FOXP1 | R513C | PRKDC | P78527 | -1.75456303 | 0.2547319 | -6.887881 | 8 | 0.0001261 | 0 | 3 | 3 | down |
| STXBP1 | R551C | PRPF8 | Q6P2Q9 | -1.78054923 | 0.5189705 | -3.430926 | 6 | 0.0139553 | 0 | 3 | 3 | down |
| MYT1L | H522Q | POLR1A | O95602 | -1.7970483 | 0.2122881 | -8.46514 | 6 | 0.0001485 | 0 | 3 | 3 | down |
| MKX | L89F | HIRA | P54198 | -1.7995409 | 2.2771362 | -0.790265 | 1 | 0.5742435 | 0 | 1 | 2 | not sig |
| TBR1 | K228E | SAV1 | Q9H4B6 | -1.80460257 | 0.595609 | -3.029844 | 4 | 0.0387865 | 0 | 3 | 2 | down |
| MYT1L | H522Q | POLR1E | Q9GZS1 | -1.83632738 | 0.1781697 | -10.30662 | 6 | 4.87E-05 | 0 | 3 | 3 | down |
| FOXP1 | R513C | FOXP4 | Q8IVH2 | -1.88017619 | 0.1243622 | -15.11855 | 8 | 3.63E-07 | 0 | 3 | 3 | down |
| MYT1L | C504R | SPRYD3 | Q8NCJ5 | -1.93175936 | 0.5414258 | -3.567912 | 6 | 0.0118139 | 0 | 3 | 3 | down |
| TBR1 | N374H | ODF2 | Q5BJF6 | -1.96273525 | 0.9395418 | -2.089034 | 3 | 0.1278947 | 0 | 1 | 3 | not sig |
| FOXP2 | R570C | FOXP4 | Q8IVH2 | -1.96942498 | 0.0475019 | -41.45993 | 4 | 2.02E-06 | 0 | 3 | 3 | down |
| RORB | Y22D | PRKDC | P78527 | -2.0485985 | 0.3500143 | -5.852899 | 4 | 0.0042515 | 0 | 3 | 3 | down |
| MKX | L89F | XRCC5 | P13010 | -2.08294052 | 0.9967617 | -2.089708 | 5 | 0.0909336 | 0 | 3 | 2 | not sig |
| MYT1L | C504R | MYCBP2 | O75592 | -2.10226788 | 0.5672311 | -3.706193 | 6 | 0.0100146 | 0 | 3 | 3 | down |
| KCNQ3 | R230C | PEX19 | P40855 | -2.11230659 | 0.2184886 | -9.667813 | 10 | 2.16E-06 | 0 | 3 | 3 | down |
| CREBBP | R413Q | RELA | Q04206 | -2.11349032 | 0.2983769 | -7.083292 | 4 | 0.002097 | 0 | 3 | 3 | down |
| DPYSL2 | R238H | DPYSL5 | Q9BPU6 | -2.11539249 | 0.5910325 | -3.579148 | 7 | 0.0089864 | 0 | 3 | 3 | down |
| STXBP1 | R551C | SNRNP200 | O75643 | -2.13674288 | 0.6047479 | -3.533279 | 6 | 0.0123187 | 0 | 3 | 3 | down |
| MYT1L | C504R | RAE1 | P78406 | -2.14895103 | 0.6507933 | -3.302048 | 6 | 0.016366 | 0 | 3 | 3 | down |
| FOXP1 | R513H | FOXP4 | Q8IVH2 | -2.1680141 | 0.1243622 | -17.43306 | 8 | 1.20E-07 | 0 | 3 | 3 | down |
| VEZF1 | Q209P | ANKRD17 | O75179 | -2.20487555 | 0.2652138 | -8.313577 | 4 | 0.0011435 | 0 | 3 | 3 | down |
| STXBP1 | R551C | PRPF6 | O94906 | -2.26391074 | 0.5834019 | -3.880534 | 3 | 0.0303126 | 0 | 2 | 3 | down |
| AP2S1 | G64D | HUWE1 | Q7Z6Z7 | -2.29953933 | 2.2702833 | -1.012887 | 2 | 0.4177216 | 0 | 1 | 1 | not sig |
| MYT1L | C504R | FBXO45 | P0C2W1 | -2.31765713 | 0.5400383 | -4.291653 | 6 | 0.0051395 | 0 | 3 | 3 | down |
| TBR1 | K228E | ODF2 | Q5BJF6 | -2.35566468 | 0.742773 | -3.171446 | 3 | 0.0504245 | 0 | 2 | 3 | not sig |
| IRFBPL | F30L | CRTAP | O75718 | -2.37848039 | 0.0227964 | -104.3358 | 1 | 0.0061015 | 0 | 1 | 2 | down |
| PPP2R5D | E198K | PPP2CA | P67775 | -2.38871415 | 0.9691612 | -2.464723 | 3 | 0.0904871 | 0 | 3 | 2 | not sig |
| DPYSL2 | R496C | DPYSL4 | O14531 | -2.40775422 | 1.502358 | -1.60265 | 5 | 0.1699141 | 0 | 2 | 3 | not sig |
| CHD8 | L834P | PTCD1 | O75127 | -2.43624748 | 0.5408585 | -4.504408 | 10 | 0.0011353 | 0 | 1 | 3 | down |
| IRFBPL | F30L | RADX | Q6NSI4 | -2.46250626 | 0.1261431 | -19.52153 | 4 | 4.06E-05 | 0 | 3 | 3 | down |
| VEZF1 | Q209P | ANKHD1 | Q8IWZ3 | -2.53903719 | 0.313226 | -8.106087 | 4 | 0.0012592 | 0 | 3 | 3 | down |
| IRFBPL | F30L | SLTM | Q9NWH9 | -2.65298724 | 0.5739121 | -4.622637 | 1 | 0.135628 | 0 | 1 | 2 | not sig |
| PPP2R5D | E198K | PPP2R1A | P30153 | -2.81396419 | 0.1805988 | -15.5813 | 4 | 9.91E-05 | 0 | 3 | 3 | down |
| PPP2R5D | E198K | PPP2CB | P62714 | -3.48968386 | 0.4498822 | -7.756883 | 2 | 0.0162166 | 0 | 2 | 2 | down |
| RORB | Y22D | LEMD2 | Q8NC56 | -3.77595055 | 0.4618222 | -8.1762 | 3 | 0.0038275 | 0 | 2 | 3 | down |
| PPP2R5D | E198K | PPP2R1B | P30154 | -4.00287477 | 0.6987708 | -5.728452 | 4 | 0.0045978 | 0 | 3 | 3 | down |
| SLC6A1 | G360S | NPTN | Q9Y639 | -4.04899778 | 0.2861499 | -14.14992 | 6 | 7.78E-06 | 0 | 1 | 3 | down |
| RORB | Y22D | VRK3 | Q8IV63 | -4.09616175 | 0.549068 | -7.460208 | 3 | 0.0049867 | 0 | 2 | 3 | down |
| PPP2R5D | E198K | PPP4C | P60510 | -4.11765136 | 0.6422689 | -6.411102 | 2 | 0.0234762 | 0 | 1 | 3 | down |
| SLC6A1 | G307R | NPTN | Q9Y639 | -4.84913556 | 0.2861499 | -16.94613 | 6 | 2.70E-06 | 0 | 1 | 3 | down |
| AP2S1 | G64D | GPX8 | Q8TED1 | -8 | | | | | oneConditionMissi | 0 | 2 | down |
| AP2S1 | G64D | EIF3G | O75821 | -8 | | | | | oneConditionMissi | 0 | 1 | down |
| AP2S1 | G64D | RHOT2 | Q8IXI1 | -8 | | | | | oneConditionMissi | 0 | 1 | down |
| AP2S1 | G64D | CKAP4 | Q07065 | -8 | | | | | oneConditionMissi | 0 | 2 | down |
| AP2S1 | G64D | AFG3L2 | Q9Y4W6 | -8 | | | | | oneConditionMissi | 0 | 2 | down |
| AP2S1 | G64D | UMPS | P11172 | -8 | | | | | oneConditionMissi | 0 | 2 | down |
| CHD8 | R1580W | COPS3 | Q9UNS2 | -8 | | | | | oneConditionMissi | 0 | 3 | down |
| IRFBPL | F30L | P3H1 | Q32P28 | -8 | | | | | oneConditionMissi | 0 | 3 | down |
| IRFBPL | F30L | ELF1 | P32519 | -8 | | | | | oneConditionMissi | 0 | 3 | down |
| KCNQ3 | R230C | GBA | P04062 | -8 | | | | | oneConditionMissi | 0 | 3 | down |
| KCNQ3 | R230H | GBA | P04062 | -8 | | | | | oneConditionMissi | 0 | 3 | down |
| KCNQ3 | R230H | PIP4P2 | Q8N4L2 | -8 | | | | | oneConditionMissi | 0 | 1 | down |
| KCNQ3 | R230L | GBA | P04062 | -8 | | | | | oneConditionMissi | 0 | 3 | down |
| KCNQ3 | R236C | GBA | P04062 | -8 | | | | | oneConditionMissi | 0 | 3 | down |
| MKX | R93G | HIRA | P54198 | -8 | | | | | oneConditionMissi | 0 | 2 | down |
| MKX | R93G | SAMD1 | Q6SPF0 | -8 | | | | | oneConditionMissi | 0 | 2 | down |
| MKX | R93G | BEND3 | Q5T5X7 | -8 | | | | | oneConditionMissi | 0 | 2 | down |
| NSD1 | C1350R | PRPF6 | O94906 | -8 | | | | | oneConditionMissi | 0 | 1 | down |
| RFX3 | A508E | CRTC2 | Q53ET0 | -8 | | | | | oneConditionMissi | 0 | 3 | down |
| RFX3 | A508E | ZNF410 | Q86VK4 | -8 | | | | | oneConditionMissi | 0 | 3 | down |
| RORB | Y22D | LIG3 | P49916 | -8 | | | | | oneConditionMissi | 0 | 3 | down |
| SLC6A1 | G299V | NPTN | Q9Y639 | -8 | | | | | oneConditionMissi | 0 | 3 | down |
| SLC6A1 | H198P | NPTN | Q9Y639 | -8 | | | | | oneConditionMissi | 0 | 3 | down |
| STXBP1 | A251T | HINT1 | P49773 | -8 | | | | | oneConditionMissi | 0 | 3 | down |
| STXBP1 | R551C | DDX23 | Q9BUQ8 | -8 | | | | | oneConditionMissi | 0 | 3 | down |
| STXBP1 | R551C | HINT1 | P49773 | -8 | | | | | oneConditionMissi | 0 | 3 | down |
| TBR1 | K228E | ACAD11 | Q709F0 | -8 | | | | | oneConditionMissi | 0 | 2 | down |

| | | | | | | | | |
|---|---|---|---|---|---|---|---|---|
| TBR1 | K228E | AP1B1 | Q10567 | -8 | oneConditionMissi | 0 | 3 | down |
| TBR1 | K228E | AP2A1 | O95782 | -8 | oneConditionMissi | 0 | 2 | down |
| TBR1 | N374H | POMGNT2 | Q8NAT1 | -8 | oneConditionMissi | 0 | 2 | down |
| TBR1 | N374H | HSD17B4 | P51659 | -8 | oneConditionMissi | 0 | 1 | down |
| TRAF7 | R655Q | CASK | O14936 | -8 | oneConditionMissi | 0 | 1 | down |
